# Rapid isothermal amplification of the diatom RuBisCO gene from eDNA and eRNA reveals their abundance and photosynthetic physiology

**DOI:** 10.64898/2026.08.30.748096

**Authors:** Frédéric G. Verret, Katherine Hartle-Mougiou, Christos Chantzaras, Alexandra Peltekis, Francesca Margiotta, Diana Sarno, Ulisse Cardini, Marco Alba, Valeria Pizziol, Ioannis Markopoulos, Isavella Papadopoulou, Roberto Gallia, Isabella Percopo, Ferdinando Tramontano, Maira Maselli, Antonio Novellino, Stella Psarra, Marina Montresor, Matthew C. Mowlem, Electra Gizeli, Martha Valiadi

**Affiliations:** Foundation for Research and Technology Hellas, Institute of Molecular Biology and Biotechnology, Nikolaou Plastira 100, 70013, Heraklion, Crete, Greece; Department of Biology, University of Crete, Voutes Campus, 70013, Heraklion, Crete, Greece; Hellenic Centre for Marine Research, Institute of Oceanography, P.O.Box 2214, 71601, Heraklion, Crete, Greece; Department of Research Infrastructures for Marine Biological esources (RIMAR), Stazione Zoologica Anton Dohrn, Naples, Italy; Department of Integrative Marine Ecology (EMI), Stazione Zoologica Anton Dohrn, Genoa Marine Center, Genoa, Italy; ETT|DEDAGroup, Via Enrico Albareto 21, 16153, Genova, Italy; Department of Integrative Marine Ecology (EMI), Stazione Zoologica Anton Dohrn, Naples, Italy; Ocean Technology and Engineering Group, National Oceanography Centre, European Way, SO14 3ZH, Southampton, United Kingdom

**Keywords:** Diatoms, Rubisco, Chloroplast, Carbon fixation, ocean biomolecular observing, photosynthesis, phytoplankton, recombinase polymerase amplification

## Abstract

Diatoms are major contributors to marine primary production, yet current approaches for monitoring their abundance and function rely on coarse satellite chlorophyll estimates or sparse cell counts and bulk carbon fixation measurements. Molecular markers are a promising approach for high-resolution measurement of both abundance and metabolic activity through analysis of environmental DNA (eDNA) and RNA (eRNA). We present an isothermal quantitative recombinase polymerase amplification (qRPA) assay targeting *rbc*L gene copies and transcripts of marine diatoms, operating at low temperature and producing results in less than 15 min. We demonstrate specificity and calibration across diverse diatom taxa, then apply the assay to eDNA and eRNA samples from the Long-Term Ecological Research Mare Chiara site in the Gulf of Naples, Italy, alongside microscopy, chlorophyll, physicochemical, and carbon-fixation data. Diatom *rbc*L DNA tracked abundance across five orders of magnitude despite seasonal shifts in community composition. Combining molecular and optical data revealed increased cellular *rbc*L copies and chlorophyll in low-light winter populations, suggesting enhanced photosynthetic capacity despite lower abundance. Furthermore, *rbc*L RNA reflected total carbon fixation rates and identified populations with differing carbon fixation activity. These results support rapid, RPA-based *rbc*L quantification as a robust approach for biomolecular ocean observing.

**Graphical abstract:** 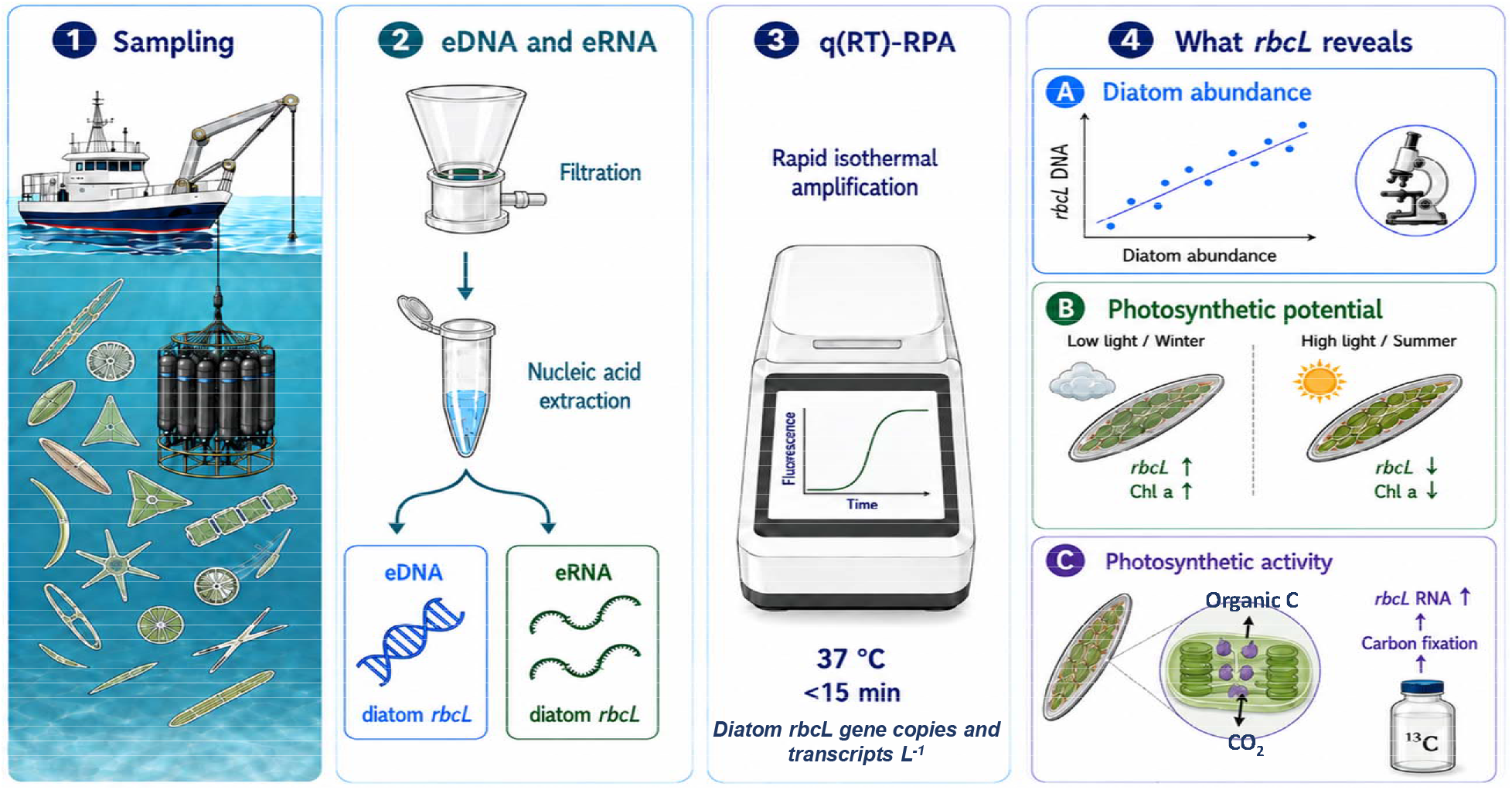

## INTRODUCTION

Marine primary productivity accounts for approximately half of global primary production, playing a central role in the Earth’s carbon cycle and climate regulation ^1–3^. Approximately half of this is contributed by diatoms, a taxonomically diverse, abundant and widespread eukaryotic phytoplankton taxon with a major contribution to oceanic carbon fixation and export ^4–7^. Given their ecological importance, monitoring diatom abundance, biomass and photosynthetic carbon fixation in the field is essential for understanding marine carbon cycling and its response to environmental change. However, the rapid temporal turnover of phytoplankton communities and the vast spatial scales of marine ecosystems present substantial observational challenges.

Traditional approaches to quantify phytoplankton abundance rely on microscopy-based cell identification, enumeration, and conversion to biovolume and carbon equivalents. However, the requirement for manual sampling and taxonomic expertise makes this approach difficult to scale across temporal and spatial gradients ^8^. High-performance liquid chromatography (HPLC) of phytoplankton pigment signatures offers an alternative but has similar practical limitations ^9^. Remote sensing provides a global perspective of phytoplankton biomass through pigment-based estimates, but ground-truthing against *in situ* measurements is still required ^10, 11^. Global and local primary productivity are also derived from integrating satellite-based bulk chlorophyll estimates ^12^ and need to be calibrated against technically demanding, and therefore sparse, carbon isotope assimilation measurements. All these methods provide bulk values for the phytoplankton community, with efforts focusing on satellite-based resolution of phytoplankton functional types ^13, 14^ and accounting for seasonal adaptations in photosynthetic physiology ^15^. Novel *in situ* measurement technologies aim to significantly improve data on phytoplankton dynamics, by achieving high-resolution monitoring capabilities and accurate datasets. Existing instruments include *in situ* fluorimetry to quantify bulk community photosynthetic activity (e.g. LabSTAF, microSTAF, Chelsea Technologies, UK) and *in situ* flow cytometric imaging for taxonomic characterization and abundance estimation (e.g. CytoBuoy, NL, and IFCB, McLane Research Labs, USA). Profiling abundance and function of key phytoplankton groups is also pursued through broader development and deployment of *in situ* biomolecular analyzers of environmental DNA and RNA (eDNA/eRNA), in line with the main goals of the Ocean Biomolecular Observing Network (OBON) and its European component EMO BON ^16–18^.

Molecular approaches targeting specific marker genes have emerged as powerful tools for inferring phytoplankton abundance and community composition. Taxon-specific quantification of the 18S ribosomal DNA (rDNA) gene is widely used due to its taxonomic conservation. However, highly variable gene copy numbers across species necessitate the application of species-specific correction factors, complicating quantitative interpretation ^19–21^. Gene proxies for biogeochemical rates are highly sought after, to obviate the need for specialized measurements using radioactive isotopes and to provide data at a much higher resolution, particularly when integrated on *in situ* analyzers. A particularly informative molecular target is the gene encoding the large subunit of ribulose-1,5-bisphosphate carboxylase/oxygenase (RuBisCO), *rbc*L. It is chloroplast-encoded, broadly conserved across phytoplankton, and directly involved in CO fixation. Cellular *rbc*L gene copies are known to vary across species and physiological conditions, due to a higher chloroplast content of larger or light-limited cells. Nevertheless, it can act as a dual proxy for both diatom abundance (via eDNA) ^21–25^ and cellular photosynthetic activity due to its transcriptional regulation (via eRNA) ^23, 26–31^. Both gene probe hybridization and qPCR approaches have established a positive relationship between phytoplankton *rbc*L transcripts L^-1^ and carbon fixation rates in the field ^26–28, 31–38^.

High-resolution and real time measurement of organism function requires molecular analysis to be performed autonomously using portable or underwater samplers and analyzers. Devices such as the Environmental Sample Processor have proven promising for *in situ* quantification of microbial metabolic function ^39–42^. However, wide-scale rollout of *in situ* biomolecular analysis instruments is hindered by the high energy demand of thermal cycling in quantitative PCR, resulting in expensive and bulky devices with limited operational time. Isothermal nucleic acid amplification technologies overcome these challenges in field-deployable systems by enabling rapid, robust, and quantitative detection of nucleic acid targets at constant temperature. In particular, recombinase polymerase amplification (RPA) operates at a low, stable temperature (37 – 42 °C), has high tolerance to impurities and a fast time-to-result in less than 15 min ^43–45^. RNA can be amplified in one step simply by adding reverse transcriptase to the RPA reaction ^46^. RPA has been applied for qualitative (yes/no) eDNA-based detection of marine organisms, for example, using lateral flow strips to detect harmful algae ^47^, or CRISPR-Cas to detect fish species ^48^. A one-pot reaction with additional reverse transcriptase amplifies RNA, showing high sensitivity for microbes and viruses ^46^. Inclusion of a fluorescently labelled exonuclease probe allows real-time detection for target quantification and is compatible with simple portable analyzers ^44^. This approach has achieved rapid quantification of waterborne pathogens down to approximately 100 transcript copies, similar to the general sensitivity of RT-qPCR, but in less than 20 min ^49^.

We set out to develop and validate a rapid, isothermal molecular assay to rapidly quantify marine diatom abundance and photosynthetic function (**Figure 1**). We designed a taxon-wide assay targeting the *rbc*L gene of diatoms and used quantitative RPA (qRPA) with optional addition of reverse transcriptase (qRT-RPA) to quantify *rbc*L gene copies and transcripts, respectively. The assay is specific to diatoms and can quantify the target gene/transcript over a wide range of field-relevant concentrations. We applied this approach to analyze seasonally and taxonomically diverse samples from the Mare Chiara Long-Term Ecological Research (LTER-MC) station in the Gulf of Naples (GoN), Italy. Using concurrent taxonomic and physicochemical datasets, we show that *rbc*L eDNA quantifies diatom abundance and identifies light-driven changes in photosynthetic physiology. Using *rbc*L eRNA, we demonstrate its potential as an indicator of diatom carbon fixation. The assay is fast and field-able, thus representing a significant advance towards real-time monitoring of diatom abundance and photosynthetic activity in natural marine environments.

**Figure 1.**
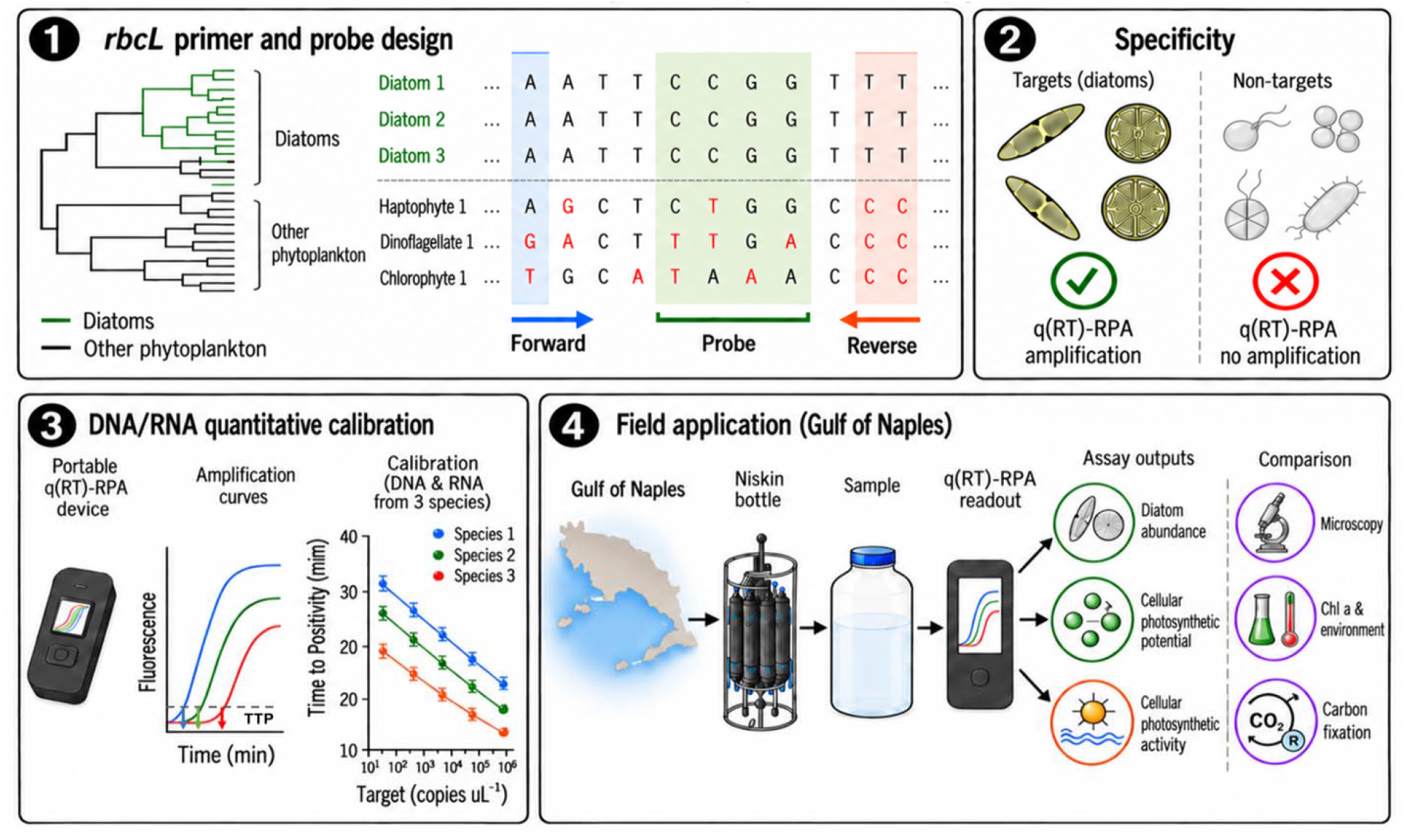
Diatom *rbc*L q(RT)-RPA assay development and validation. (1) Identification of suitable primer and probe target regions through *in silico* analysis of diverse diatom and non-diatom *rbc*L sequences. (2) Assay testing on DNA of diatom and non-diatom species representing closely and distantly related taxonomic groups. (3) Calibration curves for target quantification in unknown samples using three species, representing sequence diversity and different *rbc*L gene copy numbers per cell. Time to positivity (TTP) is the time taken for the fluorescence signal to rise above a threshold, and is proportional to the starting amount of target. (4) Application of the assay to field samples from the Gulf of Naples to predict diatom abundance, photosynthetic potential and photosynthetic activity. Validation against cell counts by microscopy and carbon fixation rates, enabling investigation of environmental controls on diatom abundance and physiology.

## MATERIALS AND METHODS

### Phytoplankton species and culturing

Most of the phytoplankton species grown in the laboratory were obtained from national culture collections, and additional diatom species were isolated from Heraklion, Crete, Greece (**Spreadsheet S1, Materials and Methods S1**). Monoclonal batch cultures were maintained at 18 °C in natural sea water based f/2 medium ^50^ and exposed to a 12:12 h light:dark photoperiod of 70 μmol photons m^-2^ s^-1^. Cell counts were carried out using a light microscope (Olympus CX43, Tokyo, Japan) and a haemacytometer for cells below 10 μm or a Sedgewick-Rafter counting chamber for cells above 10 μm in length. Cell biovolumes of *Thalassiosira rotula*, *Cyclotella cryptica* and *Hyalodiscus stelliger* were calculated based on measured cell dimensions and using the volume formula for a cylinder.

### Collection of field samples from Naples and Crete

Details of the sampling locations and sample types collected in Naples and Crete are provided in **Spreadsheet S2**. A total of 23 samples were collected from LTER-MC station ( https://deims.org/0b87459a-da3c-45af-a3e1-cb1508519411) situated in the GoN, Italy, over a period of 2 years. Samples were collected using a Niskin bottle mounted on a CTD rosette from 0.5 m depth. For nucleic acid extraction, 2 - 4 L of surface seawater were filtered through a cellulose ester filter (47 mm diameter, 1.2 μm pore-size, EMD Millipore, USA) and stored at −80 °C. For phytoplankton identification and counts, samples were immediately fixed with basic Lugol’s solution at a final concentration of 1%. Samples for nutrient analyses were collected from Niskin bottles into high-density polypropylene vials and frozen at −20 °C. For chlorophyll *a* (Chl *a*), a variable volume of seawater was filtered onto a Whatman glass fiber filter (GF/F, Ø 25 mm). The filters were immediately stored in liquid nitrogen until analysis.

Additional samples were collected along the northern coast of the island of Crete, Greece, namely Padanassa, Amoudara and Gournes. Surface seawater (1 m depth) was collected manually with a 10 L plastic bottle and brought to the lab within 20 minutes from the time of collection. Samples for nucleic acid extraction and phytoplankton identification and counting followed the same method as for the Naples samples.

### Microscopy

Identification and counts of phytoplankton species were performed using a Zeiss Axiovert 200 inverted light microscope (Carl Zeiss, Germany) at 400x magnification, following the Utermöhl method (Edler & Elbrächter, 2010). Carbon content was estimated by approximating each species to a geometric shape, calculating its mean biovolume, and converting biovolume to carbon according to Menden-Deuer & Lessard (2000) ^51^.

### Environmental parameters, nutrients and chlorophyll a analyses

Vertical profiles of temperature (°C) and salinity (PSU) were acquired using a Sea-Bird Electronics SBE 911plus v2 multi-parameter profiler, all mounted on the CTD rosette. Analyses of PO_4_, NO_3_, Si(OH)_4_ were performed using a Flow-Sys III Systea auto-analyzer using the methods reported by ^52^. Chl *a* was extracted in acetone 90% and its concentration determined according to (Holm-Hansen et al., 1965) with two different Shimadzu spectrofluorometers (Shimadzu RF-5301 PC and Shimadzu RF-6000, Shimadzu, Kyoto, Japan). Photosynthetically Active Radiation (PAR) data at 5 m depth were obtained from the Copernicus Marine Service product MULTIOBS_GLO_BIO_BGC_3D_REP_015_010 (dataset: cmems_obs-mob_glo_bgc- chl-poc_my_0.25deg_P7D-m). As PAR data were not available at the exact LTER-MC station coordinates, PAR values were extracted across a broader area encompassing the sampling location (longitude: 13.20°–15.30°E, latitude: 39.83°–41.81°N; n = 64 grid points) and averaged across all available grid points at 5 m depth for each sampling date. Where PAR data were available for the exact sampling date, those values were used directly; where no exact match existed, the temporally closest available weekly composite was selected (maximum offset: 2 days). The resulting mean PAR values were used in all subsequent statistical analyses.

### Carbon fixation rates

Carbon fixation rates of natural planktonic communities were determined by laboratory incubation experiments using the ¹³C tracer method ^53, 54^ on surface seawater samples collected in May 2025 at three different locations in the GoN. Seawater samples were pre-filtered through a 200 μm mesh to remove large mesozooplankton while retaining the phytoplankton community and incubated in six replicates in 2.5 L polycarbonate bottles (Nalgene). Prior to incubation, plankton biomass samples were collected by filtration onto pre-combusted GF/F filters (nominal pore size 0.7 μm) to determine the natural ¹³C abundance of the community. NaH¹³CO tracer solution was added to each bottle to achieve a final concentration of ∼7% of the ambient inorganic carbon pool. Incubations lasted 6 h at an *in situ* surface temperature (20 °C). Three replicates were maintained under saturating irradiance (200 µmol photons m ² s ¹) and three in the dark to correct for non-photosynthetic ¹³C uptake. At the end of each incubation, samples were filtered onto pre-combusted GF/F filters and stored frozen until analysis. Carbon isotopic composition and particulate organic carbon concentration (POC, mg C L ¹) were determined by isotope ratio mass spectrometry coupled to an elemental analyzer (EA-IRMS). Carbon fixation rates (P, pg C L^-1^ h^-1^) were calculated according to ^55^ and ^56^.

### Phytoplankton community composition

A subset of 17 out of 23 field samples collected from LTER-MC was selected for community-level analysis based on the availability of matched environmental metadata. Analyses were performed in R v4.5.0 ^57^ using the vegan ^58^ and ggplot2 ^59^ packages. Diatom community composition was analyzed from a taxon-by-sample abundance matrix of microscopy-derived cell counts. Bray-Curtis dissimilarities were calculated using vegdist(), and hierarchical clustering was used to group samples into two clusters based on community composition. Abundance data were Hellinger transformed using decostand() ^60^, and non-metric multidimensional scaling (NMDS) was performed using metaMDS() on Bray-Curtis dissimilarities calculated from the transformed data. Environmental drivers of community structure were investigated by fitting temperature, salinity, nutrients (NO□, Si(OH)□, PO□), and PAR onto the NMDS ordination using envfit(), with significance assessed by 999 permutations. One cluster was further divided into two subclusters based on contrasting *rbc*L DNA copies cell ¹ values. Differences in environmental variables, microscopy abundance, and molecular abundance metrics between all clusters and subclusters were assessed using Mann-Whitney U tests. To examine whether *rbc*L DNA copies cell ¹ were associated with larger taxa, abundance-weighted density distributions of diatom cell biovolume were compared across all clustering categories.

### Nucleic acid extraction

Phytoplankton cells were collected by vacuum filtration on Nucleopore Track-Etch membrane filters (Whatman, Germany) of 0.2 μm, 3 μm and 5 μm pore size according to the cell size of the phytoplankton species. Filters were flash-frozen in liquid nitrogen and stored at −80 °C until nucleic acid extraction. Nucleic acids were extracted using the DNeasy Plant Mini Kit and RNeasy Plant Mini Kit (Qiagen, Germany), respectively and eluted in molecular biology grade water. RNA samples were treated with TURBO DNase (Ambion, USA) to remove residual DNA through two successive incubations at 37 °C for 30 min with 2 units of DNase each. DNase was inactivated by incubation at 65 °C for 10 min in the presence of EDTA at a final concentration of 5 mM. Nucleic acid quantity was measured using Qubit 1X dsDNA HS and QubitRNA HS Assay Kits (Invitrogen, USA) and purity was assessed using a Nanodrop spectrophotometer (Thermo Fisher Scientific, USA).

### In vitro synthesis of diatom rbcL transcript fragments

Diatom *rbc*L transcript fragments were generated *in vitro* for absolute quantification of *rbc*L transcripts in cultured strains and field samples. To generate a *C. cryptica rbc*L transcript fragment, a 730 nt DNA fragment of the *C. cryptica rbc*L gene was PCR amplified with the KAPA HiFi HotStart ReadyMix (Roche, Switzerland) following manufacturer’s instructions using 10 ng of *C. cryptica* DNA as template and 0.375 μM each of forward primer *rbc*L T7 253 For (incorporating the T7 promoter sequence) and reverse primer *rbc*L 959 Rev (**Table 1**). The PCR cycling program was as follows: initial denaturation at 95 °C for 3 min, followed by 35 cycles at 98 °C for 20 s, 60 °C for 15 s and 72 °C for 1 min, with a final extension at 72 °C for 2 min. The PCR product was resolved by electrophoresis, purified and quantified as described above for the *rbc*L gene fragments. Using 0.6 μg of the purified PCR product as template, the *C. cryptica rbc*L transcript fragment was synthesized with the TranscriptAid T7 High Yield Transcription Kit (Thermo Fisher, USA) following the manufacturer’s instructions. The *C. cryptica rbc*L RNA fragment was DNase treated, purified and quantified as described above for total RNA extraction. For a *T. rotula rbc*L transcript fragment, a 315 nt RNA fragment was synthesized by Synbio Technologies (USA) based on the sequence information of the Sanger-sequenced *T. rotula rbc*L gene fragment described above. Predicted nucleotide sequences of *C. cryptica* and *T. rotula rbc*L transcript fragments are provided in **Text S1**.

**Table 1.**
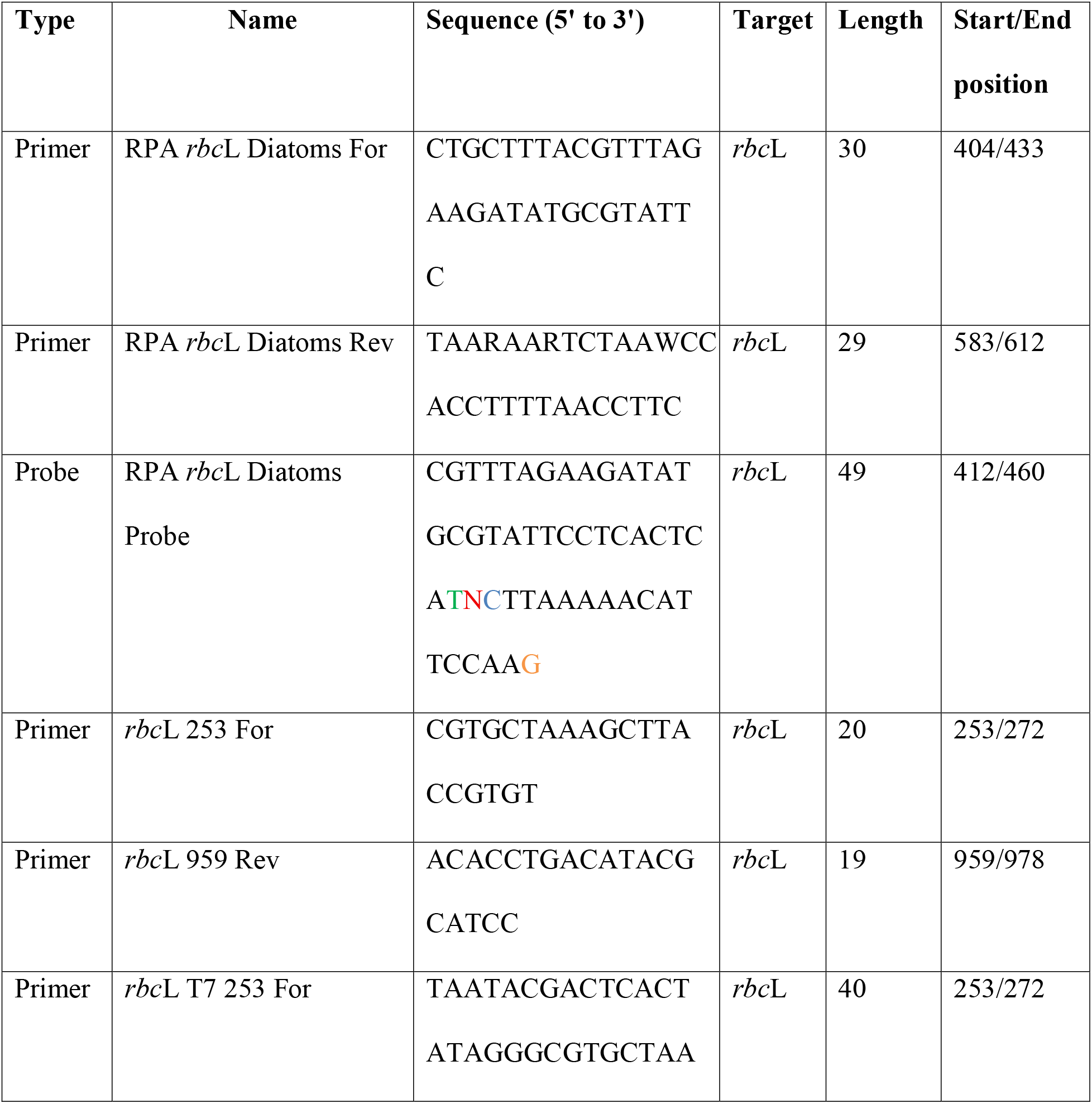
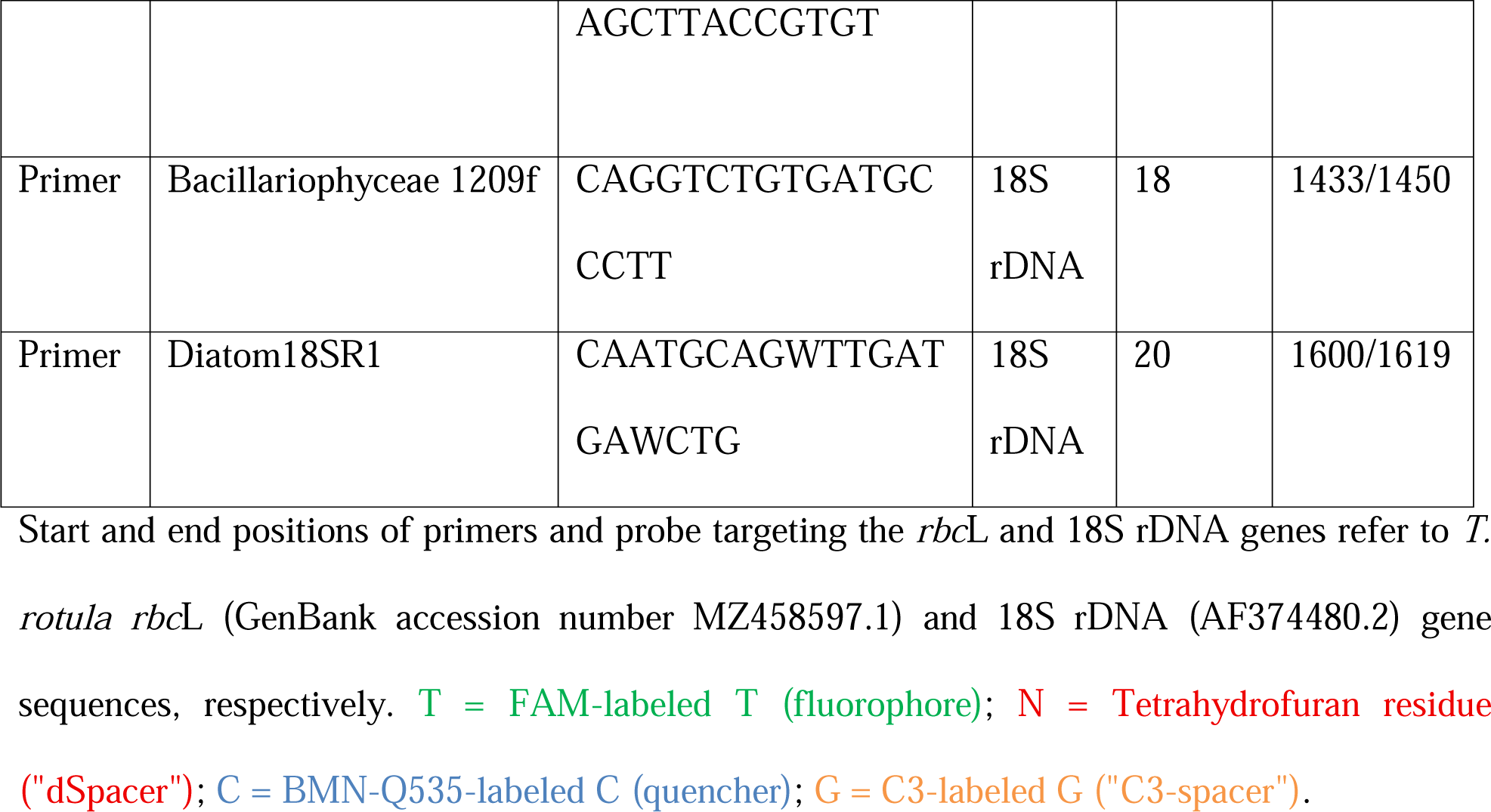
Name and nucleotide sequences of the probe and primers used in this study.

### PCR amplification and Sanger sequencing of rbcL gene fragments from diatom isolates

A 730-nt-long fragment of the *rbc*L gene was retrieved from cultured diatom species by PCR reactions that contained KAPA HiFi HotStart ReadyMix (Roche, Switzerland), 10 ng of diatom DNA and a final concentration of 0.375 μM each of forward primer *rbc*L 253For and reverse primer *rbc*L 959 Rev (**Table 1**). The PCR cycling program was as follows: initial denaturation at 95 °C for 3 min, followed by 35 cycles at 98 °C for 20 s, 60 °C for 15 s and 72 °C for 1 min, and final extension at 72 °C for 2 min. PCR products were resolved by electrophoresis on 1.5% agarose gels, excised, and purified using the Nucleospin Gel and PCR Clean-Up kit (Macherey-Nagel, Germany). Purified PCR products were quantified as described above for nucleic acid extractions and Sanger sequenced by GENEWIZ (Azenta Life Sciences, USA) using primer *rbc*L 253 For. Nucleotide sequences of the sequenced *rbc*L fragments have been deposited in the European Nucleotide Archive (https://www.ebi.ac.uk/ena/browser/home) under the study accession number PRJEB107962 (**Spreadsheet S1**). Nucleotide sequence alignments of the PCR-amplified *rbc*L fragments with the qRPA probe were carried out to identify the number of mismatches between the probe and *rbc*L target of each species. Sequence alignments are provided in **Text S2** and the number of mismatches is indicated in **Spreadsheet S3**.

### qRPA probe and primer design

The qRPA probe and primer target region was identified by analysis of multiple sequence alignment of *rbc*L sequences retrieved from the GenBank database (https://www.ncbi.nlm.nih.gov/). Multiple sequence alignment was carried out with the program MUSCLE ^61^ implemented in the software AliView ^62^ (**Text S3**). The 49-nucleotides-long FAM-labelled probe (RPA *rbc*L Diatoms Probe) was designed to hybridize to the positions 412 to 460 of the 1474-nucleotides-long diatom *rbc*L whole open reading frame sequence. The 30- nucleotides-long forward primer (RPA *rbc*L Diatoms For) was designed to hybridize from the positions 404 to 433 overlapping the 22 nucleotides of the probe 5’ end sequence. The 30 nucleotides-long reverse primer (RPA *rbc*L Diatoms Rev) was designed to hybridize from the positions 583 to 612 and contained degenerated bases at the internal positions 4, 7 and 13 (**Table 1**). The expected size of the RPA-amplified amplicon was 209 nucleotides in length spanning from positions 404 to 612. The probe and primers were synthesized by Biomers.net (Germany) and Eurofins (Luxembourg) respectively.

### Diatom rbcL qRPA assay

Assays were carried out with the lyophilized TwistAmp Exo Kit (TwistDX, UK) in a reaction with a final volume of 50 μL with the following composition: 29.5 μL rehydration buffer, a lyophilized pellet of RPA enzymes, 420 nM each of primers RPA *rbc*L Diatoms For and Rev, 120 nM of RPA *rbc*L Diatoms Probe, DNA template in water, and molecular biology grade water up to 47.5 μL. For RNA amplification, the reaction was modified as follows: a final reverse-primer concentration of 630 nM, DTT 10 mM, 200 units of reverse transcriptase (Minotech, Greece), and RNA template in water. Reactions were initiated immediately before the amplification incubation, by the addition of 2.5 μL of MgOAc to 5.6 mM final concentration. Triplicates were incubated at 37 °C for 20 min in a CFX-Connect real-time PCR detection system (Bio-Rad, USA) with SYBR/FAM fluorescence detection at 15 s intervals. Parallel no-template (water only) and no-RT (RT-qRPA in the absence of reverse transcriptase) negative controls confirmed the absence of DNA contaminants. Time-to-positivity (TTP) was defined as the time at which fluorescence increased above a threshold value, calculated as the sum of the mean value plus three times the standard deviation of the fluorescence background signal detected in the no-template negative control reactions.

### qPCR assay of diatom 18S rDNA

To quantify diatom 18S copy numbers in field samples, we first generated amplicons of the target region to use as standards for absolute quantification. The 189-nt-long fragment of the *C. cryptica* 18S rDNA gene was PCR amplified with the KAPA HiFi HotStart ReadyMix (Roche, Switzerland) using the forward primer Bacillariophyceae 1209f and reverse primer Diatom18SR1 previously described by Godhe et al., 2008 (**Table 1**) and purified by gel electrophoresis. The PCR reaction was carried out at a final volume of 25 μL with 10 ng of *C. cryptica* DNA. For qPCR, amplification used the SYBR fast qPCR kit (KAPA, Switzerland) and quantification curves were generated using a 10-fold serial dilution series ranging 10^3^–10^7^ copies per reaction. Efficiency of the 18S rDNA qPCR was 90% (R^2^ = 99).

### qRT-PCR assay of T. rotula in vitro rbcL transcript

qRT-PCR of *T. rotula in vitro rbc*L transcript fragment was carried out with the Luna universal one-step qRT-PCR kit (New England Biolab, USA) using the same primers RPA *rbc*L Diatoms For and Rev used in qRT-RPA. Reactions were run in a final volume of 20 μL with 440 nM each of forward and reverse primer and a 10-fold serial dilution of *T. rotula in vitro rbc*L transcript as template spanning from 5.45 x 10^3^ to 5.45 x 10^9^ molecules per reaction. qRT-PCR was run in a CFX-Connect real-time PCR detection system (Bio-Rad, USA). The qRT-PCR program was as follows: reverse transcription at 50 °C for 10 min, then PCR amplification with initial denaturation at 95 °C for 1 min followed by 40 cycles of denaturation at 95 °C for 10 s and hybridization/extension at 60 °C for 30 s, followed by 95 °C for 10 sec and a melt curve set from 60 to 90 °C with an increment of 0.5 °C. Efficiency of *T. rotula in vitro rbc*L transcript qRT-PCR was calculated to be 74% (R^2^ = 0.99).

## RESULTS AND DISCUSSION

### Assay inclusivity and specificity towards diatoms

We designed a taxon-wide qRPA assay for the diatom *rbc*L to efficiently amplify the gene from the majority of marine pelagic diatoms. The design targeted the most conserved *rbc*L sequence loci within diatoms but with sufficient divergence to other phytoplankton species. Primer and probe sequences were designed to match the most common sequence variants found across diatoms; limited degenerate bases were used in the primers only. In silico primer design was based on a multiple sequence alignment of the *rbc*L nucleotide sequences from 660 phytoplankton species, including 70 diatom sequences (**Text S3**). Previous studies have demonstrated that RPA efficiency decreases with increasing numbers of mismatches between the target sequence and the probe, and to a lesser extent, the primer sequences ^63–65^. In our design, both forward and reverse primer sequences exactly matched diatom *rbc*L gene sequences (**Text S3).** The probe sequence presented 0 mismatches with 55 of 70 diatom sequences, representing the majority (80%) of diatoms, 1 mismatch with 13 of 70 sequences and 3 mismatches to only 2 sequences (**Spreadsheet S4 and Text S3**). The final primer and probe set are shown in **Table 1**.

First, we assessed the assay inclusivity by testing amplification of DNA from 14 phylogenetically diverse diatom species, including both pelagic and benthic species (**Figure 2A, Spreadsheet S1**). Robust amplification was detected in 13 out of the 14 diatom species tested, with detection times (Time to Positivity, TTP) ranging from 4 to 7.3 minutes. To investigate the effect of probe and primer mismatches on assay efficiency, we PCR-amplified and Sanger-sequenced the qRPA-targeted region of each tested diatom species (**Spreadsheet S3** and **Text S2**). Amplification was delayed with increasing numbers of mismatches between the *rbc*L gene and both the probe (0 to 4 mismatches) and forward primer sequences (0 to 2 mismatches) (**Figures 2B**, **Spreadsheet S3,** and **Text S2**), in line with the literature ^63–65^. *Phaeodactylum tricornutum* was the only diatom species not to amplify, presumably due to 4 sequence mismatches with the probe. Since almost 80% of diatom sequences retrieved from GenBank presented no mismatch with the probe, with all pelagic species presenting less than 4 mismatches, the qRPA assay is expected to be inclusive across diatoms.

**Figure 2.**
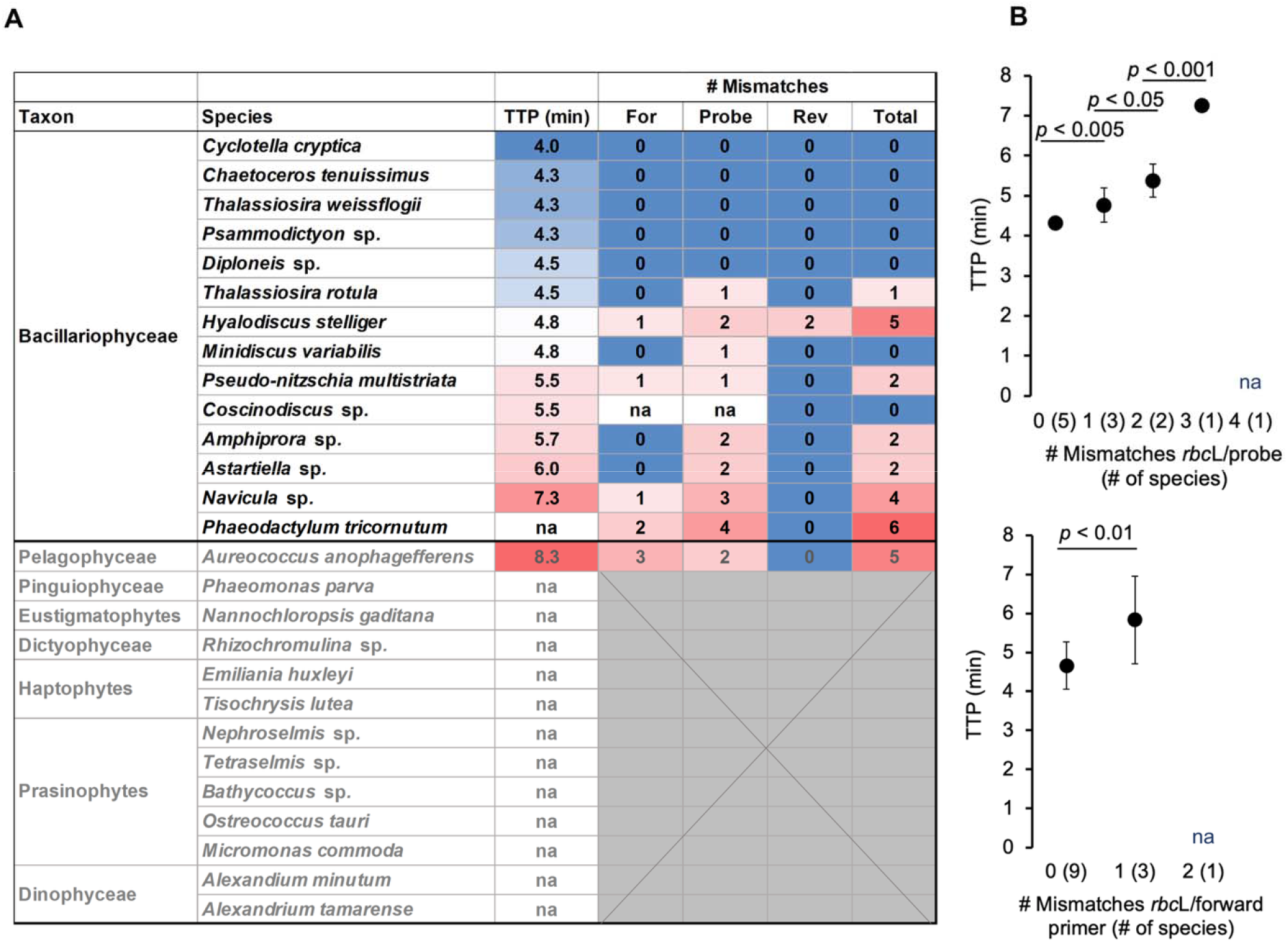
Inclusivity and specificity of the qRPA assay, and effect of primer and probe mismatches to the diatom *rbc*L gene on time to positivity (TTP). TTP was obtained using 1 ng DNA reaction^-1^ for all tests. **A** TTP against number of sequence mismatches for primers and probe. Blue to red heatmap highlights low to high TTP and mismatches. na in TTP: no amplification. na in number of mismatches: Sanger reads of insufficient quality to determine *rbc*L sequence with confidence. Grey cells: *rbc*L sequences not available for these species/strains; **B** TTP against number of sequence mismatches at the probe (top) and forward primer (bottom) binding regions. na: no amplification. The number of diatom species tested are indicated in brackets. P-values are for two-samples *t*-test. Error bars indicate standard deviation (*N* = 3).

Next, we assessed assay specificity towards diatoms by testing amplification on DNA from 13 diverse non-diatom phytoplankton species (**Figure 2A** and **Spreadsheet S1**). Non-diatom amplification was detected only in the Pelagophyceae species *Aureococcus anophagefferens*, due to its *rbc*L sequence having only 2 mismatches to the probe. This species is phylogenetically closely related to diatoms, and this is reflected in the phylogeny of its *rbc*L gene ^24^. Collectively, our results support adequate inclusivity of our *rbc*L qRPA assay towards diatoms, but cross- amplification with *rbc*L from phylogenetically closely related phytoplankton groups is an inherent challenge, as previously shown for qPCR ^26, 27, 32^. Nevertheless, diatoms are expected to dominate the phytoplankton community on a global scale ^20^.

### Assay performance and quantification metrics

Across species and molecule types, qRPA showed rapid and sensitive amplification with both DNA and RNA templates, and linear response across several orders of magnitude of input serial dilutions (**Figure 3** and **Table 2**). Benchmarking for DNA quantification included 3 widely distributed diatom species, each representing the most common *rbc*L sequence variants: *C. cryptica, T. rotula* and *H. stelliger*, which represented *rbc*L variants with 0, 1 and 2 mismatches with the qRPA probe respectively (**Spreadsheet S3** and **Text S2**). We generated calibration curves to enable *rbc*L quantification in unknown samples, using dilution series of 1) PCR-generated *rbc*L amplicons (aDNA), 2) genomic DNA and 3) cells followed by extraction, as representative realistic matrices. For DNA, the linear quantitative ranges spanned five orders of magnitude, with a limit of quantification (LoQ) of 500 copies per reaction within 11 minutes (**Figure 3A** and **Table 2**). TTP for equivalent template inputs varied between species from 1-3 min, particularly at lower input amounts (**Figure 3A**). Delayed amplification due to sequence mismatches with the probe measured using amplicon templates may have been countered by differences in *rbc*L copy numbers in their DNA (**Figure 3A** and **3B**). For example, *C. cryptica* and *T. rotula* amplified at the same time at 1 pg reaction^-1^ with the former containing 10^2^ gene copies cell^-1^ and 0 mismatches and the latter containing 10^3^ gene copies cell^-1^ and 1 mismatch. In contrast, *H. stelliger* containing 2 mismatches with the probe and 10^2^ gene copies cell^-1^ amplified significantly later than the other two species. The cellular *rbc*L copies estimated here agree with previous reports showing that, in diatoms, cellular *rbc*L copies span four orders of magnitude and correlate positively with cell biovolume under standard (or laboratory) growth conditions ^22,66^. Therefore, quantification curves to quantify diverse diatom *rbc*L sequences in natural populations should utilize the most representative species and sequence variants for a given region.

**Figure 3.**
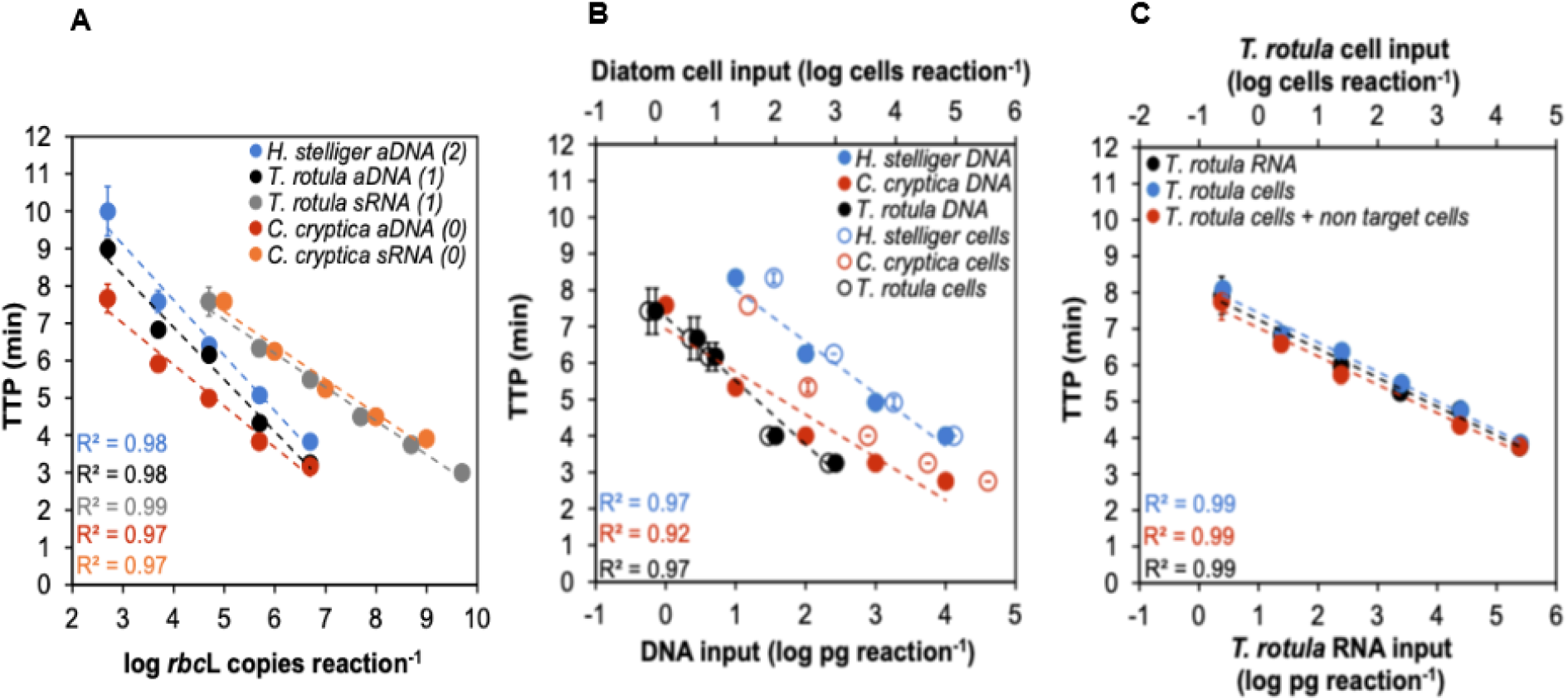
Standard curves of TTP for **A** *H. stelliger*, *T. rotula* and *C. cryptica rbc*L gene copies based on amplicon DNA (aDNA) and *T. rotula* and *C. cryptica rbc*L synthetic transcripts (sRNA) reaction^-1^. The number of mismatches between the *rbc*L gene and RPA probe sequences is indicated in brackets for each species; **B** *H. stelliger*, *T. rotula* and *C. cryptica* DNA reaction^-1^ (closed circles) and corresponding number of cells reaction^-1^ (open circles); **C** *T. rotula RNA* reaction^-1^ (bottom x-axis) and corresponding cells reaction^-1^ (top x-axis) (black circles), *T. rotula* RNA extracted from serial cell pellets of *T. rotula* either alone (blue circles) or mixed with non-target cells (red circles). Error bars indicate standard deviation (*N* = 3). The coefficients of determination of the linear regressions (dashed lines) are shown.

**Table 2.** Assay performance metrics for different input types and species. Abbreviations: *Cyclotella cryptica* (Cc), *Thalassiossira rotula* (Tr), *Hyalodiscus stelliger* (Hs). Parentheses indicate the number of sequence mismatches with the probe.

| Input type | Species | Quantitative range |
| --- | --- | --- |
| **DNA** |  |  |
| Amplicon | Cc (0), Tr (1), Hs (2) | 500 – 5,000,000 copies reaction<sup>-1</sup> |
| Genomic DNA | Cc (0) | 0.5 – 10,000 pg reaction<sup>-1</sup> |
|  | Tr (1) | 0.5 – 500 pg reaction<sup>-1</sup> |
|  | Hs (2) | 10 – 10,000 pg reaction<sup>-1</sup> |
| Cells | Cc (0) | 34 – 10,000 cells reaction<sup>-1</sup> |
|  | Tr (1) | 1 – 500 cells reaction<sup>-1</sup> |
|  | Hs (2) | 90 – 10,000 cells reaction<sup>-1</sup> |
| **RNA** |  |  |
| Synthetic RNA | Cc (0) | 10<sup>5</sup> – 10<sup>9</sup> transcripts reaction<sup>-1</sup> |
|  | Tr (1) | 10<sup>4</sup> – 10<sup>10</sup> transcripts reaction<sup>-1</sup> |
| Total RNA | Tr (1) | 2 – 200,000 pg RNA reaction<sup>-1</sup> |
| Cells | Tr (1) | 0.5 – 50,000 cells reaction<sup>-1</sup> |
| Cells + nontarget | Tr (1) | 0.5 – 50,000 cells reaction<sup>-1</sup> |

Calibration curves for RNA focused on the species with the most common sequence variants, *C. cryptica* and *T. rotula*. We also compared amplification by qRT-RPA assay against qRT-PCR as the gold standard, using *T. rotula* synthetic *rbc*L RNA (sRNA) (**Figure S1**). qRT-RPA was one order of magnitude less sensitive, which may be due to reverse transcriptase efficiency in the one-pot reaction. Finally, *T. rotula* cell extracts with and without additional RNA from non- target species, showed that background nucleic acids did not interfere with target amplification (**Figure 3C**).

Our assay was designed to quantify diatoms, a species-rich phytoplankton group with large spatial and temporal variability in cell abundance (1 – 7 x 10^7^ cells L^-1^) ^20, 67^. Therefore, optimization prioritized high inclusivity and large quantification range while retaining adequate specificity and sensitivity for relevant *in situ* concentrations. This is supported by RPA’s significant flexibility with primer and probe design enabling efficient taxon-wide amplification, compared to other isothermal approaches like loop-mediated isothermal amplification (LAMP), which requires 4-6 highly specific primers. Despite sequence variability, amplification of *rbc*L from both DNA and RNA completed in < 15 min, in line with typical amplification times of species-specific RPA assays optimized for highly sensitivity assays for early warning of harmful algal bloom species ^47, 68–71^. This versatility and rapid amplification make RPA the most suitable isothermal approach for field-based biomolecular monitoring of ecological and biogeochemical gene markers from both specific taxa and diverse functional groups.

### *Rbc*L DNA as a marker of diatom abundance in natural populations

A universal molecular marker for specific phytoplankton taxa is highly sought after to enable automated *in situ* plankton monitoring. We tested diatom *rbc*L as a marker of abundance across diverse natural communities, by comparing qRPA amplification against taxonomy information and cell counts by microscopy. Surface water samples from the LTER-MC site in the GoN, Italy ^72, 73^ spanned a 7-year period (2019-2025) (**Figure 4A** and **Spreadsheet S2**). Diatom absolute abundance varied over four orders of magnitude across the sample set (10^4^ – 10^7^ diatom cells L^-1^) and relative abundance ranged from 15% to 84% of the total phytoplankton (diatoms, coccolithophores, dinoflagellates, and other flagellates). Additional seawater samples were collected at three sites along the northern coast of Crete, Greece (**Spreadsheet S2**) to represent ultra-oligotrophic conditions with low diatom abundances (10^3^ – 10^4^ diatom cells L^-1^) ^74^.

**Figure 4.**
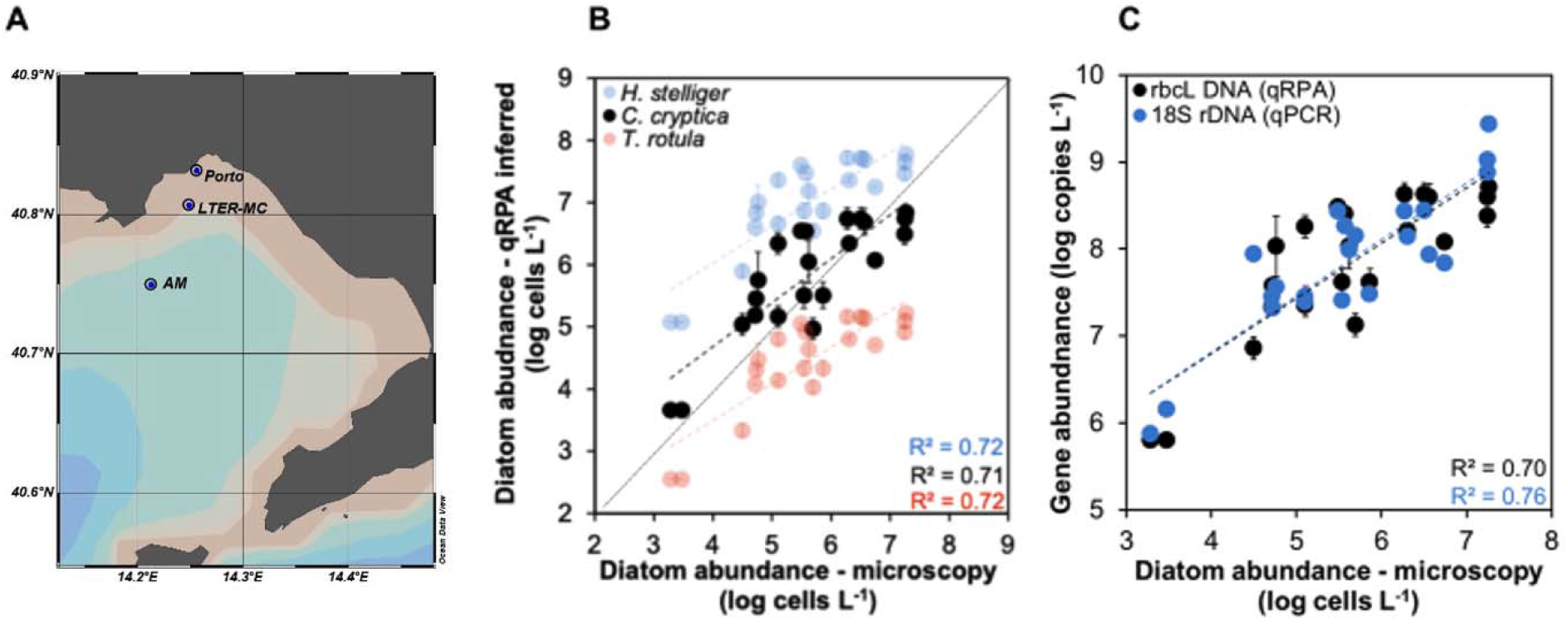
Quantification of diatom abundance in natural samples by qRPA. **A** Map of the Gulf of Naples, Italy, indicating the sample collection sites, Ammontatura (AM), LTER-MC, and Porto. The map was created using the Ocean Data View Software (https://odv.awi.de) ^77^; **B** Comparison of diatom abundance determined by microscopy to that inferred by qRPA using each diatom species-specific calibration curves. The dotted line indicates the line of identity; **C** Comparison of diatom abundance determined by microscopy to that inferred by qRPA and 18S qPCR. Dashed lines are linear regressions and their coefficients are shown. Error bars indicate standard deviation (*N* = 3).

We inferred diatom absolute abundance in field samples using our three species-specific qRPA standard curves correlating *rbc*L gene copies to cell numbers L^-1^ (**Figure 4B)**. Absolute abundance inferred from the *C. cryptica* qRPA standard curve best matched the microscopy cell counts (Pearson’s r = 0.84*, p* = 1.24 x 10^-6^, *N* = 22). In contrast, diatom absolute abundance values inferred from the *T. rotula* and *H. stelliger* standard curves were consistently higher and lower than microscopy counts, respectively (**Figure 4B**). The *C. cryptica rbc*L sequence at the probe target region is representative for most diatom species (**Spreadsheet S3**), making it the species of choice for calibrated *rbc*L-based diatom abundance estimates in natural samples.

We further validated *rbc*L as an abundance marker by comparing the degree of correlation to diatom abundance of diatom *rbc*L gene and 18S rDNA copies L^-1^, estimated by our qRPA and a widely used diatom qPCR assay, respectively ^19^. The correlation of diatom abundance to 18S rDNA copies L^-1^ (Pearson’s r *=* 0.87*, p* = 1.19 x 10^-7^, *N* = 22) was only marginally higher than to *rbc*L gene copies L^-1^ (Pearson’s *r =* 0.84*, p* = 1.24 x 10^-6^, *N* = 22) (**Figure 4C** and **Spreadsheet S5**). Our results confirm previous reports of a positive correlation between qPCR-based quantification of diatom *rbc*L gene and 18S rDNA copies L^-1^ in natural diatom communities ^31^.

Diatom 18S rDNA and *rbc*L genes are well-known markers of diatom abundance in the field ^19–22, 24, 25^. However, substantial variation in 18S rDNA and *rbc*L gene copies cell^-1^ across diatom species, which has been shown to be broadly positively correlated with cell biovolume, can lead to inaccurate estimates of diatom abundance ^19, 22, 75, 76^. Gene marker-based quantification of diatom abundance has therefore recently focused on targeting single-copy genes such as *psb*O ^75^. In our analysis, the average diatom biovolume and *rbc*L gene copies cell^-1^ across the analyzed field samples ranged from 3.95 × 10^2^ to 3.43 × 10^3^ μm^3^ and 1 × 10^1^ to 2 × 10^3^, respectively. However, no correlation was found between the average diatom biovolume and *rbc*L gene copies cell^-1^ (Pearson’s r = 0.13*, p* = 0.64, *N* = 15). Our results therefore support that quantification of diatom *rbc*L gene copies L^-1^ in field samples, when based on a robust calibration of *rbc*L gene copies cell^-1^, is a powerful approach to infer diatom absolute abundance on a large scale.

### Diatom *rbc*L DNA quantification reveals enhanced photosynthetic potential in winter diatom populations

We combined molecular analyses and abundance data to investigate temporal environmental controls on photosynthetic physiology of natural populations. Our dataset was complemented by measurements of chlorophyll *a*, nutrients (nitrate, phosphate and silicate) and physical parameters (temperature, salinity and PAR) (**Spreadsheet S2**). Temperature and PAR revealed a strong seasonal signal (**Figure 5B**). Macronutrients and salinity did not vary significantly, but silicate was higher in winter (**Figure 5B**).

**Figure 5.**
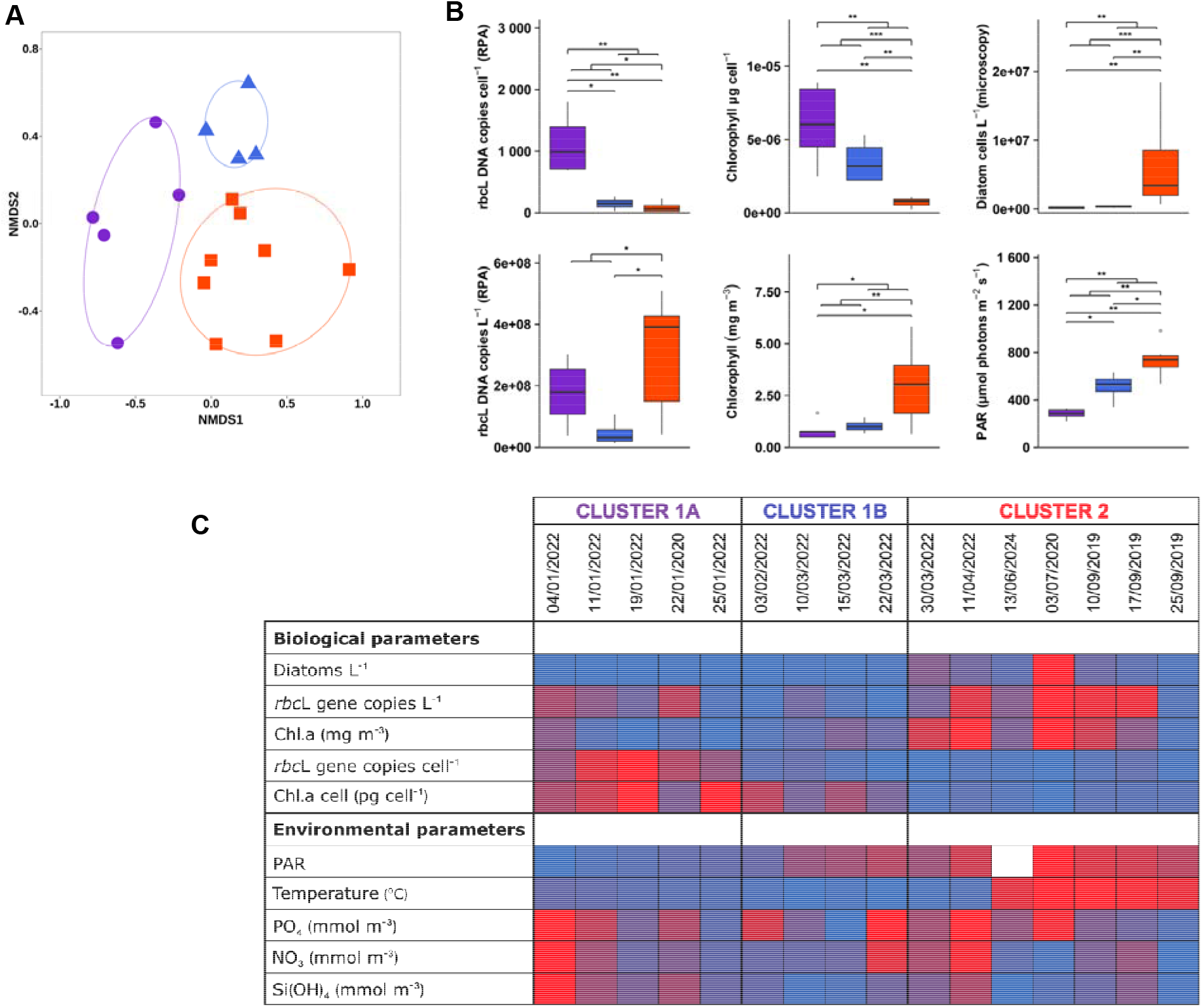
Diatom community composition and differences in environmental and biological variables across community clusters. **A** NMDS ordination of diatom community composition across field samples, with ellipses denoting cluster groupings. Points represent individual samples coloured by cluster: Cluster 1A (purple circles; January), Cluster 1B (blue triangles; February-March), and Cluster 2 (orange squares; late March-September). Clusters 1A and 1B form the broader Cluster 1, distinguished from Cluster 2 by diatom taxonomic composition and further resolved by contrasting *rbc*L DNA copies cell ¹ values; **B** Distribution of environmental and biological variables for each cluster; central lines indicate medians and boxes represent the interquartile range (IQR). Differences between clusters were assessed by Mann-Whitney U tests; statistically significant differences are indicated by brackets and asterisks. Nested brackets compare pooled Cluster 1 (1A + 1B) vs Cluster 2, and Cluster 1A vs pooled Cluster 1B + Cluster 2. Significance levels: *p* < 0.05 = *, *p* < 0.01 = **, *p* < 0.001 = ***; **C** Heatmap of qRPA inferred values of *rbc*L L^-1^ and *rbc*L diatom cell^-1^, and parallel measurements of biological and physicochemical variables (chl a: chlorophyll a; PAR: Photosynthetic Active Radiation).

Multivariate statistical analyses identified two major clusters of diatom communities based on taxonomic composition. Clusters 1 (A and B) comprised winter and early spring samples, and Cluster 2 comprised late spring, summer and autumn samples (**Figure 5A** and **5B**). These clusters were in line with changes in temperature, PAR and silicate (**Figure 5C**), reflecting the strong seasonality of phytoplankton community dynamics previously reported in the Gulf of Naples ^78–81^.

Within Cluster 1, a distinct subcluster 1A was composed exclusively of winter samples, and contained diatoms with 10.4-fold higher *rbc*L DNA copies cell^-1^ and 6.5-fold higher chlorophyll *a* cell^-1^, compared to all other samples (**Figure 5B, 5C** and **Spreadsheet S6**). This increase in cellular photosynthetic machinery was not a consequence of a seasonal shift towards larger cells (**Figure S2**). Instead, the main environmental feature distinguishing Cluster 1A was a significantly, 2.3-fold lower PAR level. Silicate concentrations were also 2.98-fold higher in cluster 1A relative to cluster 1B but not significantly different to Cluster 2. This finding is consistent with the view that winter blooms are primarily controlled by physical forcing, particularly light availability and water column stability, rather than by chemical (nutrient concentrations) or biological factors (grazing) ^79, 82^. These results indicate that winter diatom communities with broadly similar taxonomic composition can differ substantially in physiological state. Increased cellular *rbc*L gene and chlorophyll *a* content represents a photoacclimation response to reduced light availability. This is consistent with established low-light acclimation responses in phytoplankton, including enlargement of chloroplasts and increased cellular content of light-harvesting pigments and RuBisCO protein ^15, 83, 84^. Photoacclimation of phytoplankton has also been reported across the Mediterranean based on satellite data, with winter populations containing almost double chlorophyll per unit biomass ^15^. These physiological changes are relevant for satellite-based estimates of primary productivity, which are based on chlorophyll concentrations, and may underestimate the enhanced photosynthetic potential of low-light-acclimated communities ^10, 15^.

### Diatom *rbc*L RNA as a proxy for carbon fixation in natural populations

We investigated the application of *rbc*L transcript quantification for inferring diatom carbon fixation rate. First, we compared *rbc*L transcripts L^-1^ to carbon fixation rates measured by ^14^C incorporation in *T. rotula* cultures across cell abundances ranging three orders of magnitude (**Figure S3**). *Rbc*L transcript abundance correlated well with carbon fixation rates (Pearson’s r = 0.98, *p-value* < 0.05, *N* = 4), which ranged from 1.20 to 2.77 pg C h^-1^ cell ^-1^, comparable to that reported previously for the same species (1.25 to 5 pg C h^-1^ cell ^-1^) ^85^.

We then evaluated the capacity of *rbc*L expression as a proxy for primary productivity in field samples. As *rbc*L transcripts L^-1^ correlated less strongly with diatom abundance than *rbc*L gene copies L^-1^ (**Figure S4**), reflecting the more dynamic regulation of *rbc*L expression, we focused our analyses on a set of three samples with parallel carbon fixation rate measurements using ^13^C incorporation. These samples represented different locations in the GoN (Ammontatura, LTER-MC, and Porto) (**Spreadsheet S2**). Carbon fixation rates ranged from 1.81 x 10^6^ to 8.76 x 10^6^ pg C L^-1^ h^-1^, consistent with previously reported values across the Mediterranean Sea ^54, 86–91^. *rbc*L transcripts L^-1^ followed a similar trend to carbon fixation rates across the three samples **(Figure 6A**), suggesting that diatoms could be responsible for most of the measured carbon fixation. We tested whether molecular data could predict carbon fixation values by applying the only published empirical relationship between phytoplankton *rbc*L copies L^-1^ and carbon fixation rates established by John et al. (2007) ^27^. However, the predicted diatom-specific carbon fixation rate was 9–18--fold larger than the measured values for the whole community. The equation used is largely based on data from the Mississippi Delta and Gulf of Mexico, with high riverine nutrient loads ^92, 93^, and likely a fundamentally different phytoplankton community ^94^. Predicting carbon fixation values from molecular data requires a well-characterized relationship of *rbc*L transcription to carbon fixation rate from different regions, seasons and taxonomic groups. Recent approaches measuring RuBisCo protein concentrations in natural populations ^95^, combined with isotope-based carbon fixation rates and *rbc*L RNA quantification, would provide essential calibration datasets for using *rbc*L RNA as a molecular proxy for primary productivity.

**Figure 6.**
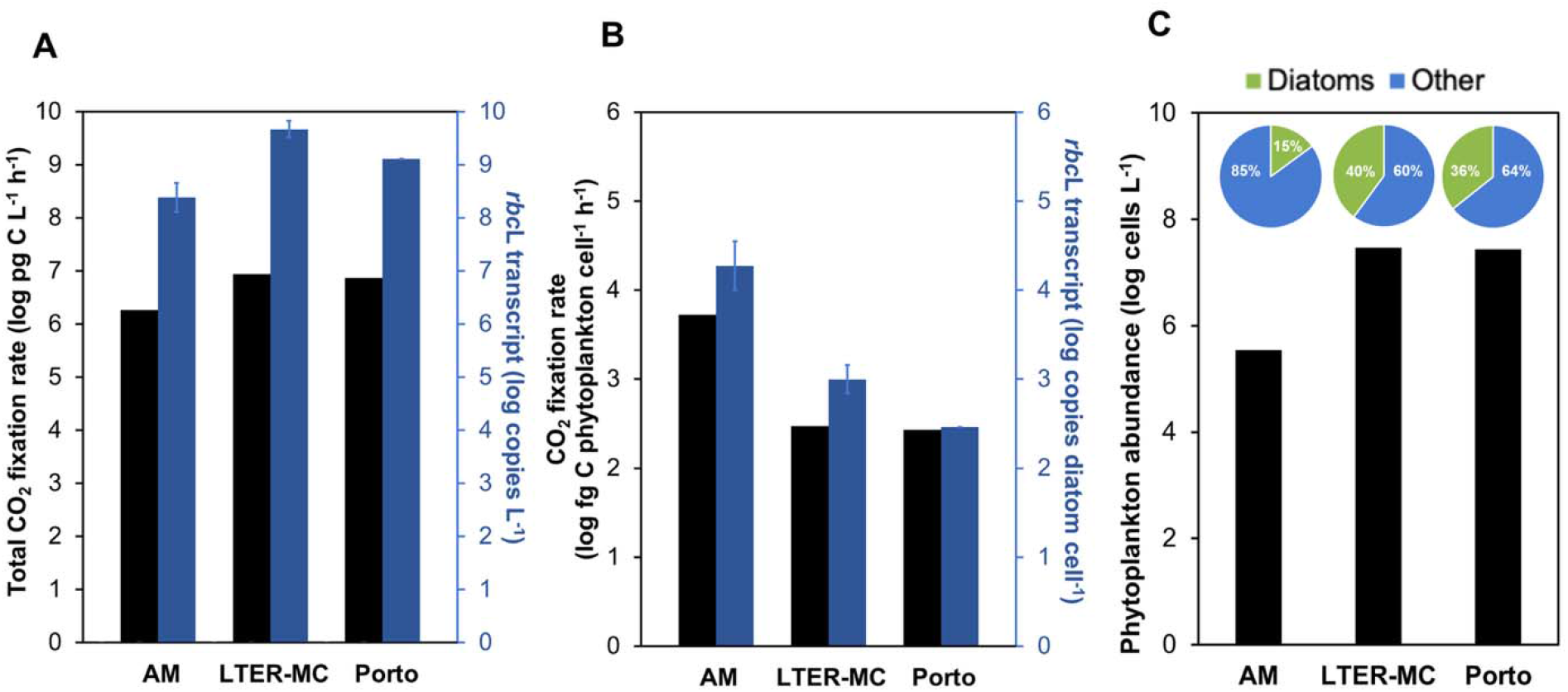
Diatom carbon fixation rate estimates from diatom *rbc*L transcript abundance in diatom communities. **A** Volumetric carbon fixation rate measured by carbon isotope incorporation (^13^C) and *rbc*L transcript L^-1^ inferred by qRT-RPA in Ammontatura (Am), LTER-MC and Porto samples. Error bars indicate standard deviation (*N* = 3); **B** Average carbon fixation rate per phytoplankton cell (left y-axis) and diatom *rbc*L transcripts diatom cell^-1^ (right y-axis) in Ammontatura (Am), LTER-MC and Porto samples. Error bars indicate standard deviation (*N* = 3); **C** Total phytoplankton absolute abundance (y-axis) and relative abundance of diatom and other phytoplankton cells (pie charts) determined by microscopy in Ammontatura (Am), LTER-MC and Porto samples.

Finally, we examined whether *rbc*L RNA can provide a measure of diatom cellular photosynthetic activity. For comparative analyses, we scaled bulk ^13^C-derived carbon fixation rates to total phytoplankton abundance and diatom *rbc*L transcripts L^-1^ to diatom abundance; we did not scale *rbc*L transcripts to gene copies, as our results showed these are also physiologically dynamic (**Figure 5**). Ammontatura stood out compared to LTER-MC and Porto, with two orders of magnitude lower total phytoplankton abundance, and 40% lower relative diatom abundance (**Figure 6C**), while maintaining comparable productivity to the other sites (**Figure 6A**). When scaled to phytoplankton abundance, Ammontatura had the highest phytoplankton cellular carbon fixation rate of 5.27 pg C h^-1^ cell^-1^, while LTER-MC and Porto samples had ca. 17-fold lower values of 0.30 and 0.27 pg C h^-1^ cell^-1^, respectively (**Figure 6B**), all within the range of values reported for diatom cultured isolates ^85, 96, 97^. The higher productivity at Ammontatura was reflected in diatom *rbc*L transcripts L^-1^ being 22-fold and 65-fold higher than LTER-MC and Porto, respectively (**Figure 6B**). This suggests that diatoms were more photosynthetically active at this site, resulting in an almost equivalent primary productivity despite a markedly lower abundance. These results demonstrate that *rbc*L eRNA, when combined with abundance data, is a powerful tool to estimate diatom-specific cellular photosynthetic activity in natural populations.

### Technological advance

We have developed a new molecular approach employing RPA-based isothermal amplification to rapidly (< 15 min) quantify *rbc*L gene copies and transcripts from the majority of marine diatoms. With calibration across diatom diversity, and with validation against microscopy-based cell counts, we demonstrate its application to estimate diatom abundance and photosynthetic physiology in natural samples. We use this approach to reveal taxon-specific seasonal responses in photosynthetic physiology, demonstrate good agreement with bulk carbon fixation measurements, and enable observations of differing diatom photosynthetic activity associated with different locations and phytoplankton community composition. These measurements are valuable for improving our monitoring capability and understanding of phytoplankton dynamics and contribute to primary productivity estimates from satellite-based chlorophyll measurements. Furthermore, our results support the case for integrating molecular and optical approaches in long-term monitoring ^98^ as a powerful combination for phytoplankton ecology. *In situ* imaging cytometry instruments are already in widespread use and the isothermal amplification approach presented here is ideally suited for implementation in autonomous *in situ* eDNA and eRNA analyzers that are currently being developed for ocean biomolecular observation.

## ASSOCIATED CONTENT

### Supporting Information

The following files are available free of charge:

Supplementary information (docx): Figures S1-4, Material and Methods S1-3, and Text S1-3.

Supplementary spreadsheets (xls): Spreadsheets 1-6.

## AUTHOR INFORMATION

### Author Contributions

Conceptualization: Martha Valiadi; Methodology: Frédéric G. Verret, Katherine Hartle- Mougiou, Alexandra Peltekis, Stella Psarra, Martha Valiadi; Investigation: Frédéric G. Verret, Christos Chantzaras, Alexandra Peltekis, Stella Psarra, Martha Valiadi; Formal Analysis: Frédéric G. Verret, Katherine Hartle-Mougiou, Alexandra Peltekis, Stella Psarra; Resources: Frédéric G. Verret, Francesca Margiotta, Diana Sarno, Ulisse Cardini, Marco Alba, Valeria Pizziol, Roberto Gallia, Isabella Percopo, Ferdinando Tramontano, Maira Maselli, Antonio Novellino, Marina Montresor; Supervision: Martha Valiadi; Funding Acquisition: Matthew C. Mowlem, Electra Gizeli, Martha Valiadi; Writing – Original Draft: Frédéric G. Verret, Katherine Hartle-Mougiou, Martha Valiadi; Review & Editing: Frédéric G. Verret, Katherine Hartle- Mougiou, Francesca Margiotta, Diana Sarno, Marina Montresor, Matthew C. Mowlem, Electra Gizeli, Martha Valiadi. All authors have read and agreed to the published version of the manuscript.

## ACKNOWLEDGMENTS

The site LTER-MC (DEIMS.ID:https://deims.org/0b87459a-da3c-45af-a3e1-cb1508519411) belongs to the Long Term Ecological Research national and international networks: LTER-Italy (https://ror.org/05ma8mw15), LTER-Europe and ILTER. We thank the LTER-MC team for their support.

## ABBREVIATIONS

eDNA/eRNA: environmental deoxyribonucleic/ribonucleic acid;
*rbc*L: ribulose-1,5-bisphosphate carboxylase/oxygenase large subunit
RuBisCO: Ribulose-1,5-bisphosphate carboxylase/oxygenase
qRT-RPA: quantitative reverse transcription recombinase polymerase amplification
TTP: time to positivity
qRT-PCR: quantitative reverse transcription polymerase chain amplification
LAMP: Loop-mediated isothermal amplification
LTER-MC: long term ecological research MareChiara
PAR: photosynthetic active radiation
Chl.a: chlorophyll a

