## Supplementary Data for "Rapid isothermal amplification of the diatom RuBisCO gene from eDNA and eRNA reveals their abundance and photosynthetic physiology"

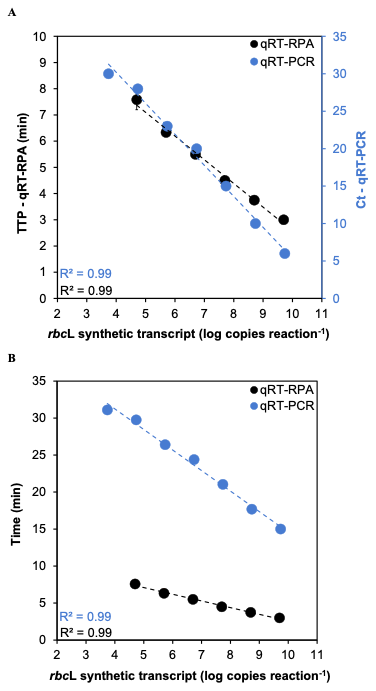

**Figure S1.** qRT-RPA versus qRT-PCR amplification of *T. rotula rbc*L transcript. **A** qRT-RPA time to positivity (TTP) and qRT-PCR cycle threshold (Ct) as a function of *T. rotula rbc*L transcripts reaction^-1^. **B** Same values as in A but after conversion of the Ct to time (in min) considering t = 0 at the start of the RT step of the qRT-PCR. The coefficients of determination of the linear regressions (dashed lines) are shown. Error bars indicate standard deviation (N = 3).

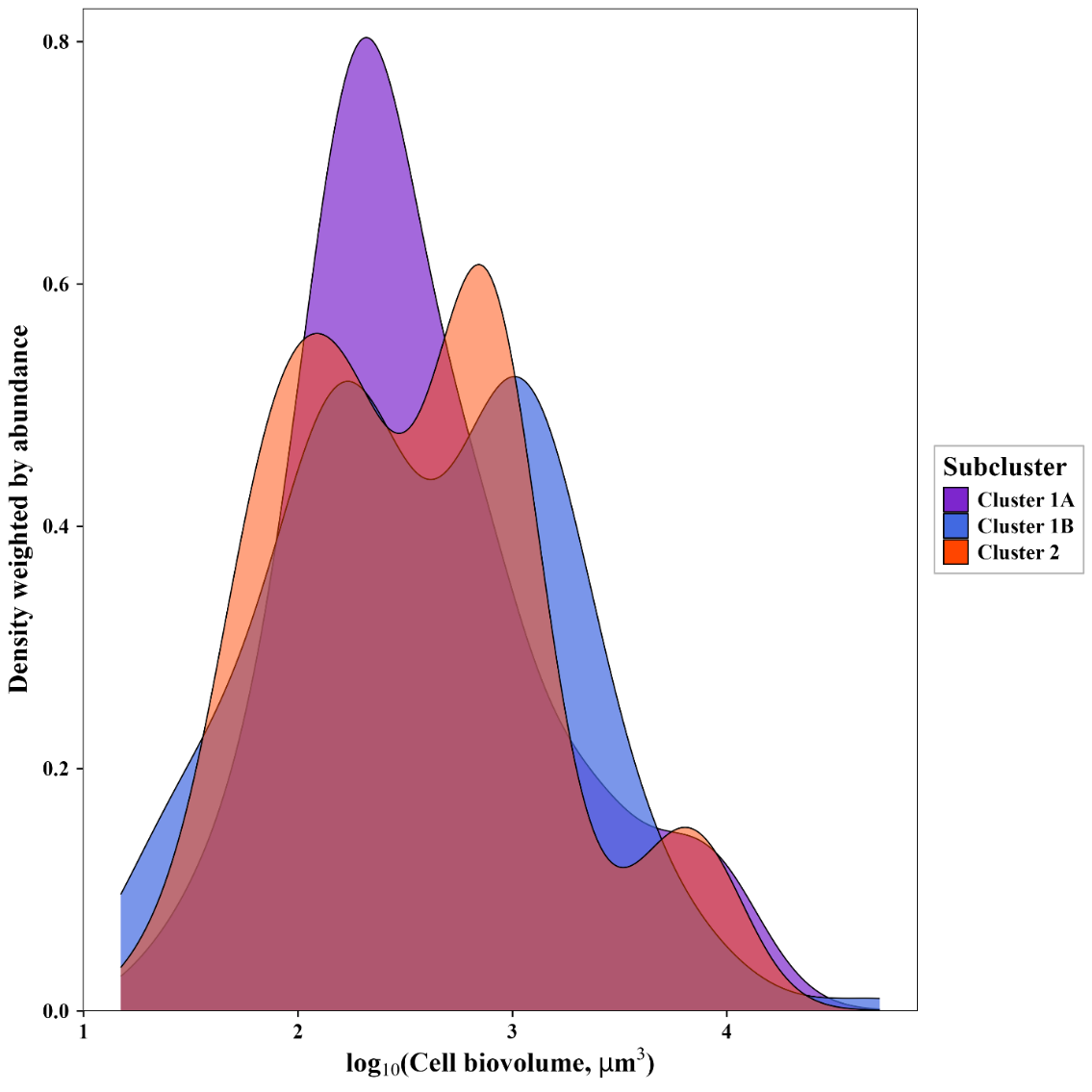

**Figure S2.** Density distribution of diatom cell biovolume across DNA per cell-defined subclusters. Curves show abundance-weighted kernel density distributions of log₁₀-transformed cell biovolume (µm³) for diatom taxa identified by microscopy within each subcluster (Cluster 1A, Cluster 1B, and Cluster 2). Density estimates were weighted by taxon abundance to reflect the dominant community cell-size structure within each subcluster. Overlapping distributions indicate broadly similar community biovolume structure among subclusters despite differences in *rbc*L DNA per microscopy-estimated cell.

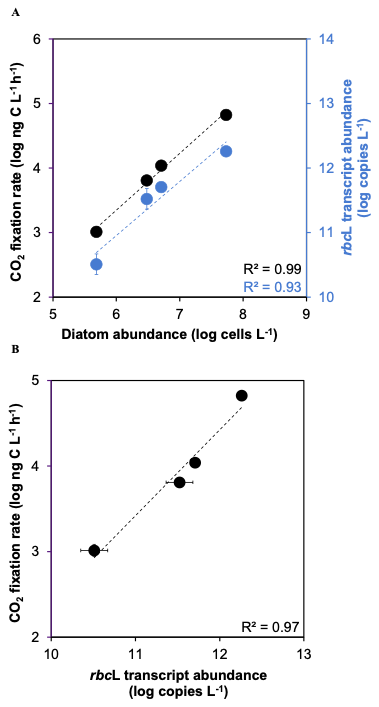

**Figure S3. A** Carbon fixation rate measured by ^14^C incorporation (left y-axis) and *rbc*L transcripts L^-1^ inferred by qRT-RPA (right y-axis) as a function of *T. rotula* cells L^-1^ counted by microscopy (x-axis). **B** Standard curve of carbon fixation rate versus *rbc*L transcripts L^-1^ derived from A. Error bars for *rbc*L transcript L^-1^ and carbon fixation rate indicate standard deviation and standard error respectively (*N* = 3). The coefficients of determination of the linear regressions are shown.

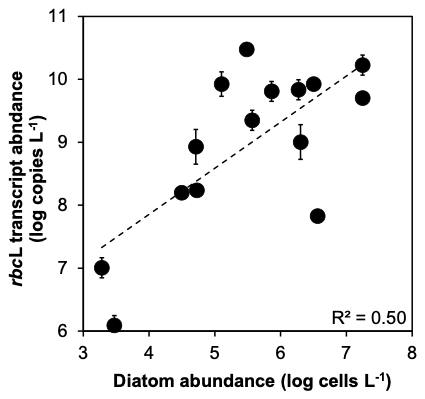

**Figure S4.** *Rbc*L transcripts abundance inferred by qRT-qRPA versus diatom abundance determined by microscopy in field samples. The equations and coefficients of determination of the linear regression (dashed line) are shown. Error bars indicate standard deviation (*N* = 3).

**Materials and Methods S1.** Isolation and identification of diatom species collected from the bay of Heraklion, Crete, Greece.

Sea water samples were collected during a cruise expedition organized by the Hellenic Center for Marine Research (HCMR) in the bay of Heraklion, Crete, Greece (35°23'31.81"N; 25°12'27.33"E) in summer 2014. One milliliter of sampled sea water was incubated in 20 mL of natural sea water based f/2 media ^1^. Phytoplankton cells were grown at 18 ºC exposed to a 12:12 h light:dark photoperiod of 70 *μ*mol photons m^-2^ s^-1^ using fluorescent tubes. After one week of culture, single cell diatoms were identified and isolated under microscope using a micropipette and propagated in monoclonal cultures. Genomic DNA extraction, PCR amplification, purification and Sanger sequencing of their *rbc*L gene fragments was carried out as described in Materials and Methods. Blastn alignments of the sequenced *rbc*L fragments was carried out on the GeneBank database (<https://www.ncbi.nlm.nih.gov/>) and genus of the Blastn best hits were assigned to the corresponding diatom isolates. Complementary inspection of cell morphology under light-microscopy was carried out to confirm the identity of the diatom isolates at the genus level. Details of the diatom species isolated from Greece are provided in **Spreadsheet S1.**

**Materials and Methods S2.** Validation of qRT-RPA based quantification of diatom *rbc*L transcript L^-1^ versus CO_2_ fixation rate in *T. rotula* laboratory culture.

*Culture conditions*

*T. rotula* was cultured in f/2+Si medium using natural seawater (salinity 38.8 PSU) collected from Sitia, Crete, which was autoclaved and filtered through a 200 μm nylon net to remove large particles and potential contaminants. Cultures were grown under 19 ± 1°C with a light intensity of 70 μmol m⁻² s⁻¹ provided by white neon light with a 12/12 h light/dark photoperiod (light phase from 7:00 to 19:00). The light intensity was set to 70 μmol m⁻² s⁻¹ to ensure optimal photosynthetic activity without inducing photoinhibition, a condition where excess light damages photosystem II ^2^ and generates reactive oxygen species ^3^. Light intensity was monitored with a Hansatech QSPAR quantum sensor (Hansatech Instruments, United Kingdom). One day before cell counting, measurement of CO_2_ fixation rate and cellular chlorophyll a content, four dilutions of the main culture were prepared in triplicate (biological replicates) to target 40000, 4000, 2000, and 400 cells/ml in 100 mL final volume (12 cultures in total). This cell dilution range was selected as to fall within the previously established linear quantitative range of the *rbc*L RT-qRPA assay for TTP versus number of cells per reaction. To determine cell abundance, 2 mL samples of each culture were fixed with 2% Lugol’s solution (20 μL) and stored at 4°C for one to two days prior to counting. On the day of counting, 1 mL from each fixed sample was transferred to a Sedgwick-Rafter counting chamber and observed under a light microscope.

*RNA extraction and rbcL qRT-RPA assay*

Seventy mL of each culture were sampled by filtration as described in Material and Method. RNAs were extracted using the magMAX-96 total RNA isolation kit (Applied Biosystems, Thermo Fisher Scientific, Waltham, Massachusetts, United States). RNAs were DNase treated and quantified as described in Material and Methods. Diatom *rbc*L qRT-RPA assays were carried as described in Material and Methods.

*Measurement of gross primary productivity by ^14^C incorporation*

Primary production was assessed using the ^14^C incorporation method originally described by Nielsen 1952 ^4^, based on the *in-situ* field protocol of Lagaria et al 2017 ^5^ and the *in vitro* protocol of Rivkin 1989 ^6^, which were modified and adapted for the present experiments using laboratory cultures. For these measurements, a single replicate was used across all four dilution levels, with four subsamples (2.10 mL each) prepared for each dilution: three light replicates (in transparent Eppendorf tubes to allow photosynthesis under light conditions and thus quantify photosynthetic ^14^C uptake) and one dark control (in an Eppendorf tube wrapped in black tape to create dark conditions in order to quantify non-photosynthetic ^14^C uptake), making a total of 16 subsamples. The 16 subsamples were inoculated with H^14^CO_3_^-^ (4.8 μCi mL^-^¹ working solution) to achieve a final activity of 0.1 μCi mL^-^¹ in each Eppendorf tube, following the conditions described by Rivkin 1989 ^6^. Incubation was initiated approximately 6 h after the light period was initiated, allowing the microalgal cultures in all dilution treatments to reach a comparable physiological state. Before the start of the incubation, the tubes were thoroughly mixed, and 50 μL subsamples were immediately taken from each dilution for total counts (×4) and placed in 5-mL vials containing 2 mL of scintillation cocktail (Ultimagold). Light-exposed samples were incubated for approximately 2 h at a constant irradiance of 70 μmol photons m^-^² s^-^¹, while the corresponding dark controls were incubated simultaneously under dark conditions. Following incubation, all samples were collected under low-light conditions and filtered through 25-mm GF/F filters. The filters were subsequently rinsed with seawater filtered through 0.2-μm pore-size polycarbonate filters and acidified overnight with 400 μL of 5% HCl to remove residual inorganic ¹^4^C. The retained ¹^4^C activity was then quantified using a Packard A2100TR liquid scintillation counter after addition of 2 mL of a scintillation fluor (Ultimagold).

**Text S1.** Nucleotide sequences of *T. rotula* and *C. cryptica RbcL* RNA fragments synthesized *in vitro*.

>T. rotula RbcL RNA fragment

TAATTTAACAGCGTCTATTATTGGTAACGTTTTTGGATTTAAAGCAGTTGCTGCTTTACGTTTAGAAGATATGCGTATTCCTCACTCATATTTAAAAACATTCCAAGGTCCTGCTACAGGTATTATTGTAGAACGTGAACGTTTAAATAAATATGGTACACCATTATTAGGTGCAACTGTAAAACCTAAATTAGGTCTTTCTGGTAAAAACTATGGTCGTGTAGTTTATGAAGGTTTAAAAGGTGGTTTAGACTTCTTAAAAGATGATGAAAACATTAACTCTCAACCATTCATGCGTTGGAGAGAACG

>C. cryptica RbcL RNA fragment

TTTGGATTTAAAGCAGTTGCTGCTTTACGTTTAGAAGATATGCGTATTCCTCACTCATACTTAAAAACATTCCAAGGTCCTGCTACAGGTATTGTTGTAGAACGTGAACGTTTAAACAAATATGGTACTCCATTATTAGGTGCTACTGTAAAACCTAAATTAGGTCTTTCTGGTAAAAACTATGGTCGTGTAGTTTATGAAGGTTTAAAAGGTGGTTTAGACTTCTTAAAAGATGATGAAAACATTAACTCTCAACCATTTATGCGTTGGAGAGAACGTTTCTTAAACTGTTTAGAAGGTATTAATCGTGCATCTGCTGCAACTGGTGAAGTTAAAGGTTCTTACTTAAACGTTACAGCAGCTACAATGGAAGAAGTATACAAACGTGCTGAGTATGCTAAAACGATCGGTTCTGTAATTATCATGATCGATTTAGTAATGGGTTACACTGCAATTCAATCAATTGCTTACTGGGCTCGTGAAAATGATATGATTTTACATTTACACCGTGCAGGTAACTCAACTTACGCTCGTCAAAAAAATCATGGTATTAACTTCCGTGTTATTTGTAAATAGATGC

**Text S2.** Multiple alignment (fasta format) of Sanger sequenced *rbc*L gene sequences from the diatom species grown in laboratory and used to identify the number of mismatches with the qRPA probe and primers.

>RPA *rbc*L Diatoms For

---------------------------------------------------------------------------------------------------------CTGCTTTACGTTTAGAAGATATGCGTATTC----------------------------------------------------------------------------------------------------------------------------------------------------------------------------------------------------------------------------------------

>RPA *rbc*L Diatoms Probe

-----------------------------------------------------------------------------------------------------------------CGTTTAGAAGATATGCGTATTCCTCACTCATACTTAAAAACATTCCAAG-------------------------------------------------------------------------------------------------------------------------------------------------------------------------------------------------------------

>RPA *rbc*L Diatoms Rev

--------------------------------------------------------------------------------------------------------------------------------------------------------------------------------------------------------------------------------------------------------------------------------------------GAAGGTTTAAAAGGTGGWTTAGAYTTYTTA-----------------------------------------------------

>*Phaeodactylum tricornutum*

---------------------CGCTTTTATCGCATATGAATGTGATTTATTTGAATAACTTAACAGCTTCTATTATTGGTAACGTATTCGGTTTCAAAGCTGTATCTGCGTTACGTTTAGAAGATATGCGTATCCCTCATTCTTACTTAAAAACGTTCCAAGGTCCTGCTACTGGTGTAATTGTAGAACGTGAACGTTTAAACAAATATGGTATTCCATTATTAGGTGCTACAGTAAAACCTAAATTAGGTTTATCTGGTAAAAACTATGGTCGTGTAGTTTATGAAGGTTTAAAAGGTGGTTTAGACTTCTTAAAAGATGATGAGAATATTAACTCACAACCCTTCATGCGTTGGAGAGAACG---

>*Pseudonitzschia multistriata*

-----------GATGTTTTCTTCGCTTTCTCGCATATGAATGTGATTTATTTGAAGAACTTAACAGCGTCTATTATTGGTAACGTATTCGGTTTCAAAGCTGTATCAGCTTTACGTTTAGAAGACATGCGTATTCCTCACTCATACTTAAAAACATTCCAAGGTCCTGCAACAGGTATCGTTGTAGAACGTGAACGTTTAAACAAATATGGTACTCCTTTATTAGGTGCTACAGTAAAACCTAAATTAGGTTTATCTGGTAAAAACTACGGTCGTGTAGTATTTGAAGGTTTAAAAGGTGGTTTAGACTTCTTAAAAGATGATGAAAACATTAACTCACAACCATTCATGCGTTGGAGAGAGCG---

>*Navicula sp*

-------------CAATACTTCGCTTTCATCGCATACGAATGTGATTTATTTGAATAACTTAACAGCTTCTATTATTGGTAACGTTTTCGGATTTAAGGCAATTTCTGCTTTACGTTTAGAAGACATGCGTATTCCTCACTCATACTTAAAGACTTTCCAAGGTCCTGCGACAGGTGTTATCGTAGAACGTGAGCGTTTAAACAAATACGGTGTACCATTATTAGGTGCTACAGTAAAGCCTAAATTAGGTTTATCTGGTAAGAACTACGGTCGTGTAGTTTTCGAAGGTTTAAAAGGTGGTTTAGACTTCTTAAAAGATGATGAGAATATTAACTCACAACCATTCATGCGTTGGAGAGAGCG---

>*Astartiella sp.*

------------ATCATTTTTCGCTTTCATCGCATACGAATGTGATTTATTCGAATAACTTAACAGCGTCTATCATTGGTAACGTATTCGGTTTTAAAGCCGTAGCTGCTTTACGTTTAGAAGATATGCGTATTCCTCATTCTTACTTAAAAACATTCCAAGGTCCAGCTACAGGTGTAGTTGTAGAACGTGAACGTTTAAACAAGTATGGTGCACCATTATTAGGTGCAACAGTAAAACCTAAATTAGGTTTATCTGGTAAAAACTACGGTCGTGTAGTATATGAAGGTTTAAAAGGTGGTTTAGACTTCTTAAAAGATGATGAGAACATTAACTCTCAACCATTCATGCGTTGGAGAGAGCGA--

>*Amphiprora sp.*

-------ACTGATCAATACTTCGCATTTGTTGCTTACGAATGTGACTTATTCGAATAACATCACAGCATCTATTATTGGTAACGTATTCGGTTTCAAAGCTATCTCTGCGTTACGTTTAGAAGATATGCGTATTCCTCACTCTTACTTAAAAACTTTCCAAGGTCCAGCTACTGGTATCATTGTAGAGCGTGAGCGTATGAACAAATACGGTATCCCATTATTAGGTGCTACAGTAAAACCAAAATTAGGTTTATCTGGTAAAAACTACGGTCGTGTAGTATATGAAGGTTTAAAAGGTGGTTTAGACTTCTTAAAAGATGATGAAAACATTAACTCACAACCATTCATGCGTTGGAGAGAACG---

>*Diploneis sp.*

------------CAATAACTTCGCTTTTATCGCATACGAATGTGATTTATTCGAATAACTTAACTGCATCTATCATCGGTAATGTATTCGGTTTCAAAGCAGTATCTGCTTTACGTTTAGAAGATATGCGTATTCCTCACTCATACTTAAAAACATTCCAAGGTCCTGCTACAGGTGTTATCGTTGAACGTGAACGTTTAAACAAATATGGTGTTCCTTTATTAGGTGCTACAGTAAAACCAAAATTAGGCTTATCTGGTAAAAACTACGGTCGTGTAGTATACGAAGGTTTAAAAGGTGGTTTAGACTTCTTAAAAGATGATGAGAACATTAACTCTCAACCATTCATGCGTTGGAGAGAGCG---

>*Psammodictyon sp.*

---AAAACACTGATCATACTTCGCTTTCATCGCATACGAATGTGATTTATTTGAAAAACTTAACAGCTTCTATTATTGGTAACGTATTCGGTTTCAAAGCTGTAGCAGCGTTACGTTTAGAAGATATGCGTATTCCTCACTCATACTTAAAAACATTCCAAGGTCCTGCAACAGGTATCATTGTAGAGCGTGAGCGTTTAAACAAATACGGTGCTCCATTATTAGGTGCAACAGTAAAACCAAAATTAGGTTTATCTGGTAAAAACTATGGTCGTGTAGTATACGAAGGTTTAAAAGGTGGTTTAGACTTCTTAAAAGATGATGAAAACATTAACTCTCAACCATTCATGCGTTGGAGAGAGCG---

>*Coscinodiscus sp.*

----------------------------------------------------------------------------------------------------------------------AGAAGATATGCGTATTCCTCACACATACTTAAAAACATTCCAAGGTCCTGCAACAGGTATTATTGTAGAACGTGAACGTTTAAACAAATATGGTACTCCTTTATTAGGTGCAACTGTAAAACCTAAATTAGGTCTTTCTGGTAAAAACTATGGTCGTGTAGTTTATGAAGGTTTAAAAGGTGGTTTAGACTTCTTAAAAGATGATGAAAACATTAACTCTCAACCATTCATGCGTTGGAGAGAGCG---

>*Chaetoceros tenuissimus*

------ACTCCGAACATACTTTGCTTTCATCGCATACGAATGTGATTTATTCGAATAACTTAACTGCGTCTATCATTGGTAACGTATTTGGTTTCAAAGCAGTATCTGCTTTACGTTTAGAAGATATGCGTATTCCTCACTCATACTTAAAAACATTCCAAGGTCCTGCTACAGGTATCGTTGTAGAACGTGAACGTTTAAACAAATACGGTATCCCATTATTAGGTGCTACTGTAAAACCTAAGTTAGGTTTATCTGGTAAAAACTACGGTCGTGTAGTTTACGAAGGTTTAAAAGGTGGTTTAGACTTCTTAAAAGATGATGAGAACATTAACTCTCAACCATTCATGCGTTGGAGAGAGCG---

>*Hyalodiscus stelliger*

CCTAAACTNAGATCAATACTTTGCATTTATCGCTTATGAATGTGATTTATTCGAAGAATTTAACTGCATCAATCATTGGTAACGTATTTGGTTTCAAAGCAGTATCTGCTTTACGTTTAGAAGATATGCGTATCCCTCACTCATATTTAAAAACATTCCAAGGCCCTGCTACAGGTGTTATCGTAGAACGTGAACGTTTAAATAAATACGGTACTCCATTATTAGGTGCAACTGTAAAACCTAAATTAGGTCTTTCAGGTAAAAATTACGGTCGTGTAATTTATGAAGGTCTTAAAGGTGGTTTAGACTTCTTAAAAGATGATGAGAATATTAACTCACAACCATTTATGCGTTGGAGAGAGCGGTA

>*Cyclotella cryptica*

--------------------------------------------------------------------------------------TTTGGATTTAAAGCAGTTGCTGCTTTACGTTTAGAAGATATGCGTATTCCTCACTCATACTTAAAAACATTCCAAGGTCCTGCTACAGGTATTGTTGTAGAACGTGAACGTTTAAACAAATATGGTACTCCATTATTAGGTGCTACTGTAAAACCTAAATTAGGTCTTTCTGGTAAAAACTATGGTCGTGTAGTTTATGAAGGTTTAAAAGGTGGTTTAGACTTCTTAAAAGATGATGAAAACATTAACTCTCAACCATTTATGCGTTGGAGAGAACG---

>*Thalassiosira weissflogii*

-----------------------------------------------------------------------------------------------AAAGCAATCACTGCTTTACGTTTAGAAGATATGCGTATTCCTCACTCATACTTAAAAACATTCCAAGGTCCTGCTACAGGTATTATTGTAGAACGTGAACGTTTAAATAAATACGGTATTCCATTATTAGGTGCTACTGTAAAACCTAAATTAGGTCTTTCTGGTAAAAACTACGGTCGTGTAGTTTACGAAGGTTTAAAAGGTGGTTTAGACTTCTTAAAAGATGATGAAAACATTAACTCTCAACCATTTATGCGTTGGAGAGAGCG---

>*Thalassiosira rotula*

----------GATCAATACTTTGCATTTATCGCATACGAATGTGATTTATTTGAATAATTTAACAGCGTCTATTATTGGTAACGTTTTTGGATTTAAAGCAGTTGCTGCTTTACGTTTAGAAGATATGCGTATTCCTCACTCATATTTAAAAACATTCCAAGGTCCTGCTACAGGTATTATTGTAGAACGTGAACGTTTAAATAAATATGGTACACCATTATTAGGTGCAACTGTAAAACCTAAATTAGGTCTTTCTGGTAAAAACTATGGTCGTGTAGTTTATGAAGGTTTAAAAGGTGGTTTAGACTTCTTAAAAGATGATGAAAACATTAACTCTCAACCATTCATGCGTTGGAGAGAACG---

>*Minidiscus variabilis*

--------------------------------------------------------------------------------------TTTGGATTTAAAGCAATCTCTGCTTTACGTTTAGAAGATATGCGTATTCCTCACACATACTTAAAAACATTCCAAGGTCCTGCAACAGGTATTATTGTAGAACGTGAACGTTTAAACAAATATGGTACTCCTTTATTAGGTGCAACTGTAAAACCTAAATTAGGTCTTTCTGGTAAGAACTATGGTCGTGTAGTTTATGAAGGTTTAAAAGGTGGTTTAGACTTCTTAAAAGATGATGAAAACATTAACTCTCAACCATTCATGCGTTGGAGAGAGCG---

**Text S3.** Multiple alignment (fasta format) of phytoplankton *rbc*L sequences retrieved from GenBank used to design the diatom *rbc*L qRPA assay probe and primers.

>RPA *rbc*L Diatoms For

--------------------------------------------------CTGCTTTACGTTTAGAAGATATGCGTATTC-------------------------------------------------------------------------------------------------------------------------------------------------------------------------------------------------------------------------------------

>RPA *rbc*L Diatoms Probe

----------------------------------------------------------CGTTTAGAAGATATGCGTATTCCTCACTCATACTTAAAAACATTCCAAG----------------------------------------------------------------------------------------------------------------------------------------------------------------------------------------------------------

> RPA *rbc*L Diatoms Rev 1

-------------------------------------------------------------------------------------------------------------------------------------------------------------------------------------------------------------------------------------GAAGGTTTAAAAGGTGGWTTAGAYTTYTTA--------------------------------------------------

>MZ458597.1 Thalassiosira rotula isolate CNS00051 ribulose-1,5-bisphosphate carboxylase/oxygenase large subunit (rbcL) gene, complete cds; chloroplast

TAATTTAACAGCGTCTATTATTGGTAACGTTTTTGGATTTAAAGCAGTTGCTGCTTTACGTTTAGAAGATATGCGTATTCCTCACTCATATTTAAAAACATTCCAAGGTCCTGCTACAGGTATTATTGTAGAACGTGAACGTTTAAATAAATATGGTACACCATTATTAGGTGCAACTGTAAAACCTAAATTAGGTCTTTCTGGTAAAAACTATGGTCGTGTAGTTTATGAAGGTTTAAAAGGTGGTTTAGACTTCTTAAAAGATGATGAAAACATTAACTCTCAACCATTCATGCGTTGGAGAGAACG

>DQ514805.1 Thalassiosira rotula strain CCMP1812 ribulose-1,5-bisphosphate carboxylase/oxygenase large subunit (rbcL) gene, partial cds; chloroplast

TAATTTAACAGCGTCTATTATTGGTAACGTTTTTGGATTTAAAGCAGTTGCTGCTTTACGTTTAGAAGATATGCGTATTCCTCACTCATATTTAAAAACATTCCAAGGTCCTGCTACAGGTATTATTGTAGAACGTGAACGTTTAAATAAATATGGTACACCATTATTAGGTGCAACTGTAAAACCTAAATTAGGTCTTTCTGGTAAAAACTATGGTCGTGTAGTTTATGAAGGTTTAAAAGGTGGTTTAGACTTCTTAAAAGATGATGAAAACATTAACTCTCAACCATTCATGCGTTGGAGAGAACG

>DQ514799.1 Thalassiosira oceanica strain CCMP1001 ribulose-1,5-bisphosphate carboxylase/oxygenase large subunit (rbcL) gene, partial cds; chloroplast

TAACTTAACAGCATCTATTATTGGTAACGTTTTTGGATTTAAAGCAGTTTCTGCTTTACGTTTAGAAGATATGCGTATTCCTCACTCATATTTAAAAACATTCCAAGGTCCTGCGACAGGTATCATTGTAGAGCGTGAACGTTTAAACAAATATGGTACTCCATTATTAGGTGCAACTGTAAAACCAAAATTAGGTCTTTCTGGTAAAAACTACGGTCGTGTAGTTTATGAAGGTTTAAAAGGTGGTTTAGATTTCTTAAAAGATGATGAAAACATTAACTCTCAACCATTCATGCGTTGGAGAGAACG

>DQ514811.1 Thalassiosira weissflogii strain L1296 ribulose-1,5-bisphosphate carboxylase/oxygenase large subunit (rbcL) gene, partial cds; chloroplast

TAACTTAACAGCATCTATTATTGGTAACGTTTTTGGGTTTAAAGCAATTTCTGCTTTACGTTTAGAAGATATGCGTATTCCTCACTCATACTTAAAAACATTCCAAGGTCCTGCTACAGGTATTATTGTAGAACGTGAACGTTTAAATAAATACGGTATTCCATTATTAGGTGCTACTGTAAAACCTAAATTAGGTCTTTCTGGTAAAAACTACGGTCGTGTAGTTTACGAAGGTTTAAAAGGTGGTTTAGACTTCTTAAAAGATGATGAAAACATTAACTCTCAACCATTTATGCGTTGGAGAGAGCG

>DQ514794.1 Thalassiosira aestivalis strain CCMP976 ribulose-1,5-bisphosphate carboxylase/oxygenase large subunit (rbcL) gene, partial cds; chloroplast

TAATTTAACAGCGTCTATTATTGGTAACGTTTTTGGATTTAAAGCAATTTCTGCTTTACGTTTAGAAGATATGCGTATTCCTCACTCATACTTAAAAACATTCCAAGGTCCTGCTACAGGTATCATTGTAGAACGTGAACGTTTAAACAAATATGGTACTCCATTATTAGGTGCTACTGTAAAACCTAAATTAGGTCTTTCTGGTAAAAACTACGGTCGTGTAGTTTATGAAGGTTTAAAAGGTGGTTTAGACTTCTTAAAAGATGATGAAAACATTAACTCTCAACCATTCATGCGTTGGAGAGAACG

>DQ514797.1 Thalassiosira minima strain CCMP990 ribulose-1,5-bisphosphate carboxylase/oxygenase large subunit (rbcL) gene, partial cds; chloroplast

TAACTTAACAGCGTCTATTATTGGTAACGTATTTGGTTTCAAAGCAATCTCTGCTTTACGTTTAGAAGATATGCGTATTCCTCACTCATACTTAAAAACATTCCAAGGTCCTGCTACAGGTATCGTTGTAGAACGTGAACGTTTAAACAAATATGGTACTCCATTATTAGGTGCAACTGTAAAACCTAAATTAGGTTTATCTGGTAAAAACTACGGTCGTGTAGTTTACGAAGGTTTAAAAGGTGGTTTAGACTTCTTAAAAGATGATGAAAACATTAACTCTCAACCATTCATGCGTTGGAGAGAGCG

>DQ514784.1 Thalassiosira pseudonana strain NEPC709 ribulose-1,5-bisphosphate carboxylase/oxygenase large subunit (rbcL) gene, partial cds; chloroplast

TAACTTAACAGCTTCTATCATCGGTAACGTTTTTGGATTCAAAGCAATTTCTGCTTTACGTTTAGAAGATATGCGTATTCCTCACTCATATTTAAAAACATTCCAAGGTCCTGCTACAGGTATCGTTGTAGAACGTGAACGTTTAAACAAATATGGTACTCCATTATTAGGTGCTACTGTAAAACCTAAATTAGGTCTTTCTGGTAAAAACTATGGTCGTGTAGTTTATGAAGGTTTAAAAGGTGGTTTAGACTTCTTAAAAGATGATGAAAACATTAACTCTCAACCATTCATGCGTTGGAGAGAGCG

>DQ514806.1 Thalassiosira mediterranea strain CS16 ribulose-1,5-bisphosphate carboxylase/oxygenase large subunit (rbcL) gene, partial cds; chloroplast

TAACTTAACAGCGTCTATTATTGGTAACGTTTTTGGTTTTAAAGCAGTTGCTGCTTTACGTTTAGAAGATATGCGTATTCCTCACTCATACTTAAAAACATTCCAAGGTCCTGCTACAGGTATCGTTGTAGAACGTGAACGTTTAAACAAATATGGTACTCCATTATTAGGTGCAACTGTAAAACCTAAATTAGGTTTATCTGGTAAAAACTATGGTCGTGTAGTTTATGAAGGTTTAAAAGGTGGTTTAGACTTCTTAAAAGATGATGAAAACATTAACTCTCAACCATTCATGCGTTGGAGAGAACG

>DQ514807.1 Thalassiosira punctigera strain FB02-06 ribulose-1,5-bisphosphate carboxylase/oxygenase large subunit (rbcL) gene, partial cds; chloroplast

TAATTTAACAGCATCTATTATCGGTAACGTTTTTGGATTTAAAGCAGTTGCTGCTTTACGTTTAGAAGATATGCGTATTCCTCACTCATATTTAAAAACATTCCAAGGTCCTGCTACAGGTATTATTGTAGAACGTGAACGTTTAAACAAATACGGTACTCCATTATTAGGTGCAACTGTAAAACCTAAATTAGGTCTTTCTGGTAAAAACTATGGTCGTGTAGTTTATGAAGGTTTAAAAGGTGGTTTAGACTTCTTAAAAGATGATGAAAACATTAACTCTCAACCATTCATGCGTTGGAGAGAACG

>DQ514809.1 Thalassiosira minuscula strain FB02-31 ribulose-1,5-bisphosphate carboxylase/oxygenase large subunit (rbcL) gene, partial cds; chloroplast

TAACTTAACAGCGTCTATTATTGGTAACGTTTTCGGATTTAAAGCAATTTCTGCTTTACGTTTAGAAGATATGCGTATTCCTCACTCATACTTAAAAACATTCCAAGGTCCTGCTACAGGTATCATTGTAGAACGTGAACGTTTAAACAAATATGGTACACCATTATTAGGTGCAACTGTAAAACCTAAATTAGGTCTTTCTGGTAAAAACTATGGTCGTGTAGTTTATGAAGGTTTAAAAGGTGGTTTAGACTTCTTAAAAGATGATGAAAACATTAACTCTCAACCATTCATGCGTTGGAGAGAACG

>DQ514783.1 Thalassiosira pseudonana strain ETC1 ribulose-1,5-bisphosphate carboxylase/oxygenase large subunit (rbcL) gene, partial cds; chloroplast

TAACTTAACAGCTTCTATCATCGGTAACGTTTTTGGATTCAAAGCAATTTCTGCTTTACGTTTAGAAGATATGCGTATTCCTCACTCATATTTAAAAACATTCCAAGGTCCTGCTACAGGTATCGTTGTAGAACGTGAACGTTTAAACAAATATGGTACTCCATTATTAGGTGCTACTGTAAAACCTAAATTAGGTCTTTCTGGTAAAAACTATGGTCGTGTAGTTTATGAAGGTTTAAAAGGTGGTTTAGACTTCTTAAAAGATGATGAAAACATTAACTCTCAACCATTCATGCGTTGGAGAGAGCG

>DQ514801.1 Thalassiosira pseudonana strain CCMP1057 ribulose-1,5-bisphosphate carboxylase/oxygenase large subunit (rbcL) gene, partial cds; chloroplast

TAACTTAACAGCTTCTATCATCGGTAACGTTTTTGGATTCAAAGCAATTTCTGCTTTACGTTTAGAAGATATGCGTATTCCTCACTCATATTTAAAAACATTCCAAGGTCCTGCTACAGGTATCGTTGTAGAACGTGAACGTTTAAACAAATATGGTACTCCATTATTAGGTGCTACTGTAAAACCTAAATTAGGTCTTTCTGGTAAAAACTATGGTCGTGTAGTTTATGAAGGTTTAAAAGGTGGTTTAGACTTCTTAAAAGATGATGAAAACATTAACTCTCAACCATTCATGCGTTGGAGAGAGCG

>DQ514800.1 Thalassiosira weissflogii strain CCMP1010 ribulose-1,5-bisphosphate carboxylase/oxygenase large subunit (rbcL) gene, partial cds; chloroplast

TAACTTAACAGCATCTATTATTGGTAACGTTTTTGGGTTTAAAGCAATTTCTGCTTTACGTTTAGAAGATATGCGTATTCCTCACTCATACTTAAAAACATTCCAAGGTCCTGCTACAGGTATTATTGTAGAACGTGAACGTTTAAATAAATACGGTATTCCATTATTAGGTGCTACTGTAAAACCTAAATTAGGTCTTTCTGGTAAAAACTACGGTCGTGTAGTTTACGAAGGTTTAAAAGGTGGTTTAGACTTCTTAAAAGATGATGAAAACATTAACTCTCAACCATTTATGCGTTGGAGAGAGCG

>MZ458599.1 Thalassiosira tenera isolate CNS00472 ribulose-1,5-bisphosphate carboxylase/oxygenase large subunit (rbcL) gene, complete cds; chloroplast

TAACTTAACAGCGTCTATTATTGGTAACGTTTTTGGATTTAAAGCAATTTCTGCTTTACGTTTAGAAGATATGCGTATTCCTCACTCATACTTAAAAACATTCCAAGGTCCTGCGACAGGTATTATTGTAGAACGTGAACGTTTAAACAAATATGGTACTCCATTATTAGGTGCAACTGTAAAACCTAAATTAGGTTTATCTGGTAAAAACTATGGTCGTGTAGTTTATGAAGGTTTAAAAGGTGGTTTAGACTTCTTAAAAGATGATGAAAACATTAACTCTCAACCATTCATGCGTTGGAGAGAACG

>DQ514803.1 Thalassiosira minuscula strain CCMP1093 ribulose-1,5-bisphosphate carboxylase/oxygenase large subunit (rbcL) gene, partial cds; chloroplast

TAACTTAACAGCGTCTATTATTGGTAACGTTTTCGGATTTAAAGCAATTTCTGCTTTACGTTTAGAAGATATGCGTATTCCTCACTCATACTTAAAAACATTCCAAGGTCCTGCTACAGGTATCATTGTAGAACGTGAACGTTTAAACAAATATGGTACACCATTATTAGGTGCAACTGTAAAACCTAAATTAGGTCTTTCTGGTAAAAACTATGGTCGTGTAGTTTATGAAGGTTTAAAAGGTGGTTTAGACTTCTTAAAAGATGATGAAAACATTAACTCTCAACCATTCATGCGTTGGAGAGAACG

>MW478286.1 Thalassiosira profunda strain CNS00050 ribulose-1,5-bisphosphate carboxylase/oxygenase large subunit (rbcL) gene, complete cds; chloroplast

TAACTTAACAGCGTCTATCATTGGTAACGTTTTCGGATTTAAAGCAATTTCTGCTTTACGTTTAGAAGATATGCGTATTCCTCACTCATATTTAAAAACATTCCAAGGTCCTGCTACAGGTATCATTGTAGAACGTGAACGTTTAAACAAATATGGTGTTCCATTATTAGGTGCAACTGTAAAACCTAAATTAGGTCTTTCTGGTAAAAACTATGGTCGTGTAGTTTATGAAGGTTTAAAAGGTGGTTTAGACTTCTTAAAAGATGATGAAAACATTAACTCTCAACCATTCATGCGTTGGAGAGAACG

>KY320298.1 Navicula incertata strain TA414 ribulose-1,5-bisphosphate carboxylase/oxygenase large subunit (rbcL) gene, complete cds; chloroplast

TAACTTAACAGCTTCGATCATTGGTAACGTATTCGGATTTAAAGCAATTTCTGCTTTACGTTTAGAAGATATGCGTATTCCTCACTCATACTTAAAAACTTTCCAAGGTCCAGCTACAGGTATCATTGTAGAACGTGAGCGTTTAAACAAATATGGTATTCCATTATTAGGTGCTACTGTAAAACCTAAATTAGGTTTATCTGGTAAAAACTACGGTCGTGTAGTTTTCGAAGGTTTAAAAGGTGGTTTAGACTTCTTAAAAGATGATGAAAACATTAACTCACAACCATTCATGCGTTGGAGAGAACG

>MW848495.1 Chaetoceros socialis ribulose-1,5-bisphosphate carboxylase/oxygenase large subunit (rbcl) gene, complete cds; chloroplast

TAACTTAACTGCGTCTATCATTGGTAACGTATTTGGTTTTAAAGCGGTTGCTGCTTTACGTTTAGAAGATATGCGTATTCCTCACTCATACTTAAAAACATTCCAAGGTCCTGCTACAGGTATCGTTGTAGAACGTGAACGTTTAAACAAATATGGTGTTCCTTTATTAGGTGCTACTGTAAAACCTAAATTAGGTTTATCTGGTAAAAACTACGGTCGTGTAGTTTTCGAAGGTTTAAAAGGTGGTTTAGACTTCTTAAAAGATGATGAGAACATTAACTCTCAACCATTCATGCGTTGGAGAGAGCG

>MK331991.1 Chaetoceros tenuissimus ribulose-1,5-bisphosphate carboxylase/oxygenase large subunit gene, partial cds; chloroplast

TAACTTAACTGCGTCTATCATTGGTAACGTATTTGGTTTCAAAGCAGTATCTGCTTTACGTTTAGAAGATATGCGTATTCCTCACTCATACTTAAAAACATTCCAAGGTCCTGCTACAGGTATCGTTGTAGAACGTGAGCGTTTAAACAAATACGGTATCCCATTATTAGGTGCTACTGTAAAACCTAAATTAGGTTTATCTGGTAAAAACTACGGTCGTGTAGTTTATGAAGGTTTAAAAGGTGGTTTAGACTTCTTAAAAGATGATGAGAACATTAACTCTCAACCATTCATGCGTTGGAGAGAACG

>MW848497.1 Chaetoceros tenuissimus ribulose-1,5-bisphosphate carboxylase/oxygenase large subunit (rbcl) gene, complete cds; chloroplast

TAACTTAACTGCGTCTATCATTGGTAACGTATTTGGTTTCAAAGCAGTATCTGCTTTACGTTTAGAAGATATGCGTATTCCTCACTCATACTTAAAAACATTCCAAGGTCCTGCTACAGGTATCGTTGTAGAACGTGAACGTTTAAACAAATACGGTATCCCATTATTAGGTGCTACTGTAAAACCTAAGTTAGGTTTATCTGGTAAAAACTACGGTCGTGTAGTTTACGAAGGTTTAAAAGGTGGTTTAGACTTCTTAAAAGATGATGAGAACATTAACTCTCAACCATTCATGCGTTGGAGAGAGCG

>KY751726.1 Phaeodactylum tricornutum strain 1 ribulose-1,5-bisphosphate carboxylase/oxygenase large subunit (rbcL) gene, partial cds; chloroplast

TAACTTAACAGCATCTATTATTGGTAACGTTTTTGGGTTTAAAGCAATTTCTGCTTTACGTTTAGAAGATATGCGTATTCCTCACTCATACTTAAAAACATTCCAAGGTCCTGCTACAGGTATTATTGTAGAACGTGAACGTTTAAATAAATACGGTATTCCATTATTAGGTGCTACTGTAAAACCTAAATTAGGTCTTTCTGGTAAAAACTACGGTCGTGTAGTTTACGAAGGTTTAAAAGGTGGTTTAGACTTCTTAAAAGATGATGAAAACATTAACTCTCAACCATTTATGCGTTGGAGAGAGCG

>DQ514766.1 Ditylum brightwellii strain CCMP1810 ribulose-1,5-bisphosphate carboxylase/oxygenase large subunit (rbcL) gene, partial cds; chloroplast

AAATTTAACTGCATCAATCATTGGTAACGTTTTCGGATTTAAAGCAATTTCTGCTTTACGTTTAGAAGATATGCGTATTCCTCACTCATACTTAAAAACATTCCAAGGTCCTGCGACAGGTATTATTGTAGAACGTGAACGTTTAAACAAATACGGTATTCCTTTATTAGGTGCTACTGTAAAACCTAAATTAGGTCTTTCTGGTAAAAACTACGGTCGTGTAGTATATGAAGGTTTAAAAGGTGGTTTAGACTTCTTAAAAGATGATGAAAACATTAACTCACAACCATTCATGCGTTGGAGAGAGCG

>HQ656824.1 Ditylum brightwellii culture-collection PCC:609 ribulose-1,5-bisphosphate carboxylase/oxygenase large subunit (rbcL) gene, partial cds; chloroplast

AAATTTAACTGCATCAATCATTGGTAACGTTTTCGGATTTAAAGCAATTTCTGCTTTACGTTTAGAAGATATGCGTATTCCTCACTCATACTTAAAAACATTCCAAGGTCCTGCGACAGGTATTATTGTAGAACGTGAACGTTTAAACAAATACGGTATTCCTTTATTAGGTGCTACTGTAAAACCTAAATTAGGTCTTTCTGGTAAAAACTACGGTCGTGTAGTATATGAAGGTTTAAAAGGTGGTTTAGACTTCTTAAAAGATGATGAAAACATTAACTCACAACCATTCATGCGTTGGAGAGAGCG

>KC309566.1 Ditylum brightwellii isolate ECT3884Ditylum ribulose-1,5-bisphosphate carboxylase/oxygenase large subunit (rbcL) gene, partial cds; chloroplast

AAATTTAACTGCATCAATCATTGGTAACGTTTTCGGATTTAAAGCAATTTCTGCTTTACGTTTAGAAGATATGCGTATTCCTCACTCATACTTAAAAACATTCCAAGGTCCTGCGACAGGTATTATTGTAGAACGTGAACGTTTAAACAAATACGGTATTCCTTTATTAGGTGCTACTGTAAAACCTAAATTAGGTCTTTCTGGTAAAAACTACGGTCGTGTAGTATATGAAGGTTTAAAAGGTGGTTTAGACTTCTTAAAAGATGATGAAAACATTAACTCACAACCATTCATGCGTTGGAGAGAGCG

>HQ912519.1 Lithodesmioides polymorpha strain ECT3772-Lithodesmioides ribulose-1,5-bisphosphate carboxylase/oxygenase large subunit (rbcL) gene, partial cds; chloroplast

AAACTTAACTGCATCTATTATTGGTAACGTTTTTGGATTTAAAGCAGTAGCTGCTTTACGTTTAGAAGATATGCGTATTCCTCACTCATACTTAAAAACATTCCAAGGTCCTGCTACAGGTATCGTTGTAGAGCGTGAACGTTTAAACAAATATGGTATTCCTTTATTAGGTGCTACTGTAAAACCTAAATTAGGTCTTTCTGGTAAAAACTACGGTCGTGTAGTATACGAAGGTTTAAAAGGTGGTTTAGACTTCTTAAAAGATGATGAAAACATTAACTCACAACCATTCATGCGTTGGAGAGAGCG

>DQ514764.1 Helicotheca tamesis strain CCMP1760 ribulose-1,5-bisphosphate carboxylase/oxygenase large subunit (rbcL) gene, partial cds; chloroplast

AAACTTAACTGCATCTATTATTGGTAACGTTTTTGGATTTAAAGCAATCTCTGCTTTACGTTTAGAAGATATGCGTATTCCTCACTCATACTTAAAAACATTCCAAGGTCCTGCTACAGGTATCGTTGTAGAACGTGAACGTTTAAACAAATATGGTATTCCTTTATTAGGTGCTACTGTAAAACCGAAATTAGGTCTTTCTGGTAAAAACTACGGTCGTGTAGTATATGAAGGTTTAAAAGGTGGTTTAGACTTCTTAAAAGATGATGAAAACATTAACTCACAACCATTCATGCGTTGGAGAGAGCG

>HQ912423.1 Lithodesmium undulatum strain CCMP1797 ribulose-1,5-bisphosphate carboxylase/oxygenase large subunit (rbcL) gene, complete cds; chloroplast

AAACTTAACTGCATCTATTATTGGTAACGTTTTTGGATTTAAAGCAGTAGCTGCTTTACGTTTAGAAGATATGCGTATTCCTCACTCATACTTAAAAACATTCCAAGGTCCTGCTACAGGTATCGTTGTAGAGCGTGAACGTTTAAACAAATATGGTATTCCTTTATTAGGTGCTACTGTAAAACCTAAATTAGGTCTTTCAGGTAAAAACTACGGTCGTGTAGTATATGAAGGTTTAAAAGGTGGTTTAGACTTCTTAAAAGATGATGAAAACATTAACTCACAACCATTCATGCGTTGGAGAGAACG

>HQ912536.1 Bellerochea horologicalis strain ECT3829-Bellerochea ribulose-1,5-bisphosphate carboxylase/oxygenase large subunit (rbcL) gene, complete cds; chloroplast

AAACTTAACTGCATCTATTATTGGTAACGTTTTCGGATTTAAAGCAATCTCTGCTTTACGTTTAGAAGATATGCGTATTCCTCACTCATACTTAAAAACATTCCAAGGTCCTGCTACAGGTATCGTTGTAGAGCGTGAACGTTTAAACAAATATGGTATTCCTTTATTAGGTGCTACTGTAAAACCTAAATTAGGTCTTTCTGGTAAAAACTATGGTCGTGTAGTATACGAAGGTTTAAAAGGTGGTTTAGACTTCTTAAAAGATGATGAAAACATCAACTCACAACCATTCATGCGTTGGAGAGAACG

>DQ514765.1 Lithodesmium undulatum strain CCMP1806 ribulose-1,5-bisphosphate carboxylase/oxygenase large subunit (rbcL) gene, partial cds; chloroplast

AAACTTAACTGCATCTATTATTGGTAACGTTTTTGGATTTAAAGCAGTAGCTGCTTTACGTTTAGAAGATATGCGTATTCCTCACTCATACTTAAAAACATTCCAAGGTCCTGCTACAGGTATCGTTGTAGAGCGTGAACGTTTAAACAAATATGGTATTCCTTTATTAGGTGCTACTGTAAAACCTAAATTAGGTCTTTCAGGTAAAAACTACGGTCGTGTAGTATATGAAGGTTTAAAAGGTGGTTTAGACTTCTTAAAAGATGATGAAAACATTAACTCACAACCATTCATGCGTTGGAGAGAACG

>KC309565.1 Ditylum sol isolate Har-1Ditylum ribulose-1,5-bisphosphate carboxylase/oxygenase large subunit (rbcL) gene, partial cds; chloroplast

AAACTTAACTGCATCAATCATTGGTAACGTTTTCGGATTTAAAGCAGTAGCTGCTTTACGTTTAGAAGATATGCGTATTCCTCACTCATACTTAAAAACATTCCAAGGTCCTGCTACAGGTATTGTTGTAGAGCGTGAACGTTTAAACAAATACGGTATTCCTTTATTAGGTGCTACTGTAAAACCTAAATTAGGTCTTTCTGGTAAAAACTACGGTCGTGTAGTATATGAAGGTTTAAAAGGTGGTTTAGACTTCTTAAAAGATGATGAAAACATTAACTCACAACCATTCATGCGTTGGAGAGAGCG

>KJ577895.1 Helicotheca tamesis isolate 25VI12-2AHelico ribulose-1,5-bisphosphate carboxylase/oxygenase large subunit (rbcL) gene, partial cds; chloroplast

AAACTTAACTGCATCTATTATTGGTAACGTTTTTGGATTTAAAGCAGTAGCTGCTTTACGTTTAGAAGATATGCGTATTCCTCACTCATACTTAAAAACATTCCAAGGTCCTGCTACAGGTATCATCGTAGAACGTGAACGTTTAAACAAATATGGTATTCCTTTATTAGGTGCTACTGTAAAACCTAAATTAGGTCTTTCTGGTAAAAACTACGGTCGTGTAGTATACGAAGGTTTAAAAGGTGGTTTAGACTTCTTAAAAGATGATGAAAACATTAACTCACAACCATTCATGCGTTGGAGAGAGCG

>MK817351.1 Lithodesmium intricatum strain Azo12Litho-J2 ribulose 1,5 bisphosphate carboxylase/oxygenase large subunit (rbcL) gene, partial cds; chloroplast

GAACTTAACTGCATCTATCATTGGTAACGTTTTCGGATTTAAAGCAGTAGCTGCTTTACGTTTAGAAGATATGCGTATTCCTCACTCATACTTAAAAACATTCCAAGGTCCTGCTACAGGTGTCGTTGTAGAGCGTGAACGTTTAAACAAATATGGTATTCCTTTATTAGGTGCTACTGTAAAACCTAAATTAGGTCTTTCAGGTAAAAACTACGGTCGTGTAGTATATGAAGGTTTAAAAGGTGGTTTAGACTTCTTAAAAGATGATGAAAACATTAACTCACAACCATTCATGCGTTGGAGAGAGCG

>MK817350.1 Lithodesmium intricatum strain Azo12Litho-5 ribulose 1,5 bisphosphate carboxylase/oxygenase large subunit (rbcL) gene, partial cds; chloroplast

GAACTTAACTGCATCTATCATTGGTAACGTTTTCGGATTTAAAGCAGTAGCTGCTTTACGTTTAGAAGATATGCGTATTCCTCACTCATACTTAAAAACATTCCAAGGTCCTGCTACAGGTGTCGTTGTAGAGCGTGAACGTTTAAACAAATATGGTATTCCTTTATTAGGTGCTACTGTAAAACCTAAATTAGGTCTTTCAGGTAAAAACTACGGTCGTGTAGTATATGAAGGTTTAAAAGGTGGTTTAGACTTCTTAAAAGATGATGAAAACATTAACTCACAACCATTCATGCGTTGGAGAGAGCG

>MK817349.1 Lithodesmium intricatum strain 25VI12-1ALitho ribulose 1,5 bisphosphate carboxylase/oxygenase large subunit (rbcL) gene, partial cds; chloroplast

GAACTTAACTGCATCTATCATTGGTAACGTTTTCGGATTTAAAGCAGTAGCTGCTTTACGTTTAGAAGATATGCGTATTCCTCACTCATACTTAAAAACATTCCAAGGTCCTGCTACAGGTGTCGTTGTAGAGCGTGAACGTTTAAACAAATATGGTATTCCTTTATTAGGTGCTACTGTAAAACCTAAATTAGGTCTTTCAGGTAAAAACTACGGTCGTGTAGTATATGAAGGTTTAAAAGGTGGTTTAGACTTCTTAAAAGATGATGAAAACATTAACTCACAACCATTCATGCGTTGGAGAGAGCG

>HQ912542.1 Lithodesmium intricatum strain ECT3850 ribulose-1,5-bisphosphate carboxylase/oxygenase large subunit (rbcL) gene, partial cds; chloroplast

GAACTTAACTGCATCTATCATTGGTAACGTTTTCGGATTTAAAGCAGTAGCTGCTTTACGTTTAGAAGATATGCGTATTCCTCACTCATACTTAAAAACATTCCAAGGTCCTGCTACAGGTATCGTTGTAGAGCGTGAACGTTTAAACAAATATGGTATTCCTTTATTAGGTGCTACTGTAAAACCTAAATTAGGTCTTTCAGGTAAAAACTACGGTCGTGTAGTATATGAAGGTTTAAAAGGTGGTTTAGACTTCTTAAAAGATGATGAAAACATTAACTCACAACCATTCATGCGTTGGAGAGAGCG

>KC309557.1 Bellerochea malleus isolate Har-1Bellmall ribulose-1,5-bisphosphate carboxylase/oxygenase large subunit (rbcL) gene, partial cds; chloroplast

AAACTTAACTGCATCTATTATTGGTAACGTTTTCGGATTTAAAGCAATCTCTGCTTTACGTTTAGAAGATATGCGTATTCCTCACTCATACTTAAAAACATTCCAAGGTCCTGCTACAGGTATCGTTGTAGAGCGTGAACGTTTAAACAAATATGGTATTCCTTTATTAGGTGCTACTGTAAAACCTAAATTAGGTCTTTCTGGTAAAAACTACGGTCGTGTAGTATACGAAGGTTTAAAAGGTGGTTTAGACTTCTTAAAAGATGATGAAAACATCAACTCACAACCATTCATGCGTTGGAGAGAACG

>HQ912534.1 Lithodesmium intricatum strain ECT3836-Lithodesmium ribulose-1,5-bisphosphate carboxylase/oxygenase large subunit (rbcL) gene, partial cds; chloroplast

GAACTTAACTGCATCTATCATTGGTAACGTTTTCGGATTTAAAGCAGTAGCTGCTTTACGTTTAGAAGATATGCGTATTCCTCACTCATACTTAAAAACATTCCAAGGTCCTGCTACAGGTATCGTTGTAGAGCGTGAACGTTTAAACAAATATGGTATTCCTTTATTAGGTGCTACTGTAAAACCTAAATTAGGTCTTTCAGGTAAAAACTACGGTCGTGTAGTATATGAAGGTTTAAAAGGTGGTTTAGACTTCTTAAAAGATGATGAAAACATTAACTCACAACCATTCATGCGTTGGAGAGAGCG

>FJ002135.1 Helicotheca tamesis isolate C60 ribulose-1,5-bisphosphate carboxylase/oxygenase large subunit gene, partial cds; chloroplast

AAACTTAACTGCATCTATTATTGGTAACGTTTTTGGATTTAAAGCAATCTCTGCTTTACGTTTAGAAGATATGCGTATTCCTCACTCATACTTAAAAACATTCCAAGGTCCTGCTACAGGTATCGTTGTAGAACGTGAACGTTTAAACAAATATGGTATTCCTTTATTAGGTGCTACTGTAAAACCGAAATTAGGTCTTTCTGGTAAAAACTACGGTCGTGTAGTATATGAAGGTTTAAAAGGTGGTTTAGACTTCTTAAAAGATGATGAAAACATTAACTCACAACCATTCATGCGTTGGAGAGAGCG

>MZ458600.1 Conticribra weissflogii isolate CNS00439 ribulose-1,5-bisphosphate carboxylase/oxygenase large subunit (rbcL) gene, complete cds; chloroplast

TAACTTAACAGCATCTATTATTGGTAACGTTTTTGGGTTTAAAGCAGTTGCTGCTTTACGTTTAGAAGATATGCGTATTCCTCACTCATACTTAAAAACATTCCAAGGTCCTGCTACAGGTATTATTGTAGAACGTGAACGTTTAAATAAATACGGTATTCCATTATTAGGTGCTACTGTAAAACCTAAATTAGGTCTTTCTGGTAAAAACTACGGTCGTGTAGTTTATGAAGGTTTAAAAGGTGGTTTAGACTTCTTAAAAGATGATGAAAACATTAACTCTCAACCATTCATGCGTTGGAGAGAGCG

>MK817348.1 Lithodesmium intricatum strain ECT3850Litho ribulose 1,5 bisphosphate carboxylase/oxygenase large subunit (rbcL) gene, partial cds; chloroplast

GAACTTAACTGCATCTATCATTGGTAACGTTTTCGGATTTAAAGCAGTAGCTGCTTTACGTTTAGAAGATATGCGTATTCCTCACTCATACTTAAAAACATTCCAAGGTCCTGCTACAGGTATCGTTGTAGAGCGTGAACGTTTAAACAAATATGGTATTCCTTTATTAGGTGCTACTGTAAAACCTAAATTAGGTCTTTCAGGTAAAAACTACGGTCGTGTAGTATATGAAGGTTTAAAAGGTGGTTTAGACTTCTTAAAAGATGATGAAAACATTAACTCACAACCATTCATGCGTTGGAGAGAGCG

>MK817347.1 Lithodesmium intricatum strain ECT3836Litho ribulose 1,5 bisphosphate carboxylase/oxygenase large subunit (rbcL) gene, partial cds; chloroplast

GAACTTAACTGCATCTATCATTGGTAACGTTTTCGGATTTAAAGCAGTAGCTGCTTTACGTTTAGAAGATATGCGTATTCCTCACTCATACTTAAAAACATTCCAAGGTCCTGCTACAGGTATCGTTGTAGAGCGTGAACGTTTAAACAAATATGGTATTCCTTTATTAGGTGCTACTGTAAAACCTAAATTAGGTCTTTCAGGTAAAAACTACGGTCGTGTAGTATATGAAGGTTTAAAAGGTGGTTTAGACTTCTTAAAAGATGATGAAAACATTAACTCACAACCATTCATGCGTTGGAGAGAGCG

>MF001952.1 Cymatosira belgica isolate s0289 ribulose-1,5-bisphosphate carboxylase/oxygenase large subunit (rbcL) gene, complete cds; chloroplast

TAACTTAACTGCATCTATCATCGGTAACGTATTCGGATTTAAAGCAGTAGCTGCTTTACGTTTAGAAGATATGCGTATTCCTCACTCATACTTAAAAACATTCCAAGGTCCGGCTACAGGTATCGTTGTAGAGCGTGAGCGTTTAAACAAATATGGTACTCCATTATTAGGTGCTACTGTAAAACCTAAATTAGGTCTTTCTGGTAAAAACTACGGTCGTGTAGTATACGAAGGTTTAAAAGGTGGTTTAGACTTCTTAAAAGATGATGAAAACATTAACTCTCAACCATTCATGCGTTGGAGAGAACG

>HQ912487.1 Campylosira cymbelliformis strain CCC-1 ribulose-1,5-bisphosphate carboxylase/oxygenase large subunit (rbcL) gene, complete cds; chloroplast

TAACTTAACTGCATCTATCATCGGTAACGTATTCGGATTTAAAGCAGTATCTGCTTTACGTTTAGAAGATATGCGTATTCCTCACTCATACTTAAAAACATTCCAAGGTCCGGCTACAGGTATTGTTGTAGAGCGTGAGCGTTTAAACAAATACGGTACTCCATTATTAGGTGCTACTGTAAAACCTAAATTAGGTCTTTCTGGTAAAAACTACGGTCGTGTAGTATATGAAGGTTTAAAAGGTGGTTTAGACTTCTTAAAGGATGATGAAAACATTAACTCTCAACCATTCATGCGTTGGAGAGAACG

>MK454991.1 Bellerochea malleus isolate SZCZE468 ribulose-1,5-bisphosphate carboxylase/oxygenase large subunit (rbcL) gene, partial cds; chloroplast

AAACTTAACTGCATCTATCATTGGTAACGTTTTCGGATTTAAAGCAATCTCTGCTTTACGTTTAGAAGATATGCGTATTCCTCACTCATACTTAAAAACATTCCAAGGTCCTGCTACAGGTATCGTTGTAGAGCGTGAACGTTTAAACAAATATGGTATTCCTTTATTAGGTGCTACTGTAAAACCTAAATTAGGTCTTTCTGGTAAAAACTACGGTCGTGTAGTATACGAAGGTTTAAAAGGTGGTTTAGACTTCTTAAAAGATGATGAAAACATCAACTCACAACCATTCATGCGTTGGAGAGAACG

>HQ912428.1 Odontella sinensis strain CCMP1815 ribulose-1,5-bisphosphate carboxylase/oxygenase large subunit (rbcL) gene, complete cds; chloroplast

TAACTTAACTGCATCAATCATTGGTAACGTATTCGGATTTAAAGCGGTAGCTGCTTTACGTTTAGAAGATATGCGTATTCCTTATGCATACTTAAAAACATTCCAAGGTCCTGCTACAGGTATCGTTGTTGAACGTGAGCGTTTAAACAAATATGGTGCACCATTATTAGGTGCTACTGTAAAACCTAAATTAGGTCTTTCTGGTAAAAACTACGGTCGTGTAGTATATGAAGGTTTAAAAGGTGGTTTAGACTTTTTAAAAGATGATGAAAATATTAACTCACAACCATTCATGCGTTGGAGAGAACG

>JX413566.1 Odontella mobiliensis isolate ECT3829Odmob ribulose-1,5-bisphosphate carboxylase/oxygenase large subunit (rbcL) gene, partial cds; chloroplast

GAATTTAACTGCGTCTATCATTGGTAACGTATTTGGATTTAAAGCGGTTGCTGCTTTACGTTTAGAAGATATGCGTATTCCTTATGCATACTTAAAAACATTCCAAGGTCCTGCTACAGGTATCGTTGTTGAACGTGAGCGTTTAAACAAATATGGTACTCCATTATTAGGTGCTACTGTAAAACCTAAATTAGGTCTTTCTGGTAAAAACTACGGTCGTGTAGTATACGAAGGTTTAAAAGGTGGTTTAGACTTCTTAAAAGATGATGAAAACATTAACTCACAACCATTCATGCGTTGGAGAGAACG

>DQ514821.1 Skeletonema menzellii strain CCMP787 ribulose-1,5-bisphosphate carboxylase/oxygenase large subunit (rbcL) gene, partial cds; chloroplast

TAACTTAACAGCATCTATTATTGGTAACGTTTTTGGATTTAAAGCAGTTGCTGCTTTACGTTTAGAAGATATGCGTATTCCTCACTCATACTTAAAAACATTCCAAGGTCCTGCTACAGGTATCGTTGTAGAACGTGAACGTTTAAACAAATATGGTACTCCATTATTAGGTGCAACTGTAAAACCTAAATTAGGTCTTTCTGGTAAAAACTATGGTCGTGTAGTTTATGAAGGTTTAAAAGGTGGTTTAGACTTCTTAAAAGATGATGAAAACATTAACTCTCAACCATTCATGCGTTGGAGAGAACG

>DQ514822.1 Skeletonema japonicum strain NB02-45 ribulose-1,5-bisphosphate carboxylase/oxygenase large subunit (rbcL) gene, partial cds; chloroplast

TAACTTAACAGCATCTATTATTGGTAACGTTTTCGGATTTAAAGCAGTTGCTGCTTTACGTTTAGAAGATATGCGTATTCCTCACTCATACTTAAAAACATTCCAAGGTCCTGCTACAGGTATCATTGTAGAACGTGAACGTATGAACAAATATGGTACTCCATTATTAGGTGCAACTGTAAAACCTAAATTAGGTCTTTCTGGTAAAAACTATGGTCGTGTAGTTTATGAAGGTTTAAAAGGTGGTTTAGACTTCTTAAAAGATGATGAAAATATTAACTCTCAACCATTCATGCGTTGGAGAGAACG

>DQ514817.1 Skeletonema grethae strain CCAP1077/3 ribulose-1,5-bisphosphate carboxylase/oxygenase large subunit (rbcL) gene, partial cds; chloroplast

TAACTTAACAGCATCTATTATTGGTAACGTTTTCGGATTTAAAGCAGTTGCTGCTTTACGTTTAGAAGATATGCGTATTCCTCACTCATACTTAAAAACATTCCAAGGTCCTGCTACAGGTATCATTGTAGAACGTGAACGTATGAACAAATACGGTACTCCATTATTAGGTGCAACTGTAAAACCTAAATTAGGTCTTTCTGGTAAAAACTATGGTCGTGTAGTTTATGAAGGTTTAAAAGGTGGTTTAGACTTCTTAAAAGATGATGAAAATATTAACTCTCAACCATTCATGCGTTGGAGAGAACG

>KJ081746.1 Skeletonema potamos strain AJA010-19 ribulose-1,5-bisphosphate carboxylase/oxygenase large subunit (rbcL) gene, complete cds; chloroplast

TAACTTAACAGCATCTATTATTGGTAACGTTTTTGGATTTAAAGCGGTTTCTGCTTTACGTTTAGAAGATATGCGTATTCCTCACTCATACTTAAAAACATTCCAAGGTCCTGCTACAGGTATCATCGTAGAACGTGAACGTTTAAACAAATATGGTACTCCATTATTAGGTGCAACTGTAAAACCTAAATTAGGTCTTTCTGGTAAAAACTACGGTCGTGTAGTTTACGAAGGTTTAAAAGGTGGTTTAGACTTCTTAAAAGATGATGAAAATATTAACTCTCAACCATTCATGCGTTGGAGAGAACG

>MW848494.1 Chaetoceros costatus ribulose-1,5-bisphosphate carboxylase/oxygenase large subunit (rbcl) gene, complete cds; chloroplast

TAACTTAACTGCTTCTATTATTGGTAACGTATTTGGTTTTAAAGCAGTTGCTGCTTTACGTTTAGAAGATATGCGTATTCCTCACTCATACTTAAAAACATTCCAAGGTCCAGCTACAGGTATCGTTGTAGAACGTGAACGTTTAAACAAATATGGTGTACCTTTATTAGGTGCTACTGTAAAACCTAAATTAGGTTTATCTGGTAAAAACTACGGTCGTGTAGTTTTCGAAGGTTTAAAAGGTGGTTTAGACTTCTTAAAAGATGATGAGAACATTAACTCTCAACCGTTTATGCGTTGGAGAGAGCG

>HQ912440.1 Cyclotella meneghiniana strain Waco1 ribulose-1,5-bisphosphate carboxylase/oxygenase large subunit (rbcL) gene, complete cds; chloroplast

TAATTTAACAGCATCTATTATCGGTAACGTTTTTGGATTTAAAGCAGTTGCTGCTTTACGTTTAGAAGATATGCGTATTCCTCACTCATACTTAAAAACATTCCAAGGTCCTGCTACAGGTATTGTTGTAGAACGTGAACGTTTAAACAAATATGGTACTCCATTATTAGGTGCTACTGTAAAACCTAAATTAGGTCTTTCTGGTAAAAACTATGGTCGTGTAGTTTATGAAGGTTTAAAAGGTGGTTTAGACTTCTTAAAAGATGATGAAAACATTAACTCTCAACCATTTATGCGTTGGAGAGAACG

>DQ514820.1 Skeletonema subsalsum strain CCAP1077/8 ribulose-1,5-bisphosphate carboxylase/oxygenase large subunit (rbcL) gene, partial cds; chloroplast

TAACTTAACAGCATCTATTATTGGTAACGTTTTTGGATTTAAAGCAGTTTCTGCTTTACGTTTAGAAGATATGCGTATTCCTCACTCATACTTAAAAACATTCCAAGGTCCTGCTACAGGTATCATCGTAGAACGTGAACGTTTAAACAAATATGGTACTCCATTATTAGGTGCAACTGTAAAACCTAAATTAGGTCTTTCTGGTAAAAACTACGGTCGTGTAGTTTACGAAGGTTTAAAAGGTGGTTTAGACTTCTTAAAAGATGATGAAAATATTAACTCTCAACCATTCATGCGTTGGAGAGAACG

>MH687900.1 Nitzschia adhaerens isolate BIOTAII-18 ribulose-1,5-bisphosphate carboxylase/oxygenase large subunit (rbcL) gene, partial cds

TAACTTAACAGCTTCTATTATTGGTAACGTTTTCGGATTCAAAGCAATCTCTGCTTTACGTTTAGAAGATATGCGTATTCCTCACTCATACTTAAAAACATTCCAAGGTCCTGCTACAGGTATCATTGTAGAACGTGAACGTTTAAACAAATACGGTATTCCTTTATTAGGTGCAACAGTAAAACCAAAATTAGGTTTATCTGGTAAAAACTACGGTCGTGTAGTATATGAAGGTTTAAAAGGTGGTTTAGACTTCTTAAAAGATGATGAAAACATTAACTCTCAACCATTCATGCGTTGGAGAGAGCG

>MW417228.1 Minutocellus polymorphus isolate CNS00095 ribulose-1,5-bisphosphate carboxylase/oxygenase large subunit (rbcL) gene, complete cds; chloroplast

TAACTTAACTGCATCTATCATTGGTAACGTATTTGGATTCAAAGCAATTTCTGCTTTACGTTTAGAAGATATGCGTATTCCTCACTCATACTTAAAAACATTCCAAGGTCCAGCTACAGGTATCATTGTAGAGCGTGAGCGTTTAAACAAATATGGTGTTCCATTATTAGGTGCTACTGTAAAACCTAAATTAGGTCTTTCTGGTAAAAACTACGGTCGTGTAGTATACGAAGGTTTAAAAGGTGGTTTAGACTTCTTAAAAGATGATGAAAACATTAACTCTCAACCATTCATGCGTTGGAGAGAACG

>HQ912489.1 Cyclotella sp. LO4-2 ribulose-1,5-bisphosphate carboxylase/oxygenase large subunit (rbcL) gene, complete cds; chloroplast

TAATTTAACAGCATCTATTATTGGTAACGTTTTTGGATTTAAAGCAATCTCTGCTTTACGTTTAGAAGATATGCGTATTCCTCACTCATATTTAAAAACATTCCAAGGTCCTGCTACAGGTATCGTTGTAGAACGTGAACGTTTAAACAAATACGGTACTCCATTATTAGGTGCAACTGTAAAACCTAAATTAGGTCTTTCTGGTAAAAACTATGGTCGTGTAGTTTATGAAGGTTTAAAAGGTGGTTTAGACTTCTTAAAAGATGATGAAAACATTAACTCTCAACCATTCATGCGTTGGAGAGAACG

>HQ912432.1 Minutocellus polymorphus strain CCMP497 ribulose-1,5-bisphosphate carboxylase/oxygenase large subunit (rbcL) gene, complete cds; chloroplast

TAACTTAACTGCATCTATCATTGGTAACGTATTTGGATTCAAAGCAATTTCTGCTTTACGTTTAGAAGATATGCGTATTCCTCACTCATACTTAAAAACATTCCAAGGTCCAGCTACAGGTATCATTGTAGAGCGTGAGCGTTTAAACAAATATGGTGTTCCATTATTAGGTGCTACTGTAAAACCTAAATTAGGTCTTTCTGGTAAAAACTACGGTCGTGTAGTATACGAAGGTTTAAAAGGTGGTTTAGACTTCTTAAAAGATGATGAAAACATTAACTCTCAACCATTCATGCGTTGGAGAGAACG

>HQ912429.1 Brockmanniella brockmannii strain CCMP151 ribulose-1,5-bisphosphate carboxylase/oxygenase large subunit (rbcL) gene, complete cds; chloroplast

TAACTTAACTGCATCTATCATCGGTAACGTATTCGGATTCAAAGCAATCTCTGCTTTACGTTTAGAAGATATGCGTATTCCTCACTCATACTTAAAAACATTCCAAGGTCCTGCTACAGGTATCGTTGTAGAGCGTGAGCGTTTAAACAAATATGGTGCTCCATTATTAGGTGCTACTGTAAAGCCTAAATTAGGTCTTTCTGGTAAAAACTATGGTCGTGTAGTATATGAAGGTTTAAAAGGTGGTTTAGACTTTTTAAAAGATGATGAAAACATTAACTCTCAACCATTTATGCGTTGGAGAGAACG

>DQ514769.1 Lauderia annulata strain CS30 ribulose-1,5-bisphosphate carboxylase/oxygenase large subunit (rbcL) gene, partial cds; chloroplast

GAACTTAACTGCGTCTATTATTGGTAACGTTTTTGGATTTAAAGCAGTTGCTGCTTTACGTTTAGAAGATATGCGTATTCCTCACTCATACTTAAAAACATTCCAAGGTCCAGCTACAGGTATTATTGTAGAACGTGAACGTTTAAACAAATATGGTACTCCATTATTAGGTGCTACTGTTAAGCCTAAATTAGGTCTTTCTGGTAAAAACTATGGTCGTGTAGTATATGAAGGTTTAAAAGGTGGTTTAGACTTCTTAAAAGATGATGAAAACATTAACTCTCAACCATTCATGCGTTGGAGAGAACG

>MH687903.1 Nitzschia adhaerens isolate PMFBION1 ribulose-1,5-bisphosphate carboxylase/oxygenase large subunit (rbcL) gene, partial cds

TAACTTAACAGCTTCTATTATTGGTAACGTTTTCGGATTCAAAGCAATCTCTGCTTTACGTTTAGAAGATATGCGTATTCCTCACTCATACTTAAAAACATTCCAAGGTCCTGCTACAGGTATCATTGTAGAACGTGAACGTTTAAACAAATACGGTATTCCTTTATTAGGTGCAACAGTAAAACCAAAATTAGGTTTATCTGGTAAAAACTACGGTCGTGTAGTATATGAAGGTTTAAAAGGTGGTTTAGACTTCTTAAAAGATGATGAAAACATTAACTCTCAACCATTCATGCGTTGGAGAGAGCG

>DQ514826.1 Cyclostephanos tholiformis strain FHTC15 ribulose-1,5-bisphosphate carboxylase/oxygenase large subunit (rbcL) gene, partial cds; chloroplast

TAATTTAACAGCATCTATTATTGGTAACGTTTTCGGATTTAAAGCAGTTGCTGCTTTACGTTTAGAAGATATGCGTATTCCTCACTCATATTTAAAAACATTCCAAGGTCCGGCTACAGGTATTATTGTAGAACGTGAACGTTTAAACAAATATGGTACTCCTTTATTAGGTGCTACTGTAAAACCTAAATTAGGTCTTTCTGGTAAAAACTATGGTCGTGTAGTTTATGAAGGTTTAAAAGGTGGTTTAGACTTCTTAAAAGATGATGAAAACATTAACTCTCAACCATTTATGCGTTGGAGAGAACG

>MW417229.1 Minutocellus polymorphus isolate CNS00217 ribulose-1,5-bisphosphate carboxylase/oxygenase large subunit (rbcL) gene, complete cds; chloroplast

TAACTTAACTGCATCTATCATTGGTAACGTATTTGGATTCAAAGCAATTTCTGCTTTACGTTTAGAAGATATGCGTATTCCTCACTCATACTTAAAAACATTCCAAGGTCCAGCTACAGGTATCATTGTAGAGCGTGAGCGTTTAAACAAATATGGTGTTCCATTATTAGGTGCTACTGTAAAACCTAAATTAGGTCTTTCTGGTAAAAACTACGGTCGTGTAGTATACGAAGGTTTAAAAGGTGGTTTAGACTTCTTAAAAGATGATGAAAACATTAACTCTCAACCATTCATGCGTTGGAGAGAACG

>DQ514782.1 Cyclotella meneghiniana strain WC03-01 ribulose-1,5-bisphosphate carboxylase/oxygenase large subunit (rbcL) gene, partial cds; chloroplast

TAATTTAACAGCATCTATTATTGGTAACGTTTTTGGATTTAAAGCAGTTGCTGCTTTACGTTTAGAAGATATGCGTATTCCTCACTCATACTTAAAAACATTCCAAGGTCCTGCTACAGGTATTGTTGTAGAACGTGAACGTTTAAACAAATATGGTACTCCATTATTAGGTGCTACTGTAAAACCTAAATTAGGTCTTTCTGGTAAAAACTATGGTCGTGTAGTTTATGAAGGTTTAAAAGGTGGATTAGACTTCTTAAAAGATGATGAAAACATTAACTCTCAACCATTTATGCGTTGGAGAGAACG

>DQ514777.1 Cyclotella meneghiniana strain LS03-01 ribulose-1,5-bisphosphate carboxylase/oxygenase large subunit (rbcL) gene, partial cds; chloroplast

TAATTTAACAGCATCTATTATTGGTAACGTTTTTGGATTTAAAGCAGTTGCTGCTTTACGTTTAGAAGATATGCGTATTCCTCACTCATACTTAAAAACATTCCAAGGTCCTGCTACAGGTATTGTTGTAGAACGTGAACGTTTAAACAAATATGGTACTCCATTATTAGGTGCTACTGTAAAACCTAAATTAGGTCTTTCTGGTAAAAACTATGGTCGTGTAGTTTATGAAGGTTTAAAAGGTGGTTTAGACTTCTTAAAAGATGATGAAAACATTAACTCTCAACCATTTATGCGTTGGAGAGAACG

>MH687902.1 Nitzschia adhaerens isolate BIOTAII-60 ribulose-1,5-bisphosphate carboxylase/oxygenase large subunit (rbcL) gene, partial cds

TAACTTAACAGCTTCTATTATTGGTAACGTTTTCGGATTCAAAGCAATCTCTGCTTTACGTTTAGAAGATATGCGTATTCCTCACTCATACTTAAAAACATTCCAAGGTCCTGCTACAGGTATCATTGTAGAACGTGAACGTTTAAACAAATACGGTATTCCTTTATTAGGTGCAACAGTAAAACCAAAATTAGGTTTATCTGGTAAAAACTACGGTCGTGTAGTATATGAAGGTTTAAAAGGTGGTTTAGACTTCTTAAAAGATGATGAAAACATTAACTCTCAACCATTCATGCGTTGGAGAGAGCG

>MH687901.1 Nitzschia adhaerens isolate BIOTAII-59 ribulose-1,5-bisphosphate carboxylase/oxygenase large subunit (rbcL) gene, partial cds

TAACTTAACAGCTTCTATTATTGGTAACGTTTTCGGATTCAAAGCAATCTCTGCTTTACGTTTAGAAGATATGCGTATTCCTCACTCATACTTAAAAACATTCCAAGGTCCTGCTACAGGTATCATTGTAGAACGTGAACGTTTAAACAAATACGGTATTCCTTTATTAGGTGCAACAGTAAAACCAAAATTAGGTTTATCTGGTAAAAACTACGGTCGTGTAGTATATGAAGGTTTAAAAGGTGGTTTAGACTTCTTAAAAGATGATGAAAACATTAACTCTCAACCATTCATGCGTTGGAGAGAGCG

>HQ912433.1 Arcocellulus mammifer strain CCMP132 ribulose-1,5-bisphosphate carboxylase/oxygenase large subunit (rbcL) gene, complete cds; chloroplast

TAACTTAACTGCATCTATCATTGGTAACGTATTCGGATTCAAAGCAATTTCTGCTTTACGTTTAGAAGATATGCGTATTCCTCACTCATACTTAAAAACATTCCAAGGTCCAGCTACAGGTATCATTGTAGAGCGTGAGCGTTTAAACAAATATGGTGTTCCATTATTAGGTGCTACTGTAAAACCTAAATTAGGTCTTTCTGGTAAAAACTACGGTCGTGTAGTATACGAAGGTTTAAAAGGTGGTTTAGACTTCTTAAAAGATGATGAAAACATTAACTCTCAACCATTCATGCGTTGGAGAGAACG

>DQ514833.1 Discostella pseudostelligera strain ROR01-1 ribulose-1,5-bisphosphate carboxylase/oxygenase large subunit (rbcL) gene, partial cds; chloroplast

TAATTTAACAGCATCTATTATTGGTAACGTTTTTGGATTTAAAGCAATTTCTGCTTTACGTTTAGAAGATATGCGTATTCCTCACTCATATTTAAAAACATTCCAAGGTCCTGCTACAGGTATCGTTGTAGAACGTGAACGTTTAAACAAATACGGTACTCCATTATTAGGTGCAACTGTAAAACCTAAATTAGGTCTTTCTGGTAAAAACTATGGTCGTGTAGTTTATGAAGGTTTAAAAGGTGGTTTAGACTTCTTAAAAGATGATGAAAACATTAACTCTCAACCATTCATGCGTTGGAGAGAGCG

>DQ514814.1 Detonula pumila strain NB48 ribulose-1,5-bisphosphate carboxylase/oxygenase large subunit (rbcL) gene, partial cds; chloroplast

TAATTTAACAGCGTCTATTATTGGTAACGTTTTTGGATTTAAAGCAGTTGCTGCTTTACGTTTAGAAGATATGCGTATTCCTCACTCATACTTAAAAACATTCCAAGGTCCTGCTACAGGTATTGTTGTAGAACGTGAACGTTTAAACAAATATGGTACTCCTTTATTAGGTGCTACTGTAAAACCTAAATTAGGTCTTTCTGGTAAAAACTATGGTCGTGTAGTTTATGAAGGTTTAAAAGGTGGTTTAGACTTCTTAAAAGATGATGAAAACATTAACTCTCAACCATTCATGCGTTGGAGAGAACG

>AB831882.1 Stephanodiscus hantzschii chloroplast rbcL gene for ribulose-1,5-bisphosphate carboxylase/oxygenase large subunit, partial cds, specimen_voucher: TNS:AL-57172

TAATTTAACAGCATCTATTATTGGTAACGTTTTCGGATTTAAAGCAGTTGCTGCTTTACGTTTAGAAGATATGCGTATTCCTCACTCATATTTAAAAACATTCCAAGGCCCTGCTACAGGTATCGTTGTAGAGCGTGAACGTTTAAACAAATATGGTACTCCTTTATTAGGTGCTACTGTAAAACCTAAATTAGGTCTTTCTGGTAAAAACTACGGTCGTGTAGTTTATGAAGGTTTAAAAGGTGGTTTAGACTTCTTAAAAGATGATGAAAACATTAACTCTCAACCATTTATGCGTTGGAGAGAACG

>LC192328.1:97-1485 Dinophyta sp. HG180 chloroplast rbcL gene for ribulose-1,5-bisphosphate carboxylase/oxygenase large subunit, complete cds

TAACTTAACAGCATCAATTATTGGTAACGTATTCGGTTTCAAAGCTGTCTCTGCATTACGTTTAGAAGATATGCGTATTCCTCACTCATACTTAAAAACATTCCAAGGTCCTGCTACAGGTATCATTGTAGAACGTGAGCGTTTAAACAAATACGGTGTTCCTTTATTAGGTGCTACAGTAAAACCAAAATTAGGTTTATCAGGTAAAAACTACGGTCGTGTAGTATATGAAGGTTTAAAAGGTGGTTTAGACTTCTTAAAAGATGATGAAAACATTAACTCACAACCATTCATGCGTTGGAGAGAGCG

>LC192329.1:77-1465 Dinophyta sp. HG204 chloroplast rbcL gene for ribulose-1,5-bisphosphate carboxylase/oxygenase large subunit, partial cds

TAACTTAACAGCATCTATTATTGGTAACGTATTCGGTTTCAAAGCTGTATCTGCGTTACGTTTAGAAGATATGCGTATTCCTCACTCATACTTAAAAACATTCCAAGGTCCTGCTACAGGTATCGTTGTAGAACGTGAGCGTTTAAACAAATACGGTGTTCCTTTATTAGGTGCTACAGTAAAACCAAAATTAGGTTTATCTGGTAAAAACTACGGTCGTGTAGTATACGAAGGTTTAAAAGGTGGTTTAGACTTCTTAAAAGATGATGAAAACATTAACTCACAACCATTCATGCGTTGGAGAGAGCG

>LC583317.1:49-1382 Dinothrix paradoxa 2020-HS01 chloroplast rbcL gene for ribulose-1,5-bisphosphate carboxylase/oxygenase large subunit, partial cds

TAACTTAACAGCATCTATTATTGGTAACGTATTCGGTTTCAAAGCTGTATCTGCGTTACGTTTAGAAGATATGCGTATTCCTCACTCATACTTAAAAACATTCCAAGGTCCTGCTACAGGTATCGTTGTAGAACGTGAGCGTTTAAACAAATACGGTGTTCCTTTATTAGGTGCTACAGTAAAACCAAAATTAGGTTTTNNNNNNNNNNNNNNNNNNNNNNNNNTATATGAAGGTTTAAAAGGTGGTTTAGACTTCTTAAAAGATGATGAGAACATTAACTCACAACCATTCATGCGTTGGAGAGAGCG

>AB195669.1:49-1379 Galeidinium rugatum chloroplast rbcL gene for ribulose-1,5-bisphosphate carboxylase/oxygenase large subunit, partial cds

TAACTTAACAGCATCTATCATCGGTAACGTATTTGGTTTCAAAGCTGTAGCTGCATTACGTTTAGAAGATATGCGTATTCCTCACTCATACTTAAAAACATTCCAAGGTCCTGCTACAGGTATCGTTGTAGAACGTGAACGTTTAAACAAATACGGTGTACCTTTATTAGGTGCAACAGTAAAACCGAAATTAGGTTTACTGGGTAAAAACTACGGTCGNGTAGTATTCGAAGGTTTAAAAGGTGGTTTAGACTTTTTAAAAGATGATGAGAATATTAACTCTCAACCATTCATGCGTTGGAGAGAGCG

>LC192335.1:117-1505 Durinskia cf. baltica HG171 chloroplast rbcL gene for ribulose-1,5-bisphosphate carboxylase/oxygenase large subunit, complete cds

TAACTTAACAGCGTCTATTATTGGTAACGTATTCGGTTTCAAAGCTGTATCAGCTTTACGTTTAGAAGATATGCGTATTCCTCACTCATACTTAAAAACATTCCAAGGTCCTGCGACAGGTATCATCGTAGAACGTGAACGTTTAAACAAATACGGTGTTCCTTTATTAGGTGCTACAGTAAAACCAAAATTAGGTTTATCTGGTAAAAACTACGGTCGTGTAGTATACGAAGGTTTAAAAGGTGGTTTAGACTTCTTAAAAGATGATGAAAACATTAACTCTCAACCATTTATGCGTTGGAGAGAACG

>AB246745.1:49-1382 Peridinium quinquecorne chloroplast rbcL gene for ribulose-1,5-bisphosphate carboxylase/oxygenase large subunit, partial cds

TAACTTAACTGCGTCTATCATTGGTAACGTATTTGGTTTCAAAGCAGTATCTGCTTTACGTTTAGAAGATATGCGTATTCCTCACTCATACTTAAAAACATTCCAAGGTCCTGCTACAGGTATCGTTGTAGAACGTGAGCGTTTAAACAAATACGGTATCCCATTATTAGGTGCTACTGTAAAACCTAAATTAGGTTTATCTGGTAAAAACTACGGTCGTGTAGTTTATGAAGGTTTAAAAGGTGGTTTAGACTTCTTAAAAGATGATGAGAACATTAACTCTCAACCATTCATGCGTTGGAGAGAGCG

>LC192336.1:76-1464 Durinskia cf. baltica HG265 chloroplast rbcL gene for ribulose-1,5-bisphosphate carboxylase/oxygenase large subunit, partial cds

TAACTTAACAGCGTCTATTATTGGTAACGTATTCGGTTTCAAAGCTGTATCAGCTTTACGTTTAGAAGATATGCGTATTCCTCACTCATACTTAAAAACATTCCAAGGTCCTGCGACAGGTATCATCGTAGAACGTGAACGTTTAAACAAATACGGTGTTCCTTTATTAGGTGCTACAGTAAAACCAAAATTAGGTTTATCTGGTAAAAACTACGGTCGTGTAGTATACGAAGGTTTAAAAGGTGGTTTAGACTTCTTAAAAGATGATGAAAACATTAACTCTCAACCATTTATGCGTTGGAGAGAACG

>LC192325.1:117-1505 Durinskia kwazulunatalensis chloroplast rbcL gene for ribulose-1,5-bisphosphate carboxylase/oxygenase large subunit, complete cds

TAACTTAACAGCATCTATCATCGGTAACGTATTTGGTTTCAAAGCTGTATCAGCGTTACGTTTAGAAGATATGCGTATTCCTCACTCATACTTAAAAACATTCCAAGGTCCTGCTACAGGTATCGTTGTAGAACGTGAGCGTTTAAACAAATATGGTACTCCATTATTAGGTGCTACAGTAAAACCAAAATTAGGTTTATCAGGTAAAAACTATGGTCGTGTAGTATACGAAGGTTTAAAAGGTGGTTTAGACTTCTTAAAAGATGATGAGAACATTAACTCTCAACCATTTATGCGTTGGAGAGAGCG

>LC192326.1:117-1505 Durinskia kwazulunatalensis chloroplast rbcL gene for ribulose-1,5-bisphosphate carboxylase/oxygenase large subunit, complete cds

TAACTTAACAGCATCTATCATCGGTAACGTATTTGGTTTCAAAGCTGTATCAGCGTTACGTTTAGAAGATATGCGTATTCCTCACTCATACTTAAAAACATTCCAAGGTCCTGCTACAGGTATCGTTGTAGAACGTGAGCGTTTAAACAAATATGGTACTCCATTATTAGGTGCTACAGTAAAACCAAAATTAGGTTTATCAGGTAAAAACTATGGTCGTGTAGTATACGAAGGTTTAAAAGGTGGTTTAGACTTCTTAAAAGATGATGAGAACATTAACTCTCAACCATTTATGCGTTGGAGAGAGCG

>LC192327.1:117-1505 Durinskia kwazulunatalensis chloroplast rbcL gene for ribulose-1,5-bisphosphate carboxylase/oxygenase large subunit, complete cds

TAACTTAACAGCATCTATCATCGGTAACGTATTTGGTTTCAAAGCTGTATCAGCGTTACGTTTAGAAGATATGCGTATTCCTCACTCATACTTAAAAACATTCCAAGGTCCTGCTACAGGTATCGTTGTAGAACGTGAGCGTTTAAACAAATATGGTACTCCATTATTAGGTGCTACAGTAAAACCAAAATTAGGTTTATCAGGTAAAAACTATGGTCGTGTAGTATACGAAGGTTTAAAAGGTGGTCTAGACTTCTTAAAAGATGATGAGAACATTAACTCTCAACCATTTATGCGTTGGAGAGAGCG

>LC192332.1:111-1499 Durinskia capensis chloroplast rbcL gene for ribulose-1,5-bisphosphate carboxylase/oxygenase large subunit, complete cds, strain: Kommetjie 2-B

TAACTTAACAGCATCTATCATTGGTAACGTATTCGGTTTCAAAGCAGTAGCTGCTCTACGTTTAGAAGATATGCGTATTCCACACTCATATTTAAAAACATTCCAAGGTCCTGCAACAGGTATTGTTGTAGAACGTGAACGTTTAAACAAATATGGTGTTCCTTTATTAGGTGCTACAGTAAAACCAAAATTAGGTTTATCTGGTAAAAACTATGGTCGTGTAGTATACGAAGGTTTAAAAGGTGGTTTAGACTTCTTAAAAGATGATGAGAACATTAACTCTCAACCATTCATGCGTTGGAGAGAACG

>LC192334.1:113-1501 Durinskia capensis chloroplast rbcL gene for ribulose-1,5-bisphosphate carboxylase/oxygenase large subunit, complete cds, strain: Kommetjie 6-A

TAACTTAACAGCATCTATCATTGGTAACGTATTCGGTTTCAAAGCAGTAGCTGCTCTACGTTTAGAAGATATGCGTATTCCACACTCATATTTAAAAACATTCCAAGGTCCTGCAACAGGTATCGTTGTAGAACGTGAACGTTTAAACAAATATGGTGTTCCTTTATTAGGTGCTACAGTAAAACCAAAATTAGGTTTATCTGGTAAAAACTATGGTCGTGTAGTATACGAAGGTTTAAAAGGTGGTTTAGACTTCTTAAAAGATGATGAGAACATTAACTCTCAACCATTCATGCGTTGGAGAGAACG

>LC192333.1:111-1499 Durinskia capensis chloroplast rbcL gene for ribulose-1,5-bisphosphate carboxylase/oxygenase large subunit, complete cds, strain: Kommetjie 2-A

TAACTTAACAGCATCTATCATTGGTAACGTATTCGGTTTCAAAGCAGTAGCTGCTCTACGTTTAGAAGATATGCGTATTCCACACTCATATTTAAAAACATTCCAAGGTCCTGCAACAGGTATCGTTGTAGAACGTGAACGTTTAAACAAATATGGTGTTCCTTTATTAGGTGCTACAGTAAAACCAAAATTAGGTTTATCTGGTAAAAACTATGGTCGTGTAGTATACGAAGGTTTAAAAGGTGGTTTAGACTTCTTAAAAGATGATGAGAACATTAACTCTCAACCATTCATGCGTTGGAGAGAACG

>LC192331.1:110-1498 Durinskia capensis chloroplast rbcL gene for ribulose-1,5-bisphosphate carboxylase/oxygenase large subunit, complete cds, strain: Kommetjie 6-B

TAACTTAACAGCATCTATCATTGGTAACGTATTCGGTTTCAAAGCAGTAGCTGCTCTACGTTTAGAAGATATGCGTATTCCACACTCATATTTAAAAACATTCCAAGGTCCTGCAACAGGTATCGTTGTAGAACGTGAACGTTTAAACAAATATGGTGTTCCTTTATTAGGTGCTACAGTAAAACCAAAATTAGGTTTATCTGGTAAAAACTATGGTCGTGTAGTATACGAAGGTTTAAAAGGTGGTTTAGACTTCTTAAAAGATGATGAGAACATTAACTCTCAACCATTCATGCGTTGGAGAGAACG

>LC385878.1:108-1496 Durinskia capensis chloroplast NY066 rbcL gene for ribulose-1,5-bisphosphate carboxylase/oxygenase large subunit, complete cds

TAACTTAACAGCATCTATCATTGGTAACGTATTCGGTTTCAAAGCAGTAGCTGCTCTACGTTTAGAAGATATGCGTATTCCACACTCATATTTAAAAACATTCCAAGGTCCTGCAACAGGTATCGTTGTAGAACGTGAACGTTTAAACAAATATGGTGTTCCTTTATTAGGTGCTACAGTAAAACCAAAATTAGGTTTATCTGGTAAAAACTATGGTCGTGTAGTATACGAAGGTTTAAAAGGTGGTTTAGACTTCTTAAAAGATGATGAGAACATTAACTCTCAACCATTCATGCGTTGGAGAGAACG

>LC192330.2:112-1500 Durinskia capensis SaldanhaBay chloroplast rbcL gene for ribulose-1,5-bisphosphate carboxylase/oxygenase large subunit, complete cds

TAACTTAACAGCATCTATTATTGGTAACGTATTCGGTTTCAAAGCAGTAGCTGCTTTACGTTTAGAAGATATGCGTATTCCACACTCATATTTAAAAACATTCCAAGGTCCTGCGACAGGTATCGTTGTAGAACGTGAGCGTTTAAACAAATATGGTGTTCCTTTATTAGGTGCTACAGTAAAACCAAAATTAGGTTTATCTGGTAAAAACTATGGTCGTGTAGTATACGAAGGTTTAAAAGGTGGTTTAGACTTCTTAAAAGATGATGAGAACATTAACTCTCAACCATTCATGCGTTGGAGAGAGCG

>AB271108.1:1-1271 Durinskia capensis rbcL gene for large subunit of ribulose 1,5-bisphosphate carboxylase/oxygenase, partial cds

TAACTTAACAGCATCTATTATTGGTAACGTATTCGGTTTCAAAGCAGTAGCTGCTTTACGTTTAGAAGATATGCGTATTCCACACTCATATTTAAAAACATTCCAAGGTCCTGCGACAGGTATCGTTGTAGAACGTGAGCGTTTAAACAAATATGGTGTTCCTTTATTAGGTGCTACAGTAAAACCAAAATTAGGTTTNNNNNNNNNNNNNNNNNNNNNNGTAGTATACGAAGGTTTAAAAGGTGGTTTAGACTTCTTAAAAGATGATGAGAACATTAACTCTCAACCATTCATGCGTTGGAGAGAACG

>AF372696.1:49-1437 Bolidomonas pacifica strain CCMP 1866 ribulose-1,5-bisphosphate carboxylase/oxygenase large subunit gene, partial cds; chloroplast gene for chloroplast product

TAACTTAACAGCTTCTATTATTGGTAACGTATTCGGTTTCAAAGCTGTATCTGCGTTACGTTTAGAAGATATGCGTATCCCTCACTCATACTTAAAAACATTCCAAGGTCCTGCAACTGGTATTATTGTAGAACGTGAACGTTTAGACAAATATGGTCGTCCTTTATTAGGTGCAACTGTAAAACCTAAATTAGGTCTTTCAGGTAAAAACTACGGTCGTGTAGTATATGAAGGTCTTAAAGGTGGTTTAGATTTCTTAAAAGATGATGAAAACATCAACTCTCAACCATTCATGCGTTACAAAGAACG

>KR998416.1:1-1358 Bolidomonas pacifica strain RCC206 ribulose-1,5-bisphosphate carboxylase/oxygenase large subunit (rbcL) gene, partial cds; chloroplast

TAACTTAACAGCTTCTATTATTGGTAACGTATTCGGTTTCAAAGCTGTATCTGCGTTACGTTTAGAAGATATGCGTATCCCTCACTCATACTTAAAAACATTCCAAGGTCCTGCAACTGGTATTATTGTAGAACGTGAACGTTTAGACAAATATGGTCGTCCTTTATTAGGTGCAACTGTAAAACCTAAATTAGGTCTTTCAGGTAAAAACTACGGTCGTGTAGTATATGAAGGTCTTAAAGGTGGTTTAGATTTCTTAAAAGATGATGAAAACATCAACTCTCAACCATTCATGCGTTACAAAGAACG

>HQ912421.1:1-1389 Bolidomonas pacifica strain CCMP1866 ribulose-1,5-bisphosphate carboxylase/oxygenase large subunit (rbcL) gene, partial cds; chloroplast

TAACTTAACAGCTTCTATTATTGGTAACGTATTCGGTTTCAAAGCTGTATCTGCGTTACGTTTAGAAGATATGCGTATCCCTCACTCATACTTAAAAACATTCCAAGGTCCTGCAACTGGTATTATTGTAGAACGTGAACGTTTAGACAAATATGGTCGTCCTTTATTAGGTGCAACTGTAAAACCTAAATTAGGTCTTTCAGGTAAAAACTACGGTCGTGTAGTATATGAAGGTCTTAAAGGTGGTTTAGATTTCTTAAAAGATGATGAAAACATCAACTCTCAACCATTCATGCGTTACAAAGAACG

>KR998419.1:1-1234 Bolidomonas mediterranea strain RCC239 ribulose-1,5-bisphosphate carboxylase/oxygenase large subunit (rbcL) gene, partial cds; chloroplast

TAACTTAACAGCGTCTATCATTGGTAACGTATTCGGTTTCAAAGCTGTATCTGCGTTACGTTTAGAAGATATGCGTATTCCTCATTCATACTTAAAAACATTCCAAGGTCCTGCAACAGGTATTATTGTAGAACGTGAACGTTTAGATAAGTACGGTCGTCCTGTATTAGGTGCTACTGTGAAACCTAAATTAGGTCTTTCTGGTAAAAACTACGGTCGTGTAGTTTATGAAGGTTTAAAAGGTGGTTTAGATTTCTTAAAAGATGATGAAAACATTAACTCTCAACCATTCATGCGTTGGAGAGAGCG

>AF333979.1:39-1427 Bolidomonas pacifica var. eleuthera ribulose-1,5-bisphosphate carboxylase/oxygenase large subunit (rbcL) gene, partial cds; plastid gene for plastid product

TAACTTAACAGCTTCTATTATTGGTAACGTATTTGGTTTCAAAGCAGTATCTGCTTTACGTTTAGAAGATATGCGTATCCCTCACTCATACTTAAAAACTTTCCAAGGTCCTGCAACTGGTATTATTGTAGAACGTGAACGTTTAGATAAATATGGTCGTCCTGTTTTAGGTGCTACTGTAAAACCTAAATTAGGTCTTTCTGGTAAAAACTACGGTCGTGTAGTTTATGAAGGTTTAAAAGGTGGTTTAGATTTCTTAAAAGATGATGAAAACATTAACTCTCAACCATTCATGCGTTGGAGAGAGCG

>AB430698.1:49-1428 Bolidomonas pacifica chloroplast rbcL gene for ribulose 1,5-bisphosphate carboxylase/oxygenase large subunit, partial cds, strain: p380

TAACTTAACAGCTTCTATTATTGGTAACGTATTTGGTTTCAAAGCAGTATCTGCTTTACGTTTAGAAGATATGCGTATCCCTCACTCATACTTAAAAACTTTCCAAGGTCCTGCAACTGGTATTATTGTAGAACGTGAACGTTTAGATAAATATGGTCGTCCTGTTTTAGGTGCTACTGTAAAACCTAAATTAGGTCTTTCTGGTAAAAACTACGGTCGTGTAGTTTATGAAGGTTTAAAAGGTGGTTTAGATTTCTTAAAAGATGATGAAAACATTAACTCTCAACCATTCATGCGTTGGAGAGAGCG

>AF333978.1:39-1427 Bolidomonas pacifica var. eleuthera ribulose-1,5-bisphosphate carboxylase/oxygenase large subunit (rbcL) gene, partial cds; plastid gene for plastid product

TAACTTAACAGCTTCTATTATTGGTAACGTATTTGGTTTCAAAGCAGTATCTGCTTTACGTTTAGAAGATATGCGTATCCCTCACTCATACTTAAAAACTTTCCAAGGTCCTGCAACTGGTATTATTGTAGAACGTGAACGTTTAGATAAATATGGTCGTCCTGTTTTAGGTGCTACTGTAAAACCTAAATTAGGTCTTTCTGGTAAAAACTACGGTCGTGTAGTTTATGAAGGTTTAAAAGGTGGTTTAGATTTCTTAAAAGATGATGAAAACATTAACTCTCAACCATTCATGCGTTGGAGAGAGCG

>KR998417.1:1-1200 Bolidomonas pacifica strain RCC210 ribulose-1,5-bisphosphate carboxylase/oxygenase large subunit (rbcL) gene, partial cds; chloroplast

TAACTTAACAGCTTCTATTATTGGTAACGTATTTGGTTTCAAAGCAGTATCTGCTTTACGTTTAGAAGATATGCGTATCCCTCACTCATACTTAAAAACTTTCCAAGGTCCTGCAACTGGTATTATTGTAGAACGTGAACGTTTAGATAAATATGGTCGTCCTGTTTTAGGTGCTACTGTAAAACCTAAATTAGGTCTTTCTGGTAAAAACTACGGTCGTGTAGTTTATGAAGGTTTAAAAGGTGGTTTAGATTTCTTAAAAGATGATGAAAACATTAACTCTCAACCATTCATGCGTTGGAGAGAGCG

>KR998420.1:4-1365 Bolidomonas sp. ALdS-2015 ribulose-1,5-bisphosphate carboxylase/oxygenase large subunit (rbcL) gene, partial cds; chloroplast

TAACTTAACTGCATCTATTATTGGTAACGTATTCGGTTTCAAAGCTGTTTCTGCGTTACGTTTAGAAGATATGCGTATTCCTCACTCATACTTAAAAACTTTCCAAGGTCCTGCAACTGGTATTATTGTAGAACGTGAACGTTTAGATAAGTACGGTCGTCCTGTATTAGGTGCTACTGTAAAACCTAAATTAGGTCTTTCTGGTAAAAACTACGGTCGTGTAGTTTATGAAGGTCTTAAAGGTGGTTTAGATTTCTTAAAAGATGATGAAAACATTAACTCTCAACCATTCATGCGTTGGAGAGAACG

>KR998418.1:1-1200 Bolidomonas pacifica strain RCC216 ribulose-1,5-bisphosphate carboxylase/oxygenase large subunit (rbcL) gene, partial cds; chloroplast

TAACTTAACAGCTTCTATTATTGGTAACGTATTCGGTTTCAAAGCTGTATCTGCGTTACGTTTAGAAGATATGCGTATCCCTCACTCATACTTAAAAACATTCCAAGGTCCTGCAACTGGTATTATTGTAGAACGTGAACGTTTAGACAAATATGGTCGTCCTTTATTAGGTGCAACTGTAAAACCTAAATTAGGTCTTTCAGGTAAAAACTACGGTCGTGTAGTATATGAAGGTCTTAAAGGTGGTTTAGATTTCTTAAAAGATGATGAAAACATCAACTCTCAACCATTCATGCGTTACAAAGAACG

>AF333977.1:53-1441 Bolidomonas mediterranea ribulose-1,5-bisphosphate carboxylase/oxygenase large subunit (rbcL) gene, partial cds; plastid gene for plastid product

TAACTTAACAGCGTCTATCATTGGTAACGTATTCGGTTTCAAAGCTGTATCTGCGTTACGTTTAGAAGATATGCGTATTCCTCATTCATACTTAAAAACATTCCAAGGTCCTGCAACAGGTATTATTGTAGAACGTGAACGTTTAGATAAGTACGGTCGTCCTGTATTAGGTGCTACTGTGAAACCTAAATTAGGTCTTTCTGGTAAAAACTACGGTCGTGTAGTTTATGAAGGTTTAAAAGGTGGTTTAGATTTCTTAAAAGATGATGAAAACATTAACTCTCAACCATTCATGCGTTGGAGAGAGCG

>KR998415.1:1-1136 Triparma strigata strain NIES-3701 ribulose-1,5-bisphosphate carboxylase/oxygenase large subunit (rbcL) gene, partial cds; chloroplast

TAACTTAACTGCATCTATTATTGGTAACGTATTCGGTTTCAAAGCTGTTTCTGCGTTACGTTTAGAAGATATGCGTATTCCTCACTCATACTTAAAAACTTTCCAAGGTCCTGCAACTGGTATTATTGTAGAACGTGAACGTTTAGATAAGTACGGTCGTCCTGTATTAGGTGCTACTGTAAAACCTAAATTAGGTCTTTCTGGTAAAAACTACGGTCGTGTAGTTTATGAAGGTCTTAAAGGTGGTTTAGATTTCTTAAAGGATGATGAAAACATTAACTCTCAACCATTCATGCGTTGGAGAGAACG

>MK659576.1:1-1129 Triparma retinervis voucher TRNT-1512-15 ribulose-1,5-bisphosphate carboxylase/oxygenase large subunit (rbcL) gene, partial cds; chloroplast

TAACTTAACAGCTTCTATTATTGGTAACGTATTCGGTTTCAAAGCTGTATCTGCGTTACGTTTAGAAGATATGCGTATCCCTCACTCATACTTAAAAACATTCCAAGGTCCTGCAACTGGTATTATTGTAGAACGTGAACGTTTAGACAAGTACGGTCGTCCTTTATTAGGTGCTACTGTAAAACCTAAATTAGGTCTTTCTGGTAAAAACTACGGTCGTGTAGTATATGAAGGTCTTAAAGGTGGTTTAGATTTCTTAAAAGATGATGAAAACATCAACTCTCAACCATTCATGCGTTACAAAGAACG

>KR998423.1:1-1361 Triparma aff. verrucosa ALdS-2015 strain NIES-3700 ribulose-1,5-bisphosphate carboxylase/oxygenase large subunit (rbcL) gene, partial cds; chloroplast

TAACTTAACTGCATCTATTATTGGTAACGTATTCGGTTTCAAAGCTGTTTCTGCGTTACGTTTAGAAGATATGCGTATTCCTCACTCATACTTAAAAACTTTCCAAGGTCCTGCAACTGGTATTATTGTAGAACGTGAACGTTTAGATAAGTACGGTCGTCCTGTATTAGGTGCTACTGTAAAACCTAAATTAGGTCTTTCTGGTAAAAACTACGGTCGTGTAGTTTATGAAGGTCTTAAAGGTGGTTTAGATTTCTTAAAAGATGATGAAAACATTAACTCTCAACCATTCATGCGTTGGAGAGAACG

>MK659575.1:1-1356 Triparma retinervis voucher TRNT-1512-10 ribulose-1,5-bisphosphate carboxylase/oxygenase large subunit (rbcL) gene, partial cds; chloroplast

TAACTTAACAGCTTCTATTATTGGTAACGTATTCGGTTTCAAAGCTGTATCTGCGTTACGTTTAGAAGATATGCGTATCCCTCACTCATACTTAAAAACATTCCAAGGTCCTGCAACTGGTATTATTGTAGAACGTGAACGTTTAGACAAGTACGGTCGTCCTTTATTAGGTGCTACTGTAAAACCTAAATTAGGTCTTTCTGGTAAAAACTACGGTCGTGTAGTATATGAAGGTCTTAAAGGTGGTTTAGATTTCTTAAAAGATGATGAAAACATCAACTCTCAACCATTCATGCGTTACAAAGAACG

>KR998421.1:1-1200 Triparma eleuthera strain RCC1730 ribulose-1,5-bisphosphate carboxylase/oxygenase large subunit (rbcL) gene, partial cds; chloroplast

TAACTTAACWGCTTCTATTATTGGTAACGTATTTGGTTTCAAAGCAGTATCTGCTTTACGTTTAGAAGATATGCGTATCCCTCACTCATACTTAAAAACTTTCCAAGGTCCTGCAACTGGTATTATTGTAGAACGTGAACGTTTAGATAAATATGGTCGTCCTGTTTTAGGTGCTACTGTAAAACCTAAATTAGGTCTTTCTGGTAAAAACTACGGTCGTGTAGTTTATGAAGGTTTAAAAGGTGGTTTAGATTTCTTAAAAGATGATGAAAACATTAACTCTCAACCATTCATGCGTTGGAGAGAGCG

>AB546640.1:79-1467 Triparma sp. TOY-0807 chloroplast rbcL gene for ribulose bisphosphate carboxylase large chain, partial cds

TAACTTAACTGCATCTATTATTGGTAACGTATTCGGTTTCAAAGCTGTTTCTGCGTTACGTTTAGAAGATATGCGTATCCCTCACTCGTACTTAAAAACTTTCCAAGGTCCTGCAACTGGTATTATTGTAGAACGTGAACGTTTAGATAAGTACGGTCGTCCTGTATTAGGTGCTACTGTAAAACCTAAATTAGGTCTTTCTGGTAAAAACTACGGTCGTGTAGTTTATGAAGGTCTTAAAGGTGGTTTAGACTTCTTAAAAGATGATGAAAACATTAACTCTCAACCATTCATGCGTTGGAGAGAACG

>NC_027746.1:51169-52557 Triparma laevis chloroplast DNA, complete sequence

TAACTTAACTGCATCTATTATTGGTAACGTATTCGGTTTCAAAGCTGTTTCTGCGTTACGTTTAGAAGATATGCGTATCCCTCACTCGTACTTAAAAACTTTCCAAGGTCCTGCAACTGGTATTATTGTAGAACGTGAACGTTTAGATAAGTACGGTCGTCCTGTATTAGGTGCTACTGTAAAACCTAAATTAGGTCTTTCTGGTAAAAACTACGGTCGTGTAGTTTATGAAGGTCTTAAAGGTGGTTTAGACTTCTTAAAAGATGATGAAAACATTAACTCTCAACCATTCATGCGTTGGAGAGAACG

>KR998414.1:1-1200 Triparma laevis f. longispina strain NIES-3699 ribulose-1,5-bisphosphate carboxylase/oxygenase large subunit (rbcL) gene, partial cds; chloroplast

TAACTTAACTGCATCTATTATTGGTAACGTATTCGGTTTCAAAGCTGTTTCTGCGTTACGTTTAGAAGATATGCGTATCCCTCACTCGTACTTAAAAACTTTCCAAGGTCCTGCAACTGGTATTATTGTAGAACGTGAACGTTTAGATAAGTACGGTCGTCCTGTATTAGGTGCTACTGTAAAACCTAAATTAGGTCTTTCTGGTAAAAACTACGGTCGTGTAGTTTATGAAGGTCTTAAAGGTGGTTTAGACTTCTTAAAAGATGATGAAAACATTAACTCTCAACCATTCATGCGTTGGAGAGAACG

>GQ231541.1:60100-61566 Aureococcus anophagefferens strain CCMP 1984 chloroplast, complete genome

-AACATGACTGCTTCTATTATTGGTAACGTATTCGGTTTCAAGGCCGTTAAAGCGTTACGTTTAGAAGATATGCGTATTCCACACTCTTACTTAAAAACATTCCAAGGTCCTGCGACTGGGGTTGTTGTTGAGCGTGAGCGTATGGATAAATTCGGTCGTCCATTATTAGGTGCTACTGTTAAGCCTAAGTTAGGTTTATCTGGTAAGAACTACGGACGTGTAGTATTCGAAGGTTTAAAAGGTGGTTTAGACTTCTTAAAGGATGATGAGAACATTAACTCACAACCATTCATGCGTTGGCGTGAGCG

>PQ791844.1 Sungminbooa capricornica ribulose-1,5-bisphosphate carboxylase/oxygenase large subunit (rbcL) gene, complete cds; chloroplast

TAACATGACAGCTTCTATTATTGGTAACGTATTCGGTTTCAAAGCCGTTAAAGCATTACGTTTAGAAGATATGCGTATTCCTCACTCATACTTAAAAACTTTCCAAGGTCCTGCAACTGGTGTAATTGTAGAGCGTGAGCGTTTAGATACGTTTGGTCGTCCACTTTTAGGTGCAACAGTTAAGCCTAAATTAGGTTTATCAGGTAAAAACTACGGTCGTGTAGTTTTCGAAGGTTTAAAAGGTGGTTTAGACTTCTTAAAAGATGATGAGAACATTAACTCACAACCATTCATGCGTTGGCGTGAGCG

>AF117904.1 Aureococcus anophagefferens strain CCMP1707 ribulose-1,5-bisphosphate carboxylase/oxygenase large subunit (rbcL) gene, chloroplast gene encoding chloroplast protein, partial cds

GAACATGACTGCTTCTATTATTGGTAACGTATTCGGTTTCAAGGCCGTTAAAGCGTTACGTTTAGAAGATATGCGTATTCCACACTCTTACTTAAAAACATTCCAAGGTCCTGCGACTGGGGTTGTTGTTGAGCGTGAGCGTATGGATAAATTCGGTCGTCCATTATTAGGTGCTACTGTTAAGCCTAAGTTAGGTTTATCTGGTAAGAACTACGGACGTGTAGTATTCGAAGGTTTAAAAGGTGGTTTAGACTTCTTAAAGGATGATGAGAACATTAACTCACAACCATTCATGCGTTGGCGTGAGCG

>AF117785.1 Aureococcus anophagefferens strain CCMP1785 ribulose-1,5-bisphosphate carboxylase/oxygenase large subunit (rbcL) gene, chloroplast gene encoding chloroplast protein, partial cds

GAACATGACTGCTTCTATTATTGGTAACGTATTCGGTTTCAAGGCCGTTAAAGCGTTACGTTTAGAAGATATGCGTATTCCACACTCTTACTTAAAAACATTCCAAGGTCCTGCGACTGGGGTTGTTGTTGAGCGTGAGCGTATGGATAAATTCGGTCGTCCATTATTAGGTGCTACTGTTAAGCCTAAGTTAGGTTTATCTGGTAAGAACTACGGACGTGTAGTATTCGAAGGTTTAAAAGGTGGTTTAGACTTCTTAAAGGATGATGAGAACATTAACTCACAACCATTCATGCGTTGGCGTGAGCG

>AF117784.1 Aureococcus anophagefferens strain CCMP1708 ribulose-1,5-bisphosphate carboxylase/oxygenase large subunit (rbcL) gene, chloroplast gene encoding chloroplast protein, partial cds

GAACATGACTGCTTCTATTATTGGTAACGTATTCGGTTTCAAGGCCGTTAAAGCGTTACGTTTAGAAGATATGCGTATTCCACACTCTTACTTAAAAACATTCCAAGGTCCTGCGACTGGGGTTGTTGTTGAGCGTGAGCGTATGGATAAATTCGGTCGTCCATTATTAGGTGCTACTGTTAAGCCTAAGTTAGGTTTATCTGGTAAGAACTACGGACGTGTAGTATTCGAAGGTTTAAAAGGTGGTTTAGACTTCTTAAAGGATGATGAGAACATTAACTCACAACCATTCATGCGTTGGCGTGAGCG

>AF117783.1 Aureococcus anophagefferens strain CCMP1706 ribulose-1,5-bisphosphate carboxylase/oxygenase large subunit (rbcL) gene, chloroplast gene encoding chloroplast protein, partial cds

GAACATGACTGCTTCTATTATTGGTAACGTATTCGGTTTCAAGGCCGTTAAAGCGTTACGTTTAGAAGATATGCGTATTCCACACTCTTACTTAAAAACATTCCAAGGTCCTGCGACTGGGGTTGTTGTTGAGCGTGAGCGTATGGATAAATTCGGTCGTCCATTATTAGGTGCTACTGTTAAGCCTAAGTTAGGTTTATCTGGTAAGAACTACGGACGTGTAGTATTCGAAGGTTTAAAAGGTGGTTTAGACTTCTTAAAGGATGATGAGAACATTAACTCACAACCATTCATGCGTTGGCGTGAGCG

>HQ710615.1 Aureococcus anophagefferens culture CCMP:1984 ribulose-1,5-bisphosphate carboxylase/oxygenase large subunit (rbcL) gene, partial cds; plastid

GAACATGACTGCTTCTATTATTGGTAACGTATTCGGTTTCAAGGCCGTTAAAGCGTTACGTTTAGAAGATATGCGTATTCCACACTCTTACTTAAAAACATTCCAAGGTCCTGCGACTGGGGTTGTTGTTGAGCGTGAGCGTATGGATAAATTCGGTCGTCCATTATTAGGTGCTACTGTTAAGCCTAAGTTAGGTTTATCTGGTAAGAACTACGGACGTGTAGTATTCGAAGGTTTAAAAGGTGGTTTAGACTTCTTAAAGGATGATGAGAACATTAACTCACAACCATTCATGCGTTGGCGTGAGCG

>ON988288.1 Wyeophycus julieharrissiae isolate Wye36 ribulose-1,5-bisphosphate carboxylase/oxygenase large subunit (rbcL) gene, complete cds; plastid

AAATATGACAGCTTCTATCATTGGTAACGTATTCGGCTTCAAGGCTGTTAAGGCGTTACGTTTAGAAGATATGCGTATTCCACACTCATACTTAAAGACATTCCAAGGTCCTGCGACTGGTGTAATTGTTGAGCGTGAGCGTTTAGACAAGTTCGGCCGTCCGCTTTTAGGTGCAACTGTTAAGCCTAAGTTAGGTCTTTCTGGTAAGAACTACGGTCGTGTAGTTTTCGAAGGTTTAAAAGGTGGTTTAGACTTCTTAAAGGATGATGAGAACATTAACTCACAACCATTCATGCGTTGGAGAGAGCG

>HQ710614.1 Veerella venusta voucher RCC-286 ribulose-1,5-bisphosphate carboxylase/oxygenase large subunit (rbcL) gene, partial cds; plastid

AAATATGACAGCTTCTATTATTGGTAACGTATTCGGTTTCAAAGCCGTTAAAGCGTTACGTTTAGAAGATATGCGTATTCCACACTCATACTTAAAGACATTCCAAGGTCCTGCAACTGGTGTAGTTGTAGAGCGTGAGCGTTTAGATAAGTTTGGTCGTCCACTTTTAGGTGCTACTGTTAAGCCTAAGTTAGGTTTATCTGGTAAGAACTACGGTCGTGTAGTTTTTGAAGGTTTAAAAGGTGGTTTAGATTTCTTAAAGGATGATGAGAACATTAACTCACAGCCATTCATGCGTTGGAGAGAGCG

>PQ791843.1 Sungminbooa tropica ribulose-1,5-bisphosphate carboxylase/oxygenase large subunit (rbcL) gene, complete cds; chloroplast

AAACATGACAGCTTCTATTATCGGTAACGTATTTGGTTTCAAGGCCGTTAAAGCGTTACGTTTAGAAGATATGCGTATTCCTCACTCATACTTAAAGACTTTCCAAGGTCCTGCAACTGGTGTAATTGTAGAACGTGAGCGTTTAGATACGTTTGGTCGTCCACTTTTAGGTGCAACAGTTAAGCCTAAGTTAGGTTTATCAGGTAAGAACTACGGTCGTGTAGTTTTTGAAGGTTTAAAAGGTGGTTTAGATTTCTTAAAGGATGATGAAAACATCAACTCACAACCATTCATGCGTTGGCGTGAGCG

>MF927466.1 Chrysoreinhardia muelleri strain Chryso 1 ribulose-1,5-bisphosphate carboxylase/oxygenase large subunit (rbcL) gene, partial cds; plastid

AAACATGACTGCTTCAATTATTGGTAACGTATTTGGTTTCAAAGCAGTTAAAGCATTACGTTTAGAGGATATGCGTATTCCTCATTCGTACTTAAAAACATTCCAAGGACCTGCTACAGGTGTAATTGTAGAACGTGAGCGTTTAGATACTTTCGGTCGTCCACTTTTAGGTGCGACAGTTAAGCCTAAGTTAGGTTTATCTGGTAAAAACTACGGTCGTGTAGTATTTGAAGGTTTAAAAGGTGGTCTAGATTTCTTAAAGGATGATGAGAACATCAACTCACAACCATTTATGCGCTGGCGTGAGCG

>ON988286.1 Pituiglomerulus carpicornicus isolate HI1 ribulose-1,5-bisphosphate carboxylase/oxygenase large subunit (rbcL) gene, partial cds; plastid

AAATATGACAGCTTCTATTATTGGTAACGTATTCGGTTTCAAAGCCGTAAAAGCATTACGTTTAGAAGATATGCGTATTCCTCATTCATACTTAAAGACTTTCCAAGGTCCTGCAACTGGTGTGATTGTAGAGCGTGAGCGTTTAGATACTTTTGGTCGTCCACTTTTAGGTGCTACCGTTAAACCTAAGTTAGGTTTATCTGGAAAGAACTATGGTCGTGTAGTTTTTGAAGGTTTAAAAGGTGGTTTAGACTTCTTAAAGGATGATGAGAATATTAACTCACAACCATTCATGCGTTGGCGTGAGCG

>ON988287.1 Chromopallida australis isolate Och39 ribulose-1,5-bisphosphate carboxylase/oxygenase large subunit (rbcL) gene, complete cds; plastid

AAATATGACAGCTTCTATCATCGGTAACGTATTCGGTTTCAAAGCTGTTAAAGCGTTACGTTTAGAAGATATGCGTATCCCTCACTCTTACTTAAAGACTTTCCAAGGTCCAGCAACTGGTGTAGTTGTAGAGCGTGAGCGTTTAGACAAGTTCGGTCGTCCACTTTTAGGTGCTACTGTTAAGCCTAAGTTAGGTTTATCAGGTAAGAACTACGGTCGTGTAGTATTTGAAGGTTTAAAAGGTGGTTTAGACTTCTTAAAGGATGATGAGAACATTAACTCACAACCATTCATGCGTTGGAGAGAGCG

>HQ710618.1 Pelagococcus subviridis culture CCMP:1429 ribulose-1,5-bisphosphate carboxylase/oxygenase large subunit (rbcL) gene, partial cds; plastid

TAACATGACTGCTTCTATTATTGGTAACGTATTCGGTTTCAAAGCTGTTAAAGCGTTACGTTTAGAAGATATGCGTATCCCACACTCATACTTAAAGACTTTCCAAGGTCCTGCTTCTGGTGTAGTTGTAGAGCGTGAGCGTTTAGACAAGTTCGGTCGCCCTCTTTTAGGTGCTACAGTAAAGCCTAAGTTAGGTTTATCTGGTAAGAACTACGGTCGTGTAGTATTCGAAGGTTTAAAAGGTGGTTTAGACTTCTTAAAGGATGACGAGAACATTAACTCACAACCATTCATGCGTTGGCGTGAGCG

>MT439864.1 Gazia saundersii strain CS-1320 ribulose-1,5-bisphosphate carboxylase/oxygenase large subunit (rbcL) gene, complete cds; chloroplast

AAACATGACTGCATCTATCATTGGTAATGTATTCGGTTTCAAGGCTGTAAAAGCATTACGTTTAGAAGATATGCGTATTCCACACTCTTACTTAAAGACTTTCCAAGGTCCTGCTACAGGTGTAGTTGTAGAGCGTGAGCGTTTAGATACTTTCGGTCGTCCTCTTTTAGGTGCAACAGTAAAGCCTAAGTTAGGTCTTTCTGGTCGTAACTACGGTCGTGTAGTTTTCGAAGGTTTAAAAGGTGGTTTAGACTTCTTAAAGGATGATGAGAATATTAACTCACAACCATTTATGCGTTGGCGTGAGCG

>PQ791845.1 Revolvomonas australis ribulose-1,5-bisphosphate carboxylase/oxygenase large subunit (rbcL) gene, complete cds; chloroplast

GAACATGACAGCTTCTATTATTGGTAACGTATTTGGTTTCAAAGCTGTTAAAGCGTTACGTTTAGAAGATATGCGTATCCCTCATTCATACTTAAAGACTTTCCAAGGTCCTGCAACAGGTGTAATTGTAGAACGTGAGCGTTTAGATACATTTGGTCGTCCACTTTTAGGTGCAACAGTTAAGCCTAAATTAGGTTTATCTGGTAAAAACTACGGTCGTGTAGTTTTCGAAGGCTTAAAGGGTGGTTTAGATTTCTTAAAAGATGATGAGAATATTAACTCTCAACCATTTATGCGCTGGCGTGAGCG

>MT469981.1 Aureoumbra geitleri strain CS-1323 ribulose-1,5-bisphosphate carboxylase/oxygenase large subunit (rbcL) gene, complete cds; chloroplast

TAACATGACAGCTTCTATTATTGGTAACGTATTCGGTTTCAAAGCTGTAAAAGCGTTACGTTTAGAAGATATGCGTATTCCACACTCTTACTTAAAGACTTTCCAAGGTCCTGCTACTGGTGTGATTGTAGAGCGTGAGCGTTTAGACACTTTTGGACGTCCACTTTTAGGTGCTACAGTAAAACCTAAGTTAGGTCTTTCTGGTAAGAACTACGGTCGTGTAGTTTTTGAAGGTTTAAAAGGTGGTTTAGACTTCTTAAAGGATGATGAGAATATTAACTCACAACCTTTCATGCGTTGGCGTGAGCG

>PQ791841.1 Aureoumbra geitleri ribulose-1,5-bisphosphate carboxylase/oxygenase large subunit (rbcL) gene, complete cds; chloroplast

TAACATGACAGCTTCTATTATTGGTAACGTATTCGGTTTCAAAGCTGTAAAAGCGTTACGTTTAGAAGATATGCGTATTCCACACTCTTACTTAAAGACTTTCCAAGGTCCTGCTACTGGTGTGATTGTAGAGCGTGAGCGTTTAGACACTTTTGGACGTCCACTTTTAGGTGCTACAGTAAAACCTAAGTTAGGTCTTTCTGGTAAGAACTACGGTCGTGTAGTTTTTGAAGGTTTAAAAGGTGGTTTAGACTTCTTAAAGGATGATGAGAATATTAACTCACAACCTTTCATGCGTTGGCGTGAGCG

>MT439865.1 Glomerochrysis psammophila strain CS-1322 ribulose-1,5-bisphosphate carboxylase/oxygenase large subunit (rbcL) gene, complete cds; chloroplast

GAATATGACAGCTTCTATCATCGGTAACGTATTTGGTTTCAAAGCCGTTAAAGCGTTACGTTTAGAAGATATGCGTATTCCACACTCTTACTTAAAGACTTTCCAAGGTCCTGCAACAGGTGTAATTGTAGAGCGTGAGCGTTTAGATACTTATGGCCGTCCTCTTTTAGGTGCTACAGTTAAGCCTAAGTTAGGTCTTTCTGGTCGTAACTACGGTCGTGTAGTATTTGAAGGTTTAAAGGGTGGTTTAGACTTCTTAAAGGATGATGAGAACATTAACTCTCAACCTTTCATGCGTTGGCGTGAGCG

>AF117786.1 Aureoumbra lagunensis ribulose-1,5-bisphosphate carboxylase/oxygenase large subunit (rbcL) gene, chloroplast gene encoding chloroplast protein, partial cds

TAATATGACAGCATCTATCATTGGTAACGTATTTGGTTTCAAAGCTGTAAAAGCGTTACGTTTAGAAGATATGCGTATTCCACACTCTTACTTAAAGACTTTCCAAGGTCCTGCTACTGGTGTGATCGTAGAGCGTGAGCGTTTAGACACATTCGGTCGTCCACTTTTAGGTGCTACAGTAAAACCTAAATTAGGTTTATCTGGTAAGAACTACGGTCGTGTAGTTTTCGAAGGTTTAAAAGGTGGTTTAGACTTTTTAAAGGATGATGAAAACATTAACTCGCAACCTTTCATGCGTTGGCGTGAGCG

>AF118136.1 Aureoumbra lagunensis strain CCMP1510 ribulose-1,5-bisphosphate carboxylase/oxygenase large subunit (rbcL) gene, chloroplast gene encoding chloroplast protein, partial cds

TAATATGACAGCATCTATCATTGGTAACGTATTTGGTTTCAAAGCTGTAAAAGCGTTACGTTTAGAAGATATGCGTATTCCACACTCTTACTTAAAGACTTTCCAAGGTCCTGCTACTGGTGTGATCGTAGAGCGTGAGCGTTTAGACACATTCGGTCGTCCACTTTTAGGTGCTACAGTAAAACCTAAATTAGGTTTATCTGGTAAGAACTACGGTCGTGTAGTTTTCGAAGGTTTAAAAGGTGGTTTAGACTTTTTAAAGGATGATGAAAACATTAACTCGCAACCTTTCATGCGTTGGCGTGAGCG

>HQ710616.1 Aureoumbra lagunensis culture CCMP:1510 ribulose-1,5-bisphosphate carboxylase/oxygenase large subunit (rbcL) gene, partial cds; plastid

TAATATGACAGCATCTATCATTGGTAACGTATTTGGTTTCAAAGCTGTAAAAGCGTTACGTTTAGAAGATATGCGTATTCCACACTCTTACTTAAAGACTTTCCAAGGTCCTGCTACTGGTGTGATCGTAGAGCGTGAGCGTTTAGACACATTCGGTCGTCCACTTTTAGGTGCTACAGTAAAACCTAAATTAGGTTTATCTGGTAAGAACTACGGTCGTGTAGTTTTCGAAGGTTTAAAAGGTGGTTTAGACTTTTTAAAGGATGATGAAAACATTAACTCGCAACCTTTCATGCGTTGGCGTGAGCG

>PQ791842.1 Chrysoreinhardia giraudii ribulose-1,5-bisphosphate carboxylase/oxygenase large subunit (rbcL) gene, complete cds; chloroplast

AAATATGACTGCTTCTATTATTGGTAACGTATTTGGTTTCAAAGCGGTGAAAGCATTACGTTTAGAGGATATGCGTATTCCTCATTCATACTTAAAGACTTTCCAAGGTCCTGCGACTGGTGTAATCGTAGAGCGTGAGCGTTTAGATACTTTCGGTCGTCCACTTTTAGGTGCGACAGTTAAGCCTAAATTAGGTTTATCAGGTAAAAACTACGGTCGTGTAGTATTCGAAGGTTTAAAAGGTGGTTTAGATTTCTTAAAGGATGATGAGAACATCAACTCACAACCATTCATGCGTTGGCGTGAGCG

>HQ710622.1 Sarcinochrysis sp. HSY-2011 culture CCMP:770 ribulose-1,5-bisphosphate carboxylase/oxygenase large subunit (rbcL) gene, partial cds; plastid

AAATATGACTGCTTCTATTATTGGTAACGTATTTGGTTTCAAAGCGGTGAAAGCATTACGTTTAGAGGATATGCGTATTCCTCATTCATACTTAAAGACTTTCCAAGGTCCTGCGACTGGTGTAATCGTAGAGCGTGAGCGTTTAGATACTTTCGGTCGTCCACTTTTAGGTGCGACAGTTAAGCCTAAATTAGGTTTATCAGGTAAAAACTACGGTCGTGTAGTATTCGAAGGTTTAAAAGGTGGTTTAGATTTCTTAAAGGATGATGAGAACATCAACTCACAACCATTCATGCGTTGGCGTGAGCG

>HQ710621.1 Sarcinochrysis sp. HSY-2011 voucher A 11.864 ribulose-1,5-bisphosphate carboxylase/oxygenase large subunit (rbcL) gene, partial cds; plastid

AAATATGACAGCTTCTATTATTGGTAACGTATTTGGTTTCAAAGCCGTTAAAGCGTTACGTTTAGAAGACATGCGGATTCCTCACTCATACCTAAAAACTTTCCAAGGTCCTGCAACTGGTGTAATTGTAGAACGTGAACGTTTAGATACGTTTGGTCGTCCACTTTTAGGTGCAACAGTTAAGCCTAAATTAGGTTTATCAGGTAAGAACTATGGTCGCGTAGTTTTCGAAGGTTTAAAAGGTGGTTTAGATTTCTTAAAAGATGATGAAAACATCAATTCACAACCATTCATGCGTTGGCGTGAGCG

>PQ791840.1 Pelagomonas calceolata ribulose-1,5-bisphosphate carboxylase/oxygenase large subunit (rbcL) gene, complete cds; chloroplast

GAACATGACTGCTTCGATTATTGGTAACGTATTCGGTTTCAAAGCTGTTAAAGCGTTACGTTTAGAAGATATGCGTATCCCACACACTTACTTAAAGACTTTCCAAGGTCCTGCTTCTGGTGTAATTGTAGAGCGTGAGCGTTTAGACACATTCGGACGTCCACTTTTAGGTGCTACAGTAAAGCCTAAGTTAGGTTTATCTGGTAAGAACTACGGTCGTGTAGTATTCGAAGGTTTAAAAGGTGGTTTAGACTTCTTAAAGGATGATGAGAACATTAACTCACAACCATTCATGCGTTGGAGAGAGCG

>MT439863.1 Gazia australica strain CS-1321 ribulose-1,5-bisphosphate carboxylase/oxygenase large subunit (rbcL) gene, complete cds; chloroplast

AAACATGACTGCATCTATCATTGGTAATGTATTCGGTTTCAAGGCTGTAAAAGCATTACGTTTAGAGGATATGCGTATCCCACACTCTTACTTAAAGACTTTCCAAGGTCCTGCTACAGGTGTAATTGTAGAGCGTGAGCGTTTAGATACTTTCGGTCGTCCTCTTTTAGGTGCAACAGTAAAGCCTAAGTTAGGTCTTTCTGGTCGTAACTACGGTCGTGTAGTTTTCGAAGGTTTAAAAGGTGGTTTAGACTTCTTAAAGGATGATGAGAACATTAACTCACAACCATTTATGCGTTGGCGTGAGCG

>KM010150.1 Pelagomonas calceolata strain CCMP 1756 ribulose-1,5-bisphosphate carboxylase/oxygenase large subunit (rbcL) gene, partial cds; plastid

GAACATGACTGCTTCGATTATTGGTAACGTATTCGGTTTCAAAGCTGTTAAAGCGTTACGTTTAGAAGATATGCGTATCCCACACAGTTACTTAAAGACTTTCCAAGGTCCTGCTTCTGGTGTAGTTGTAGAGCGTGAGCGTTTAGACACATTCGGACGTCCACTTTTAGGTGCTACAGTAAAGCCTAAGTTAGGTTTATCTGGTAAGAACTACGGTCGTGTAGTATTCGAAGGTTTAAAAGGTGGTTTAGACTTCTTAAAGGATGATGAGAACATTAACTCACAACCATTCATGCGTTGGAGAGAGCG

>gi|125617018|gb|EF165204.1| Ochromonas sp. AC514 ribulose-1,5-bisphosphate carboxylase/oxygenase large subunit (rbcL) gene, partial cds; chloroplast

-AACTTAACGGCTTCAATTATCGGAAACGTATTTGGATTTAAAGCTGTAAAAGCATTACGTTTAGAAGATATGCGTATTCCTTATGCATACTTAAAAACATTCCAAGGTCCTGCGACTGGAGTAGTTGTAGAACGTGAAAGATTAGATATTTTTGGACGTCCTTTTTTAGGAGCTACTGTAAAACCAAAATTAGGTCTTTCAGGTAAAAACTATGGTCGTGTAGTTTATGAAGGATTACAAGGTGGTTTAGACTTCTTAAAAGATGATGAGAATATTAACTCACAACCATTTATGCGTTGGCGTGAAAG

>gi|125617020|gb|EF165205.1| Ochromonas sp. CCMP1147 ribulose-1,5-bisphosphate carboxylase/oxygenase large subunit (rbcL) gene, partial cds; chloroplast

AAACTTAACAGCGTCAATCATTGGAAACGTATTTGGATTTAAAGCTGTAAAAGCATTACGTTTAGAAGATATGCGTATTCCTTATGCATACTTAAAAACATTCCAAGGTCCTGCAACTGGTGTAGTTGTAGAACGTGAAAGATTAGATATCTTCGGACGTCCTTTCTTAGGAGCTACTGTTAAACCAAAATTAGGTCTTTCAGGAAAAAATTACGGTCGTGTAGTTTATGAAGGATTACAAGGTGGTTTAGATTTCTTAAAAGATGATGAGAATATTAACTCACAACCATTTATGCGATGGCGTGAAAG

>gi|125617010|gb|EF165200.1| Ochromonas sp. CCMP2761 ribulose-1,5-bisphosphate carboxylase/oxygenase large subunit (rbcL) gene, partial cds; chloroplast

TAACTTAACAGCTTCTATTATCGGAAACGTTTTTGGTTTTAAAGCAGTAAAATGTTTACGTTTAGAAGATATGCGTATTCCTTATGCATATTTAAAAACTTTCATTGGTCCTGCTACAGGTGTAGTTGTAGAACGTGAAAGATTAGACGTTTTTGGACGTCCTCTTTTAGGTGCAACAGTAAAACCAAAATTAGGTCTTTCTGGAAAAAATTATGGTCGTGTAGTTTATGAAGGATTAAAAGGTGGTTTAGACTTCTTAAAAGATGACGAAAATATTAACTCTCAACCATTCATGCGTTGGCGTGAACG

>gi|1152729548|gb|KY575275.1| Ochromonas triangulata strain A14 651 ribulose-1,5-bisphosphate carboxylase/oxygenase large subunit (rbcL) gene, partial cds; chloroplast

TAACTTAACAGCATCTATTATCGGAAACGTATTCGGATTTAAAGCAGTTAAATGTCTTCGTTTAGAAGATATGCGTCTTCCTTATGCTTATCTAAAAACATTTATCGGGCCAGCAGCAGGTGTAATCGTTGAACGTGAAAGATTAGACGTTTTTGGTCGTCCACTTTTAGGAGCTACTGTAAAACCGAAATTAGGTCTTTCTGGTAAAAACTATGGTCGTGTAGTTTATGAAGGATTACGTGGTGGTTTAGATTTCTTAAAAGATGATGAAAACATCAACTCACAACCATTCATGAGATGGCGTGAAAG

>gi|125616956|gb|EF165173.1| Ochromonas sp. ACOI-1258 ribulose-1,5-bisphosphate carboxylase/oxygenase large subunit (rbcL) gene, partial cds; chloroplast

AAACTTAACTGCTTCTATTATAGGAAACGTTTTTGGTTTTAAAGCTGTTAAATGTTTAAGACTTGAAGATATGAGAATTCCTTATGCATATTTAAAAACATTCATAGGTCCAGCAACTGGAGTTATCGTTGAACGTGAAAGATTAGATGTTTTTGGTAGACCTTTATTAGGGGCTACAGTTAAACCAAAATTAGGTTTATCTGGAAAAAACTATGGTCGTGTTGTTTATGAAGGATTAAGAGGTGGATTAGATTTCTTAAAAGATGATGAAAATATTAACTCACAACCATTCATGAGATGGCGTGAAAG

>gi|125616958|gb|EF165174.1| Ochromonas sp. CCMP1278 ribulose-1,5-bisphosphate carboxylase/oxygenase large subunit (rbcL) gene, partial cds; chloroplast

TAACTTAACAGCATCTATTATCGGTAACGTTTTTGGATTCAAAGCCGTAAAATGTCTTCGTTTAGAAGATATGCGTCTTCCTTATGCTTACTTAAAAACTTTCATTGGACCAGCATCTGGAGTTATTGTTGAACGTGAAAGACTTGATATTTTTGGTCGTCCACTTTTAGGAGCTACAGTAAAACCGAAATTAGGTCTTTCTGGTAAAAACTATGGACGTGTTGTATACGAAGGATTAAAAGGTGGATTAGACTTCTTAAAAGATGACGAAAACATCAATTCACAACCATTCATGAGATGGCGTGAAAG

>gi|125616960|gb|EF165175.1| Ochromonas sp. aestuarii strain AC24 ribulose-1,5-bisphosphate carboxylase/oxygenase large subunit (rbcL) gene, partial cds; chloroplast

TAACTTAACAGCATCTATTATCGGTAACGTTTTTGGATTCAAAGCCGTAAAATGTCTTCGTTTAGAAGATATGCGTCTTCCTTATGCTTACTTAAAAACTTTCATTGGACCTGCGTCTGGAGTTATCGTTGAACGTGAAAGACTTGACGTTTTCGGTCGTCCACTTTTAGGAGCTACAGTAAAACCTAAATTAGGTCTTTCTGGTAAAAACTATGGTCGTGTTGTATATGAAGGATTAAAAGGTGGATTAGACTTCTTAAAAGATGACGAAAACATCAACTCACAACCATTCATGAGATGGCGTGAAAG

>gi|125616902|gb|EF185314.1| Ochromonas sp. CCMP1393 ribulose-1,5-bisphosphate carboxylase/oxygenase large subunit (rbcL) gene, partial cds; chloroplast

TAACCTTACTGCATCTATTATCGGTAACGTATTTGGTTTCAAAGCGGTTAAAGCTCTTCGTTTAGAAGATATGCGTTTACCATATGCATACTTAAAAACTTTCATCGGACCAGCTTCTGGTGTTATCGTAGAACGTGAAAGACTTGATGTTTTTGGACGTCCTCTTTTAGGTGCTACAGTTAAGCCAAAATTAGGTCTTTCTGGTAAAAACTACGGTCGTGTAGTATATGAAGGTTTAAGAGGTGGTTTAGATTTCTTAAAAGATGATGAGAATATTAACTCACAACCATTCATGAGATGGCGTGAAAG

>gi|125616904|gb|EF185315.1| Ochromonas tuberculata strain CCMP1861 ribulose-1,5-bisphosphate carboxylase/oxygenase large subunit (rbcL) gene, partial cds; chloroplast

TAACTTAACAGCATCTATCATCGGTAACGTTTTTGGATTCAAAGCCGTAAAATGTCTTCGTTTAGAAGATATGCGTATTCCTTATGCTTATTTAAAAACTTTCCAAGGTCCTGCTTCAGGAGTTATCGTAGAACGTGAAAGACTTGACGTTTTTGGACGTCCTTTGTTAGGAGCTACTGTTAAACCTAAATTAGGTCTTTCAGGTAAAAACTACGGTCGCGTAGTTTATGAAGGATTAAGAGGTGGTTTAGACTTTTTAAAAGATGACGAAAATATTAACTCACAACCATTCATGAGATGGCGTGAACG

>gi|125616980|gb|EF165185.1| Ochromonas sphaerocystis strain CCMP586 ribulose-1,5-bisphosphate carboxylase/oxygenase large subunit (rbcL) gene, partial cds; chloroplast

AAACTTAACAGCTTCTATCATTGGTAACGTTTTTGGATTTAAAGCTGTAAAAGCGCTGCGTTTAGAAGATATGCGTCTTCCATATGCATATTTAAAAACATTTATTGGTCCTGCATCTGGAGTAGTTGTAGAACGCGAAAGATTAGACGTATTTGGACGCCCTCTATTAGGAGCAACTGTAAAACCTAAGTTAGGTTTATCTGGAAAAAACTATGGTCGTGTAGTATATGAAGGATTACGTGGTGGATTAGATTTCTTAAAAGACGATGAAAATATTAATTCACAACCATTCATGAGATGGCGTGAAAG

>gi|125616982|gb|EF165186.1| Ochromonas cf. sphaerocystis strain CCMP2061 ribulose-1,5-bisphosphate carboxylase/oxygenase large subunit (rbcL) gene, partial cds; chloroplast

AAACTTAACAGCTTCTATCATTGGTAACGTTTTTGGATTTAAAGCTGTAAAAGCGCTGCGTTTAGAAGATATGCGTCTTCCATATGCATATTTAAAAACATTTATTGGTCCTGCATCTGGAGTAGTTGTAGAACGCGAAAGATTAGACGTATTTGGACGCCCTCTATTAGGAGCAACTGTAAAACCTAAGTTAGGTTTATCTGGAAAAAACTATGGTCGTGTAGTATATGAAGGATTACGTGGTGGATTAGATTTCTTAAAAGACGATGAAAATATTAATTCACAACCATTCATGAGATGGCGTGAAAG

>gi|125616984|gb|EF165187.1| Ochromonas perlata strain CCMP2732 ribulose-1,5-bisphosphate carboxylase/oxygenase large subunit (rbcL) gene, partial cds; chloroplast

AAACTTAACAGCTTCTATCATTGGTAACGTTTTCGGATTTAAAGCTGTAAAAGCGCTTCGTTTAGAAGATATGCGCCTTCCATATGCATATTTAAAAACATTTATTGGTCCTGCATCTGGAGTAGTTGTAGAACGCGAAAGATTAGACGTATTTGGACGCCCTTTATTAGGAGCAACTGTAAAACCTAAGTTAGGTTTATCTGGAAAAAACTATGGTCGTGTAGTATATGAAGGATTACGTGGTGGATTAGATTTCTTAAAAGATGATGAAAATATTAATTCACAACCATTCATGAGATGGCGTGAAAG

>gi|323690598|gb|GU935657.1| Ochromonas danica strain SAG 933.7 ribulose-1,5-bisphosphate carboxylase/oxygenase large subunit (rbcL) gene, partial cds; chloroplast

AAATTTAACAGCTTCAATTATTGGTAATGTATTTGGATTTAAAGCAGTAAAAGCTCTTCGTCTAGAAGATATGCGTCTTCCTTATGCTTATTTAAAAACATTTATTGGTCCTGCTTCTGGTGTTGTTGTAGAACGTGAAAGATTAGATGTATTTGGACGTCCTCTTTTAGGTGCAACAGTAAAACCTAAATTAGGTTTATCAGGAAAAAATTATGGACGTGTTGTTTATGAAGGATTAAAAGGTGGACTTGATTTTTTAAAAGATGATGAAAATATAAATTCTCAACCATTCATGAGATGGCGTGAAAG

>gi|323690600|gb|GU935658.1| Ochromonas sp. SAG 933.10 ribulose-1,5-bisphosphate carboxylase/oxygenase large subunit (rbcL) gene, partial cds; chloroplast

AAACTTAACGGCTTCAATTATTGGAAATGTTTTTGGATTTAAAGCTGTTAAAGCTCTTCGTTTAGAAGATATGCGTATTCCATATGCTTATTTAAAAACATTTTTAGGTCCTGCATCAGGAGTAATTGTAGAACGTGAACGTTTAGATGTCTTTGGTCGTCCTTTATTAGGAGCAACAGTAAAACCTAAATTAGGATTATCTGGAAAAAACTATGGGCGTGTAGTTTATGAAGGATTAAGAGGCGGTTTAGATTTTTTAAAAGATGATGAGAATATAAATTCTCAACCTTTTATGAGATGGCGTGAACG

>gi|354779689|gb|HQ710598.1| Ochromonas tuberculata culture CCMP:1861 ribulose-1,5-bisphosphate carboxylase/oxygenase large subunit (rbcL) gene, partial cds; plastid

TAACTTAACAGCATCTATCATCGGTAACGTTTTTGGATTCAAAGCCGTAAAATGTCTTCGTTTAGAAGATATGCGTATTCCTTATGCTTATTTAAAAACTTTCCAAGGTCCTGCTTCAGGAGTTATCGTAGAACGTGAAAGACTTGACGTTTTTGGACGTCCTTTGTTAGGAGCTACTGTTAAACCTAAATTAGGTCTTTCAGGTAAAAACTACGGTCGCGTAGTTTATGAAGGATTAAGAGGTGGTTTAGACTTTTTAAAAGATGACGAAAATATTAACTCACAACCATTCATGAGATGGCGTGAACG

>gi|125617012|gb|EF165201.1| Ochromonas sp. CCMP592 ribulose-1,5-bisphosphate carboxylase/oxygenase large subunit (rbcL) gene, partial cds; chloroplast

TAACTTAACAGCATCTATTATTGGTAATGTATTTGGATTTAAAGCAGTTAAAGCATTACGTTTAGAAGATATGCGTATGCCATATGCATACTTAAAAACTTTCTTAGGACCAGCGTCTGGTGTTATCGTAGAACGTGAAAGAATGGATGTTTTTGGACGCCCATTATTAGGTGCTACAGTAAAACCAAAATTAGGTTTATCTGGTAAAAACTATGGTCGTGTAGTTTATGAAGGTTTAGCTGGTGGTTTAGACTTCTTAAAAGATGACGAAAACATTAACTCGCAACCATTCATGAGATGGCGTGAAAG

>gi|125617014|gb|EF165202.1| Ochromonas sp. CCMP1149 ribulose-1,5-bisphosphate carboxylase/oxygenase large subunit (rbcL) gene, partial cds; chloroplast

AAACTTAACAGCATCGATTATTGGTAACGTATTTGGTTTCAAAGCAGTAAAATCTTTACGTTTAGAAGATATGCGTTTACCTTATGCATACTTAAAAACATTCATTGGACCAGCATCTGGTGTTGTTGTAGAACGTGAAAGACTTGATGTATTTGGACGTCCTCTTTTAGGAGCTACAGTAAAACCTAAGTTAGGTCTTTCTGGTAAAAACTATGGTCGTGTAGTATACGAAGGATTACGTGGTGGACTAGATTTCTTAAAGGATGATGAAAACATTAACTCACAACCATTTATGCGTTGGCGTGAGCG

>gi|125617016|gb|EF165203.1| Ochromonas marina strain AC22 ribulose-1,5-bisphosphate carboxylase/oxygenase large subunit (rbcL) gene, partial cds; chloroplast

AAACTTAACGGCTTCAATTATCGGAAACGTATTTGGATTTAAAGCTGTAAAAGCATTACGTTTAGAAGATATGCGTATTCCTTATGCATACTTAAAAACATTCCAAGGTCCTGCGACTGGAGTAGTTGTAGAACGTGAAAGATTAGATATTTTTGGACGTCCTTTTTTAGGAGCTACTGTAAAACCAAAATTAGGTCTTTCAGGTAAAAACTATGGTCGTGTAGTTTATGAAGGATTACAAGGTGGTTTAGACTTCTTAAAAGATGATGAGAATATTAACTCACAACCATTTATGCGTTGGCGTGAAAG

>gi|125616950|gb|EF165170.1| Ochromonas cf. gloeopara strain CCMP2718 ribulose-1,5-bisphosphate carboxylase/oxygenase large subunit (rbcL) gene, partial cds; chloroplast

CAATTTAACAGCATCGATTATCGGAAACGTTTTTGGTTTTAAAGCTGTAAAATGTTTACGACTTGAAGATATGAGAATTCCTTACGCTTATCTAAAAACATTTATAGGTCCAGCAACGGGAGTTATCGTTGAACGCGAACGATTAGATGTTTTTGGTAGACCTCTTTTAGGAGCTACGGTTAAGCCAAAATTAGGTTTATCTGGAAAAAACTATGGACGTGTTGTTTATGAGGGTTTACGAGGTGGATTAGATTTTTTAAAAGATGATGAAAATATCAACTCGCAACCATTCATGAGATGGCGTGAAAG

>gi|125616928|gb|EF165159.1| Ochromonas sp. CCMP1899 ribulose-1,5-bisphosphate carboxylase/oxygenase large subunit (rbcL) gene, partial cds; chloroplast

TAACTTAACTGCTTCTATTATTGGAAACGTTTTTGGATTTAAAGCTGTAAAGTGTCTTCGTTTAGAAGATATGCGTATGCCTTATGCTTACTTAAAGACATTCTTAGGACCGGCTGCAGGTGTTATTGTAGAACGTGAAAGACTTGACGTTTTTGGACGTCCTCTATTAGGTGCAACTGTTAAGCCTAAGTTAGGACTTTCTGGTAAAAACTATGGACGTGTAGTTTATGAAGGATTAAAAGGTGGTTTAGACTTCTTAAAGGATGACGAGAACATTAACTCTCAACCATTTATGCGTTGGCGTGAAAG

>gi|125616952|gb|EF165171.1| Ochromonas cf. gloeopara strain CCMP2060 ribulose-1,5-bisphosphate carboxylase/oxygenase large subunit (rbcL) gene, partial cds; chloroplast

CAATTTAACAGCATCGATTATCGGAAACGTTTTTGGTTTTAAAGCTGTAAAATGTTTACGACTTGAAGATATGAGAATTCCTTACGCTTATCTAAAAACATTTATAGGTCCAGCAACGGGAGTTATCGTTGAACGCGAACGATTAGATGTTTTTGGTAGACCTCTTTTAGGAGCTACGGTTAAGCCAAAATTAGGTTTATCTGGAAAAAACTATGGACGTGTTGTTTATGAGGGTTTACGAGGTGGATTAGATTTTTTAAAAGATGATGAAAATATCAACTCGCAACCATTCATGAGATGGCGTGAAAG

>gi|125616946|gb|EF165168.1| Ochromonas sp. MCIB-10896 ribulose-1,5-bisphosphate carboxylase/oxygenase large subunit (rbcL) gene, partial cds; chloroplast

TAACTTAACTGCATCTATCATTGGTAACGTTTTTGGATTTAAAGCCGTTAAATGTTTACGTTTAGAAGATATGCGTTTACCTTATGCATACTTAAAAACATTCATTGGTCCAGCTTCTGGGGTAATCGTAGAACGTGAAAGACTTGATGTATTCGGACGTCCATTATTAGGAGCTACAGTAAAACCAAAATTAGGTCTTTCTGGTAAAAACTACGGACGTGTTGTATATGAAGGATTACGTGGTGGATTAGACTTCTTAAAAGATGACGAAAATATTAACTCTCAACCATTCATGAGATGGCGTGAAAG

>gi|283099386|gb|GU325411.1| Ochromonas tuberculata strain CCMP1861 ribulose-1,5-bisphosphate carboxylase/oxygenase large subunit (rbcL) gene, partial cds; plastid

TAACTTAACAGCATCTATCATCGGTAACGTTTTTGGATTCAAAGCCGTAAAATGTCTTCGTTTAGAAGATATGCGTATTCCTTATGCTTATTTAAAAACTTTCCAAGGTCCTGCTTCAGGAGTTATCGTAGAACGTGAAAGACTTGACGTTTTTGGACGTCCTTTGTTAGGAGCTACTGTTAAACCTAAATTAGGTCTTTCAGGTAAAAACTACGGTCGCGTAGTTTATGAAGGATTAAGAGGTGGTTTAGACTTTTTAAAAGATGACGAAAATATTAACTCACAACCATTCATGAGATGGCGTGAACG

>gi|125616900|gb|EF185313.1| Chrysosphaerella sp. winter ribulose-1,5-bisphosphate carboxylase/oxygenase large subunit (rbcL) gene, partial cds; chloroplast

TAATTTAACAGCATCTATTATTGGAAACGTATTTGGATTTAAAGCGGTTAAATGTCTTCGTTTAGAAGATATGCGCATTCCTTATGCTTACTTAAAAACGTTCCAAGGACCTGCTTCTGGGGTTATTGTAGAACGTGAAAGATTAGATGTATTTGGACGTCCTTTATTAGGAGCTACTGTTAAACCTAAATTAGGTCTTTCTGGAAAAAATTATGGTCGTGTTGTTTATGAAGGATTAAAAGGTGGTTTAGATTTCTTAAAAGATGATGAAAATATCAACTCTCAACCTTTTATGAGATGGAGAGAAAG

>gi|125616906|gb|EF165148.1| Chrysocapsa vernalis strain CCMP278 ribulose-1,5-bisphosphate carboxylase/oxygenase large subunit (rbcL) gene, partial cds; chloroplast

TAACTTAACAGCATCTATTATTGGGAACGTATTTGGTTTCAAAGCTGTAAAATGTCTTCGTTTAGAAGATATGCGTATTCCTTTTGCCTACTTAAAAACTTTCATTGGTCCTGCTGCTGGAGTTATTGTAGAACGTGAAAGACTTGACGTATTTGGACGTCCTTTATTAGGAGCAACTGTAAAACCTAAATTAGGTCTTTCAGGAAAAAACTATGGTCGTGTAGTTTATGAAGGATTAAAAGGTGGTTTAGACTTCTTAAAAGATGACGAAAATATTAACTCTCAACCATTCATGAGATGGCGTGAAAG

>gi|125616908|gb|EF165149.1| Chrysocapsa paludosa strain CCMP380 ribulose-1,5-bisphosphate carboxylase/oxygenase large subunit (rbcL) gene, partial cds; chloroplast

TAACTTAACAGCATCTATTATTGGGAACGTATTTGGTTTCAAAGCGGTAAAATGTCTTCGTTTAGAAGATATGCGTATTCCTTTTGCATATTTAAAAACATTCATTGGTCCAGCTGCTGGAGTTATTGTAGAACGTGAAAGACTTGACGTATTTGGACGTCCTTTATTAGGAGCAACTGTAAAACCAAAATTAGGTCTTTCAGGAAAAAACTATGGTCGTGTAGTTTATGAAGGATTAAAAGGTGGTTTAGACTTCTTAAAAGATGACGAAAACATTAACTCTCAACCATTCATGAGATGGCGTGAAAG

>gi|125616910|gb|EF165150.1| Chrysonebula flava strain CCMP2765 ribulose-1,5-bisphosphate carboxylase/oxygenase large subunit (rbcL) gene, partial cds; chloroplast

AAACTTAACAGCATCTATTATCGGTAACGTATTCGGTTTCAAAGCCGTTAAATGTCTTCGTTTAGAAGATATGCGTATTCCTTATGCTTACTTAAAAACTTTCATTGGTCCAGCTGCTGGAGTTATTGTAGAACGTGAAAGACTTGACGTATTCGGACGCCCATTATTAGGAGCAACTGTTAAACCAAAATTAGGTCTTTCTGGTAAAAACTACGGTCGTGTAGTTTATGAAGGATTAAAAGGTGGTTTAGACTTCTTAAAAGATGACGAAAATATTAACTCTCAACCATTCATGAGATGGCGTGAAAG

>gi|125616912|gb|EF165151.1| Chromulina sp. SAG 17.97 strain SAG17.97 ribulose-1,5-bisphosphate carboxylase/oxygenase large subunit (rbcL) gene, partial cds; chloroplast

TAACTTAACAGCATCTATTATCGGTAACGTATTTGGTTTCAAAGCAGTTAAATGTCTTCGTTTAGAAGATATGCGTATTCCTTATGCTTACTTAAAAACTTTCATTGGTCCAGCTGCTGGAGTTATTGTAGAACGTGAAAGACTTGACGTATTCGGACGTCCTTTATTAGGAGCAACTGTTAAACCTAAATTAGGTCTTTCTGGTAAAAACTACGGACGTGTAGTTTACGAAGGATTAAAAGGTGGTTTAGACTTCTTAAAAGATGACGAAAACATCAACTCTCAACCATTCATGAGATGGCGTGAAAG

>gi|125616914|gb|EF165152.1| Kremastochrysopsis americana strain CCMP260 ribulose-1,5-bisphosphate carboxylase/oxygenase large subunit (rbcL) gene, partial cds; chloroplast

TAACTTAACAGCATCTATTATCGGAAACGTATTTGGTTTCAAAGCCGTTAAATGTCTTCGTTTAGAAGATATGCGTATTCCTTATGCTTACTTAAAAACTTTCATTGGTCCAGCTGCTGGAGTAATTGTAGAACGTGAAAGACTTGACGTATTTGGACGTCCTTTATTAGGAGCAACAGTTAAACCGAAATTAGGTCTTTCTGGTAAAAACTATGGTCGTGTAGTTTACGAAGGATTAAAAGGTGGTTTAGACTTCTTAAAAGATGACGAAAACATCAACTCTCAACCATTCATGAGATGGCGTGAAAG

>gi|125616916|gb|EF165153.1| Chrysocapsa sp. UTCC280 ribulose-1,5-bisphosphate carboxylase/oxygenase large subunit (rbcL) gene, partial cds; chloroplast

TAACTTAACAGCATCAATTATCGGGAACGTATTTGGTTTTAAAGCAGTTAAATGTCTTCGTTTAGAAGATATGCGTATTCCTTATGCTTACTTAAAAACATTCATTGGTCCAGCTGCTGGAGTTATTGTAGAACGTGAAAGACTTGATGTTTTTGGACGTCCATTATTAGGAGCAACAGTTAAACCAAAATTAGGTCTTTCTGGTAAAAACTATGGTCGTGTAGTTTATGAAGGATTAAAAGGTGGTTTAGATTTCTTAAAAGATGACGAAAATATTAACTCTCAACCATTCATGAGATGGCGTGAAAG

>gi|125616918|gb|EF165154.1| Naegeliella flagellifera strain CCMP280 ribulose-1,5-bisphosphate carboxylase/oxygenase large subunit (rbcL) gene, partial cds; chloroplast

TAACTTAACAGCGTCTATCATCGGTAACGTATTCGGATTTAAAGCAGTTAAATGTCTTCGACTAGAAGATATGCGTATTCCTTATGCTTATTTAAAAACATTCATTGGTCCTGCAGCTGGAGTTATTGTAGAACGTGAAAGACTTGACGTATTCGGACGTCCTCTTTTAGGAGCAACAGTTAAACCTAAATTAGGTCTTTCTGGTAAAAACTATGGTCGTGTAGTTTATGAAGGATTAAAAGGTGGTTTAGACTTCTTAAAAGACGATGAAAATATTAACTCTCAACCATTCATGAGATGGCGTGAAAG

>gi|125616920|gb|EF165155.1| Epipyxis aureus strain CCMP385 ribulose-1,5-bisphosphate carboxylase/oxygenase large subunit (rbcL) gene, partial cds; chloroplast

GAACTTAACTGCATCTATTATTGGTAACGTTTTTGGATTTAAAGCTGTTAAATGTCTTCGTTTAGAAGATATGCGTCTTCCTTATGCTTACTTAAAAACATTCATTGGGCCAGCTTCTGGAGTTATCGTAGAACGTGAAAGACTTGACGTTTTTGGTCGTCCATTATTAGGAGCAACTGTAAAACCGAAATTAGGTCTTTCAGGAAAAAACTATGGTCGTGTAGTTTATGAAGGATTAAGAGGTGGTTTAGACTTCTTAAAAGATGACGAAAACATTAACTCTCAACCATTCATGAGATGGCGTGAAAG

>gi|125616922|gb|EF165156.1| Dinobryon sociale var. americana strain CCMP1860 ribulose-1,5-bisphosphate carboxylase/oxygenase large subunit (rbcL) gene, partial cds; chloroplast

AAACTTAACAGCATCAATCATTGGAAACGTTTTTGGATTTAAAGCGGTTAAAGCTCTTCGTTTAGAAGATATGCGTCTTCCGTATGCATATTTAAAAACATTCTTAGGACCTGCATCTGGTGTTATTGTAGAACGTGAAAGACTTGACGTTTTTGGTCGTCCACTTTTAGGTGCAACTGTTAAACCAAAATTAGGTCTTTCAGGGAAAAACTATGGTCGTGTTGTTTATGAAGGATTAAAAGGTGGTTTAGACTTCTTAAAAGATGACGAAAACATTAACTCTCAACCATTCATGAGATGGCGTGAAAG

>gi|125616924|gb|EF165157.1| Dinobryon cylindricum strain CCMP2766 ribulose-1,5-bisphosphate carboxylase/oxygenase large subunit (rbcL) gene, partial cds; chloroplast

AAACTTAACAGCATCAATCATTGGAAACGTTTTTGGATTTAAAGCGGTTAAAGCTCTTCGTTTAGAAGATATGCGTCTTCCGTATGCATATTTAAAAACATTCTTAGGACCTGCATCTGGTGTTATTGTAGAACGTGAAAGACTTGACGTTTTTGGACGTCCACTTTTAGGTGCAACTGTTAAACCAAAATTAGGTCTTTCAGGGAAAAACTATGGTCGTGTTGTTTATGAAGGATTAAAAGGTGGTTTAGACTTCTTAAAAGATGACGAAAACATTAACTCTCAACCATTCATGAGATGGCGTGAAAG

>gi|125616926|gb|EF165158.1| Dinobryon sociale strain UTCC392 ribulose-1,5-bisphosphate carboxylase/oxygenase large subunit (rbcL) gene, partial cds; chloroplast

AAACTTAACAGCATCAATCATTGGAAACGTTTTTGGATTTAAAGCAGTTAAAGCTCTTCGTTTAGAAGATATGCGTCTGCCATATGCATATTTAAAAACATTCTTAGGACCTGCGTCTGGTGTTATTGTAGAACGTGAAAGACTTGACGTTTTTGGACGTCCACTTTTAGGTGCAACTGTTAAACCAAAATTAGGTCTTTCAGGGAAAAACTATGGTCGTGTTGTTTATGAAGGATTAAAAGGTGGTTTAGACTTCTTAAAAGATGACGAAAATATTAACTCTCAACCATTCATGAGATGGCGTGAAAG

>gi|125616930|gb|EF165160.1| Phaeoplaca thallosa strain CCMP634 ribulose-1,5-bisphosphate carboxylase/oxygenase large subunit (rbcL) gene, partial cds; chloroplast

TAATTTAACTGCATCAATTATTGGAAATGTATTTGGATTTAAAGCGGTTAAATGTCTTCGTTTAGAAGATATGCGTATTCCTTATGCTTATTTAAAAACATTCTTAGGCCCAGCTGCTGGAGTTATTGTAGAACGTGAAAGACTTGATGTATTTGGTCGTCCGTTACTTGGTGCTACTGTAAAACCTAAATTAGGTCTTTCTGGTAAAAACTACGGTCGTGTTGTTTATGAAGGATTAAAAGGTGGATTAGACTTTTTAAAAGATGACGAAAATATTAATTCTCAACCATTCATGAGATGGCGTGAAAG

>gi|125616932|gb|EF165161.1| Lagynion cf. ampullaceum strain CCMP2727 ribulose-1,5-bisphosphate carboxylase/oxygenase large subunit (rbcL) gene, partial cds; chloroplast

TAACTTAACAGCATCTATTATTGGAAACGTATTTGGATTTAAAGCTGTAAAAGCTCTTCGTTTAGAAGATATGCGTTTACCTTATGCTTATCTAAAAACATTCCAAGGACCTGCTGCGGGTGTTATTGTTGAACGTGAAAGATTAGATGTTTTTGGTCGTCCATTATTAGGAGCAACAGTTAAACCGAAATTAGGTCTTTCTGGTAAAAACTATGGTCGAGTAGTTTATGAAGGATTAAAAGGTGGTTTAGATTTCTTAAAAGATGACGAAAATATTAACTCTCAACCATTCATGAGATGGCGTGAGCG

>gi|125616934|gb|EF165162.1| Lagynion scherffelii strain CCMP465 ribulose-1,5-bisphosphate carboxylase/oxygenase large subunit (rbcL) gene, partial cds; chloroplast

TAACTTAACAGCCTCTATTATTGGAAACGTATTTGGATTTAAAGCAGTAAAAGCTCTTCGTTTAGAAGATATGCGTTTACCTTATGCTTATTTAAAAACATTCCAAGGACCTGCAGCAGGTGTTATCGTTGAACGTGAAAGACTTGATGTTTTTGGACGTCCATTATTAGGAGCAACAGTTAAACCTAAATTAGGTCTTTCTGGTAAAAATTATGGTCGAGTAGTTTATGAAGGATTAAAAGGTGGTTTAGATTTCTTAAAAGATGACGAAAATATTAATTCTCAACCATTCATGAGATGGCGTGAACG

>gi|125616936|gb|EF165163.1| Chrysophyceae sp. CCMP1161 ribulose-1,5-bisphosphate carboxylase/oxygenase large subunit (rbcL) gene, partial cds; chloroplast

GAACTTAACAGCATCTATTATTGGAAACGTATTTGGGTTCAAAGCTGTAAAATGTCTTCGTTTAGAAGATATGCGTTTACCATATGCTTACTTAAAAACATTCCAAGGTCCTGCTGCCGGAGTTATTGTTGAACGTGAAAGACTTGACGTTTTTGGACGACCTCTATTAGGTGCTACAGTAAAACCAAAATTAGGTCTTTCTGGAAAAAATTATGGTCGTGTAGTTTATGAAGGATTAAAAGGTGGTTTAGATTTCTTAAAAGATGACGAAAACATTAACTCTCAACCATTCATGAGATGGCGTGAACG

>gi|125616938|gb|EF165164.1| Chromophyton cf. rosanoffii strain CCMP2751 ribulose-1,5-bisphosphate carboxylase/oxygenase large subunit (rbcL) gene, partial cds; chloroplast

TAACTTAACAGCGTCTATCATTGGTAACGTATTTGGATTCAAAGCCGTTAAATGTCTTCGTTTAGAAGATATGCGTCTTCCATATGCTTACTTAAAAACTTTCCAAGGTCCTGCTGCAGGAGTTGTTGTAGAACGTGAAAGACTTGACGTTTTTGGACGTCCATTATTAGGTGCTACAGTAAAACCAAAATTAGGTCTTTCTGGTAAAAACTACGGACGTGTAGTTTACGAAGGATTAAGAGGTGGTTTAGACTTCTTAAAAGATGACGAAAACATTAACTCTCAACCATTCATGAGATGGCGTGAACG

>gi|125616940|gb|EF165165.1| Chromophyton cf. rosanoffii strain CCMP2753 ribulose-1,5-bisphosphate carboxylase/oxygenase large subunit (rbcL) gene, partial cds; chloroplast

TAACTTAACAGCGTCTATCATTGGTAACGTATTTGGATTTAAAGCCGTTAAATGTCTTCGTTTAGAAGATATGCGTCTTCCATACGCGTACTTAAAAACTTTCCAAGGTCCTGCTGCAGGGGTTATTGTAGAACGTGAAAGACTTGACGTTTTTGGACGTCCATTATTAGGTGCAACAGTAAAACCAAAATTAGGTCTTTCTGGTAAAAACTATGGTCGTGTAGTTTACGAAGGATTAAGAGGTGGTTTAGACTTCTTAAAAGATGACGAAAACATCAACTCTCAACCATTCATGAGATGGCGTGAACG

>gi|125616942|gb|EF165166.1| Chrysosaccus sp. CCMP368 ribulose-1,5-bisphosphate carboxylase/oxygenase large subunit (rbcL) gene, partial cds; chloroplast

TAATTTAACAGCATCTATTATTGGTAACGTATTTGGATTCAAAGCCGTTAAATGTCTTCGTTTAGAAGATATGCGTCTTCCTTATGCTTACTTAAAAACTTTCCAAGGTCCTGCTGCAGGTGTTATTGTAGAACGTGAAAGACTTGATGTTTTTGGACGTCCATTATTAGGTGCAACAGTTAAACCAAAATTAGGTCTTTCTGGTAAAAACTACGGTCGTGTAGTTTATGAAGGATTAAAAGGTGGTTTAGACTTCTTAAAAGACGATGAAAACATTAACTCTCAACCATTCATGAGATGGCGTGAACG

>gi|125616944|gb|EF165167.1| Chrysosaccus sp. CCMP1156 ribulose-1,5-bisphosphate carboxylase/oxygenase large subunit (rbcL) gene, partial cds; chloroplast

TAATTTAACAGCATCTATTATTGGTAACGTATTTGGATTCAAAGCCGTTAAATGTCTTCGTTTAGAAGATATGCGTCTTCCTTATGCTTACTTAAAAACTTTCCAAGGTCCTGCTGCAGGTGTTATTGTAGAACGTGAAAGACTTGATGTTTTTGGACGTCCATTATTAGGTGCAACAGTTAAACCAAAATTAGGTCTTTCTGGTAAAAACTACGGTCGTGTAGTTTATGAAGGATTAAAAGGTGGTTTAGACTTCTTAAAAGACGATGAAAACATTAACTCTCAACCATTCATGAGATGGCGTGAACG

>gi|125616948|gb|EF165169.1| Poterioochromonas malhamensis strain SAG933.1c ribulose-1,5-bisphosphate carboxylase/oxygenase large subunit (rbcL) gene, partial cds; chloroplast

AAATTTAACAGCTTCTATTATCGGAAATGTCTTTGGTTTTAAGGCTGTTAAATGTTTAAGACTTGAAGATATGAGAATTCCATATGCTTACCTAAAAACATTTATAGGTCCAGCAACGGGAGTTATCGTTGAGCGTGAAAGATTAGATGTTTTTGGAAGACCTCTTTTAGGAGCGACAGTTAAACCAAAATTAGGCTTATCTGGGAAAAATTATGGTCGCGTTGTTTATGAAGGGTTACGAGGTGGATTAGATTTCTTAAAGGATGACGAAAATATTAATTCACAACCATTCATGAGATGGCGTGAAAG

>gi|125616954|gb|EF165172.1| Poterioochromonas stipitata strain CCMP1862 ribulose-1,5-bisphosphate carboxylase/oxygenase large subunit (rbcL) gene, partial cds; chloroplast

AAATTTAACAGCTTCTATTATAGGAAACGTTTTTGGTTTTAAAGCTGTTAAGTGCTTAAGACTTGAAGATATGAGAATTCCTTATGCATATTTAAAAACATTTATTGGTCCAGCAACAGGAGTTGTTGTTGAACGTGAAAGATTAGATGTTTTTGGTAGACCTTTATTAGGAGCTACAGTTAAACCAAAATTAGGTTTATCTGGAAAAAACTATGGTCGTGTTGTTTATGAAGGATTAAGAGGTGGTTTAGATTTCTTAAAAGATGATGAAAATATTAACTCACAACCATTCATGAGATGGCGTGAAAG

>gi|125616962|gb|EF165176.1| Chrysoxys sp. CCMP591 ribulose-1,5-bisphosphate carboxylase/oxygenase large subunit (rbcL) gene, partial cds; chloroplast

TAACTTAACTGCTTCTATTATTGGTAACGTTTTTGGTTTCAAAGCTGTTAAATGTCTTCGTTTAGAAGATATGAGACTTCCATATGCATACTTAAAAACTTTCATCGGACCAGCTGCTGGTGTAATTGTAGAACGTGAAAGACTTGACGTTTTTGGACGTCCTCTTTTAGGTGCAACAGTAAAACCAAAATTAGGTCTTTCTGGTAAGAACTATGGACGTGTAGTATATGAAGGTTTAAAAGGTGGTTTAGACTTCTTAAAAGATGACGAAAACATCAATTCACAACCATTTATGCGTTGGCGTGAAAG

>gi|125616964|gb|EF165177.1| Ochromonas distigma AC25 ribulose-1,5-bisphosphate carboxylase/oxygenase large subunit (rbcL) gene, partial cds; chloroplast

TAACTTAACAGCATCTATTATCGGAAACGTATTCGGATTTAAAGCAGTTAAATGTCTTCGTTTAGAAGATATGCGTCTTCCTTATGCTTATCTAAAAACATTTATCGGGCCAGCAGCAGGTGTAATCGTTGAACGTGAAAGATTAGACGTTTTTGGTCGTCCACTTTTAGGAGCTACTGTAAAACCGAAATTAGGTCTTTCTGGTAAAAACTATGGTCGTGTAGTTTATGAAGGATTACGTGGTGGTTTAGATTTCTTAAAAGATGATGAAAACATCAACTCACAACCATTCATGAGATGGCGTGAAAG

>gi|125616966|gb|EF165178.1| Uroglenopsis americana strain CCMP2769 ribulose-1,5-bisphosphate carboxylase/oxygenase large subunit (rbcL) gene, partial cds; chloroplast

TAACTTAACAGCATCTATTATTGGTAACGTATTTGGATTTAAAGCCGTAAAAGCTCTTAGATTAGAAGATATGCGTCTTCCATTTGCTTACTTAAAAACTTTCATTGGTCCTGCAGCTGGTGTTATTGTTGAACGTGAAAGACTTGATGTTTTCGGACGTCCTCTTTTAGGTGCTACTGTAAAACCGAAATTAGGTCTTTCTGGAAAAAACTATGGTCGTGTAGTTTATGAAGGATTAAAAGGTGGTTTAGATTTCTTAAAAGATGATGAAAACATCAACTCACAACCATTCATGAGATGGCGTGAAAG

>gi|125616968|gb|EF165179.1| Uroglenopsis americana strain CCMP1863 ribulose-1,5-bisphosphate carboxylase/oxygenase large subunit (rbcL) gene, partial cds; chloroplast

TAACTTAACAGCATCTATTATTGGTAACGTATTTGGATTTAAAGCCGTAAAAGCTCTTAGATTAGAAGATATGCGTCTTCCATTTGCTTACTTAAAAACTTTCATTGGTCCTGCAGCTGGTGTTATTGTTGAACGTGAAAGACTTGATGTTTTCGGACGTCCTCTTTTAGGTGCTACTGTAAAACCGAAATTAGGTCTTTCTGGAAAAAACTATGGTCGTGTAGTTTATGAAGGATTAAAAGGTGGTTTAGATTTCTTAAAAGATGATGAAAACATCAACTCACAACCATTCATGAGATGGCGTGAAAG

>gi|125616970|gb|EF165180.1| Chromulina cf. nebulosa strain CCMP2719 ribulose-1,5-bisphosphate carboxylase/oxygenase large subunit (rbcL) gene, partial cds; chloroplast

TAACTTAACAGCTTCAATTATCGGAAACGTATTCGGTTTCAAAGCTGTAAAAGCTCTTCGTTTAGAAGATATGCGTATTCCATACGGATATTTAAAAACTTTCTTAGGCCCTGCAACAGGAGTTGTTGTAGAACGTGAAAGACTTGACGTTTTTGGTCGTCCTTTATTAGGTGCTACAGTTAAACCAAAATTAGGTCTTTCTGGAAAAAACTATGGTCGTGTTGTATATGAAGGATTAAGAGGTGGTCTTGATTTCTTAAAAGATGATGAGAATATCAACTCTCAACCATTCATGAGATGGCGTGAAAG

>gi|125616972|gb|EF165181.1| Chrysamoeba tenera strain UTCC273 ribulose-1,5-bisphosphate carboxylase/oxygenase large subunit (rbcL) gene, partial cds; chloroplast

CAATTTAACAGCTTCTATTATTGGAAACGTATTTGGATTCAAAGCGGTTAAAGCACTTCGTTTAGAAGACATGCGCCTTCCATATGGATACTTAAAAACTTTCTTAGGACCTGCAACAGGAGTTATTGTAGAACGTGAAAGATTAGATGTTTTTGGTCGTCCTTTATTAGGAGCTACTGTAAAACCTAAGTTAGGTCTTTCTGGTAAGAATTATGGACGTGTTGTTTATGAAGGATTAAGAGGTGGTTTAGATTTCTTAAAAGATGACGAAAACATTAACTCTCAACCATTCATGAGATGGCGTGAAAG

>gi|125616974|gb|EF165182.1| Chrysamoeba mikrokonta strain CCMP1857 ribulose-1,5-bisphosphate carboxylase/oxygenase large subunit (rbcL) gene, partial cds; chloroplast

AAACTTAACAGCTTCGATTATTGGAAACGTTTTCGGATTTAAAGCTGTAAAAGCTCTTCGTTTAGAAGATATGCGCCTTCCATACGGATATTTAAAAACATTCATTGGGCCTGCAACTGGAGTAGTTGTAGAACGTGAAAGATTAGATGTTTTTGGTCGTCCATTATTAGGAGCAACAGTAAAACCAAAATTAGGTTTATCAGGAAAAAACTATGGTCGTGTTGTTTATGAAGGATTAAGAGGTGGTTTAGACTTCTTAAAAGACGATGAAAACATTAACTCTCAACCATTCATGAGATGGCGTGAAAG

>gi|125616976|gb|EF165183.1| Ochromonas sp. CCMP2767 ribulose-1,5-bisphosphate carboxylase/oxygenase large subunit (rbcL) gene, partial cds; chloroplast

CAATTTAACGGCTTCAATTATTGGAAATGTTTTTGGATTTAAAGCTGTTAAAGCTCTTCGTTTAGAAGATATGCGTATACCATATGCTTATTTAAAAACATTCTTAGGTCCTGCATCAGGGGTAGTTGTAGAACGTGAACGTTTAGATGTTTTTGGTCGTCCTTTATTAGGAGCAACAGTAAAACCTAAATTAGGATTATCAGGAAAAAACTATGGACGTGTAGTTTATGAGGGATTAAGAGGCGGTTTAGATTTTTTAAAAGATGATGAAAATATTAATTCTCAACCTTTTATGAGATGGCGTGAAAG

>gi|125616978|gb|EF165184.1| Poteriospumella vasocystis strain CCMP2741 ribulose-1,5-bisphosphate carboxylase/oxygenase large subunit (rbcL) gene, partial cds; chloroplast

AAACTTAACGGCTTCAATTATTGGAAATGTTTTTGGATTTAAAGCTGTTAAAGCTCTTCGTTTAGAAGATATGCGTATTCCATATGCTTATTTAAAAACATTTTTAGGTCCTGCATCAGGAGTAATTGTAGAACGTGAACGTTTAGATGTCTTTGGTCGTCCTTTATTAGGAGCAACAGTAAAACCTAAATTAGGATTATCTGGAAAAAACTATGGGCGTGTAGTTTATGAAGGATTAAGAGGCGGTTTAGATTTTTTAAAAGATGATGAGAATATAAATTCTCAACCTTTTATGAGATGGCGTGAACG

>gi|125616988|gb|EF165189.1| Synura petersenii strain CCMP854 ribulose-1,5-bisphosphate carboxylase/oxygenase large subunit (rbcL) gene, partial cds; chloroplast

AAACTTAACAGCATCAATTATTGGAAACGTTTTCGGTTTTAAAGCTGTAAAATGTTTACGTTTAGAAGATATGCGTATTCCTTATGCTTATTTAAAAACGTTTATCGGACCTGCAACTGGAGTTATTGTAGAACGTGAAAGAATGGATGTATTTGGACGTCCTCTTTTAGGTGCAACTGTAAAACCAAAATTAGGTCTTTCTGGTAAAGCTTATGGTCGTGTAGTTTATGAAGGATTAAAAGGTGGTTTAGATTTCTTAAAAGACGATGAAAATATTAATTCACAACCATTCATGAGATGGCGTGAAAG

>gi|125616990|gb|EF165190.1| Synura petersenii strain CCMP857 ribulose-1,5-bisphosphate carboxylase/oxygenase large subunit (rbcL) gene, partial cds; chloroplast

AAACTTAACAGCATCAATTATTGGAAACGTTTTCGGTTTTAAAGCTGTAAAATGTTTACGTTTAGAAGATATGCGTATTCCGTATGCTTATTTAAAAACGTTTATCGGACCTGCAACTGGAGTTATTGTAGAACGTGAAAGAATGGATGTATTTGGACGTCCTCTTTTAGGTGCAACTGTAAAACCAAAATTAGGTCTTTCTGGTAAAGCTTATGGTCGTGTAGTTTATGAAGGATTAAAAGGTGGTTTAGATTTCTTAAAAGACGATGAAAATATTAATTCACAACCATTCATGAGATGGCGTGAAAG

>gi|125616992|gb|EF165191.1| Synura petersenii strain SAG24.86 ribulose-1,5-bisphosphate carboxylase/oxygenase large subunit (rbcL) gene, partial cds; chloroplast

AAACTTAACAGCATCAATTATTGGAAACGTTTTCGGTTTTAAAGCTGTAAAATGTTTACGTTTAGAAGATATGCGTATTCCGTATGCTTATTTAAAAACGTTTATCGGACCTGCAACTGGAGTTATTGTAGAACGTGAAAGAATGGATGTATTTGGACGTCCTCTTTTAGGTGCAACTGTAAAACCAAAATTAGGTCTTTCTGGTAAAGCTTATGGTCGTGTAGTTTATGAAGGATTAAAAGGTGGTTTAGATTTCTTAAAAGACGATGAAAATATTAATTCACAACCATTCATGAGATGGCGTGAAAG

>gi|125616986|gb|EF165188.1| Synura petersenii strain CCMP851 ribulose-1,5-bisphosphate carboxylase/oxygenase large subunit (rbcL) gene, partial cds; chloroplast

AAACTTAACAGCATCAATTATTGGAAACGTTTTCGGTTTTAAAGCTGTAAAATGTTTACGTTTAGAAGATATGCGTATTCCTTATGCTTATTTAAAAACGTTTATCGGACCTGCAACTGGAGTTATTGTAGAACGTGAAAGAATGGATGTATTTGGACGTCCTCTTTTAGGTGCAACTGTAAAACCAAAATTAGGTCTTTCTGGTAAAGCTTATGGTCGTGTAGTTTATGAAGGATTAAAAGGTGGTTTAGATTTCTTAAAAGACGATGAAAATATTAATTCACAACCATTCATGAGATGGCGTGAAAG

>gi|2375361750|gb|OP719269.1| Synura sphagnicola strain NIES 695 ribulose-1,5-biphosphate carboxylase/oxygenase large subunit (rbcL) gene, partial cds; chloroplast

CAATTTAACAGCTTCGATTATTGGAAACGTTTTTGGTTTTAAAGCTGTAAAATGTTTACGTTTAGAAGATATGCGTATTCCATATGCTTATTTAAAAACTTTCTTAGGCCCTGCAACAGGAGTAATCGTAGAACGTGAAAGAATGGATGTTTTTGGTCGTCCTCTTTTAGGTGCAACTGTAAAACCTAAATTAGGTCTTTCAGGAAAAAACTACGGTCGTGTTGTTTATGAAGGATTAAAAGGTGGATTAGATTTCTTAAAAGATGATGAAAATATTAATTCACAACCTTTTATGAGATGGCGTGAAAG

>gi|2375361752|gb|OP719270.1| Synura sphagnicola strain K35 ribulose-1,5-biphosphate carboxylase/oxygenase large subunit (rbcL) gene, partial cds; chloroplast

CAATTTAACAGCTTCGATTATTGGAAACGTTTTTGGTTTTAAAGCCGTAAAATGTTTACGTTTAGAAGATATGCGTATTCCATATTCTTATTTAAAAACTTTCTTAGGCCCTGCAACAGGAGTAATTGTAGAACGTGAAAGAATGGATGTTTTTGGTCGTCCTCTTTTAGGTGCAACTGTAAAACCTAAATTAGGTCTTTCAGGTAAAAACTATGGTCGTGTTGTTTATGAAGGTCTAAAAGGTGGATTAGATTTCTTAAAAGATGATGAAAATATTAATTCTCAACCTTTTATGAGATGGCGTGAAAG

>gi|2375361754|gb|OP719271.1| Synura sphagnicola strain M44 ribulose-1,5-biphosphate carboxylase/oxygenase large subunit (rbcL) gene, partial cds; chloroplast

CAATTTAACAGCTTCGATTATTGGAAACGTTTTTGGTTTTAAAGCCGTAAAATGTTTACGTTTAGAAGATATGCGTATTCCATATTCTTATTTAAAAACTTTCTTAGGCCCTGCAACAGGAGTAATTGTAGAACGTGAAAGAATGGATGTTTTTGGTCGTCCTCTTTTAGGTGCAACTGTAAAACCTAAATTAGGTCTTTCAGGTAAAAACTATGGTCGTGTTGTTTATGAAGGTCTAAAAGGTGGATTAGATTTCTTAAAAGATGATGAAAATATTAATTCTCAACCTTTTATGAGATGGCGTGAAAG

>gi|125616994|gb|EF165192.1| Synura uvella strain CCMP871 ribulose-1,5-bisphosphate carboxylase/oxygenase large subunit (rbcL) gene, partial cds; chloroplast

AAACTTAACTGCATCTATCATTGGAAACGTTTTTGGTTTCAAAGCTGTAAAATGTTTACGTTTAGAAGATATGCGTCTTCCATATGCTTACTTAAAAACTTTCTTAGGCCCTGCAACTGGAGTTATTGTAGAACGTGAAAGAATGGATGTATTTGGACGTCCTTTATTAGGAGCAACTGTTAAACCAAAACTAGGTCTTTCAGGAAAAAACTATGGTCGTGTTGTTTATGAAGGATTAAAAGGTGGTTTAGACTTCTTAAAAGATGACGAAAATATTAATTCACAACCTTTCATGAGATGGCGTGAAAG

>gi|125616996|gb|EF165193.1| Mallomonas annulata strain CCMP474 ribulose-1,5-bisphosphate carboxylase/oxygenase large subunit (rbcL) gene, partial cds; chloroplast

TAACTTAACAGCATCTATCATCGGAAACGTTTTTGGTTTCAAAGCCGTTAAAGCTTTACGTTTAGAAGATATGCGTATTCCTTATGCATACTTAAAAACTTTCCAAGGTCCAGCTACTGGAGTTATTGTTGAACGTGAAAGAATGGATGTATTTGGACGTCCTTTATTAGGAGCTACTGTTAAACCTAAATTAGGTCTTTCTGGTAAAAACTATGGTCGTGTAGTTTATGAAGGATTAAAAGGTGGTTTAGATTTCTTAAAAGATGATGAAAATATTAACTCACAACCATTCATGAGATGGCGTGAAAG

>gi|125616998|gb|EF165194.1| Mallomonas striata var. serrata strain CCMP2059 ribulose-1,5-bisphosphate carboxylase/oxygenase large subunit (rbcL) gene, partial cds; chloroplast

AAACTTAACAGCTTCTATCATCGGAAACGTTTTCGGATTTAAAGCAGTTAAAGCTTTACGTTTAGAAGATATGCGTATTCCTTTTGCTTACTTAAAAACTTTCCAAGGTCCTGCAACTGGAGTTGTTGTAGAACGTGAAAGAATGGATGTTTTTGGACGTCCTTTATTAGGAGCTACTGTTAAACCTAAATTAGGTCTTTCAGGAAAAAACTATGGTCGTGTAGTTTATGAAGGATTAAAAGGTGGTTTAGACTTTTTAAAAGATGATGAAAATATTAACTCACAACCATTCATGAGATGGCGTGAAAG

>gi|1003703568|gb|KT852946.1| Mallomonas paragrandis strain BOROK VN 827 ribulose-1,5-bisphosphate carboxylase/oxygenase large subunit (rbcL) gene, partial cds; plastid

GAACTTAACAGCTTCAATTATTGGAAACGTTTTTGGATTTAAAGCAGTAAAAGCTTTACGTTTAGAAGATATGCGTATTCCATATGCTTACCTAAAAACTTTCTTAGGACCAGCGACTGGAGTTATTGTTGAACGTGAAAGAATGGATGTATTTGGACGTCCATTATTAGGAGCAACTGTTAAACCAAAATTAGGTCTTTCAGGAAAAAATTATGGTCGTGTTGTTTATGAAGGATTAAAAGGTGGTTTAGACTTCTTAAAAGACGACGAAAATATCAATTCTCAAGCCTTCATGAGATGGCGTGAAAG

>gi|125617000|gb|EF165195.1| Mallomonas rasilis strain CCMP479 ribulose-1,5-bisphosphate carboxylase/oxygenase large subunit (rbcL) gene, partial cds; chloroplast

TAACTTAACAGCATCAATTATCGGAAACGTTTTTGGTTTCAAAGCTGTTAAAGCATTACGTTTAGAAGATATGCGTATTCCTTTTGCTTACCTAAAAACTTTCCAAGGTCCTGCAACAGGAGTTGTTGTAGAACGTGAAAGAATGGATGTTTTTGGACGTCCTTTATTAGGTGCTACTGTTAAACCAAAATTAGGTCTTTCTGGTAAAAACTATGGTCGTGTAGTTTATGAAGGATTAAAAGGTGGTTTAGATTTCTTAAAAGATGATGAAAATATTAATTCACAACCATTCATGAGATGGCGTGAAAG

>gi|125617006|gb|EF165198.1| Mallomonas insignis strain CCMP2549 ribulose-1,5-bisphosphate carboxylase/oxygenase large subunit (rbcL) gene, partial cds; chloroplast

AAACTTAACTGCTTCTCTTATCGGAAACGTTTTTGGTTTTAAAGCAGTAAAAGCATTACGTCTAGAAGATATGCGTCTTCCTTATGCTTACCTTAAAACTTTCCTTGGCCCAGCAACAGGAGTTATTGTAGAACGTGAAAGAATGGATGTTTTTGGACGTCCTTTATTAGGTGCTACTGTAAAACCAAAACTAGGTCTTTCAGGAAAAAACTATGGTCGTGTAGTTTATGAAGGATTAAAAGGTGGTTTAGATTTTTTAAAAGATGACGAAAATATCAATTCTCAACCATTCATGAGATGGCGTGAAAG

>gi|125617002|gb|EF165196.1| Synura curtispina strain CCMP847 ribulose-1,5-bisphosphate carboxylase/oxygenase large subunit (rbcL) gene, partial cds; chloroplast

AAACTTAACAGCTTCTATTATTGGTAACGTTTTTGGATTTAAAGCTGTAAAATGTTTACGTTTAGAAGATATGCGTATTCCTTATGCTTATTTAAAAACTTTCTTAGGCCCAGCAACAGGAGTTATCGTAGAACGTGAAAGAATGGATGTTTTTGGACGTCCTTTATTAGGTGCTACTGTTAAACCTAAATTAGGTCTTTCAGGAAAAAATTATGGTCGTGTAGTTTATGAAGGATTAAAAGGTGGTTTAGACTTCTTAAAAGACGATGAAAATATTAATTCACAACCATTCATGAGATGGCGTGAAAG

>gi|125617004|gb|EF165197.1| Synura sphagnicola strain CCMP1705 ribulose-1,5-bisphosphate carboxylase/oxygenase large subunit (rbcL) gene, partial cds; chloroplast

CAATTTAACAGCTTCGATTATTGGAAACGTTTTTGGTTTTAAAGCCGTAAAATGTTTACGTTTAGAAGATATGCGTATTCCATATTCTTATTTAAAAACTTTCTTAGGCCCTGCAACAGGAGTAATTGTAGAACGTGAAAGAATGGATGTTTTTGGTCGTCCTCTTTTAGGTGCAACTGTAAAACCTAAATTAGGTCTTTCAGGTAAAAACTATGGTCGTGTTGTTTATGAAGGTCTAAAAGGTGGATTAGATTTCTTAAAAGATGATGAAAATATTAATTCTCAACCTTTTATGAGATGGCGTGAAAG

>gi|125617008|gb|EF165199.1| Tessellaria volvocina strain CCMP1781 ribulose-1,5-bisphosphate carboxylase/oxygenase large subunit (rbcL) gene, partial cds; chloroplast

TAACTTAACAGCTTCTATTATCGGAAACGTTTTTGGTTTTAAAGCAGTAAAATGTTTACGTTTAGAAGATATGCGTATTCCTTATGCATATTTAAAAACTTTCATTGGTCCTGCTACAGGTGTAGTTGTAGAACGTGAAAGATTAGACGTTTTTGGACGTCCTCTTTTAGGTGCAACAGTAAAACCAAAATTAGGTCTTTCTGGAAAAAATTATGGTCGTGTAGTTTATGAAGGATTAAAAGGTGGTTTAGACTTCTTAAAAGATGACGAAAATATTAACTCTCAACCATTCATGCGTTGGCGTGAACG

>gi|156534942|gb|EF589143.1| Chrysocapsa paludosa strain LCR-MP ribulose-1,5-bisphosphate carboxylase/oxygenase large subunit (rbcL) gene, partial cds; chloroplast

TAACTTAACAGCATCTATTATTGGGAACGTATTTGGTTTCAAAGCGGTAAAATGTCTTCGTTTAGAAGATATGCGTATTCCTTTTGCATATTTAAAAACATTCATTGGTCCAGCTGCTGGAGTTATTGTAGAACGTGAAAGACTTGACGTATTTGGACGTCCTTTATTAGGAGCAACTGTAAAACCAAAATTAGGTCTTTCAGGAAAAAACTATGGTCGTGTAGTTTATGAAGGATTAAAAGGTGGTTTAGACTTCTTAAAAGATGACGAAAACATTAACTCTCAACCATTCATGAGATGGCGTGAAAG

>gi|1658264175|gb|MK153250.1| Uroglena volvox isolate U26-3 ribulose-1,5-bisphosphate carboxylase/oxygenase large subunit (rbcL) gene, partial cds; chloroplast

TAACTTAACAGCATCAATTATTGGTAACGTTTTTGGTTTTAAAGCTGTTAAATGCCTTCGTTTAGAAGACATGCGTCTTCCTTATGCTTATTTAAAAACATTTATTGGCCCTGCAGCTGGAGTTATTGTAGAACGCGAAAGACTTGACATTTTTGGTCGACCTCTTTTAGGTGCAACAGTAAAACCAAAATTAGGTCTTTCTGGTAAAAACTATGGTCGTGTAGTTTATGAAGGATTAAAAGGTGGTTTAGACTTCTTAAAAGATGATGAGAATATTAATTCTCAAGCATTTATGAGATGGCGTGAAAG

>gi|1658264177|gb|MK153251.1| Uroglena sp. isolate U29-5 ribulose-1,5-bisphosphate carboxylase/oxygenase large subunit (rbcL) gene, partial cds; chloroplast

TAACTTAACAGCATCAATTATTGGTAACGTTTTTGGTTTTAAAGCTGTTAAATGCCTTCGTTTAGAAGACATGCGTCTTCCTTATGCTTATTTAAAAACATTTATTGGCCCTGCAGCTGGAGTTATTGTAGAGCGTGAAAGACTTGACGTTTTTGGTCGACCTCTTTTAGGTGCAACAGTAAAACCAAAATTAGGTCTTTCTGGTAAAAACTATGGTCGTGTAGTTTATGAAGGATTAAAAGGTGGGTTAGACTTCTTAAAAGATGATGAGAATATTAATTCTCAAGCATTTATGAGATGGCGTGAAAG

>gi|1658264179|gb|MK153252.1| Uroglena sp. isolate UK-37 ribulose-1,5-bisphosphate carboxylase/oxygenase large subunit (rbcL) gene, partial cds; chloroplast

TAACTTAACAGCATCAATTATTGGTAACGTTTTTGGTTTTAAAGCTGTTAAATGCCTTCGTTTAGAAGACATGCGTCTTCCTTATGCTTATTTAAAAACATTTATTGGCCCTGCAGCTGGAGTTATTGTAGAGCGTGAAAGACTTGACGTTTTTGGTCGACCTCTTTTAGGTGCAACAGTAAAACCAAAATTAGGTCTTTCTGGTAAAAACTATGGTCGTGTAGTTTATGAAGGATTAAAAGGTGGGTTAGACTTCTTAAAAGATGATGAGAATATTAATTCTCAAGCATTTATGAGATGGCGTGAAAG

>gi|1658264181|gb|MK153253.1| Uroglenopsis sp. isolate UJ-6 ribulose-1,5-bisphosphate carboxylase/oxygenase large subunit (rbcL) gene, partial cds; chloroplast

TAACTTAACAGCATCTATTATTGGTAACGTATTTGGATTTAAAGCCGTAAAAGCTCTTAGATTAGAAGATATGCGTATTCCATTTGCTTACTTAAAAACTTTCATTGGTCCTGCATCTGGTGTTATTGTTGAACGTGAAAGACTTGATGTTTTCGGACGTCCTCTTTTAGGTGCTACTGTAAAACCGAAATTAGGTCTTTCTGGAAAAAACTATGGTCGTGTAGTTTATGAAGGATTAAAAGGTGGTTTAGATTTCTTAAAAGATGATGAAAATATCAACTCACAACCATTCATGAGATGGCGTGAAAG

>gi|1658264183|gb|MK153254.1| Uroglenopsis sp. isolate UK-25 ribulose-1,5-bisphosphate carboxylase/oxygenase large subunit (rbcL) gene, partial cds; chloroplast

TAACTTAACAGCATCTATTATTGGTAACGTATTTGGATTTAAAGCCGTAAAAGCTCTTAGATTAGAAGATATGCGTCTTCCATTTGCTTACTTAAAAACTTTCATTGGTCCTGCATCTGGTGTTATTGTTGAACGTGAAAGACTTGATGTTTTCGGACGTCCTCTTTTAGGTGCTACTGTAAAACCGAAATTAGGTCTTTCTGGAAAAAACTATGGTCGTGTAGTTTATGAAGGATTAAAAGGTGGTTTAGATTTCTTAAAAGATGATGAAAATATCAACTCACAACCATTCATGAGATGGCGTGAAAG

>gi|1658264185|gb|MK153255.1| Uroglenopsis sp. isolate U19 ribulose-1,5-bisphosphate carboxylase/oxygenase large subunit (rbcL) gene, partial cds; chloroplast

TAACTTAACAGCATCTATTATTGGTAACGTATTTGGATTTAAAGCCGTAAAAGCTCTTAGATTAGAAGATATGCGTCTTCCATATGCTTACTTAAAAACTTTCATTGGTCCTGCATCTGGTGTTATTGTTGAACGTGAAAGACTTGATGTTTTCGGACGTCCTCTTTTAGGTGCTACTGTAAAACCGAAATTAGGTCTTTCTGGAAAAAACTATGGTCGTGTAGTTTATGAAGGATTAAAAGGTGGTTTAGATTTCTTAAAAGATGATGAAAACATCAACTCACAACCATTCATGAGATGGCGTGAAAG

>gi|1658264187|gb|MK153256.1| Uroglenopsis americana isolate UK-4 ribulose-1,5-bisphosphate carboxylase/oxygenase large subunit (rbcL) gene, partial cds; chloroplast

TAACTTAACAGCATCTATTATTGGTAACGTATTTGGATTTAAAGCCGTAAAAGCTCTTAGATTAGAAGATATGCGTCTTCCATTTGCTTACTTAAAAACTTTCATTGGTCCTGCAGCTGGTGTTATTGTTGAACGTGAAAGACTTGATGTTTTCGGACGTCCTCTTTTAGGTGCTACTGTAAAACCGAAATTAGGTCTTTCTGGAAAAAACTATGGTCGTGTAGTTTATGAAGGATTAAAAGGTGGTTTAGATTTCTTAAAAGATGATGAAAACATCAACTCACAACCATTCATGAGATGGCGTGAAAG

>gi|1658264189|gb|MK153257.1| Uroglenopsis turfosa isolate UN-28 ribulose-1,5-bisphosphate carboxylase/oxygenase large subunit (rbcL) gene, partial cds; chloroplast

TAACTTAACAGCATCTATTATTGGTAACGTATTTGGATTTAAAGCCGTAAAAGCTCTTAGATTAGAAGATATGCGTCTTCCATTTGCTTACTTAAAAACGTTTATTGGTCCTGCATCTGGTGTTATCGTTGAACGTGAAAGACTTGATGTTTTCGGACGTCCTCTTTTAGGTGCTACTGTAAAACCAAAATTAGGTCTTTCTGGAAAAAACTATGGTCGTGTAGTTTATGAAGGATTAAAAGGTGGTTTAGATTTCTTAAAAGATGATGAAAACATCAACTCACAACCATTCATGAGATGGCGTGAAAG

>gi|1658264191|gb|MK153258.1| Uroglenopsis turfosa isolate UK-81 ribulose-1,5-bisphosphate carboxylase/oxygenase large subunit (rbcL) gene, partial cds; chloroplast

TAACTTAACAGCATCTATTATTGGTAACGTATTTGGATTTAAAGCCGTAAAAGCTCTTAGATTAGAAGATATGCGTCTTCCATTTGCTTACTTAAAAACGTTTATTGGTCCTGCATCTGGTGTTATCGTTGAACGTGAAAGACTTGATGTTTTCGGACGTCCTCTTTTAGGTGCTACTGTAAAACCAAAATTAGGTCTTTCTGGAAAAAACTATGGTCGTGTAGTTTATGAAGGATTAAAAGGTGGTTTAGATTTCTTAAAAGATGATGAAAACATCAACTCACAACCATTCATGAGATGGCGTGAAAG

>gi|1658264193|gb|MK153259.1| Urostipulosphaera sp. isolate U7-1 ribulose-1,5-bisphosphate carboxylase/oxygenase large subunit (rbcL) gene, partial cds; chloroplast

AAACTTAACAGCATCTCTTATTGGGAATGTTTTTGGGTTTAAAGCAGTTAAATGTCTTCGTTTAGAAGATATGCGCCTTCCTTATGCTTACTTAAAAACTTTTATTGGTCCTGCAACTGGTGTTATCGTAGAACGTGAAAGATTAGATATCTTTGGTCGCCCACTTTTAGGAGCAACAGTAAAACCTAAATTAGGTCTTTCAGGAAAAAATTACGGTCGTGTAGTTTATGAAGGTCTTAGAGGTGGATTAGACTTTTTAAAAGATGATGAAAATATTAACTCTCAACCATTCATGAGATGGCGTGAAAG

>gi|1658264195|gb|MK153260.1| Urostipulosphaera sp. isolate U5-5 ribulose-1,5-bisphosphate carboxylase/oxygenase large subunit (rbcL) gene, partial cds; chloroplast

AAACTTAACTGCTTCTTTAATTGGTAACGTTTTTGGATTTAAAGCTGTTAAATGTCTTCGTTTAGAAGATATGCGCCTTCCTTATGCTTACTTAAAAACTTTCATTGGTCCTGCAACTGGTGTTATTGTAGAACGTGAAAGATTAGATGTATTTGGTCGTCCTCTTTTAGGAGCAACAGTAAAACCTAAATTAGGTCTTTCAGGAAAAAACTATGGTCGTGTAGTATATGAAGGTCTTAGAGGTGGTTTAGATTTCTTAAAAGATGACGAAAATATTAACTCTCAACCATTCATGAGATGGCGTGAAAG

>gi|1658264197|gb|MK153261.1| Urostipulosphaera notabilis isolate U12-1 ribulose-1,5-bisphosphate carboxylase/oxygenase large subunit (rbcL) gene, partial cds; chloroplast

AAACTTAACTGCTTCTTTAATTGGTAACGTTTTTGGATTTAAAGCTGTTAAATGTCTTCGTTTAGAAGATATGCGCCTTCCCTATGCTTATTTAAAAACTTTCATTGGTCCTGCAAGTGGTGTTATTGTAGAACGTGAAAGATTAGATGTATTTGGTCGTCCTCTTTTAGGAGCAACAGTAAAACCTAAATTAGGTCTTTCAGGAAAAAACTATGGTCGTGTAGTATATGAAGGTCTTAGAGGTGGTTTAGATTTCTTAAAAGATGATGAAAATATTAACTCTCAACCATTCATGAGATGGCGTGAAAG

>gi|1658264199|gb|MK153262.1| Urostipulosphaera sp. isolate U10-6 ribulose-1,5-bisphosphate carboxylase/oxygenase large subunit (rbcL) gene, partial cds; chloroplast

AAACTTAACTGCTTCTCTAATTGGTAACGTTTTTGGATTTAAAGCTGTTAAATGTCTTCGTTTAGAAGATATGCGCCTTCCTTATGCTTACTTAAAAACTTTCATTGGTCCTGCAAGTGGTGTTATTGTAGAACGTGAAAGATTAGATGTATTTGGTCGTCCTCTTTTAGGAGCAACAGTAAAACCTAAATTAGGTCTTTCAGGAAAAAACTATGGTCGTGTAGTATATGAAGGTCTTAGAGGTGGTTTAGATTTCTTAAAAGATGATGAAAATATTAACTCTCAACCATTCATGAGATGGCGTGAAAG

>gi|1658264201|gb|MK153263.1| Urostipulosphaera sp. isolate UP-34 ribulose-1,5-bisphosphate carboxylase/oxygenase large subunit (rbcL) gene, partial cds; chloroplast

AAACTTAACTGCTTCTTTAATTGGTAACGTTTTTGGATTTAAAGCTGTTAAATGTCTTCGTTTAGAAGATATGCGCCTTCCTTATGCTTACTTAAAAACTTTCATTGGTCCTGCAAGTGGTGTTATTGTAGAACGTGAAAGATTAGATGTATTTGGTCGTCCTCTTTTAGGAGCAACAGTAAAACCTAAATTAGGTCTTTCAGGGAAAAACTACGGTCGTGTAGTATATGAAGGTCTTAGAGGTGGTTTAGATTTCTTAAAAGATGATGAAAATATTAACTCTCAACCATTCATGAGATGGCGTGAAAG

>gi|1756564916|gb|MK949446.1| Kremastochrysopsis austriaca strain DR75b ribulose-1,5-bisphosphate carboxylase/oxygenase large subunit mRNA, complete cds; chloroplast

TAACTTAACAGCATCTATTATCGGAAACGTATTCGGTTTCAAAGCCGTTAAATGTCTTCGTTTAGAAGATATGCGTATTCCTTATGCTTATTTAAAAACTTTCATTGGTCCAGCTGCTGGAGTAATTGTAGAACGTGAAAGACTTGACGTATTTGGACGTCCTTTATTAGGAGCAACAGTTAAACCAAAATTAGGTCTTTCTGGTAAAAACTACGGTCGTGTAGTTTACGAAGGATTAAAAGGTGGTTTAGACTTCTTAAAAGATGACGAAAACATCAACTCTCAACCATTCATGAGATGGCGTGAAAG

>gi|1770647962|gb|MK614367.1| Kremastochrysopsis austriaca strain DR75b ribulose-1,5-bisphosphate carboxylase/oxygenase large subunit (rbcL) gene, partial cds; chloroplast

TAACTTAACAGCATCTATTATCGGAAACGTATTCGGTTTCAAAGCCGTTAAATGTCTTCGTTTAGAAGATATGCGTATTCCTTATGCTTATTTAAAAACTTTCATTGGTCCAGCTGCTGGAGTAATTGTAGAACGTGAAAGACTTGACGTATTTGGACGTCCTTTATTAGGAGCAACAGTTAAACCAAAATTAGGTCTTTCTGGTAAAAACTACGGTCGTGTAGTTTACGAAGGATTAAAAGGTGGTTTAGACTTCTTAAAAGATGACGAAAACATCAACTCTCAACCATTCATGAGATGGCGTGAAAG

>gi|2198617969|gb|MW251560.1| Uroglena glabra strain UG-40 ribulose-1,5-bisphosphate carboxylase/oxygenase large subunit (rbcL) gene, partial cds; chloroplast

TAACTTAACAGCATCAATTATTGGTAACGTTTTTGGTTTTAAAGCTGTTAAATGCCTTCGTTTAGAAGATATGCGTCTTCCTTATGCTTATTTAAAAACATTTATTGGCCCTGCATCTGGAGTTATTGTAGAACGTGAAAGACTTGACGTTTTTGGTCGACCTCTTTTAGGTGCAACAGTAAAACCAAAATTAGGTCTTTCTGGTAAAAACTATGGTCGTGTAGTTTATGAAGGATTAAAAGGTGGTTTAGACTTCTTAAAAGATGATGAGAATATTAATTCTCAAGCATTTATGAGATGGCGTGAAAG

>gi|2198617971|gb|MW251561.1| Uroglena glabra strain U36-7 ribulose-1,5-bisphosphate carboxylase/oxygenase large subunit (rbcL) gene, partial cds; chloroplast

TAACTTAACAGCATCAATTATTGGTAACGTTTTTGGTTTTAAAGCTGTTAAATGCCTTCGTTTAGAAGATATGCGTCTTCCTTATGCTTATTTAAAAACATTTATTGGCCCTGCATCTGGAGTTATTGTAGAACGTGAAAGACTTGACGTTTTTGGTCGACCTCTTTTAGGTGCAACAGTAAAACCAAAATTAGGTCTTTCTGGTAAAAACTATGGTCGTGTAGTTTATGAAGGATTAAAAGGTGGTTTAGACTTCTTAAAAGATGATGAGAATATTAATTCTCAAGCATTTATGAGATGGCGTGAAAG

>gi|2198617973|gb|MW251562.1| Uroglena glabra strain U24-3 ribulose-1,5-bisphosphate carboxylase/oxygenase large subunit (rbcL) gene, partial cds; chloroplast

TAACTTAACAGCATCAATTATTGGTAACGTTTTTGGTTTTAAAGCTGTTAAATGCCTTCGTTTAGAAGATATGCGTCTTCCTTATGCTTATTTAAAAACATTTATTGGCCCTGCATCTGGAGTTATTGTAGAACGTGAAAGACTTGACGTTTTTGGTCGACCTCTTTTAGGTGCAACAGTAAAACCAAAATTAGGTCTTTCTGGTAAAAACTATGGTCGTGTAGTTTATGAAGGATTAAAAGGTGGTTTAGACTTCTTAAAAGATGATGAGAATATTAATTCTCAAGCATTTATGAGATGGCGTGAAAG

>gi|2198617975|gb|MW251563.1| Uroglena imitata strain U32-1 ribulose-1,5-bisphosphate carboxylase/oxygenase large subunit (rbcL) gene, partial cds; chloroplast

TAACTTAACAGCATCAATTATTGGTAACGTTTTTGGTTTTAAAGCTGTTAAATGCCTTCGTTTAGAAGACATGCGTCTTCCTTATGCTTATTTAAAAACATTTATTGGCCCTGCATCTGGAGTTATTGTAGAGCGTGAAAGACTTGACGTTTTTGGTCGACCTCTTTTAGGTGCAACAGTAAAACCAAAATTAGGTCTTTCTGGTAAAAACTATGGTCGTGTAGTTTATGAAGGATTAAAAGGTGGGTTAGACTTCTTAAAAGATGATGAGAATATTAATTCTCAAGCATTTATGAGATGGCGTGAAAG

>gi|2198617977|gb|MW251564.1| Uroglena skujae strain U26-14-379 ribulose-1,5-bisphosphate carboxylase/oxygenase large subunit (rbcL) gene, partial cds; chloroplast

TAACTTAACAGCATCAATTATTGGTAACGTTTTTGGTTTTAAAGCTGTTAAATGCCTTCGTTTAGAAGACMTGCGTCTTCCTTATGCTTATTTAAAAACATTTATTGGCCCTGCATCYGGAGTTATTGTAGAGCGTGAAAGACTTGACGTTTTTGGTCGACCTCTTTTAGGTGCAACAGTAAAACCAAAATTAGGTCTTTCTGGTAAAAACTATGGTCGTGTAGTTTATGAAGGATTAAAAGGTGGGTTAGACTTCTTAAAAGATGATGAGAATATTAATTCTCAAGCATTTATGAGATGGCGTGAAAG

>gi|2198617979|gb|MW251565.1| Uroglena skujae strain U26-33-670 ribulose-1,5-bisphosphate carboxylase/oxygenase large subunit (rbcL) gene, partial cds; chloroplast

TAACTTAACAGCATCAATTATTGGTAACGTTTTTGGTTTTAAAGCTGTTAAATGCCTTCGTTTAGAAGACATGCGTCTTCCTTATGCTTATTTAAAAACATTTATTGGCCCTGCATCTGGAGTTATTGTAGAGCGTGAAAGACTTGACGTTTTTGGTCGACCTCTTTTAGGTGCAACAGTAAAACCAAAATTAGGTCTTTCTGGTAAAAACTATGGTCGTGTAGTTTATGAAGGATTAAAAGGTGGGTTAGACTTCTTAAAAGATGATGAGAATATTAATTCTCAAGCATTTATGAGATGGCGTGAAAG

>gi|2198617981|gb|MW251566.1| Uroglena zachariasii strain U27-2 ribulose-1,5-bisphosphate carboxylase/oxygenase large subunit (rbcL) gene, partial cds; chloroplast

TAACTTAACAGCATCAATTATTGGTAACGTTTTTGGTTTTAAAGCTGTTAAATGCCTTCGTTTAGAAGACATGCGTCTTCCTTATGCTTATTTAAAAACATTTATTGGCCCTGCAGCTGGAGTTATTGTAGAGCGTGAAAGACTTGACGTTTTTGGTCGACCTCTTTTAGGTGCAACAGTAAAACCAAAATTAGGTCTTTCTGGTAAAAACTATGGTCGTGTAGTTTATGAAGGATTAAAAGGTGGGTTAGACTTCTTAAAAGATGATGAGAATATTAATTCTCAAGCATTTATGAGATGGCGTGAAAG

>gi|2198617983|gb|MW251567.1| Uroglena zachariasii strain U29-1-499 ribulose-1,5-bisphosphate carboxylase/oxygenase large subunit (rbcL) gene, partial cds; chloroplast

TAACTTAACAGCATCAATTATTGGTAACGTTTTTGGTTTTAAAGCTGTTAAATGCCTTCGTTTAGAAGACATGCGTCTTCCTTATGCTTATTTAAAAACATTTATTGGCCCTGCAGCTGGAGTTATTGTAGAGCGTGAAAGACTTGACGTTTTTGGTCGACCTCTTTTAGGTGCAACAGTAAAACCAAAATTAGGTCTTTCTGGTAAAAACTATGGTCGTGTAGTTTATGAAGGATTAAAAGGTGGGTTAGACTTCTTAAAAGATGATGAGAATATTAATTCTCAAGCATTTATGAGATGGCGTGAAAG

>gi|2198617985|gb|MW251568.1| Uroglena zachariasii strain U38-Ua ribulose-1,5-bisphosphate carboxylase/oxygenase large subunit (rbcL) gene, partial cds; chloroplast

TAACTTAACAGCATCAATTATTGGTAACGTTTTTGGTTTTAAAGCTGTTAAATGCCTTCGTTTAGAAGACATGCGTCTTCCTTATGCTTATTTAAAAACATTTATTGGCCCTGCAGCTGGAGTTATTGTAGAGCGTGAAAGACTTGACGTTTTTGGTCGACCTCTTTTAGGTGCAACAGTAAAACCAAAATTAGGTCTTTCTGGTAAAAACTATGGTCGTGTAGTTTATGAAGGATTAAAAGGTGGGTTAGACTTCTTAAAAGATGATGAGAATATTAATTCTCAAGCATTTATGAGATGGCGTGAAAG

>gi|2198617987|gb|MW251569.1| Uroglena zachariasii strain U26-19-449 ribulose-1,5-bisphosphate carboxylase/oxygenase large subunit (rbcL) gene, partial cds; chloroplast

TAACTTAACAGCATCAATTATTGGTAACGTTTTTGGTTTTAAAGCTGTTAAATGCCTTCGTTTAGAAGACATGCGTCTTCCTTATGCTTATTTAAAAACATTTATTGGCCCTGCAGCTGGAGTTATTGTAGAGCGTGAAAGACTTGACGTTTTTGGTCGACCTCTTTTAGGTGCAACAGTAAAACCAAAATTAGGTCTTTCTGGTAAAAACTATGGTCGTGTAGTTTATGAAGGATTAAAAGGTGGGTTAGACTTCTTAAAAGATGATGAGAATATTAATTCTCAAGCATTTATGAGATGGCGTGAAAG

>gi|2198617989|gb|MW251570.1| Uroglena zachariasii strain UG-23-613 ribulose-1,5-bisphosphate carboxylase/oxygenase large subunit (rbcL) gene, partial cds; chloroplast

TAACTTAACAGCATCAATTATTGGTAACGTTTTTGGTTTTAAAGCTGTTAAATGCCTTCGTTTAGAAGACATGCGTCTTCCTTATGCTTATTTAAAAACATTTATTGGCCCTGCAGCTGGAGTTATTGTAGAGCGTGAAAGACTTGACGTTTTTGGTCGACCTCTTTTAGGTGCAACAGTAAAACCAAAATTAGGTCTTTCTGGTAAAAACTATGGTCGTGTAGTTTATGAAGGATTAAAAGGTGGGTTAGACTTCTTAAAAGATGATGAGAATATTAATTCTCAAGCATTTATGAGATGGCGTGAAAG

>gi|2198617991|gb|MW251571.1| Uroglena cf. zachariasii strain U13-8 ribulose-1,5-bisphosphate carboxylase/oxygenase large subunit (rbcL) gene, partial cds; chloroplast

TAACTTAACAGCATCAATTATTGGTAACGTTTTTGGTTTTAAAGCTGTTAAATGCCTTCGTTTAGAAGACATGCGTCTTCCTTATGCTTATTTAAAAACATTTATTGGCCCTGCATCTGGAGTTATTGTAGAGCGTGAAAGACTTGACGTTTTTGGTCGACCTCTTTTAGGTGCAACAGTAAAACCAAAATTAGGTCTTTCTGGTAAAAACTATGGTCGTGTAGTTTATGAAGGATTAAAAGGTGGGTTAGACTTCTTAAAAGATGATGAGAATATTAATTCTCAAGCATTTATGAGATGGCGTGAAAG

>gi|2198617993|gb|MW251572.1| Uroglena cf. zachariasii strain UP-21 ribulose-1,5-bisphosphate carboxylase/oxygenase large subunit (rbcL) gene, partial cds; chloroplast

TAACTTAACAGCATCAATTATTGGTAACGTTTTTGGTTTTAAAGCTGTTAAATGCCTTCGTTTAGAAGACATGCGTCTTCCTTATGCTTATTTAAAAACATTTATTGGCCCTGCATCTGGAGTTATTGTAGAGCGTGAAAGACTTGACGTTTTTGGTCGACCTCTTTTAGGTGCAACAGTAAAACCAAAATTAGGTCTTTCTGGTAAAAACTATGGTCGTGTAGTTTATGAAGGATTAAAAGGTGGGTTAGACTTCTTAAAAGATGATGAGAATATTAATTCTCAAGCATTTATGAGATGGCGTGAAAG

>gi|2198617995|gb|MW251573.1| Uroglena cf. zachariasii strain U16-15 ribulose-1,5-bisphosphate carboxylase/oxygenase large subunit (rbcL) gene, partial cds; chloroplast

TAACTTAACAGCATCAATTATTGGTAACGTTTTTGGTTTTAAAGCTGTTAAATGCCTTCGTTTAGAAGACATGCGTCTTCCTTATGCTTATTTAAAAACATTTATTGGCCCTGCATCTGGAGTTATTGTAGAGCGTGAAAGACTTGACGTTTTTGGTCGACCTCTTTTAGGTGCAACAGTAAAACCAAAATTAGGTCTTTCTGGTAAAAACTATGGTCGTGTAGTTTATGAAGGATTAAAAGGTGGGTTAGACTTCTTAAAAGATGATGAGAATATTAATTCTCAAGCATTTATGAGATGGCGTGAAAG

>gi|2198617997|gb|MW251574.1| Uroglena cf. zachariasii strain U1-2 ribulose-1,5-bisphosphate carboxylase/oxygenase large subunit (rbcL) gene, partial cds; chloroplast

TAACTTAACAGCATCAATTATTGGTAACGTTTTTGGTTTTAAAGCTGTTAAATGCCTTCGTTTAGAAGACATGCGTCTTCCTTATGCTTATTTAAAAACATTTATTGGCCCTGCATCTGGAGTTATTGTAGAGCGTGAAAGACTTGACGTTTTTGGTCGACCTCTTTTAGGTGCAACAGTAAAACCAAAATTAGGTCTTTCTGGTAAAAACTATGGTCGTGTAGTTTATGAAGGATTAAAAGGTGGGTTAGACTTCTTAAAAGATGATGAGAATATTAATTCTCAAGCATTTATGAGATGGCGTGAAAG

>gi|2198617999|gb|MW251575.1| Uroglena sp. 2 strain U34-1 ribulose-1,5-bisphosphate carboxylase/oxygenase large subunit (rbcL) gene, partial cds; chloroplast

TAACTTAACAGCATCAATTATTGGTAACGTTTTTGGTTTTAAAGCTGTTAAATGCCTTCGTTTAGAAGACATGCGTCTTCCTTATGCTTATTTAAAAACATTTATTGGCCCTGCAGCTGGAGTTATTGTAGAGCGTGAAAGACTTGACGTTTTTGGTCGACCTCTTTTAGGTGCAACAGTAAAACCAAAATTAGGTCTTTCTGGTAAAAACTATGGTCGTGTAGTTTATGAAGGATTAAAAGGTGGTTTAGACTTCTTAAAAGATGATGAGAATATTAATTCTCAAGCATTTATGAGATGGCGTGAAAG

>gi|2198618001|gb|MW251576.1| Uroglena botrys strain U2-6 ribulose-1,5-bisphosphate carboxylase/oxygenase large subunit (rbcL) gene, partial cds; chloroplast

TAACTTAACAGCATCTATTATTGGTAACGTATTTGGATTTAAAGCCGTAAAAGCTCTTAGATTAGAAGATATGCGTCTTCCATATGCTTACTTAAAAACTTTCATTGGTCCTGCATCTGGTGTTATTGTTGAACGTGAAAGACTTGATGTTTTCGGACGTCCTCTTTTAGGTGCTACTGTAAAACCGAAATTAGGTCTTTCTGGAAAAAACTATGGTCGTGTAGTTTATGAAGGATTAAAAGGTGGTTTAGATTTCTTAAAAGATGATGAAAACATCAACTCACAACCATTCATGAGATGGCGTGAAAG

>gi|2198618003|gb|MW251577.1| Uroglena botrys strain U1-6-62 ribulose-1,5-bisphosphate carboxylase/oxygenase large subunit (rbcL) gene, partial cds; chloroplast

TAACTTAACAGCATCTATTATTGGTAACGTATTTGGATTTAAAGCCGTAAAAGCTCTTAGATTAGAAGATATGCGTCTTCCATATGCTTACTTAAAAACTTTCATTGGTCCTGCATCTGGTGTTATTGTTGAACGTGAAAGACTTGATGTTTTCGGACGTCCTCTTTTAGGTGCTACTGTAAAACCGAAATTAGGTCTTTCTGGAAAAAACTATGGTCGTGTAGTTTATGAAGGATTAAAAGGTGGTTTAGATTTCTTAAAAGATGATGAAAACATCAACTCACAACCATTCATGAGATGGCGTGAAAG

>gi|2198618005|gb|MW251578.1| Uroglenopsis turfosa strain UN-9 ribulose-1,5-bisphosphate carboxylase/oxygenase large subunit (rbcL) gene, partial cds; chloroplast

TAACTTAACAGCATCTATTATTGGTAACGTATTTGGATTTAAAGCCGTAAAAGCTCTTAGATTAGAAGATATGCGTCTTCCATTTGCTTACTTAAAAACGTTTATTGGTCCTGCATCTGGTGTTATCGTTGAACGTGAAAGACTTGATGTTTTCGGACGTCCTCTTTTAGGTGCTACTGTAAAACCAAAATTAGGTCTTTCTGGAAAAAACTATGGTCGTGTAGTTTATGAAGGATTAAAAGGTGGTTTAGATTTCTTAAAAGATGATGAAAACATCAACTCACAACCATTCATGAGATGGCGTGAAAG

>gi|2198618007|gb|MW251579.1| Uroglenopsis turfosa strain UK-79 ribulose-1,5-bisphosphate carboxylase/oxygenase large subunit (rbcL) gene, partial cds; chloroplast

TAACTTAACAGCATCTATTATTGGTAACGTATTTGGATTTAAAGCTGTAAAAGCTCTTAGATTAGAAGATATGCGTCTTCCATTTGCTTACTTAAAAACGTTTATTGGTCCTGCATCTGGTGTTATCGTTGAACGTGAAAGACTTGATGTTTTCGGACGTCCTCTTTTAGGTGCTACTGTAAAACCAAAATTAGGTCTTTCTGGAAAAAACTATGGTCGTGTAGTTTATGAAGGATTAAAAGGTGGTTTAGATTTCTTAAAAGATGATGAAAACATCAACTCACAACCATTCATGAGATGGCGTGAAAG

>gi|2198618009|gb|MW251580.1| Uroglenopsis sp. 1 strain U1-6-120 ribulose-1,5-bisphosphate carboxylase/oxygenase large subunit (rbcL) gene, partial cds; chloroplast

TAACTTAACAGCATCTATTATTGGTAACGTATTTGGATTTAAAGCCGTAAAAGCTCTTAGATTAGAAGATATGCGTATTCCATTTGCTTACTTAAAAACTTTCATTGGTCCTGCATCTGGTGTTATTGTTGAACGTGAAAGACTTGATGTTTTCGGACGTCCTCTTTTAGGTGCTACTGTAAAACCGAAATTAGGTCTTTCTGGAAAAAACTATGGTCGTGTAGTTTATGAAGGATTAAAAGGTGGTTTAGATTTCTTAAAAGATGATGAAAATATCAACTCACAACCATTCATGAGATGGCGTGAAAG

>gi|2198618011|gb|MW251581.1| Uroglenopsis sp. 1 strain UJ-2 ribulose-1,5-bisphosphate carboxylase/oxygenase large subunit (rbcL) gene, partial cds; chloroplast

TAACTTAACAGCATCTATTATTGGTAACGTATTTGGATTTAAAGCCGTAAAAGCTCTTAGATTAGAAGATATGCGTATTCCATTTGCTTACTTAAAAACTTTCATTGGTCCTGCATCTGGTGTTATTGTTGAACGTGAAAGACTTGATGTTTTCGGACGTCCTCTTTTAGGTGCTACTGTAAAACCGAAATTAGGTCTTTCTGGAAAAAACTATGGTCGTGTAGTTTATGAAGGATTAAAAGGTGGTTTAGATTTCTTAAAAGATGATGAAAATATCAACTCACAACCATTCATGAGATGGCGTGAAAG

>gi|2198618013|gb|MW251582.1| Uroglenopsis sp. 1 strain U27-5 ribulose-1,5-bisphosphate carboxylase/oxygenase large subunit (rbcL) gene, partial cds; chloroplast

TAACTTAACAGCATCTATTATTGGTAACGTATTTGGATTTAAAGCCGTAAAAGCTCTTAGATTAGAAGATATGCGTATTCCATTTGCTTACTTAAAAACTTTCATTGGTCCTGCATCTGGTGTTATTGTTGAACGTGAAAGACTTGATGTTTTCGGACGTCCTCTTTTAGGTGCTACTGTAAAACCGAAATTAGGTCTTTCTGGAAAAAACTATGGTCGTGTAGTTTATGAAGGATTAAAAGGTGGTTTAGATTTCTTAAAAGATGATGAAAATATCAACTCACAACCATTCATGAGATGGCGTGAAAG

>gi|2198618015|gb|MW251583.1| Uroglenopsis sp. 2 strain UK-17-869 ribulose-1,5-bisphosphate carboxylase/oxygenase large subunit (rbcL) gene, partial cds; chloroplast

TAACTTAACAGCATCTATTATTGGTAACGTATTTGGATTTAAAGCCGTAAAAGCTCTTAGATTAGAAGATATGCGTCTTCCATTTGCTTACTTAAAAACTTTCATTGGTCCTGCATCTGGTGTTATTGTTGAACGTGAAAGACTTGATGTTTTCGGACGTCCTCTTTTAGGTGCTACTGTAAAACCGAAATTAGGTCTTTCTGGAAAAAACTATGGTCGTGTAGTTTATGAAGGATTAAAAGGTGGTTTAGATTTCTTAAAAGATGATGAAAATATCAACTCACAACCATTCATGAGATGGCGTGAAAG

>gi|2198618017|gb|MW251584.1| Uroglenopsis sp. 2 strain U26-19-451 ribulose-1,5-bisphosphate carboxylase/oxygenase large subunit (rbcL) gene, partial cds; chloroplast

TAACTTAACAGCATCTATTATTGGTAACGTATTTGGATTTAAAGCCGTAAAAGCTCTTAGATTAGAAGATATGCGTCTTCCATATGCTTACTTAAAAACTTTCATTGGTCCTGCAGCTGGTGTTATTGTTGAACGTGAAAGACTTGATGTTTTCGGACGTCCTCTTTTAGGTGCTACTGTAAAACCGAAATTAGGTCTTTCTGGAAAAAACTATGGTCGTGTAGTTTATGAAGGATTAAAAGGTGGTTTAGATTTCTTAAAAGATGATGAAAACATCAACTCACAACCATTCATGAGATGGCGTGAAAG

>gi|2198618019|gb|MW251585.1| Urostipulosphaera granulata strain U33 ribulose-1,5-bisphosphate carboxylase/oxygenase large subunit (rbcL) gene, partial cds; chloroplast

AAACTTAACAGCATCTCTTATTGGGAATGTTTTTGGGTTTAAAGCAGTTAAATGTCTTCGTTTAGAAGATATGCGCCTTCCTTATGCTTACTTAAAAACTTTTATTGGTCCTGCAACTGGTGTTATCGTAGAACGTGAAAGATTAGATATCTTTGGTCGCCCACTTTTAGGAGCAACAGTAAAACCTAAATTAGGTCTTTCAGGAAAAAATTACGGTCGTGTAGTTTATGAAGGTCTTAGAGGTGGATTAGACTTTTTAAAAGATGATGAAAATATTAACTCTCAACCATTCATGAGATGGCGTGAAAG

>gi|2198618021|gb|MW251586.1| Urostipulosphaera lindiae strain UI-18 ribulose-1,5-bisphosphate carboxylase/oxygenase large subunit (rbcL) gene, partial cds; chloroplast

AAACTTAACTGCTTCTTTAATTGGTAACGTTTTTGGATTTAAAGCTGTTAAATGTCTTCGTTTAGAAGATATGCGCCTTCCTTATGCTTACTTAAAAACTTTCATTGGTCCTGCAAGTGGTGTTATTGTAGAACGTGAAAGATTAGATGTATTTGGTCGTCCTCTTTTAGGAGCAACAGTAAAACCTAAATTAGGTCTTTCAGGGAAAAACTACGGTCGTGTAGTATATGAAGGTCTTAGAGGTGGTTTAGATTTCTTAAAAGATGATGAAAATATTAACTCTCAACCATTCATGAGATGGCGTGAAAG

>gi|2198618023|gb|MW251587.1| Urostipulosphaera lindiae strain U29-1-496 ribulose-1,5-bisphosphate carboxylase/oxygenase large subunit (rbcL) gene, partial cds; chloroplast

AAACTTAACTGCTTCTTTAATTGGTAACGTTTTTGGATTTAAAGCTGTTAAATGTCTTCGTTTAGAAGATATGCGCCTTCCTTATGCTTACTTAAAAACTTTCATTGGTCCTGCAAGTGGTGTTATTGTAGAACGTGAAAGATTAGATGTATTTGGTCGTCCTCTTTTAGGAGCAACAGTAAAACCTAAATTAGGTCTTTCAGGGAAAAACTACGGTCGTGTAGTATATGAAGGTCTTAGAGGTGGTTTAGATTTCTTAAAAGATGATGAAAATATTAACTCTCAACCATTCATGAGATGGCGTGAAAG

>gi|2198618025|gb|MW251588.1| Urostipulosphaera lindiae strain UN-17 ribulose-1,5-bisphosphate carboxylase/oxygenase large subunit (rbcL) gene, partial cds; chloroplast

AAACTTAACTGCTTCTTTAATTGGTAACGTTTTTGGATTTAAAGCTGTTAAATGTCTTCGTTTAGAAGATATGCGCCTTCCTTATGCTTACTTAAAAACTTTCATTGGTCCTGCAAGTGGTGTTATTGTAGAACGTGAAAGATTAGATGTATTTGGTCGTCCTCTTTTAGGAGCAACAGTAAAACCTAAATTAGGTCTTTCAGGGAAAAACTACGGTCGTGTAGTATATGAAGGTCTTAGAGGTGGTTTAGATTTCTTAAAAGATGATGAAAATATTAACTCTCAACCATTCATGAGATGGCGTGAAAG

>gi|2198618027|gb|MW251589.1| Urostipulosphaera notabilis strain U13-7 ribulose-1,5-bisphosphate carboxylase/oxygenase large subunit (rbcL) gene, partial cds; chloroplast

AAACTTAACTGCTTCTTTAATTGGTAACGTTTTTGGATTTAAAGCTGTTAAATGTCTTCGTTTAGAAGATATGCGCCTTCCCTATGCTTATTTAAAAACTTTCATTGGTCCTGCAAGTGGTGTTATTGTAGAACGTGAAAGATTAGATGTATTTGGTCGTCCTCTTTTAGGAGCAACAGTAAAACCTAAATTAGGTCTTTCAGGAAAAAACTATGGTCGTGTAGTATATGAAGGTCTTAGAGGTGGTTTAGATTTCTTAAAAGATGATGAAAATATTAACTCTCAACCATTCATGAGATGGCGTGAAAG

>gi|219940313|emb|AM421005.1| Phaeobotrys solitaria chloroplast partial rbcL gene for ribulose bisphosphate carboxylase large chain, strain SAG 15.95

TAATTTAACAGCATCTATCATTGGTAACGTTTTTGGTTTCAAAGCCGTTAAAGCATTACGTTTAGAAGATATGCGCATTCCTTTTGCATATTTAAAAACTTTCCAAGGTCCAGCTACAGGTTTAATTGTAGAAAGAGAAAGAATGGATAAATTTGGAAGACCATTCTTAGGTGCTACTGTAAAACCTAAACTAGGCCTTTCAGGTAAAAACTACGGTCGAGTAGTGTATGAAGGTTTACGTGGTGGTCTTGATTTCCTTAAAGATGATGAAAATATTAATTCACAACCATTCATGCGTTGGAGAGAACG

>gi|2477559156|gb|OQ466159.1| Dinobryon divergens strain CCMP3056 ribulose-1,5-bisphosphate carboxylase/oxygenase large subunit (rbcL) gene, partial cds; chloroplast

TAACTTAACAGCATCAATTATTGGAAACGTATTTGGATTCAAAGCGGTTAAAGCGCTTCGTTTAGAAGATATGCGTATTCCATATGCTTATTTAAAAACTTTCCTAGGACCAGCATCTGGGGTTATTGTAGAACGTGAAAGACTTGATGTTTTCGGACGTCCACTTTTAGGTGCAACAGTTAAACCAAAATTAGGTTTATCAGGTAAAAACTATGGTCGTGTAGTTTATGAAGGGTTAAAAGGTGGTTTAGATTTCTTAAAAGATGACGAAAATATTAACTCTCAACCATTCATGAGATGGCGTGAAAG

>gi|2477559158|gb|OQ466160.1| Dinobryon divergens strain CCMP3055 ribulose-1,5-bisphosphate carboxylase/oxygenase large subunit (rbcL) gene, partial cds; chloroplast

TAACTTAACAGCATCAATTATTGGAAACGTATTCGGATTTAAAGCGGTTAAAGCTCTTCGTCTAGAAGATATGCGTATTCCATACGCTTATTTAAAAACTTTCCTAGGACCAGCATCAGGGGTTATTGTAGAACGTGAAAGACTTGATGTTTTCGGACGTCCACTTTTAGGTGCAACAGTTAAACCAAAATTAGGTCTATCTGGTAAAAACTATGGTCGTGTAGTTTATGAAGGGTTAAAAGGTGGTTTAGATTTCTTAAAAGATGACGAAAATATTAACTCTCAACCATTCATGAGATGGCGTGAAAG

>gi|2477559160|gb|OQ466161.1| Dinobryon divergens strain CCMP2900 ribulose-1,5-bisphosphate carboxylase/oxygenase large subunit (rbcL) gene, partial cds; chloroplast

TAACTTAACAGCATCAATTATTGGAAACGTATTCGGATTTAAAGCGGTTAAAGCTCTTCGTCTAGAAGATATGCGTATTCCATACGCTTATTTAAAAACTTTCCTAGGACCAGCATCAGGGGTTATTGTAGAACGTGAAAGACTTGATGTTTTCGGACGTCCACTTTTAGGTGCAACAGTTAAACCAAAATTAGGTCTATCTGGTAAAAACTATGGTCGTGTAGTTTATGAAGGGTTAAAAGGTGGTTTAGATTTCTTAAAAGATGACGAAAATATTAACTCTCAACCATTCATGAGATGGCGTGAAAG

>gi|2477559162|gb|OQ466162.1| Dinobryon bavaricum strain CCMP3054 ribulose-1,5-bisphosphate carboxylase/oxygenase large subunit (rbcL) gene, partial cds; chloroplast

TAACTTAACAGCATCAATTATTGGAAACGTATTCGGATTTAAAGCGGTTAAAGCTCTTCGTCTAGAAGATATGCGTATTCCATATGCTTATTTAAAAACTTTCTTAGGACCTGCATCAGGGGTTATTGTAGAACGTGAAAGACTTGATGTTTTCGGACGTCCACTTTTAGGTGCAACAGTTAAACCAAAATTAGGTCTATCTGGTAAAAACTATGGTCGTGTAGTTTATGAAGGGTTAAAAGGTGGTTTAGATTTCTTAAAAGATGACGAAAATATTAACTCTCAACCATTCATGAGATGGCGTGAAAG

>gi|2477559164|gb|OQ466163.1| Dinobryon bavaricum strain CCMP3270 ribulose-1,5-bisphosphate carboxylase/oxygenase large subunit (rbcL) gene, partial cds; chloroplast

TAACTTAACAGCATCAATTATTGGAAACGTATTCGGATTTAAAGCGGTTAAAGCTCTTCGTCTAGAAGATATGCGTATTCCATATGCTTATTTAAAAACTTTCTTAGGACCTGCATCAGGGGTTATTGTAGAACGTGAAAGACTTGATGTTTTCGGACGTCCACTTTTAGGTGCAACAGTTAAACCAAAATTAGGTCTATCTGGTAAAAACTATGGTCGTGTAGTTTATGAAGGGTTAAAAGGTGGTTTAGATTTCTTAAAAGATGACGAAAATATTAACTCTCAACCATTCATGAGATGGCGTGAAAG

>gi|2477559166|gb|OQ466164.1| Dinobryon bavaricum strain CCMP2884 ribulose-1,5-bisphosphate carboxylase/oxygenase large subunit (rbcL) gene, partial cds; chloroplast

TAACTTAACAGCATCAATTATTGGAAACGTATTCGGATTTAAAGCGGTTAAAGCTCTTCGTCTAGAAGATATGCGTATTCCATATGCTTATTTAAAAACTTTCCTAGGACCAGCATCAGGGGTTATTGTAGAACGTGAAAGACTTGATGTTTTCGGACGTCCACTTTTAGGTGCAACAGTTAAACCAAAATTAGGTCTATCTGGTAAAAACTATGGTCGTGTAGTTTATGAAGGGTTAAAAGGTGGTTTAGATTTCTTAAAAGATGACGAAAATATTAACTCTCAACCATTCATGAGATGGCGTGAAAG

>gi|2477559168|gb|OQ466165.1| Dinobryon sp. MJeong-2023d strain Deokghi051818A ribulose-1,5-bisphosphate carboxylase/oxygenase large subunit (rbcL) gene, partial cds; chloroplast

TAACTTAACAGCATCAATTATTGGAAACGTATTTGGATTTAAAGCCGTTAAAGCTCTTCGTCTAGAAGATATGCGTATTCCATATGCTTATTTAAAAACTTTCCTAGGACCAGCATCAGGGGTTATTGTAGAACGTGAAAGACTTGATGTTTTCGGACGTCCACTTTTAGGTGCAACAGTTAAACCAAAATTAGGTCTATCTGGTAAAAACTATGGTCGTGTAGTTTATGAAGGATTAAAAGGTGGTTTAGATTTCTTAAAAGATGACGAAAATATTAACTCTCAACCATTCATGAGATGGCGTGAAAG

>gi|2477559170|gb|OQ466166.1| Dinobryon sp. MJeong-2023d strain Meokgol112021MS9 ribulose-1,5-bisphosphate carboxylase/oxygenase large subunit (rbcL) gene, partial cds; chloroplast

TAACTTAACAGCATCAATTATTGGAAACGTATTTGGATTTAAAGCCGTTAAAGCTCTTCGTCTAGAAGATATGCGTATTCCATATGCTTATTTAAAAACTTTCCTAGGACCAGCATCAGGGGTTATTGTAGAACGTGAAAGACTTGATGTTTTCGGACGTCCACTTTTAGGTGCAACAGTTAAACCAAAATTAGGTCTATCTGGTAAAAACTATGGTCGTGTAGTTTATGAAGGATTAAAAGGTGGTTTAGATTTCTTAAAAGATGACGAAAATATTAACTCTCAACCATTCATGAGATGGCGTGAAAG

>gi|2477559172|gb|OQ466167.1| Dinobryon sp. MJeong-2023d strain Yongghi051818A ribulose-1,5-bisphosphate carboxylase/oxygenase large subunit (rbcL) gene, partial cds; chloroplast

TAACTTAACAGCATCAATTATTGGAAACGTATTTGGATTTAAAGCCGTTAAAGCTCTTCGTCTAGAAGATATGCGTATTCCATATGCTTATTTAAAAACTTTCCTAGGACCAGCATCAGGGGTTATTGTAGAACGTGAAAGACTTGATGTTTTCGGACGTCCACTTTTAGGTGCAACAGTTAAACCAAAATTAGGTCTATCTGGTAAAAACTATGGTCGTGTAGTTTATGAAGGATTAAAAGGTGGTTTAGATTTCTTAAAAGATGACGAAAATATTAACTCTCAACCATTCATGAGATGGCGTGAAAG

>gi|2477559174|gb|OQ466168.1| Dinobryon sp. MJeong-2023d strain Oun112021MS2 ribulose-1,5-bisphosphate carboxylase/oxygenase large subunit (rbcL) gene, partial cds; chloroplast

TAACTTAACAGCATCAATTATTGGAAACGTATTTGGATTTAAAGCCGTTAAAGCTCTTCGTCTAGAAGATATGCGTATTCCATATGCTTATTTAAAAACTTTCCTAGGACCAGCATCAGGGGTTATTGTAGAACGTGAAAGACTTGATGTTTTCGGACGTCCACTTTTAGGTGCAACAGTTAAACCAAAATTAGGTCTATCTGGTAAAAACTATGGTCGTGTAGTTTATGAAGGATTAAAAGGTGGTTTAGATTTCTTAAAAGATGACGAAAATATTAACTCTCAACCATTCATGAGATGGCGTGAAAG

>gi|2477559176|gb|OQ466169.1| Dinobryon sp. MJeong-2023d strain Ori120521MS2 ribulose-1,5-bisphosphate carboxylase/oxygenase large subunit (rbcL) gene, partial cds; chloroplast

TAACTTAACAGCATCAATTATTGGAAACGTATTTGGATTTAAAGCCGTTAAAGCTCTTCGTCTAGAAGATATGCGTATTCCATATGCTTATTTAAAAACTTTCCTAGGACCAGCATCAGGGGTTATTGTAGAACGTGAAAGACTTGATGTTTTCGGACGTCCACTTTTAGGTGCAACAGTTAAACCAAAATTAGGTCTATCTGGTAAAAACTATGGTCGTGTAGTTTATGAAGGATTAAAAGGTGGTTTAGATTTCTTAAAAGATGACGAAAATATTAACTCTCAACCATTCATGAGATGGCGTGAAAG

>gi|2477559178|gb|OQ466170.1| Dinobryon cylindricollarium strain Gwangok010822MS1 ribulose-1,5-bisphosphate carboxylase/oxygenase large subunit (rbcL) gene, partial cds; chloroplast

TAACTTAACAGCATCAATTATTGGAAACGTTTTTGGATTTAAAGCAGTTAAAGCTTTACGTTTAGAAGATATGCGTATTCCATATGCTTACTTAAAAACTTTCTTAGGACCAGCATCAGGTGTTATTGTAGAACGTGAAAGACTTGACGTTTTCGGACGCCCACTTTTAGGTGCAACAGTTAAACCAAAATTAGGTCTATCTGGAAAAAATTATGGTCGTGTAGTTTATGAAGGATTAAAAGGTGGTTTAGATTTCTTAAAAGATGATGAAAATATTAACTCTCAACCATTCATGAGATGGCGTGAAAG

>gi|2477559180|gb|OQ466171.1| Dinobryon cylindricollarium strain Jeongdong010822MS1 ribulose-1,5-bisphosphate carboxylase/oxygenase large subunit (rbcL) gene, partial cds; chloroplast

TAACTTAACAGCATCAATTATTGGAAACGTTTTTGGATTTAAAGCAGTTAAAGCTTTACGTTTAGAAGATATGCGTATTCCATATGCTTACTTAAAAACTTTCTTAGGACCAGCATCAGGTGTTATTGTAGAACGTGAAAGACTTGACGTTTTCGGACGCCCACTTTTAGGTGCAACAGTTAAACCAAAATTAGGTCTATCTGGAAAAAATTATGGTCGTGTAGTTTATGAAGGATTAAAAGGTGGTTTAGATTTCTTAAAAGATGATGAAAATATTAACTCTCAACCATTCATGAGATGGCGTGAAAG

>gi|2477559182|gb|OQ466172.1| Dinobryon cylindricollarium strain Myeoseul111618D ribulose-1,5-bisphosphate carboxylase/oxygenase large subunit (rbcL) gene, partial cds; chloroplast

TAACTTAACAGCATCAATTATTGGAAACGTTTTTGGATTTAAAGCAGTTAAAGCTCTACGTTTAGAAGATATGCGTATTCCATATGCTTACTTAAAAACTTTCTTAGGACCAGCATCAGGTGTTATTGTAGAACGTGAAAGACTTGACGTTTTCGGACGCCCACTTTTAGGTGCAACAGTTAAACCAAAATTAGGTCTATCTGGAAAAAATTATGGTCGTGTAGTTTATGAAGGATTAAAAGGTGGTTTAGATTTCTTAAAAGATGATGAAAATATTAACTCTCAACCATTCATGAGATGGCGTGAAAG

>gi|2477559184|gb|OQ466173.1| Dinobryon sociale strain Angol061922MS3 ribulose-1,5-bisphosphate carboxylase/oxygenase large subunit (rbcL) gene, partial cds; chloroplast

AAACTTAACAGCATCGATTATTGGAAACGTTTTTGGATTTAAAGCTGTTAAAGCTCTTCGATTAGAAGATATGCGTATTCCATATGCTTACTTAAAAACTTTCTTAGGACCAGCATCAGGTGTTATTGTAGAACGTGAAAGACTTGACGTTTTCGGACGCCCACTTTTAGGTGCAACAGTTAAACCAAAATTAGGTCTATCTGGTAAAAACTATGGTCGTGTAGTTTATGAAGGATTAAAAGGTGGTTTAGATTTCTTAAAAGATGACGAAAATATTAACTCTCAACCATTCATGAGATGGCGTGAAAG

>gi|2477559186|gb|OQ466174.1| Dinobryon exstoundulatum strain Dallae111421MS1 ribulose-1,5-bisphosphate carboxylase/oxygenase large subunit (rbcL) gene, partial cds; chloroplast

AAACTTAACAGCATCAATTATTGGAAACGTTTTTGGATTTAAAGCTGTTAAAGCTCTTCGTTTAGAAGATATGCGTATTCCATATGCTTATTTAAAAACTTTCTTAGGACCAGCATCAGGTGTTATTGTAGAACGTGAAAGACTTGACGTTTTCGGACGTCCACTTTTAGGTGCAACAGTTAAACCAAAATTAGGTCTATCTGGTAAAAACTATGGTCGTGTAGTTTATGAAGGATTAAAAGGTGGTTTAGATTTCTTAAAAGATGACGAAAATATTAACTCTCAACCATTCATGAGATGGCGTGAAAG

>gi|2477559188|gb|OQ466175.1| Dinobryon similis strain Sanseong2ji102318A ribulose-1,5-bisphosphate carboxylase/oxygenase large subunit (rbcL) gene, partial cds; chloroplast

AAACTTAACAGCATCAATTATTGGAAACGTTTTTGGATTTAAAGCTGTTAAAGCGCTTCGTTTAGAAGATATGCGTATTCCATATGCTTATTTAAAAACTTTCTTAGGCCCAGCATCAGGTGTTATTGTAGAACGTGAAAGACTTGACGTTTTCGGACGCCCACTTTTAGGTGCAACAGTTAAACCAAAATTAGGTCTATCTGGTAAAAACTATGGTCGTGTAGTTTATGAAGGATTAAAAGGTGGTTTAGATTTCTTAAAAGATGACGAAAATATTAACTCTCAACCATTCATGAGATGGCGTGAAAG

>gi|2477559190|gb|OQ466176.1| Dinobryon similis strain Goemok010822MS2 ribulose-1,5-bisphosphate carboxylase/oxygenase large subunit (rbcL) gene, partial cds; chloroplast

AAACTTAACAGCATCAATTATTGGAAACGTTTTTGGATTTAAAGCTGTTAAAGCGCTTCGTTTAGAAGATATGCGTATTCCATATGCTTATTTAAAAACTTTCTTAGGCCCAGCATCAGGTGTTATTGTAGAACGTGAAAGACTTGACGTTTTCGGACGCCCACTTTTAGGTGCAACAGTTAAACCAAAATTAGGTCTATCTGGTAAAAACTATGGTCGTGTAGTTTATGAAGGATTAAAAGGTGGTTTAGATTTCTTAAAAGATGACGAAAATATTAACTCTCAACCATTCATGAGATGGCGTGAAAG

>gi|2477559192|gb|OQ466177.1| Dinobryon similis strain Meokgol112021MS16 ribulose-1,5-bisphosphate carboxylase/oxygenase large subunit (rbcL) gene, partial cds; chloroplast

AAACTTAACAGCATCAATTATTGGAAACGTTTTTGGATTTAAAGCTGTTAAAGCGCTTCGTTTAGAAGATATGCGTATTCCATATGCTTATTTAAAAACTTTCTTAGGCCCAGCATCAGGTGTTATTGTAGAACGTGAAAGACTTGACGTTTTCGGACGCCCACTTTTAGGTGCAACAGTTAAACCAAAATTAGGTCTATCTGGTAAAAACTATGGTCGTGTAGTTTATGAAGGATTAAAAGGTGGTTTAGATTTCTTAAAAGATGACGAAAATATTAACTCTCAACCATTCATGAGATGGCGTGAAAG

>gi|2477559194|gb|OQ466178.1| Dinobryon inclinatum strain CCMP1859 ribulose-1,5-bisphosphate carboxylase/oxygenase large subunit (rbcL) gene, partial cds; chloroplast

AAACTTAACAGCATCAATCATTGGAAACGTTTTTGGATTTAAAGCGGTTAAAGCTCTTCGTTTAGAAGATATGCGTCTTCCGTATGCATATTTAAAAACATTCTTAGGACCTGCATCTGGTGTTATTGTAGAACGTGAAAGACTTGACGTTTTTGGACGTCCACTTTTAGGTGCAACTGTTAAACCAAAATTAGGTCTTTCAGGGAAAAACTATGGTCGTGTTGTTTATGAAGGATTAAAAGGTGGTTTAGACTTCTTAAAAGATGACGAAAACATTAACTCTCAACCATTCATGAGATGGCGTGAAAG

>gi|2477559196|gb|OQ466179.1| Dinobryon inclinatum strain CCMP1860 ribulose-1,5-bisphosphate carboxylase/oxygenase large subunit (rbcL) gene, partial cds; chloroplast

AAACTTAACAGCATCAATCATTGGAAACGTTTTTGGATTTAAAGCGGTTAAAGCTCTTCGTTTAGAAGATATGCGTCTTCCGTATGCATATTTAAAAACATTCTTAGGACCTGCATCTGGTGTTATTGTAGAACGTGAAAGACTTGACGTTTTTGGTCGTCCACTTTTAGGTGCAACTGTTAAACCAAAATTAGGTCTTTCAGGGAAAAACTATGGTCGTGTTGTTTATGAAGGATTAAAAGGTGGTTTAGACTTCTTAAAAGATGACGAAAACATTAACTCTCAACCATTCATGAGATGGCGTGAAAG

>gi|2477559198|gb|OQ466180.1| Dinobryon inclinatum strain CCMP2766 ribulose-1,5-bisphosphate carboxylase/oxygenase large subunit (rbcL) gene, partial cds; chloroplast

AAACTTAACAGCATCAATCATTGGAAACGTTTTTGGATTTAAAGCGGTTAAAGCTCTTCGTTTAGAAGATATGCGTCTTCCGTATGCATATTTAAAAACATTCTTAGGACCTGCATCTGGTGTTATTGTAGAACGTGAAAGACTTGACGTTTTTGGACGTCCACTTTTAGGTGCAACTGTTAAACCAAAATTAGGTCTTTCAGGGAAAAACTATGGTCGTGTTGTTTATGAAGGATTAAAAGGTGGTTTAGACTTCTTAAAAGATGACGAAAACATTAACTCTCAACCATTCATGAGATGGCGTGAAAG

>gi|2477559200|gb|OQ466181.1| Dinobryon inclinatum strain Chojeon011219A ribulose-1,5-bisphosphate carboxylase/oxygenase large subunit (rbcL) gene, partial cds; chloroplast

AAACTTAACAGCATCAATCATTGGAAACGTTTTTGGATTTAAAGCGGTTAAAGCTCTTCGTTTAGAAGATATGCGTCTTCCGTATGCATATTTAAAAACATTCTTAGGACCTGCATCTGGTGTTATTGTAGAACGTGAAAGACTTGACGTTTTTGGACGTCCACTTTTAGGTGCAACTGTTAAACCAAAATTAGGTCTTTCAGGGAAAAACTATGGTCGTGTTGTTTATGAAGGATTAAAAGGTGGTTTAGACTTCTTAAAAGATGACGAAAATATTAACTCTCAACCATTCATGAGATGGCGTGAAAG

>gi|2477559202|gb|OQ466182.1| Dinobryon inclinatum strain Dogwan033018A ribulose-1,5-bisphosphate carboxylase/oxygenase large subunit (rbcL) gene, partial cds; chloroplast

AAACTTAACAGCATCAATCATTGGAAACGTTTTTGGATTTAAAGCGGTTAAAGCTCTTCGTTTAGAAGATATGCGTCTTCCGTATGCATACTTAAAAACATTCTTAGGACCTGCGTCTGGTGTTATTGTAGAACGTGAAAGACTTGACGTTTTTGGACGCCCACTTTTAGGTGCAACTGTTAAACCAAAATTAGGTCTTTCAGGGAAAAACTATGGTCGTGTTGTTTATGAAGGATTAAAAGGTGGTTTAGACTTCTTAAAAGATGACGAAAATATTAACTCTCAACCATTCATGAGATGGCGTGAAAG

>gi|2477559204|gb|OQ466183.1| Dinobryon inclinatum strain Hwalgol120318D ribulose-1,5-bisphosphate carboxylase/oxygenase large subunit (rbcL) gene, partial cds; chloroplast

AAACTTAACAGCATCAATCATTGGAAACGTTTTTGGATTTAAAGCGGTTAAAGCTCTTCGTTTAGAAGATATGCGTCTTCCGTATGCATACTTAAAAACATTCTTAGGACCTGCGTCTGGTGTTATTGTAGAACGTGAAAGACTTGACGTTTTTGGACGCCCACTTTTAGGTGCAACTGTTAAACCAAAATTAGGTCTTTCAGGGAAAAACTATGGTCGTGTTGTTTATGAAGGATTAAAAGGTGGTTTAGACTTCTTAAAAGATGACGAAAATATTAACTCTCAACCATTCATGAGATGGCGTGAAAG

>gi|2477559206|gb|OQ466184.1| Dinobryon inclinatum strain Jijije022318A ribulose-1,5-bisphosphate carboxylase/oxygenase large subunit (rbcL) gene, partial cds; chloroplast

AAACTTAACAGCATCAATCATTGGAAACGTTTTTGGATTTAAAGCGGTTAAAGCTCTTCGTTTAGAAGATATGCGTCTTCCGTATGCATACTTAAAAACATTCTTAGGACCTGCGTCTGGTGTTATTGTAGAACGTGAAAGACTTGACGTTTTTGGACGCCCACTTTTAGGTGCAACTGTTAAACCAAAATTAGGTCTTTCAGGGAAAAACTATGGTCGTGTTGTTTATGAAGGATTAAAAGGTGGTTTAGACTTCTTAAAAGATGACGAAAATATTAACTCTCAACCATTCATGAGATGGCGTGAAAG

>gi|2477559208|gb|OQ466185.1| Dinobryon taiyuanensis strain Dogwan180227HM1 ribulose-1,5-bisphosphate carboxylase/oxygenase large subunit (rbcL) gene, partial cds; chloroplast

AAACTTAACAGCATCAATTATTGGAAACGTTTTTGGATTTAAAGCGGTTAAAGCGCTTCGTTTAGAAGATATGCGTCTTCCATATGCTTATTTAAAAACATTCTTAGGACCTGCCGCTGGTGTTATTGTAGAACGTGAAAGACTTGACGTTTTTGGACGTCCACTTTTAGGTGCAACAGTTAAACCGAAATTAGGTCTTTCAGGAAAAAATTATGGTCGTGTTGTTTATGAAGGATTAAAAGGTGGTTTAGACTTCTTAAAAGATGACGAAAATATTAACTCTCAACCATTCATGAGATGGCGTGAAAG

>gi|2477559210|gb|OQ466186.1| Dinobryon taiyuanensis strain Josoo1ri111620MS1 ribulose-1,5-bisphosphate carboxylase/oxygenase large subunit (rbcL) gene, partial cds; chloroplast

AAACTTAACAGCATCAATTATTGGAAACGTTTTTGGATTTAAAGCGGTTAAAGCGCTTCGTTTAGAAGATATGCGTCTTCCATATGCTTATTTAAAAACATTCTTAGGACCTGCCGCTGGTGTTATTGTAGAACGTGAAAGACTTGACGTTTTTGGACGTCCACTTTTAGGTGCAACAGTTAAACCGAAATTAGGTCTTTCAGGAAAAAATTATGGTCGTGTTGTTTATGAAGGATTAAAAGGTGGTTTAGACTTCTTAAAAGATGACGAAAATATTAACTCTCAACCATTCATGAGATGGCGTGAAAG

>gi|2477559212|gb|OQ466187.1| Dinobryon taiyuanensis strain Yookhoje031321MS4 ribulose-1,5-bisphosphate carboxylase/oxygenase large subunit (rbcL) gene, partial cds; chloroplast

AAACTTAACAGCATCAATTATTGGAAACGTTTTTGGATTTAAAGCGGTTAAAGCGCTTCGTTTAGAAGATATGCGTCTTCCATATGCTTATTTAAAAACATTCTTAGGACCTGCCGCTGGTGTTATTGTAGAACGTGAAAGACTTGACGTTTTTGGACGTCCACTTTTAGGTGCAACAGTTAAACCGAAATTAGGTCTTTCAGGAAAAAATTATGGTCGTGTTGTTTATGAAGGATTAAAAGGTGGTTTAGACTTCTTAAAAGATGACGAAAATATTAACTCTCAACCATTCATGAGATGGCGTGAAAG

>gi|2477559214|gb|OQ466188.1| Dinobryon taiyuanensis strain Wonyong111618A ribulose-1,5-bisphosphate carboxylase/oxygenase large subunit (rbcL) gene, partial cds; chloroplast

AAACTTAACAGCATCAATTATTGGAAACGTTTTTGGATTTAAAGCGGTTAAAGCGCTTCGTTTAGAAGATATGCGTCTTCCATATGCTTATTTAAAAACATTCTTAGGACCTGCCGCTGGTGTTATTGTAGAACGTGAAAGACTTGACGTTTTTGGACGTCCACTTTTAGGTGCAACAGTTAAACCGAAATTAGGTCTTTCAGGAAAAAATTATGGTCGTGTTGTTTATGAAGGATTAAAAGGTGGTTTAGACTTCTTAAAAGATGACGAAAATATTAACTCTCAACCATTCATGAGATGGCGTGAAAG

>gi|2477559216|gb|OQ466189.1| Dinobryon taiyuanensis strain Cheonma111321MS4 ribulose-1,5-bisphosphate carboxylase/oxygenase large subunit (rbcL) gene, partial cds; chloroplast

AAACTTAACAGCATCAATTATTGGAAACGTTTTTGGATTTAAAGCGGTTAAAGCGCTTCGTTTAGAAGATATGCGTCTTCCATATGCTTATTTAAAAACATTCTTAGGACCTGCCGCTGGTGTTATTGTAGAACGTGAAAGACTTGACGTTTTTGGACGTCCACTTTTAGGTGCAACAGTTAAACCGAAATTAGGTCTTTCAGGAAAAAATTATGGTCGTGTTGTTTATGAAGGATTAAAAGGTGGTTTAGACTTCTTAAAAGATGACGAAAATATTAACTCTCAACCATTCATGAGATGGCGTGAAAG

>gi|2477559218|gb|OQ466190.1| Dinobryon taiyuanensis strain Dogwan022718C ribulose-1,5-bisphosphate carboxylase/oxygenase large subunit (rbcL) gene, partial cds; chloroplast

AAACTTAACAGCATCAATTATTGGAAACGTTTTTGGATTTAAAGCGGTTAAAGCGCTTCGTTTAGAAGATATGCGTCTTCCATATGCTTATTTAAAAACATTCTTAGGACCTGCCGCTGGTGTTATTGTAGAACGTGAAAGACTTGACGTTTTTGGACGTCCACTTTTAGGTGCAACAGTTAAACCGAAATTAGGTCTTTCAGGAAAAAATTATGGTCGTGTTGTTTATGAAGGATTAAAAGGTGGTTTAGACTTCTTAAAAGATGACGAAAATATTAACTCTCAACCATTCATGAGATGGCGTGAAAG

>gi|2477559220|gb|OQ466191.1| Dinobryon spinum strain Arong051421MS2 ribulose-1,5-bisphosphate carboxylase/oxygenase large subunit (rbcL) gene, partial cds; chloroplast

AAACTTAACAGCATCAATCATTGGAAACGTTTTTGGATTTAAAGCAGTTAAAGCTCTTCGTTTAGAAGATATGCGTCTTCCATATGCATATTTAAAAACATTCTTAGGACCTGCATCTGGTGTTATTGTAGAACGTGAAAGACTTGACGTTTTTGGACGTCCACTTTTAGGTGCAACTGTTAAACCAAAATTAGGTCTTTCAGGGAAAAACTATGGTCGTGTTGTTTATGAAGGATTAAAAGGTGGTTTAGACTTCTTAAAAGATGACGAAAATATTAACTCTCAACCATTCATGAGATGGCGTGAAAG

>gi|2477559222|gb|OQ466192.1| Dinobryon spinum strain Deokghi042118B ribulose-1,5-bisphosphate carboxylase/oxygenase large subunit (rbcL) gene, partial cds; chloroplast

AAACTTAACAGCATCAATCATTGGAAACGTTTTTGGATTTAAAGCAGTTAAAGCTCTTCGTTTAGAAGATATGCGTCTTCCATATGCATATTTAAAAACATTCTTAGGACCTGCATCTGGTGTTATTGTAGAACGTGAAAGACTTGACGTTTTTGGACGTCCACTTTTAGGTGCAACTGTTAAACCAAAATTAGGTCTTTCAGGGAAAAACTATGGTCGTGTTGTTTATGAAGGATTAAAAGGTGGTTTAGACTTCTTAAAAGATGACGAAAATATTAACTCTCAACCATTCATGAGATGGCGTGAAAG

>gi|2477559224|gb|OQ466193.1| Dinobryon spinum strain Gwangdaeje031221MS4 ribulose-1,5-bisphosphate carboxylase/oxygenase large subunit (rbcL) gene, partial cds; chloroplast

AAACTTAACAGCATCAATCATTGGAAACGTTTTTGGATTTAAAGCAGTTAAAGCTCTTCGTTTAGAAGATATGCGTCTTCCATATGCATATTTAAAAACATTCTTAGGACCTGCATCTGGTGTTATTGTAGAACGTGAAAGACTTGACGTTTTTGGACGTCCACTTTTAGGTGCAACTGTTAAACCAAAATTAGGTCTTTCAGGGAAAAACTATGGTCGTGTTGTTTATGAAGGATTAAAAGGTGGTTTAGACTTCTTAAAAGATGACGAAAATATTAACTCTCAACCATTCATGAGATGGCGTGAAAG

>gi|2477559226|gb|OQ466194.1| Dinobryon spinum strain Geumgok020610D ribulose-1,5-bisphosphate carboxylase/oxygenase large subunit (rbcL) gene, partial cds; chloroplast

AAACTTAACAGCATCAATCATTGGAAACGTTTTTGGATTTAAAGCAGTTAAAGCTCTTCGTTTAGAAGATATGCGTCTTCCATATGCATATTTAAAAACATTCTTAGGACCTGCATCTGGTGTTATTGTAGAACGTGAAAGACTTGACGTTTTTGGACGTCCACTTTTAGGTGCAACTGTTAAACCAAAATTAGGTCTTTCAGGGAAAAACTATGGTCGTGTTGTTTATGAAGGATTAAAAGGTGGTTTAGACTTCTTAAAAGATGACGAAAATATTAACTCTCAACCATTCATGAGATGGCGTGAAAG

>gi|2477559228|gb|OQ466195.1| Dinobryon spinum strain Myeoseul031718A ribulose-1,5-bisphosphate carboxylase/oxygenase large subunit (rbcL) gene, partial cds; chloroplast

AAACTTAACAGCATCAATCATTGGAAACGTTTTTGGATTTAAAGCAGTTAAAGCTCTTCGTTTAGAAGATATGCGTCTTCCATATGCATATTTAAAAACATTCTTAGGACCTGCATCTGGTGTTATTGTAGAACGTGAAAGACTTGACGTTTTTGGACGTCCACTTTTAGGTGCAACTGTTAAACCAAAATTAGGTCTTTCAGGGAAAAACTATGGTCGTGTTGTTTATGAAGGATTAAAAGGTGGTTTAGACTTCTTAAAAGATGACGAAAATATTAACTCTCAACCATTCATGAGATGGCGTGAAAG

>gi|2477559230|gb|OQ466196.1| Dinobryon spinum strain Wonyong122822S2 ribulose-1,5-bisphosphate carboxylase/oxygenase large subunit (rbcL) gene, partial cds; chloroplast

AAACTTAACAGCATCAATCATTGGAAACGTTTTTGGATTTAAAGCAGTTAAAGCTCTTCGTTTAGAAGATATGCGTCTTCCATATGCATATTTAAAAACATTCTTAGGACCTGCATCTGGTGTTATTGTAGAACGTGAAAGACTTGACGTTTTTGGACGTCCACTTTTAGGTGCAACTGTTAAACCAAAATTAGGTCTTTCAGGGAAAAACTATGGTCGTGTTGTTTATGAAGGATTAAAAGGTGGTTTAGACTTCTTAAAAGATGACGAAAATATTAACTCTCAACCATTCATGAGATGGCGTGAAAG

>gi|2477559232|gb|OQ466197.1| Dinobryon sertularia var. thyrsoideum strain Chosan041710C ribulose-1,5-bisphosphate carboxylase/oxygenase large subunit (rbcL) gene, partial cds; chloroplast

AAACTTAACAGCATCAATCATTGGAAACGTTTTTGGATTTAAAGCAGTTAAAGCTCTTCGTTTAGAAGATATGCGTCTTCCATATGCATATTTAAAAACATTCTTAGGACCTGCATCTGGTGTTATTGTAGAACGTGAAAGACTTGACGTTTTTGGACGTCCACTTTTAGGTGCAACTGTTAAACCAAAATTAGGTCTTTCAGGGAAAAACTATGGTCGTGTTGTTTATGAAGGATTAAAAGGTGGTTTAGACTTCTTAAAAGATGACGAAAATATTAACTCTCAACCATTCATGAGATGGCGTGAAAG

>gi|2477559234|gb|OQ466198.1| Dinobryon sertularia var. thyrsoideum strain Geumgang111321MS6 ribulose-1,5-bisphosphate carboxylase/oxygenase large subunit (rbcL) gene, partial cds; chloroplast

AAACTTAACAGCATCAATCATTGGAAACGTTTTTGGATTTAAAGCAGTTAAAGCTCTTCGTTTAGAAGATATGCGTCTTCCATATGCATATTTAAAAACATTCTTAGGACCTGCATCTGGTGTTATTGTAGAACGTGAAAGACTTGACGTTTTTGGACGTCCACTTTTAGGTGCAACTGTTAAACCAAAATTAGGTCTTTCAGGGAAAAACTATGGTCGTGTTGTTTATGAAGGATTAAAAGGTGGTTTAGACTTCTTAAAAGATGACGAAAATATTAACTCTCAACCATTCATGAGATGGCGTGAAAG

>gi|2477559236|gb|OQ466199.1| Dinobryon sertularia var. thyrsoideum strain Yanghari040921MS12 ribulose-1,5-bisphosphate carboxylase/oxygenase large subunit (rbcL) gene, partial cds; chloroplast

AAACTTAACAGCATCAATCATTGGAAACGTTTTTGGATTTAAAGCAGTTAAAGCTCTTCGTTTAGAAGATATGCGTCTTCCATATGCATATTTAAAAACATTCTTAGGACCTGCATCTGGTGTTATTGTAGAACGTGAAAGACTTGACGTTTTTGGACGTCCACTTTTAGGTGCAACTGTTAAACCAAAATTAGGTCTTTCAGGGAAAAACTATGGTCGTGTTGTTTATGAAGGATTAAAAGGTGGTTTAGACTTCTTAAAAGATGACGAAAATATTAACTCTCAACCATTCATGAGATGGCGTGAAAG

>gi|2477559238|gb|OQ466200.1| Dinobryon cylindricum var. palustre strain Joogyo2je043021MS12 ribulose-1,5-bisphosphate carboxylase/oxygenase large subunit (rbcL) gene, partial cds; chloroplast

AAACTTAACAGCATCAATCATTGGAAACGTTTTTGGATTTAAAGCAGTTAAAGCTCTTCGTTTAGAAGATATGCGTCTGCCATATGCATATTTAAAAACATTCTTAGGACCTGCGTCTGGTGTTATTGTAGAACGTGAAAGACTTGACGTTTTTGGACGTCCACTTTTAGGTGCAACTGTTAAACCAAAATTAGGTCTTTCAGGGAAAAACTATGGTCGTGTTGTTTATGAAGGATTAAAAGGTGGTTTAGACTTCTTAAAAGATGACGAAAATATTAACTCTCAACCATTCATGAGATGGCGTGAAAG

>gi|2477559240|gb|OQ466201.1| Dinobryon cylindricum var. palustre strain Gapa120521MS4 ribulose-1,5-bisphosphate carboxylase/oxygenase large subunit (rbcL) gene, partial cds; chloroplast

AAACTTAACAGCATCAATCATTGGAAACGTTTTTGGATTTAAAGCAGTTAAAGCTCTTCGTTTAGAAGATATGCGTCTGCCATATGCATATTTAAAAACATTCTTAGGACCTGCGTCTGGTGTTATTGTAGAACGTGAAAGACTTGACGTTTTTGGACGTCCACTTTTAGGTGCAACTGTTAAACCAAAATTAGGTCTTTCAGGGAAAAACTATGGTCGTGTTGTTTATGAAGGATTAAAAGGTGGTTTAGACTTCTTAAAAGATGACGAAAATATTAACTCTCAACCATTCATGAGATGGCGTGAAAG

>gi|2477559242|gb|OQ466202.1| Dinobryon cylindricum var. palustre strain Dowon111321MS2 ribulose-1,5-bisphosphate carboxylase/oxygenase large subunit (rbcL) gene, partial cds; chloroplast

AAACTTAACAGCATCAATCATTGGAAACGTTTTTGGATTTAAAGCAGTTAAAGCTCTTCGTTTAGAAGATATGCGTCTGCCATATGCATATTTAAAAACATTCTTAGGACCTGCGTCTGGTGTTATTGTAGAACGTGAAAGACTTGACGTTTTTGGACGTCCACTTTTAGGTGCAACTGTTAAACCAAAATTAGGTCTTTCAGGGAAAAACTATGGTCGTGTTGTTTATGAAGGATTAAAAGGTGGTTTAGACTTCTTAAAAGATGACGAAAATATTAACTCTCAACCATTCATGAGATGGCGTGAAAG

>gi|2477559244|gb|OQ466203.1| Dinobryon cylindricum var. palustre strain Bonghwa040718C ribulose-1,5-bisphosphate carboxylase/oxygenase large subunit (rbcL) gene, partial cds; chloroplast

AAACTTAACAGCATCAATCATTGGAAACGTTTTTGGATTTAAAGCAGTTAAAGCTCTTCGTTTAGAAGATATGCGTCTGCCATATGCATATTTAAAAACATTCTTAGGACCTGCGTCTGGTGTTATTGTAGAACGTGAAAGACTTGACGTTTTTGGACGTCCACTTTTAGGTGCAACTGTTAAACCAAAATTAGGTCTTTCAGGGAAAAACTATGGTCGTGTTGTTTATGAAGGATTAAAAGGTGGTTTAGACTTCTTAAAAGATGACGAAAATATTAACTCTCAACCATTCATGAGATGGCGTGAAAG

>gi|2477559246|gb|OQ466204.1| Dinobryon cylindricum var. palustre strain Baeteo110621MS2 ribulose-1,5-bisphosphate carboxylase/oxygenase large subunit (rbcL) gene, partial cds; chloroplast

AAACTTAACAGCATCAATCATTGGAAACGTTTTTGGATTTAAAGCAGTTAAAGCTCTTCGTTTAGAAGATATGCGTCTGCCATATGCATATTTAAAAACATTCTTAGGACCTGCGTCTGGTGTTATTGTAGAACGTGAAAGACTTGACGTTTTTGGACGTCCACTTTTAGGTGCAACTGTTAAACCAAAATTAGGTCTTTCAGGGAAAAACTATGGTCGTGTTGTTTATGAAGGATTAAAAGGTGGTTTAGACTTCTTAAAAGATGACGAAAATATTAACTCTCAACCATTCATGAGATGGCGTGAAAG

>gi|2477559248|gb|OQ466205.1| Dinobryon ningwuensis strain Yeokjae092718B ribulose-1,5-bisphosphate carboxylase/oxygenase large subunit (rbcL) gene, partial cds; chloroplast

AAACTTAACAGCATCAATCATTGGAAACGTTTTCGGATTTAAAGCCGTTAAAGCTTTACGTTTAGAAGATATGCGTCTTCCATATGCTTATTTAAAAACTTTCTTAGGACCAGCATCTGGTGTTATTGTAGAACGTGAAAGACTTGACGTTTTTGGACGTCCACTTTTAGGTGCAACAGTTAAACCAAAATTAGGTCTTTCAGGAAAAAACTATGGTCGTGTAGTATATGAAGGATTAAAAGGTGGTTTAGACTTCTTAAAAGATGACGAAAACATTAACTCTCAACCATTCATGAGATGGCGTGAAAG

>gi|2477559250|gb|OQ466206.1| Dinobryon ningwuensis strain Yeongrangho111321MS1 ribulose-1,5-bisphosphate carboxylase/oxygenase large subunit (rbcL) gene, partial cds; chloroplast

AAACTTAACAGCATCAATCATTGGAAACGTTTTCGGATTTAAAGCCGTTAAAGCTTTACGTTTAGAAGATATGCGTCTTCCATATGCTTATTTAAAAACTTTCTTAGGACCAGCATCTGGTGTTATTGTAGAACGTGAAAGACTTGACGTTTTTGGACGTCCACTTTTAGGTGCAACAGTTAAACCAAAATTAGGTCTTTCAGGAAAAAACTATGGTCGTGTAGTATATGAAGGATTAAAAGGTGGTTTAGACTTCTTAAAAGATGACGAAAACATTAACTCTCAACCATTCATGAGATGGCGTGAAAG

>gi|2477559252|gb|OQ466207.1| Dinobryon ningwuensis strain Myeoseul031718C ribulose-1,5-bisphosphate carboxylase/oxygenase large subunit (rbcL) gene, partial cds; chloroplast

AAACTTAACAGCATCAATCATTGGAAACGTTTTCGGATTTAAAGCCGTTAAAGCTTTACGTTTAGAAGATATGCGTCTTCCATATGCTTATTTAAAAACTTTCTTAGGACCAGCATCTGGTGTTATTGTAGAACGTGAAAGACTTGACGTTTTTGGACGTCCACTTTTAGGTGCAACAGTTAAACCAAAATTAGGTCTTTCAGGAAAAAACTATGGTCGTGTAGTATATGAAGGATTAAAAGGTGGTTTAGACTTCTTAAAAGATGACGAAAACATTAACTCTCAACCATTCATGAGATGGCGTGAAAG

>gi|2477559254|gb|OQ466208.1| Dinobryon balticum strain CCMP1766 ribulose-1,5-bisphosphate carboxylase/oxygenase large subunit (rbcL) gene, partial cds; chloroplast

TAACTTAACAGCATCTATTATTGGTAACGTATTTGGATTTAAGGCTGTTAAGGCTTTACGTTTAGAAGATATGCGTCTTCCATATGCTTACTTAAAAGGATTCCTAGGACCAGCTTGTGGAATTATTGTAGAACGTGAAAGACTTGATGTATTTGGACGTCCTCTTTTAGGTGCTACTGTAAAACCAAAACTAGGTCTTTCTGGTAAGAACTACGGTCGTGTAGTTTATGAAGGATTAAAAGGTGGTTTAGACTTCTTAAAAGATGATGAGAATATTAACTCTCAACCATTCATGAGATGGCGTGAAAG

>gi|2523494768|gb|OQ064798.1| Spiniferomonas trioralis strain CZ146B ribulose-1,5-biphosphate carboxylase/oxygenase large subunit (rbcL) gene, partial cds; plastid

TAACTTAACTGCTTCAATTATCGGAAACGTATTTGGATTTAAAGCTGTTAAATGTTTACGTTTAGAAGATATGCGTCTTCCTTATGCATACTTAAAAACATTCTTAGGTCCTGCTAGTGGTGTTATTGTTGAACGTGAAAGATTAGACGTATTCGGACGTCCTTTATTAGGTGCTACTGTTAAACCTAAATTAGGTTTATCTGGTAAAAACTATGGACGTGTAGTTTATGAAGGATTAAAAGGTGGTTTAGACTTCTTAAAAGATGATGAAAACATCAATTCACAACCATTCATGAGATGGCGTGAAAG

>gi|2523494770|gb|OQ064799.1| Chrysosphaerella coronacircumspina strain CZ28M ribulose-1,5-biphosphate carboxylase/oxygenase large subunit (rbcL) gene, partial cds; plastid

TAACTTAACTGCATCAATTATCGGAAACGTATTTGGTTTTAAAGCAGTTAAATGTCTTCGTTTAGAAGATATGCGTCTTCCTTACGCTTATTTAAAAACTTTTATTGGACCTGCAGCTGGGGTTATTGTAGAACGTGAAAGATTAGATGTTTTTGGACGTCCTCTATTAGGTGCTACTGTTAAACCAAAATTAGGTCTTTCTGGAAAAAACTATGGTCGTGTAGTTTATGAAGGATTAAAAGGTGGTTTAGATTTCTTAAAAGATGATGAAAATATCAATTCACAACCTTTTATGAGATGGCGTGAAAG

>gi|2771091379|gb|OR663905.1| Chrysophyceae sp. strain CZ_61A ribulose-1,5-biphosphate carboxylase/oxygenase large subunit (rbcL) gene, partial cds; chloroplast

TAACTTAACAGCATCTATCATCGGAAACGTATTTGGTTTCAAAGCCGTTAAAGCTCTTCGTTTAGAAGATATGCGTCTTCCTTACGCATACTTAAAAACATTCTTAGGTCCTGCTGCTGGTGTAATTGTAGAACGTGAAAGACTTGACGTATTCGGACGTCCTTTATTAGGAGCAACAGTAAAACCTAAATTAGGTCTTTCTGGTAAAAACTACGGACGTGTTGTTTATGAAGGTTTAAGAGGTGGTTTAGATTTCTTAAAAGATGACGAAAACATTAACTCTCAACCATTCATGAGATGGCGCGAAAG

>gi|2771091381|gb|OR663906.1| Chrysophyceae sp. strain CZ_54B ribulose-1,5-biphosphate carboxylase/oxygenase large subunit (rbcL) gene, partial cds; chloroplast

TAACTTAACAGCATCTATCATCGGAAACGTATTTGGTTTCAAAGCCGTTAAAGCTCTTCGTTTAGAAGATATGCGTCTTCCTTACGCATACTTAAAAACATTCTTAGGTCCTGCTGCTGGTGTAATTGTAGAACGTGAAAGACTTGACGTATTCGGACGTCCTTTATTAGGAGCAACAGTAAAACCTAAATTAGGTCTTTCTGGTAAAAACTACGGACGTGTTGTTTATGAAGGTTTAAGAGGTGGTTTAGATTTCTTAAAAGATGACGAAAACATTAACTCTCAACCATTCATGAGATGGCGCGAAAG

>gi|2771091383|gb|OR663907.1| Chrysotilos ferrea strain AT_26A ribulose-1,5-biphosphate carboxylase/oxygenase large subunit (rbcL) gene, partial cds; chloroplast

TAACTTAACAGCATCTATTATCGGTAACGTATTCGGATTCAAAGCGGTTAAATGTCTTCGTTTAGAAGATATGCGTATTCCTTTTGGTTACTTAAAAACTTTTATTGGACCAGCAGCAGGAGTTATTGTAGAACGTGAAAGACTTGACGTATTCGGACGTCCTCTATTAGGAGCAACTGTTAAACCAAAATTAGGTCTTTCTGGTAAAAACTACGGACGTGTAGTTTATGAAGGATTAAAAGGTGGTTTAGACTTCTTAAAAGATGACGAAAACATCAACTCTCAACCATTCATGAGATGGCGTGAAAG

>gi|6006785|gb|AF155877.1| Chrysocapsa vernalis ribulose-1,5-bisphosphate carboxylase/oxygenase large subunit (rbcL) gene, partial cds; chloroplast gene for chloroplast product

TAACTTAACAGCATCTATTATTGGGAACGTATTTGGTTTCAAAGCTGTAAAATGTCTTCGTTTAGAAGATATGCGTATTCCTTTTGCCTACTTAAAAACTTTCATTGGTCCTGCTGCTGGAGTTATTGTAGAACGTGAAAGACTTGACGTATTTGGACGTCCTTTATTAGGAGCAACTGTAAAACCTAAATTAGGTCTTTCAGGAAAAAACTATGGTCGTGTAGTTTATGAAGGATTAAAAGGTGGTTTAGACTTCTTAAAAGATGACGAAAATATTAACTCTCAACCATTCATGAGATGGCGTGAAAG

>gi|631776895|emb|HG315743.1| Chrysosphaerella rotundata plastid partial rbcL gene for ribulose-1,5-bisphosphate carboxylase/oxygenase, large subunit, strain S89.C4

TAACTTAACTGCATCAATTATCGGAAACGTATTCGGTTTCAAAGCTGTTAAATGTCTTCGTTTAGAAGATATGCGTCTTCCTTATGCTTACTTAAAAACATTTTTAGGACCTGCTGCTGGTGTTGTTGTAGAACGTGAAAGATTAGACGTATTTGGACGTCCTCTATTAGGTGCTACTGTTAAACCTAAATTAGGTCTTTCTGGAAAAAACTATGGTCGTGTAGTTTATGAAGGATTAAAAGGTGGTTTAGATTTCTTAAAAGATGATGAAAATATCAATTCGCAAGCTTTTATGAGATGGCGTGAAAG

>gi|2895139|gb|AF015570.1| Chrysolepidomonas dendrolepidota ribulose-1,5-bisphosphate carboxylase/oxygenase large subunit (rbcL) gene, chloroplast gene encoding chloroplast protein, partial cds

AAACTTAACAGCATCTATTATTGGTAACGTTTTCGGTTTTAAAGCTGTAAAATGTCTTCGTTTAGAAGATATGCGTCTTCCTTACGCTTACTTAAAAACATTCATTGGGCCAGCTAGTGGTGTTATCGTAGAACGTGAAAGACTTGACGTTTTCGGTCGTCCTCTTTTAGGTGCTACAGTTAAACCAAAATTAGGTCTTTCAGGAAAAAACTATGGTCGTGTAGTTTATGAAGGATTAAAAGGTGGTTTAGATTTCTTAAAAGATGANGAAAACATCAACTCTCAAGCTTTCATGAGATGGCGCGAAAG

>gi|2895141|gb|AF015571.1| Epipyxis pulchra ribulose-1,5-bisphosphate carboxylase/oxygenase large subunit (rbcL) gene, chloroplast gene encoding chloroplast protein, partial cds

AAACTTAACAGCATCTATTATCGGAAACGTTTTTGGATTCAAAGCTGTTAAATGTTTACGTTTAGAAGATATGCGTCTTCCTTACGCTTACTTAAAAACGTTTATTGGTCCAGCATCTGGAGTTATTGTAGAACGTGAAAGACTTGACGTTTTTGGTCGTCCATTATTAGGTGCTACTGTTAAACCAAAATTAGGTCTTTCAGGAAAAAACTATGGACGTGTAGTTTATGAAGGATTAAGAGGTGGATTAGATTTCTTAAAAGATGACGAAAACATCAACTCTCAACCATTCATGAGATGGCGTGAAAG

>gi|2895143|gb|AF015572.1| Hibberdia magna ribulose-1,5-bisphosphate carboxylase/oxygenase large subunit (rbcL) gene, chloroplast gene encoding chloroplast protein, partial cds

TAACTTAACAGCATCTATTATCGGTAACGTATTTGGATTCAAAGCAGTTAAAGCTCTTCGTTTAGAAGATATGCGTATTCCTTATGCTTACTTAAAAACTTTCATTGGGCCAGCTGCTGGAGTTATTGTAGAACGTGAAAGATTAGACGTATTCGGACGTCCTTTATTAGGAGCTACTGTTAAACCTAAATTAGGTCTTTCTGGTAAAAACTATGGTCGTGTAGTTTATGAAGGATTAAAAGGTGGTTTAGACTTCTTAAAAGATGACGAAAACATTAACTCTCAACCATTCATNNNNNGGCGTGAAAG

>gi|6006783|gb|AF155876.1| Chromulina nebulosa ribulose-1,5-bisphosphate carboxylase/oxygenase large subunit (rbcL) gene, partial cds; chloroplast gene for chloroplast product

CAATTTAACAGCCTCAATTATCGGAAACGTATTTGGTTTCAAAGCTGTAAAAGCTCTTCGTCTTGAAGATATGCGTATCCCATACGGATATTTAAAAACTTTCTTAGGCCCTGCAACAGGAGTTATCGTAGAACGTGAAAGACTTGATATTTTTGGTCGTCCATTATTAGGTGCAACAGTTAAACCAAAATTAGGTCTTTCTGGAAAAAACTATGGTCGTGTTGTATACGAAGGATTAAGAGGTGGTCTTGATTTCTTAAAAGATGACGAAAATATTAACTCTCAACCATTCATGAGATGGCGTGAAAG

>gi|631776897|emb|HG315744.1| Chrysosphaerella brevispina plastid partial rbcL gene for ribulose-1,5-bisphosphate carboxylase/oxygenase, large subunit, strain S74.D5

TAACTTAACTGCATCAATTATCGGAAACGTATTTGGTTTTAAAGCTGTTAAATGTCTTCGTTTAGAAGATATGCGTCTTCCTTATGCTTACTTAAAAACATTTTTAGGACCTGCTGCTGGTGTTGTTGTAGAACGTGAAAGATTAGACGTATTTGGACGTCCTCTATTAGGTGCTACTGTTAAACCTAAATTAGGTCTTTCTGGAAAAAACTATGGTCGTGTAGTTTATGAAGGATTAAAAGGTGGTTTAGATTTCTTAAAAGATGATGAAAATATCAATTCACAAGCTTTTATGAGATGGCGTGAAAG

>gi|666689791|gb|KF443039.1| Chrysopodocystis socialis strain AC38 ribulose-1,5-bisphosphate carboxylase/oxygenase large subunit (rbcL) gene, partial cds; chloroplast

AAATTTAACAGCTTCAATTATCGGGAACGTATTTGGCTTTAAAGCGGTAAAAGCTCTTCGTCTTGAAGACATGCGTATACCGTATGCGTACTTAAAAACTTTCCAAGGACCTGCAACTGGAGTAATTGTAGAGCGTGAGCGTCTTGATACTTTTGGTAGGCCTATATTAGGTGCCACTGTAAAACCTAAATTAGGTTTGTCCGGAAAAAATTACGGCCGAGTAGTTTATGAAGGTTTACGTGGCGGTTTAGACTTTCTAAAAGATGACGAGAATATTAACTCTCAACCGTTTATGCGCTGGCGTGAACG

>gi|2523494772|gb|OQ064800.1| Dermatochrysis reticulata strain UK18 ribulose-1,5-biphosphate carboxylase/oxygenase large subunit (rbcL) gene, partial cds; plastid

TAACTTAACAGCATCTCTTATCGGTAACGTATTCGGATTTAAAGCTGTTAAATGTCTTCGTTTAGAAGATATGCGTATTCCTTATGCTTATTTAAAAACATTTATTGGTCCAGCAGCAGGAGTTATCGTAGAACGTGAAAGACTTGACGTATTCGGACGTCCTTTATTAGGAGCAACAGTTAAACCAAAATTAGGTCTTTCTGGTAAAAACTATGGTCGTGTAGTTTATGAAGGATTAAAAGGTGGTTTAGATTTCTTAAAAGATGACGAAAATATTAACTCTCAACCATTCATGAGATGGCGTGAAAG

>gi|2557605428|gb|OR284295.1| Hydrurus foetidus strain Fenhe ribulose-1,5-bisphosphate carboxylase/oxygenase large subunit (rbcL) gene, partial cds; chloroplast

TAACTTAACTGCTTCTATTATCGGAAACGTATTTGGATTTAAAGCTGTTAAATGTCTTCGTTTAGAAGATATGCGTCTACCTTATGCTTACTTAAAAACTTTCTTAGGTCCAGCTGCTGGAGTTATCGTTGAACGTGAAAGACTTGACATTTTTGGACGTCCTTTATTAGGAGCAACTGTTAAACCTAAATTAGGTCTTTCTGGTAAAAACTACGGTCGTGTAGTTTACGAAGGATTAAAAGGTGGTTTAGACTTCTTAAAAGATGACGAAAACATTAACTCTCAACCATTCATGAGATGGCGTGAAAG

>gi|2722125399|gb|PP722689.1| Mallomonas sp. NMartynenko-2024a strain 329P ribulose-1,5-bisphosphate carboxylase/oxygenase large subunit (rbcL) gene, partial cds; plastid

GAACTTAACAGCTTCAATTATTGGAAACGTTTTCGGATTTAAAGCTGTAAAAGCTTTACGTTTAGAAGATATGCGTATTCCATTTGCTTACTTAAAAACTTTCTTAGGTCCAGCAACTGGAGTTATCGTTGAACGTGAAAGAATGGATGTTTTTGGACGTCCTCTATTAGGAGCTACTGTTAAACCAAAATTAGGTCTTTCAGGAAAAAACTATGGTCGTGTAGTTTATGAAGGATTAAAAGGTGGTTTAGATTTCTTAAAAGATGACGAAAACATTAATTCTCAACCTTTCATGAGATGGCGTGAAAG

>gi|2722125401|gb|PP722690.1| Mallomonas sp. NMartynenko-2024a strain 312P ribulose-1,5-bisphosphate carboxylase/oxygenase large subunit (rbcL) gene, partial cds; plastid

GAACTTAACAGCTTCAATTATTGGAAACGTTTTCGGATTTAAAGCTGTAAAAGCTTTACGTTTAGAAGATATGCGTATTCCATTTGCTTACTTAAAAACTTTCTTAGGTCCAGCAACTGGAGTTATCGTTGAACGTGAAAGAATGGATGTTTTTGGACGTCCTCTATTAGGAGCTACTGTTAAACCAAAATTAGGTCTTTCAGGAAAAAACTATGGTCGTGTAGTTTATGAAGGATTAAAAGGTGGTTTAGATTTCTTAAAAGATGACGAAAACATTAATTCTCAACCTTTCATGAGATGGCGTGAAAG

>gi|2722125403|gb|PP722691.1| Mallomonas sp. NMartynenko-2024b strain 892P ribulose-1,5-bisphosphate carboxylase/oxygenase large subunit (rbcL) gene, partial cds; plastid

GAACTTAACAGCGTCAATTATTGGAAACGTTTTCGGATTTAAAGCTGTAAAAGCTTTACGTTTAGAAGATATGCGCATTCCATATGCTTACTTAAAAACTTTTTTAGGTCCAGCAACTGGAGTTATCGTTGAACGTGAAAGAATGGATGTATTTGGACGTCCTCTATTAGGAGCTACTGTTAAACCAAAATTAGGTCTTTCAGGTAAAAACTATGGTCGTGTAGTTTACGAAGGATTAAAAGGTGGTTTAGATTTCTTAAAAGATGATGAAAATATCAATTCTCAACCTTTCATGAGATGGCGTGAAAG

>gi|2722125405|gb|PP722692.1| Mallomonas sp. NMartynenko-2024b strain B12/22 ribulose-1,5-bisphosphate carboxylase/oxygenase large subunit (rbcL) gene, partial cds; plastid

GAACTTAACAGCGTCAATTATTGGAAACGTTTTCGGATTTAAAGCTGTAAAAGCTTTACGTTTAGAAGATATGCGCATTCCATATGCTTACTTAAAAACTTTTTTAGGTCCAGCAACTGGAGTTATCGTTGAACGTGAAAGAATGGATGTATTTGGACGTCCTCTATTAGGAGCTACTGTTAAACCAAAATTAGGTCTTTCAGGTAAAAACTATGGTCGTGTAGTTTACGAAGGATTAAAAGGTGGTTTAGATTTCTTAAAAGATGATGAAAATATCAATTCTCAACCTTTCATGAGATGGCGTGAAAG

>gi|2722125407|gb|PP722693.1| Mallomonas sp. NMartynenko-2024b strain VNG 2136 ribulose-1,5-bisphosphate carboxylase/oxygenase large subunit (rbcL) gene, partial cds; plastid

GAACTTAACAGCGTCAATTATCGGAAACGTTTTCGGATTTAAAGCTGTAAAAGCTTTACGTTTAGAAGATATGCGCATTCCATATGCTTACTTAAAAACTTTTTTAGGTCCAGCAACTGGAGTTATCGTTGAACGTGAAAGAATGGATGTATTTGGACGTCCTCTATTAGGAGCTACTGTTAAACCAAAATTAGGTCTTTCAGGTAAAAACTATGGTCGTGTAGTTTACGAAGGATTAAAAGGTGGTTTAGATTTCTTAAAAGATGATGAAAATATCAATTCTCAACCTTTCATGAGATGGCGTGAAAG

>gi|2722125409|gb|PP722694.1| Mallomonas sp. NMartynenko-2024b strain VNG 2032 ribulose-1,5-bisphosphate carboxylase/oxygenase large subunit (rbcL) gene, partial cds; plastid

GAACTTAACAGCGTCAATTATTGGAAACGTTTTCGGATTTAAAGCTGTAAAAGCTTTACGTTTAGAAGATATGCGCATTCCATATGCTTACTTAAAAACTTTTTTAGGTCCAGCAACTGGAGTTATCGTTGAACGTGAAAGAATGGATGTATTTGGACGTCCTCTATTAGGAGCTACTGTTAAACCAAAATTAGGTCTTTCAGGTAAAAACTATGGTCGTGTAGTTTACGAAGGATTAAAAGGTGGTTTAGATTTCTTAAAAGATGATGAAAATATCAATTCTCAACCTTTCATGAGATGGCGTGAAAG

>gi|2722125411|gb|PP722695.1| Mallomonas pseudomatvienkoae strain 297Yu ribulose-1,5-bisphosphate carboxylase/oxygenase large subunit (rbcL) gene, partial cds; plastid

AAACTTAACAGCTTCAATTATTGGAAACGTTTTCGGATTTAAAGCTGTAAAAGCTTTACGTTTAGAAGATATGCGTATTCCATATGCTTACTTAAAAACTTTCTTAGGTCCAGCAACTGGAGTTATTGTTGAACGTGAAAGAATGGATGTTTTTGGACGTCCTATGTTAGGAGCTACTGTTAAACCAAAATTAGGTCTTTCAGGAAAAAACTATGGTCGTGTAGTTTATGAAGGATTAAAAGGTGGTCTAGATTTCTTAAAAGATGATGAAAACATCAATTCTCAACCTTTCATGAGATGGCGTGAACG

>gi|2722125413|gb|PP722696.1| Mallomonas pseudomatvienkoae strain VNG 2035 ribulose-1,5-bisphosphate carboxylase/oxygenase large subunit (rbcL) gene, partial cds; plastid

AAACTTAACAGCTTCAATTATTGGAAACGTTTTCGGATTTAAAGCTGTAAAAGCTTTACGTTTAGAAGATATGCGTATTCCATATGCTTACTTAAAAACTTTCTTAGGTCCAGCAACTGGAGTTATCGTTGAACGTGAAAGAATGGATGTTTTTGGACGTCCTATGTTAGGAGCTACTGTTAAACCAAAATTAGGTCTTTCAGGAAAAAACTATGGTCGTGTAGTTTATGAAGGATTAAAAGGTGGTCTAGATTTCTTAAAAGATGATGAAAACATCAATTCTCAACCTTTCATGAGATGGCGTGAACG

>gi|2722125415|gb|PP722697.1| Mallomonas pseudomatvienkoae strain VNG 2129 ribulose-1,5-bisphosphate carboxylase/oxygenase large subunit (rbcL) gene, partial cds; plastid

AAACTTAACAGCTTCAATTATTGGAAACGTTTTCGGATTTAAAGCTGTAAAAGCTTTACGTTTAGAAGATATGCGTATTCCATATGCTTACTTAAAAACTTTCTTAGGTCCAGCAACTGGAGTTATCGTTGAACGTGAAAGAATGGATGTTTTTGGACGTCCTATGTTAGGAGCTACTGTTAAACCAAAATTAGGTCTTTCAGGAAAAAACTATGGTCGTGTAGTTTATGAAGGATTAAAAGGTGGTCTAGATTTCTTAAAAGATGATGAAAACATCAATTCTCAACCTTTCATGAGATGGCGTGAACG

>gi|2755083954|gb|PP966934.1| Mallomonas matvienkoae strain R064 ribulose-1,5-bisphosphate carboxylase/oxygenase large subunit (rbcL) gene, partial cds; chloroplast

AAACTTAACAGCATCAATTATCGGAAACGTTTTTGGTTTCAAAGCTGTTAAAGCTTTACGTTTAGAAGATATGCGTATTCCTTATGCTTATTTAAAAACGTTTATCGGACCAGCAACTGGAGTTATTGTTGAACGTGAAAGAATGGATGTTTTTGGACGTCCTCTATTAGGTGCTACTGTTAAACCAAAATTAGGTCTTTCAGGAAAAAATTATGGTCGTGTAGTTTACGAAGGATTAAAAGGTGGTTTAGATTTCTTAAAAGATGATGAAAATATTAATTCTCAAGCTTTTATGAGATGGCGTGAAAG

>gi|2778273102|gb|PQ064525.1| Mallomonas mangofera var. foveata strain 130Yu ribulose-1,5-bisphosphate carboxylase/oxygenase large subunit (rbcL) gene, partial cds; chloroplast

TAACTTAACAGCATCTATTATCGGTAACGTTTTTGGATTCAAAGCAGTAAAAGCATTACGTTTAGAAGATATGCGTATTCCTTATGCATACTTAAAAACATTCCAAGGTCCAGCTACAGGAGTTGTTGTAGAACGTGAAAGAATGGATGTATTCGGACGTCCTTTATTAGGAGCAACTGTTAAACCAAAATTAGGTCTTTCTGGTAAAAACTATGGTCGTGTAGTTTATGAAGGATTAAAAGGTGGATTAGACTTCTTAAAAGATGACGAAAACATTAACTCTCAACCATTCATGAGATGGCGTGAAAG

>gi|2778273104|gb|PQ064526.1| Mallomonas mangofera var. foveata strain 25Yu ribulose-1,5-bisphosphate carboxylase/oxygenase large subunit (rbcL) gene, partial cds; chloroplast

TAACTTAACAGCATCTATTATCGGTAACGTTTTTGGATTCAAAGCCGTAAAAGCATTACGTTTAGAAGATATGCGTATTCCTTATGCATACTTAAAAACATTCCAAGGTCCAGCTACAGGAGTTGTTGTAGAACGTGAAAGAATGGATGTATTCGGACGTCCTTTATTAGGAGCAACTGTTAAACCAAAATTAGGTCTTTCTGGTAAAAACTACGGTCGTGTAGTTTACGAAGGATTAAAAGGTGGATTAGACTTCTTAAAAGATGACGAAAACATTAACTCTCAACCATTCATGAGATGGCGTGAAAG

>gi|2778273106|gb|PQ064527.1| Mallomonas mangofera strain R079 ribulose-1,5-bisphosphate carboxylase/oxygenase large subunit (rbcL) gene, partial cds; chloroplast

TAACTTAACGGCATCTATTATTGGTAACGTTTTTGGATTCAAAGCTGTAAAAGCTTTACGTTTAGAAGATATGCGTATTCCTTATGCATACTTAAAAACTTTCCAAGGTCCAGCAACAGGAGTTATTGTAGAACGTGAAAGAATGGATGTTTTTGGACGTCCTTTATTAGGAGCAACTGTTAAACCTAAATTAGGTCTTTCTGGTAAAAACTATGGTCGTGTAGTTTATGAAGGATTAAAAGGTGGATTAGACTTCTTAAAAGATGATGAAAATATTAATTCACAACCATTCATGAGATGGCGTGAAAG

>gi|2778273108|gb|PQ064528.1| Mallomonas mangofera strain R092 ribulose-1,5-bisphosphate carboxylase/oxygenase large subunit (rbcL) gene, partial cds; chloroplast

TAACTTAACGGCATCTATTATTGGTAACGTTTTTGGATTCAAAGCTGTAAAAGCTTTACGTTTAGAAGATATGCGTATTCCTTATGCATACTTAAAAACTTTCCAAGGTCCAGCAACAGGAGTTATTGTAGAACGTGAAAGAATGGATGTTTTTGGACGTCCTTTATTAGGAGCAACTGTTAAACCTAAATTAGGTCTTTCTGGTAAAAACTATGGTCGTGTAGTTTATGAAGGATTAAAAGGTGGATTAGACTTCTTAAAAGATGATGAAAATATTAATTCACAACCATTCATGAGATGGCGTGAAAG

>gi|2780169390|gb|PQ031109.1| Mallomonas actinoloma var. maramuresensis strain CZ104D ribulose-1,5-bisphosphate carboxylase/oxygenase large subunit (rbcL) gene, complete cds; chloroplast

TAACTTAACAGCATCTATCATCGGGAACGTTTTCGGATTCAAAGCCGTTAAAGCTTTACGTTTAGAAGATATGCGTATTCCTTATGCTTACTTAAAAACTTTCCAAGGTCCAGCGACTGGAGTTATTGTAGAACGTGAAAGAATGGATGTATTTGGTCGCCCTCTATTAGGTGCTACAGTTAAACCAAAATTAGGTCTTTCTGGTAAAAACTATGGACGTGTTGTTTATGAAGGATTAAAAGGTGGTTTAGACTTCTTAAAAGATGACGAAAATATTAACTCACAACCATTCATGAGATGGCGTGAAAG

>gi|2780169392|gb|PQ031110.1| Mallomonas actinoloma var. maramuresensis strain CZ105B ribulose-1,5-bisphosphate carboxylase/oxygenase large subunit (rbcL) gene, complete cds; chloroplast

TAACTTAACAGCATCTATCATCGGGAACGTTTTCGGATTCAAAGCCGTTAAAGCTTTACGTTTAGAAGATATGCGTATTCCTTTTGCTTACTTAAAAACTTTCCAAGGTCCAGCGACTGGAGTTATTGTAGAACGTGAAAGAATGGATGTATTTGGTCGTCCTCTATTAGGTGCTACAGTTAAACCAAAATTAGGTCTTTCTGGTAAAAACTATGGACGTGTTGTTTATGAAGGATTAAAAGGTGGTTTAGACTTCTTAAAAGATGACGAAAATATTAACTCACAACCATTCATGAGATGGCGTGAAAG

>gi|2780169394|gb|PQ031111.1| Mallomonas aerolata strain CZ35M ribulose-1,5-bisphosphate carboxylase/oxygenase large subunit (rbcL) gene, complete cds; chloroplast

TAATTTAACAGCATCTATCATCGGAAATGTTTTTGGATTTAAAGCTGTTAAAGCTTTACGTTTAGAAGATATGCGTATTCCTTATGCTTACTTAAAAACTTTCCAAGGTCCTGCAACTGGAGTTATTGTAGAACGTGAAAGAATGGATGTATTCGGTCGTCCTTTATTAGGTGCTACAGTTAAACCAAAATTAGGTCTTTCTGGTAAAAACTATGGACGTGTTGTTTATGAAGGATTAAAAGGTGGTTTAGACTTCTTAAAAGATGACGAAAATATTAACTCACAACCATTCATGAGATGGCGTGAAAG

>gi|2780169396|gb|PQ031112.1| Mallomonas aerolata strain CZ35O ribulose-1,5-bisphosphate carboxylase/oxygenase large subunit (rbcL) gene, complete cds; chloroplast

TAATTTAACAGCATCTATCATCGGAAATGTTTTTGGATTTAAAGCTGTTAAAGCTTTACGTTTAGAAGATATGCGTATTCCTTATGCTTACTTAAAAACTTTCCAAGGTCCTGCAACTGGAGTTATTGTAGAACGTGAAAGAATGGATGTATTCGGTCGTCCTTTATTAGGTGCTACAGTTAAACCAAAATTAGGTCTTTCTGGTAAAAACTATGGACGTGTTGTTTATGAAGGATTAAAAGGTGGTTTAGACTTCTTAAAAGATGACGAAAATATTAACTCACAACCATTCATGAGATGGCGTGAAAG

>gi|2780169398|gb|PQ031113.1| Mallomonas crassisquama strain CZ04A ribulose-1,5-bisphosphate carboxylase/oxygenase large subunit (rbcL) gene, complete cds; chloroplast

AAACTTAACAGCATCTATTATCGGTAACGTTTTTGGATTCAAAGCCGTTAAAGCTTTACGTTTAGAAGATATGCGTATTCCTTATGCTTACTTAAAAACTTTCCAAGGTCCTGCAACTGGAGTTGTTGTAGAACGTGAAAGAATGGATGTATTTGGTCGCCCTCTATTAGGTGCTACAGTAAAACCAAAATTAGGTCTTTCTGGTAAAAACTATGGACGTGTTGTTTATGAAGGATTAAAAGGTGGTTTAGACTTCTTAAAAGATGACGAAAATATTAACTCACAACCATTCATGAGATGGCGTGAAAG

>gi|2780169400|gb|PQ031114.1| Mallomonas crassisquama strain CZ07M ribulose-1,5-bisphosphate carboxylase/oxygenase large subunit (rbcL) gene, complete cds; chloroplast

AAACTTAACAGCATCTATTATCGGTAACGTTTTTGGATTCAAAGCCGTTAAAGCTTTACGTTTAGAAGATATGCGTATTCCTTATGCTTACTTAAAAACTTTCCAAGGTCCTGCAACTGGAGTTGTTGTAGAACGTGAAAGAATGGATGTATTTGGTCGCCCTCTATTAGGTGCTACAGTAAAACCAAAATTAGGTCTTTCTGGTAAAAACTATGGACGTGTTGTTTATGAAGGATTAAAAGGTGGTTTAGACTTCTTAAAAGATGACGAAAATATTAACTCACAACCATTCATGAGATGGCGTGAAAG

>gi|2780169402|gb|PQ031115.1| Mallomonas elongata strain CZ26E ribulose-1,5-bisphosphate carboxylase/oxygenase large subunit (rbcL) gene, complete cds; chloroplast

TAACTTAACAGCATCTATTATCGGTAACGTTTTCGGATTCAAAGCCGTTAAAGCTTTACGTTTAGAAGATATGCGTATTCCTTATGCTTACTTAAAAACTTTCCAAGGTCCTGCGACTGGAGTTATTGTAGAACGTGAAAGAATGGATGTTTTTGGTCGCCCTTTATTAGGTGCTACAGTTAAACCAAAATTAGGTCTTTCTGGTAAAAACTACGGGCGTGTTGTTTATGAAGGATTAAAAGGTGGTTTAGACTTCTTAAAAGATGATGAAAACATCAATTCACAACCATTCATGAGATGGCGTGAAAG

>gi|2780169404|gb|PQ031116.1| Mallomonas elongata strain CZ24B ribulose-1,5-bisphosphate carboxylase/oxygenase large subunit (rbcL) gene, complete cds; chloroplast

TAACTTAACAGCATCTATTATCGGTAACGTTTTCGGATTCAAAGCCGTTAAAGCTTTACGTTTAGAAGATATGCGTATTCCTTATGCTTACTTAAAAACTTTCCAAGGTCCTGCGACTGGAGTTATTGTAGAACGTGAAAGAATGGATGTTTTTGGTCGCCCTTTATTAGGTGCTACAGTTAAACCAAAATTAGGTCTTTCTGGTAAAAACTACGGGCGTGTTGTTTATGAAGGATTAAAAGGTGGTTTAGACTTCTTAAAAGATGATGAAAACATCAATTCACAACCATTCATGAGATGGCGTGAAAG

>gi|2780169406|gb|PQ031117.1| Mallomonas intermedia strain CZ34M ribulose-1,5-bisphosphate carboxylase/oxygenase large subunit (rbcL) gene, complete cds; chloroplast

AAACTTAACAGCATCTATCATTGGTAACGTTTTCGGATTCAAAGCTGTTAAAGCTTTACGTTTAGAAGATATGCGTATTCCTTATGCTTACTTAAAAACTTTCCAAGGTCCTGCGACTGGAGTTGTTGTAGAACGTGAAAGAATGGACTGTTTTGGTCGTCCTCTATTAGGTGCTACAGTTAAACCAAAATTAGGTCTTTCTGGTAAAAACTATGGACGTGTTGTTTATGAAGGATTAAAAGGTGGTTTAGACTTTTTAAAAGATGATGAAAATATTAACTCACAACCATTCATGAGATGGCGTGAAAG

>gi|2780169408|gb|PQ031118.1| Mallomonas intermedia strain CZ35C ribulose-1,5-bisphosphate carboxylase/oxygenase large subunit (rbcL) gene, complete cds; chloroplast

AAACTTAACAGCATCTATCATTGGTAACGTTTTCGGATTCAAAGCTGTTAAAGCTTTACGTTTAGAAGATATGCGTATTCCTTATGCTTACTTAAAAACTTTCCAAGGTCCTGCGACTGGAGTTGTTGTAGAACGTGAAAGAATGGACTGTTTTGGTCGTCCTCTATTAGGTGCTACAGTTAAACCAAAATTAGGTCTTTCTGGTAAAAACTATGGACGTGTTGTTTATGAAGGATTAAAAGGTGGTTTAGACTTTTTAAAAGATGATGAAAATATTAACTCACAACCATTCATGAGATGGCGTGAAAG

>gi|2780169410|gb|PQ031119.1| Mallomonas lelymene strain CZ29D ribulose-1,5-bisphosphate carboxylase/oxygenase large subunit (rbcL) gene, complete cds; chloroplast

AAACTTAACAGCCTCTATTATTGGTAACGTTTTTGGATTTAAAGCTGTTAAAGCTTTACGTTTAGAAGATATGCGTATTCCTTATGCTTACTTAAAAACTTTCCAAGGTCCTGCAACTGGAGTTGTTGTAGAACGTGAAAGAATGGATGTATTTGGACGTCCTTTATTAGGTGCTACAGTTAAACCAAAATTAGGTCTTTCTGGTAAAAACTATGGACGTGTAGTTTATGAAGGATTAAAAGGTGGTTTAGACTTTTTAAAAGATGATGAAAATATTAACTCACAACCATTCATGAGATGGCGTGAAAG

>gi|2780169412|gb|PQ031120.1| Mallomonas lelymene strain IT02A ribulose-1,5-bisphosphate carboxylase/oxygenase large subunit (rbcL) gene, complete cds; chloroplast

AAACTTAACAGCCTCTATTATTGGTAACGTTTTTGGATTTAAAGCTGTTAAAGCTTTACGTTTAGAAGATATGCGTATTCCTTATGCTTACTTAAAAACTTTCCAAGGTCCTGCAACTGGAGTTGTTGTAGAACGTGAAAGAATGGATGTATTTGGACGTCCTTTATTAGGTGCTACAGTTAAACCAAAATTAGGTCTTTCTGGTAAAAACTATGGACGTGTAGTTTATGAAGGATTAAAAGGTGGTTTAGACTTTTTAAAAGATGATGAAAATATTAACTCACAACCATTCATGAGATGGCGTGAAAG

>gi|2780169414|gb|PQ031121.1| Mallomonas flora strain CZ35K ribulose-1,5-bisphosphate carboxylase/oxygenase large subunit (rbcL) gene, complete cds; chloroplast

TAACTTAACAGCTTCTATTATTGGAAACGTTTTTGGATTCAAAGCCGTTAAATGTTTACGTTTAGAAGATATGCGTATTCCTTATGCATACTTAAAAACTTTCCAAGGTCCAGCAACAGGAGTGGTTGTAGAACGTGAAAGAATGGATGTTTTTGGACGTCCAATGTTAGGTGCTACTGTTAAACCTAAATTAGGTCTTTCAGGAAAAAACTATGGTCGTGTAGTTTATGAAGGATTAAAAGGTGGTTTAGACTTCTTAAAAGATGATGAAAATATTAACTCACAACCATTCATGAGATGGCGTGAAAG

>gi|2780169416|gb|PQ031122.1| Mallomonas flora strain CZ53C ribulose-1,5-bisphosphate carboxylase/oxygenase large subunit (rbcL) gene, complete cds; chloroplast

TAACTTAACAGCTTCTATTATTGGAAACGTTTTTGGATTTAAAGCCGTTAAATGTTTACGTTTAGAAGATATGCGTATTCCTTATGCATACTTAAAAACTTTCCAAGGTCCAGCAACAGGAGTTGTTGTAGAACGTGAAAGAATGGATGTTTTTGGACGTCCAATGTTAGGTGCTACTGTTAAACCTAAATTAGGTCTTTCAGGAAAAAACTATGGTCGTGTAGTTTATGAAGGATTAAAAGGTGGTTTAGACTTCTTAAAAGATGATGAAAATATTAACTCACAACCATTCATGAGATGGCGTGAAAG

>gi|2780169418|gb|PQ031123.1| Mallomonas striata strain CZ33H ribulose-1,5-bisphosphate carboxylase/oxygenase large subunit (rbcL) gene, complete cds; chloroplast

AAATTTAACAGCTTCTATTATTGGGAACGTTTTCGGATTTAAAGCCGTTAAAGCTTTACGTTTAGAAGATATGCGTATTCCGTATGCTTATTTAAAAACTTTCCAAGGTCCTGCAACAGGAGTTGTTGTAGAACGTGAAAGAATGGATGTTTTTGGACGTCCTTTATTAGGTGCTACTGTTAAACCAAAATTAGGTCTTTCTGGAAAAAACTATGGTCGTGTAGTTTATGAAGGATTAAAAGGTGGTTTAGATTTCTTAAAAGATGATGAAAATATTAACTCACAACCATTCATGAGATGGCGTGAAAG

>gi|2780169420|gb|PQ031124.1| Mallomonas kalinae strain CCMP 479 ribulose-1,5-bisphosphate carboxylase/oxygenase large subunit (rbcL) gene, complete cds; chloroplast

TAACTTAACAGCATCAATTATCGGAAACGTTTTTGGTTTCAAAGCTGTTAAAGCATTACGTTTAGAAGATATGCGTATTCCTTTTGCTTACCTAAAAACTTTCCAAGGTCCTGCAACAGGAGTTGTTGTAGAACGTGAAAGAATGGATGTTTTTGGACGTCCTTTATTAGGTGCTACTGTTAAACCAAAATTAGGTCTTTCTGGTAAAAACTATGGTCGTGTAGTTTATGAAGGATTAAAAGGTGGTTTAGATTTCTTAAAAGATGATGAAAATATTAATTCACAACCATTCATGAGATGGCGTGAAAG

>gi|2780169422|gb|PQ031125.1| Mallomonas kalinae strain CZ41B ribulose-1,5-bisphosphate carboxylase/oxygenase large subunit (rbcL) gene, complete cds; chloroplast

TAACTTAACAGCATCAATTATCGGAAACGTTTTTGGTTTCAAAGCTGTTAAAGCATTACGTTTAGAAGATATGCGTATTCCTTTTGCTTACCTAAAAACTTTCCAAGGTCCTGCAACAGGAGTTGTTGTAGAACGTGAAAGAATGGATGTTTTTGGACGTCCTTTATTAGGTGCTACTGTTAAACCAAAATTAGGTCTTTCTGGTAAAAACTATGGTCGTGTAGTTTATGAAGGATTAAAAGGTGGTTTAGATTTCTTAAAAGATGATGAAAATATTAATTCACAACCATTCATGAGATGGCGTGAAAG

>gi|2780169424|gb|PQ031126.1| Mallomonas papillosa strain CZ03F ribulose-1,5-bisphosphate carboxylase/oxygenase large subunit (rbcL) gene, complete cds; chloroplast

AAACTTAACTGCCTCTATCATTGGAAACGTTTTCGGATTTAAAGCTGTTAAAGCTTTACGTTTAGAAGATATGCGTATTCCCTATTCTTACTTAAAAACATTCCAAGGTCCTGCAACTGGAGTTATTGTAGAACGTGAAAGAATGGATGTATTTGGACGTCCTTTATTAGGAGCTACAGTTAAACCAAAATTAGGTCTTTCAGGAAAAAATTATGGTCGTGTAGTTTATGAAGGATTAAAAGGTGGTTTAGATTTCTTAAAAGATGATGAAAATATTAACTCACAACCATTCATGAGATGGCGTGAAAG

>gi|2780169426|gb|PQ031127.1| Mallomonas papillosa strain Jeongsan121209J ribulose-1,5-bisphosphate carboxylase/oxygenase large subunit (rbcL) gene, complete cds; chloroplast

AAACTTAACTGCCTCTATCATTGGAAACGTTTTCGGATTTAAAGCTGTTAAAGCTTTACGTTTAGAAGATATGCGTATTCCTTATTCTTACTTAAAAACATTCCAAGGTCCTGCAACTGGAGTTATTGTAGAACGTGAAAGAATGGATGTTTTTGGACGTCCTTTATTAGGAGCAACAGTTAAACCAAAATTAGGTCTTTCAGGAAAAAATTACGGTCGTGTAGTTTATGAAGGATTAAAAGGTGGTTTAGATTTCTTAAAAGATGATGAAAATATTAACTCACAACCATTCATGAGATGGCGTGAAAG

>gi|2780169428|gb|PQ031128.1| Mallomonas rasilis strain CZ38X ribulose-1,5-bisphosphate carboxylase/oxygenase large subunit (rbcL) gene, partial cds; chloroplast

TAACTTAACAGCTTCTATCATCGGAAACGTTTTTGGTTTCAAAGCCGTTAAAGCATTACGTTTAGAAGATATGCGTATTCCATATTCTTATCTAAAAACTTTCCAAGGTCCTGCAACAGGAGTTATTGTAGAACGTGAAAGAATGGATGTTTTTGGACGTCCTCTATTAGGAGCTACTGTTAAACCTAAATTAGGTCTTTCTGGTAAAAACTATGGTCGTGTAGTTTATGAAGGATTAAAAGGTGGTTTAGATTTCTTAAAAGACGATGAAAATATCAATTCACAACCATTCATGAGATGGCGTGAAAG

>gi|2780169430|gb|PQ031129.1| Mallomonas rasilis strain D01D ribulose-1,5-bisphosphate carboxylase/oxygenase large subunit (rbcL) gene, complete cds; chloroplast

TAACTTAACAGCTTCTATTATCGGAAACGTTTTTGGTTTCAAAGCCGTTAAAGCATTACGTTTAGAAGATATGCGTATTCCATATTCTTATCTAAAAACTTTCCAAGGTCCTGCAACAGGAGTTGTTGTAGAACGTGAAAGAATGGATGTTTTTGGACGTCCTCTATTAGGAGCTACTGTTAAACCTAAATTAGGTCTTTCTGGTAAAAACTATGGTCGTGTAGTTTATGAAGGATTAAAAGGTGGTTTAGATTTCTTAAAAGATGATGAAAATATCAATTCACAACCATTCATGAGATGGCGTGAAAG

>gi|2780169432|gb|PQ031130.1| Mallomonas annulata strain CZ34K ribulose-1,5-bisphosphate carboxylase/oxygenase large subunit (rbcL) gene, complete cds; chloroplast

TAACTTAACAGCATCTATCATCGGAAACGTTTTCGGTTTCAAAGCCGTTAAAGCTTTACGTTTAGAAGATATGCGTATTCCTTATGCATACTTAAAAACTTTCCAAGGTCCAGCTACTGGAGTTATTGTTGAACGTGAAAGAATGGATGTATTTGGACGTCCTTTATTAGGAGCTACTGTTAAACCTAAATTAGGTCTTTCTGGTAAAAACTATGGTCGTGTAGTTTATGAAGGATTAAAAGGTGGTTTAGATTTCTTAAAAGATGATGAAAATATTAACTCACAACCATTCATGAGATGGCGTGAAAG

>gi|2780169434|gb|PQ031131.1| Mallomonas annulata strain CZ23C ribulose-1,5-bisphosphate carboxylase/oxygenase large subunit (rbcL) gene, complete cds; chloroplast

TAACTTAACAGCATCTATCATCGGAAACGTTTTCGGTTTCAAAGCCGTTAAAGCTTTACGTTTAGAAGATATGCGTATTCCTTATGCATACTTAAAAACTTTCCAAGGTCCAGCTACTGGAGTTATTGTTGAACGTGAAAGAATGGATGTATTTGGACGTCCTTTATTAGGAGCTACTGTTAAACCTAAATTAGGTCTTTCTGGTAAAAACTATGGTCGTGTAGTTTATGAAGGATTAAAAGGTGGTTTAGATTTCTTAAAAGATGATGAAAATATTAACTCACAACCATTCATGAGATGGCGTGAAAG

>gi|2780169436|gb|PQ031132.1| Mallomonas doignonii strain CZ08K ribulose-1,5-bisphosphate carboxylase/oxygenase large subunit (rbcL) gene, complete cds; chloroplast

TAACTTAACAGCCTCTATTATTGGTAACGTTTTTGGTTTCAAAGCAGTAAAATGTTTACGTTTAGAAGATATGCGTATTCCTTATGCTTACTTAAAAACATTCCAAGGTCCTGCATGTGGAGTTATTGTTGAACGTGAAAGAATGGATTGTTTTGGACGTCCTTTATTAGGAGCAACTGTTAAACCAAAATTAGGTCTTTCTGGTAAAAACTATGGTCGTGTAGTTTATGAAGGATTAAAAGGTGGATTAGACTTCTTAAAAGATGATGAAAACATCAACTCACAACCATTCATGAGATGGCGTGAAAG

>gi|2780169438|gb|PQ031133.1| Mallomonas doignonii strain CZ08N ribulose-1,5-bisphosphate carboxylase/oxygenase large subunit (rbcL) gene, complete cds; chloroplast

TAACTTAACAGCCTCTATTATTGGTAACGTTTTTGGTTTCAAAGCAGTAAAATGTTTACGTTTAGAAGATATGCGTATTCCTTATGCTTACTTAAAAACATTCCAAGGTCCTGCATGTGGAGTTATTGTTGAACGTGAAAGAATGGATTGTTTTGGACGTCCTTTATTAGGAGCAACTGTTAAACCAAAATTAGGTCTTTCTGGTAAAAACTATGGTCGTGTAGTTTATGAAGGATTAAAAGGTGGATTAGACTTCTTAAAAGATGATGAAAACATCAACTCACAACCATTCATGAGATGGCGTGAAAG

>gi|2780169440|gb|PQ031134.1| Mallomonas eoa strain CZ115B ribulose-1,5-bisphosphate carboxylase/oxygenase large subunit (rbcL) gene, complete cds; chloroplast

TAACTTAACAGCATCTATTATTGGGAACGTTTTTGGTTTCAAAGCTGTAAAATGTTTACGTTTAGAAGATATGCGTATTCCTTATGCATACTTAAAAACATTCCAAGGTCCTGCGACTGGAGTAGTTGTTGAACGTGAAAGAATGGATTGTTTTGGACGTCCTTTATTAGGAGCAACTGTAAAACCAAAATTAGGTCTTTCTGGTAAAAACTATGGTCGTGTAGTTTATGAAGGATTAAAAGGTGGATTAGACTTCTTAAAAGATGATGAAAATATCAATTCTCAACCATTCATGAGATGGCGTGAAAG

>gi|2780169442|gb|PQ031135.1| Mallomonas eoa strain CZ115D ribulose-1,5-bisphosphate carboxylase/oxygenase large subunit (rbcL) gene, complete cds; chloroplast

TAACTTAACAGCATCTATTATTGGGAACGTTTTTGGTTTCAAAGCTGTAAAATGTTTACGTTTAGAAGATATGCGTATTCCTTATGCATACTTAAAAACATTCCAAGGTCCTGCGACTGGAGTAGTTGTTGAACGTGAAAGAATGGATTGTTTTGGACGTCCTTTATTAGGAGCAACTGTAAAACCAAAATTAGGTCTTTCTGGTAAAAACTATGGTCGTGTAGTTTATGAAGGATTAAAAGGTGGATTAGACTTCTTAAAAGATGATGAAAATATCAATTCTCAACCATTCATGAGATGGCGTGAAAG

>gi|2780169444|gb|PQ031136.1| Mallomonas favosa strain D02A ribulose-1,5-bisphosphate carboxylase/oxygenase large subunit (rbcL) gene, partial cds; chloroplast

TAACTTAACAGCATCTATCATTGGTAACGTTTTTGGATTTAAAGCTGTAAAAGCTTTACGTTTAGAAGATATGCGTATTCCTTATGCATACTTAAAAACTTTCCAAGGTCCAGCTACAGGAGTAATTGTAGAACGTGAAAGAATGGATGTATTTGGACGTCCTTTATTAGGAGCAACAGTTAAACCAAAATTAGGTCTTTCTGGAAAAAACTATGGTCGTGTAGTTTATGAAGGATTAAAAGGTGGATTAGACTTCTTAAAAGATGACGAAAACATTAACTCTCAACCATTCATGAGATGGCGTGAAAG

>gi|2780169446|gb|PQ031137.1| Mallomonas favosa strain D03C ribulose-1,5-bisphosphate carboxylase/oxygenase large subunit (rbcL) gene, complete cds; chloroplast

TAACTTAACAGCATCTATCATTGGTAACGTTTTTGGATTTAAAGCTGTAAAAGCTTTACGTTTAGAAGATATGCGTATTCCTTATGCATACTTAAAAACTTTCCAAGGTCCAGCTACAGGAGTAATTGTAGAACGTGAAAGAATGGATGTATTTGGACGTCCTTTATTAGGAGCAACAGTTAAACCAAAATTAGGTCTTTCTGGAAAAAACTATGGTCGTGTAGTTTATGAAGGATTAAAAGGTGGATTAGACTTCTTAAAAGATGACGAAAACATTAACTCTCAACCATTCATGAGATGGCGTGAAAG

>gi|2780169448|gb|PQ031138.1| Mallomonas munda strain CZ38H ribulose-1,5-bisphosphate carboxylase/oxygenase large subunit (rbcL) gene, complete cds; chloroplast

TAACTTAACAGCATCTATTATTGGGAACGTTTTTGGTTTCAAAGCTGTAAAATGTTTACGTTTAGAAGATATGCGTATTCCTTATGCTTATTTAAAAACATTCCAAGGTCCTGCAACTGGAGTAGTTGTTGAACGTGAAAGAATGGATTGTTTTGGACGTCCTTTATTAGGAGCAACTGTAAAACCAAAATTAGGTCTTTCTGGTAAAAACTATGGTCGTGTAGTTTATGAAGGATTAAAAGGTGGATTAGACTTCTTAAAAGATGATGAAAATATCAATTCTCAACCATTCATGAGATGGCGTGAAAG

>gi|2780169450|gb|PQ031139.1| Mallomonas munda strain CZ85G ribulose-1,5-bisphosphate carboxylase/oxygenase large subunit (rbcL) gene, complete cds; chloroplast

TAACTTAACAGCATCTATTATTGGGAACGTTTTTGGTTTCAAAGCTGTAAAATGTTTACGTTTAGAAGATATGCGTATTCCTTATGCTTATTTAAAAACATTCCAAGGTCCTGCAACTGGAGTAGTTGTTGAACGTGAAAGAATGGATTGTTTTGGACGTCCTTTATTAGGAGCAACTGTAAAACCAAAATTAGGTCTTTCTGGTAAAAACTATGGTCGTGTAGTTTATGAAGGATTAAAAGGTGGATTAGACTTCTTAAAAGATGATGAAAATATCAATTCTCAACCATTCATGAGATGGCGTGAAAG

>gi|2780169452|gb|PQ031140.1| Mallomonas schwemmlei strain CZ33F ribulose-1,5-bisphosphate carboxylase/oxygenase large subunit (rbcL) gene, complete cds; chloroplast

TAACTTAACAGCATCTATCATCGGTAACGTATTTGGTTTCAAAGCTGTAAAAGCATTACGTTTAGAAGATATGCGTATTCCTTATGCCTACTTAAAAACTTTCCAAGGTCCTGCGACTGGAGTTGTTGTAGAACGTGAAAGAATGGATGTATTCGGACGTCCTTTATTAGGAGCAACTGTTAAACCTAAATTAGGTCTTTCTGGTAAAAACTATGGTCGTGTAGTTTATGAAGGATTAAAAGGTGGATTAGACTTCTTAAAAGATGACGAAAACATTAACTCTCAACCATTCATGAGATGGCGTGAAAG

>gi|2780169454|gb|PQ031141.1| Mallomonas schwemmlei strain CZ33J ribulose-1,5-bisphosphate carboxylase/oxygenase large subunit (rbcL) gene, complete cds; chloroplast

TAACTTAACAGCATCTATCATCGGTAACGTATTTGGTTTCAAAGCTGTAAAAGCATTACGTTTAGAAGATATGCGTATTCCTTATGCCTACTTAAAAACTTTCCAAGGTCCTGCGACTGGAGTTGTTGTAGAACGTGAAAGAATGGATGTATTCGGACGTCCTTTATTAGGAGCAACTGTTAAACCTAAATTAGGTCTTTCTGGTAAAAACTATGGTCGTGTAGTTTATGAAGGATTAAAAGGTGGATTAGACTTCTTAAAAGATGACGAAAACATTAACTCTCAACCATTCATGAGATGGCGTGAAAG

>gi|2780169456|gb|PQ031142.1| Mallomonas adamas strain CZ38K ribulose-1,5-bisphosphate carboxylase/oxygenase large subunit (rbcL) gene, complete cds; chloroplast

TAACTTAACTGCATCGATTATTGGAAACGTTTTCGGTTTCAAAGCTGTAAAAGCATTACGTTTAGAAGATATGCGTATTCCTTATGCATACTTAAAAACTTTCTTAGGTCCTGCAACAGGAGTTATTGTAGAACGTGAAAGAATGGATTGTTTTGGACGTCCTCTATTAGGAGCAACAGTTAAACCTAAATTAGGTCTTTCAGGAAAAAACTATGGTCGTGTAGTTTATGAAGGATTAAAAGGTGGTCTAGATTTCTTAAAAGATGATGAAAATATTAATTCTCAACCATTCATGAGATGGCGTGAAAG

>gi|2780169458|gb|PQ031143.1| Mallomonas adamas strain CZ38P ribulose-1,5-bisphosphate carboxylase/oxygenase large subunit (rbcL) gene, complete cds; chloroplast

TAACTTAACTGCATCGATTATTGGAAACGTTTTCGGTTTCAAAGCTGTAAAAGCATTACGTTTAGAAGATATGCGTATTCCTTATGCATACTTAAAAACTTTCTTAGGTCCTGCAACAGGAGTTATTGTAGAACGTGAAAGAATGGATTGTTTTGGACGTCCTCTATTAGGAGCAACAGTTAAACCTAAATTAGGTCTTTCAGGAAAAAACTATGGTCGTGTAGTTTATGAAGGATTAAAAGGTGGTCTAGATTTCTTAAAAGATGATGAAAATATTAATTCTCAACCATTCATGAGATGGCGTGAAAG

>gi|2780169460|gb|PQ031144.1| Mallomonas splendens strain AU07B ribulose-1,5-bisphosphate carboxylase/oxygenase large subunit (rbcL) gene, complete cds; chloroplast

AAACTTAACTGCTTCAATTATTGGAAACGTTTTTGGTTTCAAAGCTGTAAAAGCATTACGTTTAGAAGATATGCGTATTCCTTATGCATATTTAAAAACTTTCTTAGGTCCTGCAACTGGAGTAGTTGTAGAACGTGAAAGAATGGATTGTTTTGGACGTCCTCTATTAGGAGCAACAGTTAAACCTAAATTAGGTCTTTCTGGAAAAAACTATGGTCGTGTAGTTTATGAAGGATTAAGAGGTGGATTAGATTTCTTAAAAGATGACGAAAATATTAACTCTCAACCATTCATGCGTTGGCGTGAAAG

>gi|2780169462|gb|PQ031145.1| Mallomonas splendens strain CCMP 1782 ribulose-1,5-bisphosphate carboxylase/oxygenase large subunit (rbcL) gene, complete cds; chloroplast

AAACTTAACTGCTTCAATTATTGGAAACGTTTTTGGTTTCAAAGCTGTAAAAGCATTACGTTTAGAAGATATGCGTATTCCTTATGCATATTTAAAAACTTTCTTAGGTCCTGCAACTGGAGTAATTGTAGAACGTGAAAGAATGGATTGTTTTGGACGTCCTTTATTAGGAGCAACAGTTAAACCTAAATTAGGTCTTTCTGGAAAAAACTATGGTCGTGTAGTTTATGAAGGATTAAGAGGTGGATTAGATTTCTTAAAAGATGACGAAAATATTAACTCTCAACCATTCATGCGTTGGCGTGAAAG

>gi|2780169464|gb|PQ031146.1| Mallomonas caudata strain CZ25J ribulose-1,5-bisphosphate carboxylase/oxygenase large subunit (rbcL) gene, complete cds; chloroplast

AAACTTGACAGCTTCAATTATCGGAAATGTATTTGGTTTTAAAGCAGTAAAAGCTTTACGTTTAGAAGATATGCGTATTCCTTATGCATATTTAAAAACATTCTTAGGCCCTGCAACAGGAGTAATCGTTGAACGTGAAAGAATGGATTGTTTTGGACGTCCTTTATTAGGAGCGACTGTTAAACCAAAATTAGGTCTTTCAGGAAAAAACTATGGTCGTGTAGTTTATGAAGGATTAAAAGGTGGATTAGATTTCTTAAAAGATGACGAAAATATCAACTCTCAAGCTTTTATGAGATGGCGTGAAAG

>gi|2780169466|gb|PQ031147.1| Mallomonas caudata strain CZ92A ribulose-1,5-bisphosphate carboxylase/oxygenase large subunit (rbcL) gene, complete cds; chloroplast

AAACTTGACAGCTTCAATTATCGGAAATGTATTTGGTTTTAAAGCAGTAAAAGCTTTACGTTTAGAAGATATGCGTATTCCTTATGCATATTTAAAAACATTCTTAGGCCCTGCAACAGGAGTAATCGTTGAACGTGAAAGAATGGATTGTTTTGGACGTCCTTTATTAGGAGCGACTGTTAAACCAAAATTAGGTCTTTCAGGAAAAAACTATGGTCGTGTAGTTTATGAAGGATTAAAAGGTGGATTAGATTTCTTAAAAGATGACGAAAATATCAACTCTCAAGCTTTTATGAGATGGCGTGAAAG

>gi|2811253644|gb|PQ300136.1| Mallomonas favosa f. gemina strain 2_18 ribulose-1,5-bisphosphate carboxylase/oxygenase large subunit (rbcL) gene, partial cds; plastid

TAACTTAACAGCATCAATTATTGGTAACGTTTTTGGATTCAAAGCTGTAAAAGCTTTACGTTTAGAAGATATGCGTATCCCTTATGCATACTTAAAAACATTCCAAGGTCCAGCTACAGGAGTTATTGTAGAACGTGAAAGAATGGATGTATTTGGACGTCCTTTATTAGGAGCTACTGTTAAACCTAAATTAGGTCTTTCTGGAAAAAACTATGGTCGTGTAGTTTATGAAGGATTAAAAGGTGGATTAGACTTCTTAAAAGATGACGAAAATATCAATTCTCAACCATTCATGAGATGGCGTGAAGG

>gi|2811253646|gb|PQ300137.1| Mallomonas favosa f. gemina strain 12_18 ribulose-1,5-bisphosphate carboxylase/oxygenase large subunit (rbcL) gene, partial cds; plastid

TAACTTAACAGCATCTATTATTGGTAACGTTTTTGGATTCAAAGCTGTAAAAGCTTTACGTTTAGAAGATATGCGTATCCCTTATGCATACTTAAAAACATTCCAAGGTCCAGCTACAGGAGTTATTGTAGAACGTGAAAGAATGGATGTATTTGGACGTCCTTTATTAGGAGCTACTGTTAAACCTAAATTAGGTCTTTCTGGAAAAAACTATGGTCGTGTAGTTTATGAAGGATTAAAAGGTGGATTAGACTTCTTAAAAGATGACGAAAATATCAATTCTCAACCATTCATGAGATGGCGTGAAAG

>gi|2811253648|gb|PQ300138.1| Mallomonas favosa strain 629Yu ribulose-1,5-bisphosphate carboxylase/oxygenase large subunit (rbcL) gene, partial cds; plastid

TAACTTAACAGCATCTATCATCGGTAACGTTTTTGGATTCAAAGCTGTAAAAGCTTTACGTTTAGAAGATATGCGTATTCCTTATGCATACTTAAAAACTTTCCAAGGTCCAGCTACAGGGGTAATTGTAGAACGTGAAAGAATGGATGTATTTGGACGTCCTTTATTAGGAGCAACAGTTAAACCAAAATTAGGTCTTTCAGGAAAAAACTATGGTCGTGTAGTTTATGAAGGATTAAAAGGTGGATTAGACTTCTTAAAAGATGACGAAAACATTAACTCTCAACCATTCATGAGATGGCGTGAAAG

>gi|703265141|emb|HG514234.1| Synura borealis partial rbcL gene for ribulose-1,5-bisphosphate carboxylase/oxygenase large subunit gene, strain S 58.C7

AAACTTAACAGCATCAATTATTGGAAACGTTTTCGGTTTTAAAGCTGTAAAATGTTTACGTTTAGAAGATATGCGTATTCCGTATGCTTATTTAAAAACGTTTATTGGACCTGCAACTGGAGTTATTGTAGAACGTGAAAGAATGGATGTATTTGGACGTCCTCTTTTAGGTGCAACTGTAAAACCAAAATTAGGTCTTTCTGGTAAAGCTTATGGTCGTGTAGTTTATGAAGGATTAAAAGGTGGTTTAGATTTCTTAAAAGACGATGAAAATATTAATTCACAACCATTCATGAGATGGCGTGAAAG

>gi|703265145|emb|HG514235.1| Synura borealis partial rbcL gene for ribulose-1,5-bisphosphate carboxylase/oxygenase large subunit gene, strain S 62.D7

AAACTTAACAGCATCAATTATTGGAAACGTTTTCGGTTTTAAAGCTGTAAAATGTTTACGTTTAGAAGATATGCGTATTCCGTATGCTTATTTAAAAACGTTTATTGGACCTGCAACTGGAGTTATTGTAGAACGTGAAAGAATGGATGTATTTGGACGTCCTCTTTTAGGTGCAACTGTAAAACCAAAATTAGGTCTTTCTGGTAAAGCTTATGGTCGTGTAGTTTATGAAGGATTAAAAGGTGGTTTAGATTTCTTAAAAGACGATGAAAATATTAATTCACAACCATTCATGAGATGGCGTGAAAG

>gi|703265150|emb|HG514236.1| Synura borealis partial rbcL gene for ribulose-1,5-bisphosphate carboxylase/oxygenase large subunit gene, strain S 90.M34

AAACTTAACAGCATCAATTATTGGAAACGTTTTCGGTTTTAAAGCTGTAAAATGTTTACGTTTAGAAGATATGCGTATTCCGTATGCTTATTTAAAAACGTTTATTGGACCTGCAACTGGAGTTATTGTAGAACGTGAAAGAATGGATGTATTTGGACGTCCTCTTTTAGGTGCAACTGTAAAACCAAAATTAGGTCTTTCTGGTAAAGCTTATGGTCGTGTAGTTTATGAAGGATTAAAAGGTGGTTTAGATTTCTTAAAAGACGATGAAAATATTAATTCACAACCATTCATGAGATGGCGTGAAAG

>gi|703265155|emb|HG514237.1| Synura borealis partial rbcL gene for ribulose-1,5-bisphosphate carboxylase/oxygenase large subunit gene, strain S 114.B8

AAACTTAACAGCATCAATTATTGGAAACGTTTTCGGTTTTAAAGCTGTAAAATGTTTACGTTTAGAAGATATGCGTATTCCGTATGCTTATTTAAAAACGTTTATTGGACCTGCAACTGGAGTTATTGTAGAACGTGAAAGAATGGATGTATTTGGACGTCCTCTTTTAGGTGCAACTGTAAAACCAAAATTAGGTCTTTCTGGTAAAGCTTATGGTCGTGTAGTTTATGAAGGATTAAAAGGTGGTTTAGATTTCTTAAAAGACGATGAAAATATTAATTCACAACCATTCATGAGATGGCGTGAAAG

>gi|703265160|emb|HG514238.1| Synura borealis partial rbcL gene for ribulose-1,5-bisphosphate carboxylase/oxygenase large subunit gene, strain S 114.C8

AAACTTAACAGCATCAATTATTGGAAACGTTTTCGGTTTTAAAGCTGTAAAATGTTTACGTTTAGAAGATATGCGTATTCCGTATGCTTATTTAAAAACGTTTATTGGACCTGCAACTGGAGTTATTGTAGAACGTGAAAGAATGGATGTATTTGGACGTCCTCTTTTAGGTGCAACTGTAAAACCAAAATTAGGTCTTTCTGGTAAAGCTTATGGTCGTGTAGTTTATGAAGGATTAAAAGGTGGTTTAGATTTCTTAAAAGACGATGAAAATATTAATTCACAACCATTCATGAGATGGCGTGAAAG

>gi|703265164|emb|HG514239.1| Synura borealis partial rbcL gene for ribulose-1,5-bisphosphate carboxylase/oxygenase large subunit gene, strain S 117.D3

AAACTTAACAGCATCAATTATTGGAAACGTTTTCGGTTTTAAAGCTGTAAAATGTTTACGTTTAGAAGATATGCGTATTCCGTATGCTTATTTAAAAACGTTTATTGGACCTGCAACTGGAGTTATTGTAGAACGTGAAAGAATGGATGTATTTGGACGTCCTCTTTTAGGTGCAACTGTAAAACCAAAATTAGGTCTTTCTGGTAAAGCTTATGGTCGTGTAGTTTATGAAGGATTAAAAGGTGGTTTAGATTTCTTAAAAGACGATGAAAATATTAATTCACAACCATTCATGAGATGGCGTGAAAG

>gi|703265170|emb|HG514240.1| Synura borealis partial rbcL gene for ribulose-1,5-bisphosphate carboxylase/oxygenase large subunit gene, strain S 114.G6

AAACTTAACAGCATCAATTATTGGAAACGTTTTCGGTTTTAAAGCTGTAAAATGTTTACGTTTAGAAGATATGCGTATTCCGTATGCTTATTTAAAAACGTTTATCGGACCTGCAACTGGAGTTATTGTAGAACGTGAAAGAATGGATGTATTTGGACGTCCTCTTTTAGGTGCAACTGTAAAACCAAAATTAGGTCTTTCTGGTAAAGCTTATGGTCGTGTAGTTTATGAAGGATTAAAAGGTGGTTTAGATTTCTTAAAAGACGATGAAAATATTAATTCACAACCATTCATGAGATGGCGTGAAAG

>gi|703265175|emb|HG514241.1| Synura borealis partial rbcL gene for ribulose-1,5-bisphosphate carboxylase/oxygenase large subunit gene, strain S 115.F4

AAACTTAACAGCATCAATTATTGGAAACGTTTTCGGTTTTAAAGCTGTAAAATGTTTACGTTTAGAAGATATGCGTATTCCGTATGCTTATTTAAAAACGTTTATCGGACCTGCAACTGGAGTTATTGTAGAACGTGAAAGAATGGATGTATTTGGACGTCCTCTTTTAGGTGCAACTGTAAAACCAAAATTAGGTCTTTCTGGTAAAGCTTATGGTCGTGTAGTTTATGAAGGATTAAAAGGTGGTTTAGATTTCTTAAAAGACGATGAAAATATTAATTCACAACCATTCATGAGATGGCGTGAAAG

>gi|703265179|emb|HG514242.1| Synura borealis partial rbcL gene for ribulose-1,5-bisphosphate carboxylase/oxygenase large subunit gene, strain S 115.G7

AAACTTAACAGCATCAATTATTGGAAACGTTTTCGGTTTTAAAGCTGTAAAATGTTTACGTTTAGAAGATATGCGTATTCCGTATGCTTATTTAAAAACGTTTATTGGACCTGCAACTGGAGTTATTGTAGAACGTGAAAGAATGGATGTATTTGGACGTCCTCTTTTAGGTGCAACTGTAAAACCAAAATTAGGTCTTTCTGGTAAAGCTTATGGTCGTGTAGTTTATGAAGGATTAAAAGGTGGTTTAGATTTCTTAAAAGACGATGAAAATATTAATTCACAACCATTCATGAGATGGCGTGAAAG

>gi|703265184|emb|HG514243.1| Synura heteropora partial rbcL gene for ribulose-1,5-bisphosphate carboxylase/oxygenase large subunit gene, strain S 20.45

AAACTTAACAGCATCAATTATTGGAAACGTTTTTGGTTTTAAAGCTGTAAAATGTTTACGTTTAGAAGATATGCGTATCCCGTATGCTTATTTAAAAACGTTTATTGGACCTGCAACTGGAGTTATTGTAGAACGTGAAAGAATGGATGTATTTGGACGTCCTCTTTTAGGTGCAACTGTAAAACCAAAATTAGGTCTTTCTGGTAAAGCTTATGGTCGTGTAGTTTATGAAGGATTAAAAGGTGGTTTAGATTTCTTAAAAGACGATGAAAATATTAATTCACAACCATTCATGAGATGGCGTGAAAG

>gi|703265188|emb|HG514244.1| Synura heteropora partial rbcL gene for ribulose-1,5-bisphosphate carboxylase/oxygenase large subunit gene, strain S 87.C6

AAACTTAACAGCATCAATTATTGGAAACGTTTTTGGTTTTAAAGCTGTAAAATGTTTACGTTTAGAAGATATGCGTATCCCGTATGCTTATTTAAAAACGTTTATTGGACCTGCAACTGGAGTTATTGTAGAACGTGAAAGAATGGATGTATTTGGACGTCCTCTTTTAGGTGCAACTGTAAAACCAAAATTAGGTCTTTCTGGTAAAGCTTATGGTCGTGTAGTTTATGAAGGATTAAAAGGTGGTTTAGATTTCTTAAAAGACGATGAAAATATTAATTCACAACCATTCATGAGATGGCGTGAAAG

>gi|703265192|emb|HG514245.1| Synura heteropora partial rbcL gene for ribulose-1,5-bisphosphate carboxylase/oxygenase large subunit gene, strain S 101.F7

AAACTTAACAGCATCAATTATTGGAAACGTTTTTGGTTTTAAAGCTGTAAAATGTTTACGTTTAGAAGATATGCGTATCCCGTATGCTTATTTAAAAACGTTTATTGGACCTGCAACTGGAGTTATTGTAGAACGTGAAAGAATGGATGTATTTGGACGTCCTCTTTTAGGTGCAACTGTAAAACCAAAATTAGGTCTTTCTGGTAAAGCTTATGGTCGTGTAGTTTATGAAGGATTAAAAGGTGGTTTAGATTTCTTAAAAGACGATGAAAATATTAATTCACAACCATTCATGAGATGGCGTGAAAG

>gi|703265198|emb|HG514246.1| Synura hibernica partial rbcL gene for ribulose-1,5-bisphosphate carboxylase/oxygenase large subunit gene, strain S IE E4

AAACTTAACAGCATCAATTATTGGAAACGTTTTCGGTTTTAAAGCTGTAAAATGTTTACGTTTAGAAGATATGCGTATCCCGTATGCTTATTTAAAAACGTTTATTGGACCTGCAACTGGAGTTATTGTAGAACGTGAAAGAATGGATGTATTTGGACGTCCTCTTTTAGGTGCAACTGTAAAACCAAAATTAGGTCTTTCTGGTAAAGCTTATGGTCGTGTAGTTTATGAAGGATTAAAAGGTGGTTTAGATTTCTTAAAAGACGATGAAAATATTAATTCACAACCATTCATGAGATGGCGTGAAAG

>gi|703265202|emb|HG514247.1| Synura hibernica partial rbcL gene for ribulose-1,5-bisphosphate carboxylase/oxygenase large subunit gene, strain S IE E8

AAACTTAACAGCATCAATTATTGGAAACGTTTTCGGTTTTAAAGCTGTAAAATGTTTACGTTTAGAAGATATGCGTATCCCGTATGCTTATTTAAAAACGTTTATTGGACCTGCAACTGGAGTTATTGTAGAACGTGAAAGAATGGATGTATTTGGACGTCCTCTTTTAGGTGCAACTGTAAAACCAAAATTAGGTCTTTCTGGTAAAGCTTATGGTCGTGTAGTTTATGAAGGATTAAAAGGTGGTTTAGATTTCTTAAAAGACGATGAAAATATTAATTCACAACCATTCATGAGATGGCGTGAAAG

>gi|703265207|emb|HG514248.1| Synura hibernica partial rbcL gene for ribulose-1,5-bisphosphate carboxylase/oxygenase large subunit gene, strain S IE 104.D11

AAACTTAACAGCATCAATTATTGGAAACGTTTTCGGTTTTAAAGCTGTAAAATGTTTACGTTTAGAAGATATGCGTATCCCGTATGCTTATTTAAAAACGTTTATTGGACCTGCAACTGGAGTTATTGTAGAACGTGAAAGAATGGATGTATTTGGACGTCCTCTTTTAGGTGCAACTGTAAAACCAAAATTAGGTCTTTCTGGTAAAGCTTATGGTCGTGTAGTTTATGAAGGATTAAAAGGTGGTTTAGATTTCTTAAAAGACGATGAAAATATTAATTCACAACCATTCATGAGATGGCGTGAAAG

>gi|703265215|emb|HG514249.1| Synura hibernica partial rbcL gene for ribulose-1,5-bisphosphate carboxylase/oxygenase large subunit gene, strain S IE 105.F6

AAACTTAACAGCATCAATTATTGGAAACGTTTTCGGTTTTAAAGCTGTAAAATGTTTACGTTTAGAAGATATGCGTATCCCGTATGCTTATTTAAAAACGTTTATTGGACCTGCAACTGGAGTTATTGTAGAACGTGAAAGAATGGATGTATTTGGACGTCCTCTTTTAGGTGCAACTGTAAAACCAAAATTAGGTCTTTCTGGTAAAGCTTATGGTCGTGTAGTTTATGAAGGATTAAAAGGTGGTTTAGATTTCTTAAAAGACGATGAAAATATTAATTCACAACCATTCATGAGATGGCGTGAAAG

>gi|703265221|emb|HG514250.1| Synura laticarina partial rbcL gene for ribulose-1,5-bisphosphate carboxylase/oxygenase large subunit gene, strain S 90.C8

AAACTTAACAGCATCAATTATTGGAAATGTTTTCGGTTTTAAAGCTGTAAAATGTTTACGTTTAGAAGATATGCGTATTCCGTATGCTTATTTAAAAACGTTTATCGGACCTGCAACTGGAGTTATTGTAGAACGTGAAAGAATGGATGTATTTGGACGTCCTCTTTTAGGTGCAACTGTAAAACCAAAATTAGGTCTTTCTGGTAAAGCTTATGGTCGTGTAGTTTATGAAGGATTAAAAGGTGGTTTAGATTTCTTAAAAGACGATGAAAATATTAATTCACAACCATTCATGAGATGGCGTGAAAG

>gi|703265225|emb|HG514251.1| Synura laticarina partial rbcL gene for ribulose-1,5-bisphosphate carboxylase/oxygenase large subunit gene, strain S 115.D2

AAACTTAACAGCATCAATTATTGGAAACGTTTTCGGTTTTAAAGCTGTAAAATGTTTACGTTTAGAAGATATGCGTATTCCGTATGCTTATTTAAAAACGTTTATCGGACCTGCAACTGGAGTTATTGTAGAACGTGAAAGAATGGATGTATTTGGACGTCCTCTTTTAGGTGCAACTGTAAAACCAAAATTAGGTCTTTCTGGTAAAGCTTATGGTCGTGTAGTTTATGAAGGATTAAAAGGTGGTTTAGATTTCTTAAAAGACGATGAAAATATTAATTCACAACCATTCATGAGATGGCGTGAAAG

>gi|703265229|emb|HG514252.1| Synura macropora partial rbcL gene for ribulose-1,5-bisphosphate carboxylase/oxygenase large subunit gene, strain S 71.B4

AAACTTAACAGCATCAATTATTGGAAACGTTTTCGGTTTTAAAGCTGTAAAATGTTTACGTTTAGAAGATATGCGTATTCCTTATGCTTATTTAAAAACGTTTATCGGACCTGCAACTGGAGTTATTGTAGAACGTGAAAGAATGGATGTATTTGGACGCCCTCTTTTAGGTGCAACTGTAAAACCAAAATTAGGTCTTTCTGGTAAAGCTTATGGTCGTGTAGTTTATGAAGGATTAAAAGGTGGTTTAGATTTCTTAAAAGACGATGAAAATATTAATTCACAACCATTCATGAGATGGCGTGAAAG

>gi|703265235|emb|HG514253.1| Synura truttae partial rbcL gene for ribulose-1,5-bisphosphate carboxylase/oxygenase large subunit gene, strain CAUP B705

AAACTTAACAGCATCAATTATTGGAAACGTTTTTGGTTTTAAAGCTGTAAAATGTTTACGTTTAGAAGATATGCGTATCCCATATGCTTATTTAAAAACATTTATTGGACCTGCAACTGGAGTTATTGTAGAACGTGAAAGAATGGATGTATTTGGACGTCCTCTTTTAGGTGCAACTGTAAAACCAAAATTAGGTCTTTCTGGTAAAGCTTATGGTCGTGTAGTTTATGAAGGATTAAAAGGTGGTTTAGATTTCTTAAAAGACGATGAAAATATTAATTCACAACCATTCATGAGATGGCGTGAAAG

>gi|703265240|emb|HG514254.1| Synura truttae partial rbcL gene for ribulose-1,5-bisphosphate carboxylase/oxygenase large subunit gene, strain S 62.B5

AAACTTAACAGCATCAATTATTGGAAACGTTTTTGGTTTTAAAGCTGTAAAATGTTTACGTTTAGAAGATATGCGTATCCCATATGCTTATTTAAAAACATTTATTGGACCTGCAACTGGAGTTATTGTAGAACGTGAAAGAATGGATGTATTTGGACGTCCTCTTTTAGGTGCAACTGTAAAACCAAAATTAGGTCTTTCTGGTAAAGCTTATGGTCGTGTAGTTTATGAAGGATTAAAAGGTGGTTTAGATTTCTTAAAAGACGATGAAAATATTAATTCACAACCATTCATGAGATGGCGTGAAAG

>HQ710604.1 Rhizochromulina sp. CCMP237 culture CCMP:237 ribulose-1,5-bisphosphate carboxylase/oxygenase large subunit (rbcL) gene, partial cds; plastid

AAACCTTACAGCTTCAATTATTGGTAACGTATTTGGTTTCAAAGCTGTAAAAGCTCTTCGTCTAGAAGATATGCGTATTCCTTACGCGTACCTACGTACCTTCCAAGGTCCAGCAACAGGTGTTATCGTAGAGCGTGAGCGTCTAAACAACTTTGGTCGTCCAATTCTAGGTGCTACTGTAAAACCTAAGCTAGGTCTTTCAGGTCGTAACTATGGTCGTGTAGTTTATGAAGGTCTAAAAGGTGGTCTTGACTTCTTAAAAGATGATGAAAACATCAACTCTCAACCATTCATGCGTTGGCGTGAGCG

>NC_043890.1:c61315-59849 Rhizochromulina marina voucher A13,803 chloroplast, complete genome

-AACCTTACAGCTTCTATCATTGGTAACGTTTTCGGTTTCAAAGCAGTTAAAGCGCTTCGTTTAGAAGATATGCGTATTCCATTTGCATACTTAAAAACTTTCCAAGGTCCAGCAACAGGTGTTGTTGTAGAACGTGAGCGTTTAAATAACTTCGGTCGTCCAATTCTTGGAGCTACTGTAAAACCTAAGCTAGGTCTTTCTGGCCGTAACTATGGTCGTGTAGTATACGAAGGTCTAAAAGGTGGTCTAGACTTCCTAAAAGATGACGAGAACATTAACTCTCAACCATTCATGCGCTGGCGTGAGCG

>NC_044407.1:85178-86644 Florenciella parvula voucher CCMP2471 chloroplast, complete genome

-AACATTACTGCATCAATCATTGGTAACGTATTTGGTTTCAAGGCTGTTAAGGCTCTTCGTCTAGAAGATATGCGTATCCCATTTGCTTACTTAAAGACTTTCCAAGGTCCAGCTACAGGTGTTGTAGTAGAGCGTGAGCGTATGGATAAGTTTGGTCGTCCTCTTCTTGGTGCTACTGTTAAGCCTAAGCTAGGTCTTTCAGGTAAGAACTACGGTCGTGTAGTATATGAAGGTCTTAAGGGTGGTCTTGATTTCCTTAAAGATGATGAGAACATTAACTCTCAAGCATTCATGCGTTGGCGTGAGCG

>NC_044408.1:14534-16000 Pseudopedinella elastica voucher CCMP716 chloroplast, complete genome

-AACATTACTGCTTCTATTATCGGTAACGTATTTGGTTTCAAAGCTGTAAAAGCTCTTCGTTTAGAAGATATGCGTATCCCATATGCTTACCTTAAGACATTCCAAGGTCCAGCTACAGGTGTAGTTGTAGAGCGTGAGCGTATGAACTCATTCGGTCGTCCACTACTTGGTGCAACTGTTAAGCCTAAATTAGGTCTTTCTGGTCGTAACTACGGTCGTGTAGTATATGAAGGTCTTAAAGGTGGTCTAGACTTCCTTAAAGATGATGAGAATATTAACTCTCAACCATTCATGCGTTGGCGTGAGCG

>NC_043929.1:119020-120486 Dictyocha speculum voucher CCMP1381 chloroplast, complete genome

-AATATGACAGCGTCTATTATTGGTAACGTTTTCGGTTTCAAAGCGGTTAAAGCTCTTCGTCTAGAAGATATGCGTATCCCATTTGCTTACCTAAAAACATTCCAAGGACCTGCAACTGGTGTTGTTGTAGAGCGTGAGCGTATGGATAAATTTGGTCGTCCTTTATTAGGTGCGACAGTTAAGCCTAAATTAGGTCTTTCTGGTAAAAACTATGGTCGTGTAGTTTTCGAGGGTCTTAAAGGTGGTCTTGATTTCCTTAAAGATGATGAGAATATTAACTCACAAGCGTTCATGCGTTGGAGAGAACG

>NC_043929.1:121176-122735 Dictyocha speculum voucher CCMP1381 chloroplast, complete genome

-AACATGGCTGCATTAATTATTGGCAACGTTTTTGGATTCAAAATGCTAAGAAGCATTCGTCTTGAAGATATGCGCATCCCTCTTGCTTATTTAACAACTTTTCAAGGTCCGGCAACAGGTGTTGAAGAAGAACGTGCACGTCTGGACAAATATGGACGACCACTTATTGGAGCCAACATTCGACCTAAGTTCGGTCTTTCGGGTAGAGATGAAGGTCGAGCTGTCTATGCAGGTCTTAAAGGTGGTCTTGATTACCTCAAAGGTGGTGAAAATAATAGTAGTCAATCATTTATGCGTTGGAAAGAACG

>Pinguiophyceae_AF438321.1 Phaeomonas parva ribulose-1,5-bisphosphate carboxylase/oxygenase large subunit (rbcL) gene, partial cds; chloroplast gene for chloroplast product

AAACTTAACTGCGTCTATCATTGGTAACGTATTTGGTTTCAAAGCTGTTAAAGCTCTTCGTTTAGAAGATATGCGTATCCCTTACGCTTACTTAAAGACTTTCCAAGGTCCTGCTACTGGTTTAGTTGTAGAACGTGAGCGTATGGATAAGTACGGTCGTCCATTATTAGGTGCAACTGTAAAACCTAAGTTAGGTTTATCTGGTAAGAACTACGGTCGTGTAGTATACGAAGGTTTACGCGGTGGTTTAGATTTCTTAAAGGATGATGAAAACATTAACTCTCAACCATTCATGCGTTGGAGAGAGCG

>Pinguiophyceae_HQ710625.1 Pinguiochrysis pyriformis voucher PP 301 ribulose-1,5-bisphosphate carboxylase/oxygenase large subunit (rbcL) gene, partial cds; plastid

AAACTTAACTGCGTCTATCATTGGTAACGTATTCGGTTTCAAGGCCGTTAAAGCTCTTCGTTTAGAAGATATGCGTATCCCTTATGCTTACTTAAAGACTTTCCAAGGTCCTGCAACTGGTTTAATTGTAGAACGTGAGCGTATGGATAAGTATGGTCGTCCATTCTTAGGTGCGACTGTAAAGCCTAAGTTAGGTTTATCTGGTAAGAACTACGGTCGTGTAGTATACGAAGGTTTACGTGGTGGTTTAGATTTCTTAAAGGATGATGAAAACATTAACTCTCAACCATTCATGCGTTGGAGAGAGCG

>Pinguiophyceae_AF438317.1 Pinguiochrysis pyriformis ribulose-1,5-bisphosphate carboxylase/oxygenase large subunit (rbcL) gene, partial cds; chloroplast gene for chloroplast product

AAACTTAACTGCGTCTATCATTGGTAACGTATTCGGTTTCAAGGCCGTTAAAGCTCTTCGTTTAGAAGATATGCGTATCCCTTATGCTTACTTAAAGACTTTCCAAGGTCCTGCAACTGGTTTAATTGTAGAACGTGAGCGTATGGATAAGTATGGTCGTCCATTCTTAGGTGCGACTGTAAAGCCTAAGTTAGGTTTATCTGGTAAGAACTACGGTCGTGTAGTATACGAAGGTTTACGTGGTGGTTTAGATTTCTTAAAGGATGATGAAAACATTAACTCTCAACCATTCATGCGTTGGAGAGAGCG

>AB034635.1:1-1024 Karenia mikimotoi chloroplast rbcL gene for large subunit of RubisCO, partial cds

-AATTTAACCGCATCAATAATTGGAAATGTTTTCGGAATGAAAGCGGTTCAGTCGCTTAGATTAGAAGACATGCGAATCCCTGTGGCTTATTTAAAAACGTTCCAAGGTCCTGCCACTGGATTAGTTGTTGAACGCGAACGACTTGACAAGTTTGGGCGTCCACTTCTTGGGGCAACAGTTAAACCAAAGCTAGGATTATCAGGTAAAAACTATGGCCGAGTCGTATATGAAGGTCTAAAAGGAGGCTTGGATTTCTTAAAGGATGATGAGAACATTAACTCACAACCGTTTATGCGTTATCGAGAACG

>JX899690.2:1-963 Karenia mikimotoi ribulose-1,5-bisphosphate carboxylase/oxygenase large subunit (rbcL) mRNA, partial cds; chloroplast

-AATTTAACCGCATCAATAATTGGAAATGTTTTCGGAATGAAAGCGGTTCAGTCGCTTAGATTAGAAGACATGCGAATCCCTGTAGCTTATCTAAAAACGTTCCAAGGTCCTGCTACGGGACTAGTTGTTGAACGCGAACGACTTGACAAATTTGGGCGTCCACTCCTCGGAGCAACAGTTAAACCAAAGCTAGGCTTATCGGGTAAAAATTACGGTAGGGTCGTGTACGAAGGTCTAAAAGGAGGATTGGATTTCCTAAAGGATGATGAGAACATTAACTCACAACCGTTTATGCGTTACCGAGAACG

>JX899685.3:32-1055 Karenia mikimotoi ribulose-1,5-bisphosphate carboxylase/oxygenase large subunit (rbcL) gene, partial cds; and tRNA-Phe gene, complete sequence; chloroplast

-AATTTAACCGCATCAATAATTGGAAATGTTTTCGGAATGAAAGCGGTTAAGTCGCTTAGATTAGAAGACATGCGAATCCCTGTAGCTTATCTAAAAACGTTTCAAGGTCCTGCTACGGGATTAGTTGTTGAACGCGAACGACTTGACAAATTTGGGCGTCCACTCCTCGGAGCAACAGTTAAACCAAAGCTAGGCTTATCGGGTAAAAATTACGGTAGGGTCGTGTACGAAGGTTTAAAAGGAGGATTGGATTTTCTAAAGGATGATGAGAACATTAATTCACAACCGTTTATGCGTTACCGAGAACG

>KR935867.1:505-1430 Karenia brevis strain CCM2281 ribulose-1,5-bisphosphate carboxylase/oxygenase large subunit (rbcL) mRNA, complete cds; chloroplast

-AATTTAACAGCCTCCATAATTGGTAATATTTTTGGTTTCAAAGCTGTAAAGTCTCTTCGGCTTGAGGATATGAGAATCCCGGTGGCTTATCTAAAAACGTTCCAAGGTCCAGCTACTGGTTTAGTCGTTGAACGAGAACGTCTAGATAAGTTCGGTCGACCTCTTTTGGGAGCGACAGTTAAACCAAAATTAGGTCTTTCCGGTAAAAATTATGGAAGAGTTGTATACGAAGGGTTAAAAGGGGGATTAGATTTCCTTAAAGATGATGAAAATATTAATTCGCAGCCTTTCATGCGGTATCGTGAACG

>AY119786.1:1-907 Karenia brevis strain CCMP 718 ribulose-1,5-bisphosphate carboxylase/oxygenase (rbcL) gene, partial cds; chloroplast

-AATTTAACAGCCTCCATAATTGGTAATATTTTTGGTTTTAAAGCTGTAAAGTCTCTTCGGCTTGAGGATATGAGAATCCCGGTGGCTTATCTAAAAACGTTTCAAGGTCCAGCTACTGGTTTAGTCGTTGAACGAGAACGTCTAGATAAGTTCGGTCGACCTCTTTTGGGAGCGACAGTTAAACCAAAATTAGGTCTTTCCGGTAAAAATTATGGAAGAGTTGTATACGAAGGGTTAAAAGGGGGATTAGATTTTCTTAAAGATGATGAAAATATTAATTCGCAGCCTTTTATGCGGTATCGTGAACG

>KT149178.1 Nannochloropsis oceanica strain CS-246 ribulose-1,5-bisphosphate carboxylase/oxygenase large subunit (rbcL) gene, partial cds; plastid

-AACTTAACAGCTTCAATTATCGGTAACGTATTTGGATTCAAAGCTGTAAAAGCATTACGTCTTGAAGATATGCGTATGCCTTACGCTTACTTAAAAACATTCCAAGGTCCAGCTACTGGTGTGATTGTTGAACGTGAGCGTTTAGACAAATTCGGACGTCCTTTATTAGGTGCAACTGTAAAACCTAAACTTGGTTTATCAGGTAAAAACTATGGACGTGTTGTATACGAAGGTTTAAAAGGTGGTTTAGACTTCTTAAAAGATGACGAAAACATTAACTCTCAACCATTCATGCGTTGGCGTGAACG

>KT149177.1 Nannochloropsis oceanica strain CS-179 ribulose-1,5-bisphosphate carboxylase/oxygenase large subunit (rbcL) gene, partial cds; plastid

-AACTTAACAGCTTCAATTATCGGTAACGTATTTGGATTCAAAGCTGTAAAAGCATTACGTCTTGAAGATATGCGTATGCCTTACGCTTACTTAAAAACATTCCAAGGTCCAGCTACTGGTGTGATTGTTGAACGTGAGCGTTTAGACAAATTCGGACGTCCTTTATTAGGTGCAACTGTAAAACCTAAACTTGGTTTATCAGGTAAAAACTATGGACGTGTTGTATACGAAGGTTTAAAAGGTGGTTTAGACTTCTTAAAAGATGACGAAAACATTAACTCTCAACCATTCATGCGTTGGCGTGAACG

>HQ710610.1 Nannochloropsis oceanica voucher EUS-001 ribulose-1,5-bisphosphate carboxylase/oxygenase large subunit (rbcL) gene, partial cds; plastid

-AACTTAACAGCTTCAATTATCGGTAACGTATTTGGATTCAAAGCTGTAAAAGCATTACGTCTTGAAGATATGCGTATGCCTTACGCTTACTTAAAAACATTCCAAGGTCCAGCTACTGGTGTGATTGTTGAACGTGAGCGTTTAGACAAATTCGGACGTCCTTTATTAGGTGCAACTGTAAAACCTAAACTTGGTTTATCAGGTAAAAACTATGGACGTGTTGTATACGAAGGTTTAAAAGGTGGTTTAGACTTCTTAAAAGATGACGAAAACATTAACTCTCAACCATTCATGCGTTGGCGTGAACG

>AB052285.1 Nannochloropsis oceanica chloroplast rbcL gene for ribulose-1,5-bisphosphate carboxylase/oxygenase large subunit, partial cds, strain:MBIC10440

-AACTTAACAGCTTCAATTATCGGTAACGTATTTGGATTCAAAGCTGTAAAAGCATTACGTCTTGAAGATATGCGTATGCCTTACGCTTACTTAAAAACATTCCAAGGTCCAGCTACTGGTGTGATTGTTGAACGTGAGCGTTTAGACAAATTCGGACGTCCTTTATTAGGTGCAACTGTAAAACCTAAACTTGGTTTATCAGGTAAAAACTATGGACGTGTTGTATACGAAGGTTTAAAAGGTGGTTTAGACTTCTTAAAAGATGACGAAAACATTAACTCTCAACCATTCATGCGTTGGCGTGAACG

>AB052284.1 Nannochloropsis oceanica chloroplast rbcL gene for ribulose-1,5-bisphosphate carboxylase/oxygenase large subunit, partial cds, strain:MBIC10426

-AACTTAACAGCTTCAATTATCGGTAACGTATTTGGATTCAAAGCTGTAAAAGCATTACGTCTTGAAGATATGCGTATGCCTTACGCTTACTTAAAAACATTCCAAGGTCCAGCTACTGGTGTGATTGTTGAACGTGAGCGTTTAGACAAATTCGGACGTCCTTTATTAGGTGCAACTGTAAAACCTAAACTTGGTTTATCAGGTAAAAACTATGGACGTGTTGTATACGAAGGTTTAAAAGGTGGTTTAGACTTCTTAAAAGATGACGAAAACATTAACTCTCAACCATTCATGCGTTGGCGTGAACG

>AB052283.1 Nannochloropsis oceanica chloroplast rbcL gene for ribulose-1,5-bisphosphate carboxylase/oxygenase large subunit, partial cds, strain:MBIC10179

-AACTTAACAGCTTCAATTATCGGTAACGTATTTGGATTCAAAGCTGTAAAAGCATTACGTCTTGAAGATATGCGTATGCCTTACGCTTACTTAAAAACATTCCAAGGTCCAGCTACTGGTGTGATTGTTGAACGTGAGCGTTTAGACAAATTCGGACGTCCTTTATTAGGTGCAACTGTAAAACCTAAACTTGGTTTATCAGGTAAAAACTATGGACGTGTTGTATACGAAGGTTTAAAAGGTGGTTTAGACTTCTTAAAAGATGACGAAAACATTAACTCTCAACCATTCATGCGTTGGCGTGAACG

>AB052281.1 Nannochloropsis oceanica chloroplast rbcL gene for ribulose-1,5-bisphosphate carboxylase/oxygenase large subunit, partial cds, strain:MBIC10090

-AACTTAACAGCTTCAATTATCGGTAACGTATTTGGATTCAAAGCTGTAAAAGCATTACGTCTTGAAGATATGCGTATGCCTTACGCTTACTTAAAAACATTCCAAGGTCCAGCTACTGGTGTGATTGTTGAACGTGAGCGTTTAGACAAATTCGGACGTCCTTTATTAGGTGCAACTGTAAAACCTAAACTTGGTTTATCAGGTAAAAACTATGGACGTGTTGTATACGAAGGTTTAAAAGGTGGTTTAGACTTCTTAAAAGATGACGAAAACATTAACTCTCAACCATTCATGCGTTGGCGTGAACG

>KP057242.1 Nannochloropsis oceanica ribulose-1,5-bisphosphate carboxylase/oxygenase large subunit gene, partial cds; chloroplast

-AACTTAACAGCTTCAATTATCGGTAACGTATTTGGATTCAAAGCTGTAAAAGCATTACGTCTTGGAGATATGCGTATGCCTTACGCTTACTTAAAAACATTCCAAGGTCCAGCTACTGGTGTGATTGTTGAACGTGAGCGTTTAGACAAATTCGGACGTCCTTTATTAGGTGCAACTGTAAAACCTAAACTTGGTTTATCAGGTAAAAACTATGGACGTGTTGTATACGAAGGTTTAAAAGGTGGTTTAGACTTCTTAAAAGATGACGAAAACATTAACTCTCAACCATTCATGCGTTGGCGTGAACG

>JX913540.1 Nannochloropsis sp. SC-2012 ribulose-1,5-bisphosphate carbxylase/oxygenase large subunit (rbcL) gene, partial cds; chromoplast

-AACTTAACAGCTTCAATTATCGGTAACGTATTTGGATTCAAAGCTGTAAAAGCATTACGTCTTGAAGATATGCGTATGCCTTACGCTTACTTAAAAACATTCCAAGGTCCAGCTACTGGTGTGATTGTTGAACGTGAGCGTTTAGACAAATTCGGACGTCCTTTATTAGGTGCAACTGTAAAACCTAAACTTGGTTTATCAGGTAAAAACTATGGACGTGTTGTATACGAAGGTTTAAAAGGTGGTTTAGACTTCTTAAAAGATGACGAAAACATTAACTCTCAACCATTCATGCGTTGGCGTGAACG

>AY680702.1 Nannochloropsis maritima ribulose-1,5-bisphosphate carbxylase/oxygenase large subunit (rbcL) gene, partial cds; chloroplast

-AACTTAACAGCTTCAATTATCGGTAACGTATTTGGATTCAAAGCTGTAAAAGCATTACGTCTTGAAGATGTGCGTATGCCTTACGCTTACTTAAAAACATTCCAAGGTCCAGCTACTGGTGTGATTGTTGAACGTGAGCGTTTAGACAAATTCGGACGTCCTTTATTAGGTGCAACTGTAAAACCTAAACTTGGTTTATCAGGTAAAAACTATGGACGTGTTGTATACGAAGGTTTAAAAGGTGGTTTAGACTTCTTAAAAGATGACGAAAACATTAACTCTCAACCATTCATGCGTTGGCGTGAACG

>HQ710611.1 Nannochloropsis sp. HSY-2011 voucher EC-009 ribulose-1,5-bisphosphate carboxylase/oxygenase large subunit (rbcL) gene, partial cds; plastid

-AACTTAACAGCTTCAATTATCGGTAACGTATTTGGATTCAAAGCTGTAAAAGCATTACGTCTTGAAGATATGCGTATGCCTTACGCTTACTTAAAAACATTCCAAGGTCCAGCTACTGGTGTGATTGTTGAACGTGAGCGTTTAGACAAATTCGGACGTCCTTTATTAGGTGCAACTGTAAAACCTAAACTTGGTTTATCAGGTAAAAACTATGGACGTGTTGTATACGAAGGTTTAAAAGGTGGTTTAGACTTCTTAAAAGATGACGAAAACATTAACTCTCAACCATTCATGCGTTGGCGTGAACG

>KF010153.1 Nannochloropsis oceanica isolate CCALA978 ribulose-1,5- bisphosphate carboxylase oxygenase large subunit (rbcL) gene, partial cds; chloroplast

-AACTTAACAGCTTCAATTATCGGTAACGTATTTGGATTCAAAGCTGTAAAAGCATTACGTCTTGAAGATATGCGTATGCCTTACGCTTACTTAAAAACATTCCAAGGTCCAGCTACTGGTGTGATTGTTGAACGTGAGCGTTTAGACAAATTCGGACGTCCTTTATTAGGTGCAACTGTAAAACCTAAACTTGGTTTATCAGGTAAAAACTATGGACGTGTTGTATACGAAGGTTTAAAAGGTGGTTTAGACTTCTTAAAAGATGACGAAAACATTAACTCTCAACCATTCATGCGTTGGCGTGAACG

>KT149187.1 Nannochloropsis australis strain CS-759 ribulose-1,5-bisphosphate carboxylase/oxygenase large subunit (rbcL) gene, partial cds; plastid

-AACTTAACAGCTTCAATTATCGGTAACGTATTTGGATTTAAAGCTGTAAAAGCATTACGTCTTGAAGATATGCGTATGCCTTACGCTTACTTAAAAACATTCCAAGGCCCAGCTACTGGTGTGATTGTTGAACGTGAGCGTTTAGACAAATTCGGACGTCCTTTATTAGGTGCAACTGTAAAACCTAAACTTGGTTTATCAGGTAAAAACTATGGACGTGTTGTATACGAAGGTCTAAAAGGTGGTTTAGACTTCTTAAAAGATGACGAAAACATTAACTCTCAACCATTCATGCGTTGGCGTGAACG

>KT149179.1 Nannochloropsis australis strain CS-416 ribulose-1,5-bisphosphate carboxylase/oxygenase large subunit (rbcL) gene, partial cds; plastid

-AACTTAACAGCTTCAATTATCGGTAACGTATTTGGATTTAAAGCTGTAAAAGCATTACGTCTTGAAGATATGCGTATGCCTTACGCTTACTTAAAAACATTCCAAGGCCCAGCTACTGGTGTGATTGTTGAACGTGAGCGTTTAGACAAATTCGGACGTCCTTTATTAGGTGCAACTGTAAAACCTAAACTTGGTTTATCAGGTAAAAACTATGGACGTGTTGTATACGAAGGTCTAAAAGGTGGTTTAGACTTCTTAAAAGATGACGAAAACATTAACTCTCAACCATTCATGCGTTGGCGTGAACG

>KT149182.1 Nannochloropsis oceanica strain CS-699 ribulose-1,5-bisphosphate carboxylase/oxygenase large subunit (rbcL) gene, partial cds; plastid

-AACTTAACAGCTTCAATTATCGGTAACGTATTTGGATTCAAAGCTGTAAAAGCATTACGTCTTGAAGATATGCGTATGCCTTACGCTTACTTAAAAACATTCCAAGGTCCAGCTACTGGTGTGATTGTTGAACGTGAGCGTTTAGACAAATTCGGACGTCCTTTATTAGGTGCAACTGTAAAACCTAAACTTGGTTTATCAGGTAAAAACTATGGACGTGTTGTATACGAAGGTTTAAAAGGTGGTTTAGACTTCTTAAAAGATGACGAAAACATTAACTCTCAACCATTCATGCGTTGGCGTGAACG

>KC128502.1 Nannochloropsis granulata strain BDH02 ribulose-1,5-bisphosphate carboxylase/oxygenase large subunit (rbcL) gene, partial cds; chloroplast

-AACTTAACAGCTTCAATTATCGGTAACGTATTTGGATTCAAAGCTGTAAAAGCATTACGTCTTGAAGATATGCGTATGCCTTACGCTTACTTAAAAACATTCCAAGGCCCAGCTACTGGTGTGATTGTTGAACGTGAGCGTTTAGACAAATTCGGACGTCCTTTATTAGGTGCAACTGTAAAACCTAAACTTGGTTTATCAGGTAAAAACTATGGACGTGTTGTATACGAAGGTTTAAAAGGTGGTTTAGACTTCTTAAAAGATGACGAAAACATTAACTCTCAACCATTCATGCGTTGGCGTGAACG

>AB052280.1 Nannochloropsis granulata chloroplast rbcL gene for ribulose-1,5-bisphosphate carboxylase/oxygenase large subunit, partial cds, strain:MBIC10054

-AACTTAACAGCTTCAATTATCGGTAACGTATTTGGATTCAAAGCTGTAAAAGCATTACGTCTTGAAGATATGCGTATGCCTTACGCTTACTTAAAAACATTCCAAGGCCCAGCTACTGGTGTGATTGTTGAACGTGAGCGTTTAGACAAATTCGGACGTCCTTTATTAGGTGCAACTGTAAAACCTAAACTTGGTTTATCAGGTAAAAACTATGGACGTGTTGTATACGAAGGTTTAAAAGGTGGTTTAGACTTCTTAAAAGATGACGAAAACATTAACTCTCAACCATTCATGCGTTGGCGTGAACG

>HQ710609.1 Nannochloropsis oculata culture CCMP:525 ribulose-1,5-bisphosphate carboxylase/oxygenase large subunit (rbcL) gene, partial cds; plastid

-AACTTAACAGCTTCAATTATCGGTAACGTATTTGGATTCAAAGCTGTAAAAGCATTACGTCTTGAAGATATGCGTATGCCTTACGCTTACTTAAAAACATTCCAAGGTCCAGCTACTGGTGTGATTGTTGAACGTGAGCGTTTAGACAAATTCGGACGTCCTTTATTAGGTGCAACTGTAAAACCTAAACTTGGTTTATCAGGTAAAAACTATGGACGTGTTGTATACGAAGGTTTAAAAGGTGGTTTAGACTTCTTAAAAGATGATGAAAACATTAACTCTCAACCATTCATGCGTTGGCGTGAACG

>AB052286.1 Nannochloropsis oculata chloroplast rbcL gene for ribulose-1,5-bisphosphate carboxylase/oxygenase large subunit, partial cds, strain:CCAP849/1

-AACTTAACAGCTTCAATTATCGGTAACGTATTTGGATTCAAAGCTGTAAAAGCATTACGTCTTGAAGATATGCGTATGCCTTACGCTTACTTAAAAACATTCCAAGGTCCAGCTACTGGTGTGATTGTTGAACGTGAGCGTTTAGACAAATTCGGACGTCCTTTATTAGGTGCAACTGTAAAACCTAAACTTGGTTTATCAGGTAAAAACTATGGACGTGTTGTATACGAAGGTTTAAAAGGTGGTTTAGACTTCTTAAAAGATGATGAAAACATTAACTCTCAACCATTCATGCGTTGGCGTGAACG

>AB280614.1 Nannochloropsis oculata chromoplast rbcL, rbcS genes for ribulose 1.5-bisphosphate carboxylase/oxygenase large subunit, ribulose 1.5-bisphosphate carboxylase/oxygenase small subunit, partial cds

-AACTTAACAGCTTCAATTATCGGTAACGTATTTGGATTCAAAGCTGTAAAAGCATTACGTCTTGAAGATATGCGTATGCCTTACGCTTACTTAAAAACATTCCAAGGTCCAGCTACTGGTGTGATTGTTGAACGTGAGCGTTTAGACAAATTCGGACGTCCTTTATTAGGTGCAACTGTAAAACCTAAACTTGGTTTATCAGGTAAAAACTATGGACGTGTTGTATACGAAGGTTTAAAAGGTGGTTTAGACTTCTTAAAAGATGATGAAAACATTAACTCTCAACCATTCATGCGTTGGCGTGAACG

>DQ977734.1 Nannochloropsis sp. JL2/4-1 ribulose-1,5-bisphosphate carbxylase/oxygenase large subunit (rbcL) gene, partial cds; chloroplast

-AACTTAACAGCTTCAATTATCGGTAACGTATTTGGATTCAAAGCTGTAAAAGCATTACGTCTTGAAGATATGCGTATGCCTTACGCTTACTTAAAAACATTCCAAGGTCCAGCTACTGGTGTGGTTGTTGAACGTGAGCGTTTAGACAAATTCGGACGTCCTTTATTAGGTGCAACTGTAAAACCTAAACTTGGTTTATCAGGTAAAAACTATGGACGTGTTGTATACGAAGGTTTAAAAGGTGGTTTAGACTTCTTAAAAGATGACGAAAACATCAACTCTCAACCATTCATGCGTTGGCGTGAACG

>HQ201713.1 Nannochloropsis oceanica strain LAMB0001 ribulose-1,5-bisphosphate carboxylase/oxygenase large subunit (rbcL) gene, partial cds; chloroplast

-AACTTAACAGCTTCAATTATCGGTAACGTATTTGGATTCAAAGCTGTAAAAGCATTACGTCTTGAAGATATGCGTATGCCTTACGCTTACTTAAAAACATTCCAAGGTCCAGCTACTGGTGTGATTGTTGAACGTGAGCGTTTAGACAAATTCGGACGTCCTTTATTAGGTGCAACTGTAAAACCTAAACTTGGTTTATCAGGTAAAAACTATGGACGTGTTGTATACGAAGGTTTAAAAGGTGGTTTAGACTTCTTAAAAGATGACGAAAACATTAACTCTCAACCATTCATGCGTTGGCGTGAACG

>DQ977744.1 Nannochloropsis sp. AN1/12-10 ribulose-1,5-bisphosphate carbxylase/oxygenase large subunit (rbcL) gene, partial cds; chloroplast

-AACTTAACAGCTTCAATTATCGGTAACGTATTTGGATTCAAAGCTGTAAAAGCATTACGTCTTGAAGATATGCGTATGCCTTACGCTTACTTAAAAACATTCCAAGGTCCAGCTACTGGTGTGGTTGTTGAACGTGAGCGTTTAGACAAATTCGGACGTCCTTTATTAGGTGCAACTGTAAAACCAAAACTTGGTTTATCAGGTAAAAACTATGGACGTGTTGTATACGAAGGTTTAAAAGGTGGATTAGACTTCTTAAAAGATGACGAAAACATCAACTCTCAACCATTCATGCGTTGGCGTGAACG

>DQ977732.1 Nannochloropsis sp. MDL3-4 ribulose-1,5-bisphosphate carbxylase/oxygenase large subunit (rbcL) gene, partial cds; chloroplast

-AACTTAACAGCTTCAATTATCGGTAACGTATTTGGATTCAAAGCTGTAAAAGCATTACGTCTTGAAGATATGCGTATGCCTTACGCTTACTTAAAAACATTCCAAGGTCCAGCTACTGGTGTGGTTGTTGAACGTGAGCGTTTAGACAAATTCGGACGTCCTTTATTAGGTGCAACTGTAAAACCTAAACTTGGTTTATCAGGTAAAAACTATGGACGTGTTGTATACGAAGGTTTAAAAGGTGGATTAGACTTCTTAAAAGATGACGAAAACATCAACTCTCAACCATTCATGCGTTGGCGTGAACG

>DQ977731.1 Nannochloropsis sp. AN1/12-5 ribulose-1,5-bisphosphate carbxylase/oxygenase large subunit (rbcL) gene, partial cds; chloroplast

-AACTTAACAGCTTCAATTATCGGTAACGTATTTGGATTCAAAGCTGTAAAAGCATTACGTCTTGAAGATATGCGTATGCCTTACGCTTACTTAAAAACATTCCAAGGTCCAGCTACTGGTGTGGTTGTTGAACGTGAGCGTTTAGACAAATTCGGACGTCCTTTATTAGGTGCAACTGTAAAACCAAAACTTGGTTTATCAGGTAAAAACTATGGACGTGTTGTATACGAAGGTTTAAAAGGTGGATTAGACTTCTTAAAAGATGACGAAAACATCAACTCTCAACCATTCATGCGTTGGCGTGAACG

>AM421006.1 Nannochloropsis limnetica chloroplast partial rbcL gene for ribulose bisphosphate carboxylase large chain, strain SAG 18.99

-AACTTAACAGCTTCAATTATCGGTAACGTATTTGGATTCAAAGCTGTAAAAGCATTACGTCTTGAAGATATGCGTATGCCTTACGCTTACTTAAAAACATTCCAAGGTCCAGCTACTGGTGTGGTTGTTGAACGTGAGCGTTTAGACAAATTCGGACGTCCTTTATTAGGTGCAACTGTAAAACCAAAACTTGGTTTATCAGGTAAAAACTATGGACGTGTTGTATACGAAGGTTTAAAAGGTGGATTAGACTTCTTAAAAGATGACGAAAACATCAACTCTCAACCATTCATGCGTTGGCGTGAACG

>DQ977743.1 Nannochloropsis sp. MDL11-16 ribulose-1,5-bisphosphate carbxylase/oxygenase large subunit (rbcL) gene, partial cds; chloroplast

-AACTTAACAGCTTCAATTATCGGTAACGTATTTGGATTCAAAGCTGTAAAAGCATTACGTCTTGAAGATATGCGTATGCCTTACGCTTACTTAAAAACATTCCAAGGTCCAGCTACTGGTGTGGTTGTTGAACGTGAGCGTTTAGACAAATTCGGACGTCCTTTATTAGGTGCAACTGTAAAACCTAAACTTGGTTTATCAGGTAAAAACTATGGACGTGTTGTATACGAAGGTTTAAAAGGTGGTTTAGACTTCTTAAAAGATGACGAAAACATCAACTCTCAACCATTCATGCGTTGGCGTGAACG

>DQ977740.1 Nannochloropsis limnetica isolate MDL11-14 ribulose-1,5-bisphosphate carbxylase/oxygenase large subunit (rbcL) gene, partial cds; chloroplast

-AACTTAACAGCTTCAATTATCGGTAACGTATTTGGATTCAAAGCTGTAAAAGCATTACGTCTTGAAGATATGCGTATGCCTTACGCTTACTTAAAAACATTCCAAGGTCCAGCTACTGGTGTGGTTGTTGAACGTGAGCGTTTAGACAAATTCGGACGTCCTTTATTAGGTGCAACTGTAAAACCAAAACTTGGTTTATCAGGTAAAAACTATGGACGTGTTGTATACGAAGGTTTAAAAGGTGGATTAGACTTCTTAAAAGATGACGAAAACATCAACTCTCAACCATTCATGCGTTGGCGTGAACG

>DQ977737.1 Nannochloropsis sp. JL11-8 ribulose-1,5-bisphosphate carbxylase/oxygenase large subunit (rbcL) gene, partial cds; chloroplast

-AACTTAACAGCTTCAATTATCGGTAACGTATTTGGATTCAAAGCTGTAAAAGCATTACGTCTTGAAGATATGCGTATGCCTTACGCTTACTTAAAAACATTCCAAGGTCCAGCTACTGGTGTGGTTGTTGAACGTGAGCGTTTAGACAAATTCGGACGTCCTTTATTAGGTGCAACTGTAAAACCTAAACTTGGTTTATCAGGTAAAAACTATGGACGTGTTGTATACGAAGGTTTAAAAGGTGGTTTAGACTTCTTAAAAGATGACGAAAACATCAACTCTCAACCATTCATGCGTTGGCGTGAACG

>DQ977733.1 Nannochloropsis sp. AN1/12-7 ribulose-1,5-bisphosphate carbxylase/oxygenase large subunit (rbcL) gene, partial cds; chloroplast

-AACTTAACAGCTTCAATTATCGGTAACGTATTTGGATTCAAAGCTGTAAAAGCATTACGTCTTGAAGATATGCGTATGCCTTACGCTTACTTAAAAACATTCCAAGGTCCAGCTACTGGTGTGGTTGTTGAACGTGAGCGTTTAGACAAATTCGGACGTCCTTTATTAGGTGCAACTGTAAAACCTAAACTTGGTTTATCAGGTAAAAACTATGGACGTGTTGTATACGAAGGTTTAAAAGGTGGTTTAGACTTCTTAAAAGATGACGAAAACATCAACTCTCAACCATTCATGCGTTGGCGTGAACG

>DQ977729.1 Nannochloropsis limnetica isolate KR1998/3 ribulose-1,5-bisphosphate carbxylase/oxygenase large subunit (rbcL) gene, partial cds; chloroplast

-AACTTAACAGCTTCAATTATCGGTAACGTATTTGGATTCAAAGCTGTAAAAGCATTACGTCTTGAAGATATGCGTATGCCTTACGCTTACTTAAAAACATTCCAAGGTCCAGCTACTGGTGTGGTTGTTGAACGTGAGCGTTTAGACAAATTCGGACGTCCTTTATTAGGTGCAACTGTAAAACCAAAACTTGGTTTATCAGGTAAAAACTATGGACGTGTTGTATACGAAGGTTTAAAAGGTGGATTAGACTTCTTAAAAGATGACGAAAACATCAACTCTCAACCATTCATGCGTTGGCGTGAACG

>DQ977741.1 Nannochloropsis limnetica isolate AS3-9 ribulose-1,5-bisphosphate carbxylase/oxygenase large subunit (rbcL) gene, partial cds; chloroplast

-AACTTAACAGCTTCAATTATCGGTAACGTATTTGGATTCAAAGCTGTAAAAGCATTACGTCTTGAAGATATGCGTATGCCTTACGCTTACTTAAAAACATTCCAAGGTCCAGCTACTGGTGTGGTTGTTGAACGTGAGCGTTTAGACAAATTCGGACGTCCTTTATTAGGTGCAACTGTAAAACCAAAACTTGGTTTATCAGGTAAAAACTATGGACGTGTTGTATACGAAGGTTTAAAAGGTGGATTAGACTTCTTAAAAGATGACGAAAACATCAACTCTCAACCATTCATGCGTTGGCGTGAACG

>DQ977739.1 Nannochloropsis limnetica isolate AS2/16-8 ribulose-1,5-bisphosphate carbxylase/oxygenase large subunit (rbcL) gene, partial cds; chloroplast

-AACTTAACAGCTTCAATTATCGGTAACGTATTTGGATTCAAAGCTGTAAAAGCATTACGTCTTGAAGATATGCGTATGCCTTACGCTTACTTAAAAACATTCCAAGGTCCAGCTACTGGTGTGGTTGTTGAACGTGAGCGTTTAGACAAATTCGGACGTCCTTTATTAGGTGCAACTGTAAAACCAAAACTTGGTTTATCAGGTAAAAACTACGGACGTGTTGTATACGAAGGTTTAAAAGGTGGATTAGACTTCTTAAAAGATGACGAAAACATCAACTCTCAACCATTCATGCGTTGGCGTGAACG

>DQ977736.1 Nannochloropsis sp. Tow 2/24 P-1w ribulose-1,5-bisphosphate carbxylase/oxygenase large subunit (rbcL) gene, partial cds; chloroplast

-AACTTAACAGCTTCAATTATCGGTAACGTATTTGGATTCAAAGCTGTAAAAGCATTACGTCTTGAAGATATGCGTATGCCTTACGCTTACTTAAAAACATTCCAAGGTCCAGCTACTGGTGTGGTTGTTGAACGTGAGCGTTTAGACAAATTCGGACGTCCTTTATTAGGTGCAACTGTAAAACCTAAACTTGGTTTATCAGGTAAAAACTATGGACGTGTTGTATACGAAGGTTTAAAAGGTGGATTAGACTTCTTAAAAGATGACGAAAACATCAACTCTCAACCATTCATGCGTTGGCGTGAACG

>DQ977730.1 Nannochloropsis limnetica isolate JL1/12-5 ribulose-1,5-bisphosphate carbxylase/oxygenase large subunit (rbcL) gene, partial cds; chloroplast

-AACTTAACAGCTTCAATTATCGGTAACGTATTTGGATTCAAAGCTGTAAAAGCATTACGTCTTGAAGATATGCGTATGCCTTACGCTTACTTAAAAACATTCCAAGGTCCAGCTACTGGTGTGGTTGTTGAACGTGAGCGTTTAGACAAATTCGGACGTCCTTTATTAGGTGCAACTGTAAAACCAAAACTTGGTTTATCAGGTAAAAACTACGGACGTGTTGTATACGAAGGTTTAAAAGGTGGATTAGACTTCTTAAAAGATGACGAAAACATCAACTCTCAACCATTCATGCGTTGGCGTGAACG

>DQ977742.1 Nannochloropsis sp. AN2/29-2 ribulose-1,5-bisphosphate carbxylase/oxygenase large subunit (rbcL) gene, partial cds; chloroplast

-AACTTAACAGCTTCAATTATCGGTAACGTATTTGGATTCAAAGCTGTAAAAGCATTACGTCTTGAAGATATGCGTATGCCTTACGCTTACTTAAAAACATTCCAAGGTCCAGCTACTGGTGTGGTTGTTGAACGTGAGCGTTTAGACAAATTCGGACGTCCTTTATTAGGTGCAACTGTAAAACCAAAACTTGGTTTATCAGGTAAAAACTATGGACGTGTTGTATACGAAGGTTTAAAAGGTGGTTTAGACTTCTTAAAAGATGACGAAAACATCAACTCTCAACCATTCATGCGTTGGCGTGAACG

>DQ977738.1 Nannochloropsis sp. AN2/29-6 ribulose-1,5-bisphosphate carbxylase/oxygenase large subunit (rbcL) gene, partial cds; chloroplast

-AACTTAACAGCTTCAATTATCGGTAACGTATTTGGATTCAAAGCTGTAAAAGCATTACGTCTTGAAGATATGCGTATGCCTTACGCTTACTTAAAAACATTCCAAGGTCCAGCTACTGGTGTGGTTGTTGAACGTGAGCGTTTAGACAAATTCGGACGTCCTTTATTAGGTGCAACTGTAAAACCAAAACTTGGTTTATCAGGTAAAAACTATGGACGTGTTGTATACGAAGGTTTAAAAGGTGGTTTAGACTTCTTAAAAGATGACGAAAACATCAACTCTCAACCATTCATGCGTTGGCGTGAACG

>DQ977735.1 Nannochloropsis sp. AS4-1 ribulose-1,5-bisphosphate carbxylase/oxygenase large subunit (rbcL) gene, partial cds; chloroplast

-AACTTAACAGCTTCAATTATCGGTAACGTATTTGGATTCAAAGCTGTAAAAGCATTACGTCTTGAAGATATGCGTATGCCTTACGCTTACTTAAAAACATTCCAAGGTCCAGCTACTGGTGTGGTTGTTGAACGTGAGCGTTTAGACAAATTCGGACGTCCTTTATTAGGTGCAACTGTAAAACCAAAACTTGGTTTATCAGGTAAAAACTATGGACGTGTTGTATACGAAGGTTTAAAAGGTGGTTTAGACTTCTTAAAAGATGACGAAAACATCAACTCTCAACCATTCATGCGTTGGCGTGAACG

>KT149185.1 Nannochloropsis oceanica strain CS-702 ribulose-1,5-bisphosphate carboxylase/oxygenase large subunit (rbcL) gene, partial cds; plastid

-AACTTAACAGCTTCAATTATCGGTAACGTATTTGGATTCAAAGCTGTAAAAGCATTACGTCTTGAAGATATGCGTATGCCTTACGCTTACTTAAAAACATTCCAAGGTCCAGCTACTGGTGTGATTGTTGAACGTGAGCGTTTAGACAAATTCGGACGTCCTTTATTAGGTGCAACTGTAAAACCTAAACTTGGTTTATCAGGTAAAAACTATGGACGTGTTGTATACGAAGGTTTAAAAGGTGGTTTAGACTTCTTAAAAGATGACGAAAACATTAACTCTCAACCATTCATGCGTTGGCGTGAACG

>KT149186.1 Nannochloropsis oceanica strain CS-703 ribulose-1,5-bisphosphate carboxylase/oxygenase large subunit (rbcL) gene, partial cds; plastid

-AACTTAACAGCTTCAATTATCGGTAACGTATTTGGATTCAAAGCTGTAAAAGCATTACGTCTTGAAGATATGCGTATGCCTTACGCTTACTTAAAAACATTCCAAGGTCCAGCTACTGGTGTGATTGTTGAACGTGAGCGTTTAGACAAATTCGGACGTCCTTTATTAGGTGCAACTGTAAAACCTAAACTTGGTTTATCAGGTAAAAACTATGGACGTGTTGTATACGAAGGTTTAAAAGGTGGTTTAGACTTCTTAAAAGATGACGAAAACATTAACTCTCAACCATTCATGCGTTGGCGTGAACG

>KT149183.1 Nannochloropsis oceanica strain CS-700 ribulose-1,5-bisphosphate carboxylase/oxygenase large subunit (rbcL) gene, partial cds; plastid

-AACTTAACAGCTTCAATTATCGGTAACGTATTTGGATTCAAAGCTGTAAAAGCATTACGTCTTGAAGATATGCGTATGCCTTACGCTTACTTAAAAACATTCCAAGGTCCAGCTACTGGTGTGATTGTTGAACGTGAGCGTTTAGACAAATTCGGACGTCCTTTATTAGGTGCAACTGTAAAACCTAAACTTGGTTTATCAGGTAAAAACTATGGACGTGTTGTATACGAAGGTTTAAAAGGTGGTTTAGACTTCTTAAAAGATGACGAAAACATTAACTCTCAACCATTCATGCGTTGGCGTGAACG

>MW665495.1 Nannochloropsis limnetica var. globosa strain Pic 8/18 T-24d ribulose-1,5-bis-phosphate carboxylase/oxygenase, large subunit (rbcL) gene, partial cds; plastid

-AACTTAACAGCTTCAATTATCGGTAACGTATTTGGATTCAAAGCTGTAAAAGCATTACGTCTTGAAGATATGCGTATGCCTTACGCTTACTTAAAAACATTCCAAGGTCCAGCTACTGGTGTGGTTGTTGAACGTGAGCGTTTAGACAAATTCGGACGTCCTTTATTAGGTGCAACTGTAAAACCTAAACTTGGTTTATCAGGTAAAAACTATGGACGTGTTGTATACGAAGGTTTAAAAGGTGGTTTAGACTTCTTAAAAGATGACGAAAACATCAACTCTCAACCATTCATGCGTTGGCGTGAACG

>AB052288.1 Nannochloropsis salina chloroplast rbcL gene for ribulose-1,5-bisphosphate carboxylase/oxygenase large subunit, partial cds, strain:CCAP849/2

-AACTTAACGGCTTCAATCATCGGTAACGTATTTGGATTCAAAGCTGTAAAAGCGTTACGTCTTGAAGATATGCGTATGCCTTATGCTTACTTAAAAACATTCCAAGGACCAGCTACTGGTGTGATTGTTGAACGTGAGCGTTTAGACAAATTCGGACGTCCTCTATTAGGTGCAACTGTAAAACCAAAATTAGGTTTATCAGGTAAAAACTATGGGCGTGTTGTTTATGAAGGTTTAAAAGGTGGATTAGACTTCTTAAAAGATGACGAAAACATCAACTCTCAACCATTCATGCGCTGGCGCGAACG

>AB052287.1 Nannochloropsis salina chloroplast rbcL gene for ribulose-1,5-bisphosphate carboxylase/oxygenase large subunit, partial cds, strain:MBIC10063

-AACTTAACGGCTTCAATCATCGGTAACGTATTTGGATTCAAAGCTGTAAAAGCGTTACGTCTTGAAGATATGCGTATGCCTTATGCTTACTTAAAAACATTCCAAGGACCAGCTACTGGTGTGATTGTTGAACGTGAGCGTTTAGACAAATTCGGACGTCCTCTATTAGGTGCAACTGTAAAACCAAAATTAGGTTTATCAGGTAAAAACTATGGGCGTGTTGTTTATGAAGGTTTAAAAGGTGGATTAGACTTCTTAAAAGATGACGAAAACATCAACTCTCAACCATTCATGCGCTGGCGCGAACG

>AB052735.1 Nannochloropsis gaditana chloroplast gene for ribulose-1,5-bisphosphate carboxylase/oxygenase large subunit, partial cds, strain:MBIC10418

-AACTTAACGGCTTCAATCATCGGTAACGTATTTGGATTCAAAGCTGTAAAAGCGTTACGTCTTGAAGATATGCGTATGCCTTATGCTTACTTAAAAACATTCCAAGGACCAGCTACTGGTGTGATTGTTGAACGTGAGCGTTTAGACAAATTCGGACGTCCTCTATTAGGTGCAACTGTAAAACCAAAATTAGGTTTATCAGGTAAAAACTATGGGCGTGTTGTTTATGAAGGTTTAAAAGGTGGATTAGACTTCTTAAAAGATGACGAAAACATCAACTCTCAACCATTCATGCGCTGGCGCGAACG

>AB052734.1 Nannochloropsis gaditana chloroplast gene for ribulose-1,5-bisphosphate carbxylase/oxygenase large subunit, partial cds, strain:MBIC10123

-AACTTAACGGCTTCAATCATCGGTAACGTATTTGGATTCAAAGCTGTAAAAGCGTTACGTCTTGAAGATATGCGTATGCCTTATGCTTACTTAAAAACATTCCAAGGACCAGCTACTGGTGTGATTGTTGAACGTGAGCGTTTAGACAAATTCGGACGTCCTCTATTAGGTGCAACTGTAAAACCAAAATTAGGTTTATCAGGTAAAAACTATGGGCGTGTTGTTTATGAAGGTTTAAAAGGTGGATTAGACTTCTTAAAAGATGACGAAAACATCAACTCTCAACCATTCATGCGCTGGCGCGAACG

>KT149180.1 Nannochloropsis salina strain CS-446 ribulose-1,5-bisphosphate carboxylase/oxygenase large subunit (rbcL) gene, partial cds; plastid

-AACTTAACGGCTTCAATCATCGGTAACGTATTTGGATTCAAAGCTGTAAAAGCGTTACGTCTTGAAGATATGCGTATGCCTTATGCTTACTTAAAAACATTCCAAGGACCAGCTACTGGTGTGATTGTTGAACGTGAGCGTTTAGACAAATTCGGACGTCCTCTATTAGGTGCAACTGTAAAACCAAAATTAGGTTTATCAGGTAAAAACTATGGGCGTGTTGTTTATGAAGGTTTAAAAGGTGGATTAGACTTCTTAAAAGATGACGAAAACATCAACTCTCAACCATTCATGCGCTGGCGCGAACG

>KT149181.1 Nannochloropsis gaditana strain CS-448 ribulose-1,5-bisphosphate carboxylase/oxygenase large subunit (rbcL) gene, partial cds; plastid

-AACTTAACGGCTTCAATCATCGGTAACGTATTTGGATTCAAAGCTGTAAAAGCGTTACGTCTTGAAGATATGCGTATGCCTTATGCTTACTTAAAAACATTCCAAGGACCAGCTACTGGTGTGATTGTTGAACGTGAGCGTTTAGACAAATTCGGACGTCCTCTATTAGGTGCAACTGTAAAACCAAAATTAGGTTTATCAGGTAAAAACTATGGGCGTGTTGTTTATGAAGGTTTAAAAGGTGGATTAGACTTCTTAAAAGATGACGAAAACATCAACTCTCAACCATTCATGCGCTGGCGCGAACG

>AB052279.1 Nannochloropsis gaditana chloroplast rbcL gene for ribulose-1,5-bisphosphate carboxylase/oxygenase large subunit, partial cds, strain:MBIC10118

-AACTTAACGGCTTCAATCATCGGTAACGTATTTGGATTCAAAGCTGTAAAAGCGTTACGTCTTGAAGATATGCGTATGCCTTATGCTTACTTAAAAACATTCCAAGGACCAGCTACTGGTGTGATTGTTGAACGTGAGCGTTTAGACAAATTCGGACGTCCTCTATTAGGTGCAACTGTAAAACCAAAATTAGGTTTATCAGGTAAAAACTATGGGCGTGTTGTTTATGAAGGTTTAAAAGGTGGATTAGACTTCTTAAAAGATGACGAAAACATCAACTCTCAACCATTCATGCGCTGGCGCGAACG

>HQ710608.1 Monodus unipapilla culture SAG:8.83 voucher SAG 8.83 ribulose-1,5-bisphosphate carboxylase/oxygenase large subunit (rbcL) gene, partial cds; plastid

-AACTTAACAGCGTCAATCATTGGTAACGTATTTGGATTTAAAGCTGTTAAAGCATTACGTCTTGAAGATATGCGTATTCCATATGCTTACTTAAAAACATTCCAAGGTCCTGCGACTGGTATAATTGTTGAACGTGAGCGTTTAGACAAATTCGGACGACCTTTCTTAGGTGCAACTGTAAAACCTAAACTTGGTTTATCAGGGAAAAACTACGGACGTGTTGTATACGAAGGTTTAAAAGGTGGATTAGACTTCTTAAAAGATGATGAAAACATTAACTCTCAACCTTTCATGCGTTGGCGTGAACG

>MK790681.1 Monodus subterranea strain UTEX 151 ribulose-1,5-bisphosphate carboxylase/oxygenase large subunit gene, partial cds; plastid

-AACTTAACAGCGTCAATCATTGGTAACGTATTTGGATTTAAAGCTGTTAAAGCATTACGTCTTGAAGATATGCGTATTCCATATGCTTACTTAAAAACATTCCAAGGTCCTGCGACTGGTATAATTGTTGAACGTGAGCGTTTAGACAAATTCGGACGACCTTTCTTAGGTGCAACTGTAAAACCTAAACTTGGTTTATCAGGGAAAAACTACGGACGTGTTGTATACGAAGGTTTAAAAGGTGGATTAGACTTCTTAAAAGATGATGAAAACATTAACTCTCAACCTTTCATGCGTTGGCGTGAACG

>ON920853.1 Pseudotetraedriella kamillae strain SAG 2056 ribulose-1,5-bisphosphate carboxylase/oxygenase large subunit (rbcL) gene, complete cds; chloroplast

-AACTTAACAGCATCAATCATTGGTAACGTATTTGGATTCAAAGCCGTTAAAGCATTACGTCTTGAAGATATGCGTATTCCATATGCTTATTTAAAAACATTCCAAGGTCCTGCGACTGGTGTAATCGTTGAACGTGAGCGTTTAGATAAATTCGGACGACCTCTATTAGGTGCAACTGTAAAACCTAAACTTGGTTTATCAGGGAAAAACTACGGACGTGTTGTATATGAAGGTTTAAAAGGTGGTTTAGACTTCTTAAAAGATGACGAAAACATCAACTCTCAACCTTTCATGCGCTGGCGCGAACG

>MW665512.1 Pseudotetraedriella sp. strain WarPS-5 ribulose-1,5-bis-phosphate carboxylase/oxygenase, large subunit (rbcL) gene, partial cds; plastid

-AACTTAACAGCATCAATCATTGGTAACGTATTTGGTTTCAAAGCCGTTAAAGCATTACGTCTTGAAGACATGCGTATTCCATATGCTTACTTAAAGACATTCCAAGGTCCTGCGACTGGTGTAATTGTTGAACGTGAGCGTTTAGACAAATTCGGACGTCCTTTATTAGGTGCAACTGTAAAACCTAAACTTGGTTTATCAGGGAAAAACTATGGACGTGTTGTATACGAAGGTTTAAAAGGTGGTTTAGACTTCTTAAAAGATGACGAAAATATCAACTCACAACCTTTCATGCGTTGGCGTGAACG

>MW665519.1 Eustigmatophyceae sp. strain WTwin_8/9_T-2m6.8 ribulose-1,5-bis-phosphate carboxylase/oxygenase, large subunit (rbcL) gene, partial cds; plastid

-AACTTAACAGCTTCAATTATTGGTAACGTATTTGGATTCAAAGCAGTTAAAGCTTTACGTCTTGAAGATATGCGTCTTCCATATGCATACCTAAAAACATTCCAAGGGCCTGCAACAGGTGTGATCGTTGAACGTGAACGTTTAGACAAATTCGGAAGACCTCTACTTGGTGCAACTGTAAAACCAAAACTTGGTTTATCAGGTAAAAACTATGGTCGTGTTGTATTCGAAGGTTTAAAAGGTGGTTTAGACTTCTTAAAAGATGATGAAAACATCAACTCTCAACCTTTCATGCGCTGGCGCGAACG

>KT149184.1 Nannochloropsis gaditana strain CS-701 ribulose-1,5-bisphosphate carboxylase/oxygenase large subunit (rbcL) gene, partial cds; plastid

-AACTTAACGGCTTCAATCATCGGTAACGTATTTGGATTCAAAGCTGTAAAAGCGTTACGTCTTGAAGATATGCGTATGCCTTATGCTTACTTAAAAACATTCCAAGGACCAGCTACTGGTGTGATTGTTGAACGTGAGCGTTTAGACAAATTCGGACGTCCTYTATTAGGTGCAACTGTAAAACCAAAATTAGGTTTATCAGGTAAAAACTATGGGCGTGTTGTTTATGAAGGTTTAAAAGGTGGATTAGACTTCTTAAAAGATGACGAAAACATCAACTCTCAACCATTCATGCGCTGGCGCGAACG

>MN401197.1 Pseudocharaciopsis ovalis strain BCCO_30_2917 ribulose-1,5-bisphosphate carboxylase/oxygenase large subunit (rbcL) gene, partial cds; chloroplast

-AACTTAACAGCTTCAATTATCGGTAACGTATTTGGATTCAAAGCAGTAAAAGCATTACGTCTTGAAGATATGCGTCTTCCTTATGCTTACTTAAAAACTTTCCAAGGTCCTGCTACAGGTATCGTTGTAGAACGTGAACGTTTAGATAAGTTCGGAAGACCTTTCTTAGGTGCAACTGTAAAACCAAAATTAGGTTTATCTGGTAAAAACTACGGTCGTGTTGTATTTGAAGGTTTAAAAGGTGGTTTAGACTTCTTAAAAGATGACGAAAACATTAACTCTCAACCATTCATGCGTTGGCGCGAACG

>MN401190.1 Characiopsiella sp. strain ACOI 2423A ribulose-1,5-bisphosphate carboxylase/oxygenase large subunit (rbcL) gene, partial cds; chloroplast

-AACTTAACAGCTTCAATTATCGGTAACGTTTTTGGATTCAAAGCAGTTAAAGCATTACGTCTTGAAGATATGCGTATTCCTTATGCTTATTTAAAAACTTTCCAAGGTCCTGCAACAGGTATCGTTGTAGAACGTGAACGTTTAGATAAGTTTGGAAGACCTTTCTTAGGTGCAACTGTAAAACCAAAATTAGGTTTATCAGGTAAAAACTACGGTCGTGTTGTATTCGAAGGTTTAAAAGGTGGTTTAGACTTCTTAAAAGATGACGAAAACATTAACTCACAACCATTCATGCGTTGGCGCGAACG

>GQ405007.3 Eustigmatophyceae sp. WTwin 8/18 T-5d ribulose-1,5-bisphosphate carboxylase/oxygenase large subunit (rbcL) gene, partial cds; chloroplast

-AACTTAACAGCTTCAATCATTGGTAACGTTTTTGGATTCAAAGCAGTTAAAGCATTACGTCTTGAAGATATGCGTATTCCTTATGCTTATTTAAAAACTTTCCAAGGTCCTGCAACAGGTATCGTTGTAGAACGTGAACGTTTAGATAAGTTTGGAAGACCTTTCTTAGGTGCAACTGTAAAACCAAAATTAGGTTTATCTGGTAAAAACTACGGTCGTGTTGTATTTGAAGGTTTAAAAGGTGGTTTAGACTTCTTAAAAGATGACGAAAACATTAACTCTCAACCTTTCATGCGTTGGCGCGAACG

>KX354384.1 Eustigmatophyceae sp. Tow 8/18 T-6d ribulose-1,5-bisphosphate carboxylase/oxygenase large subunit (rbcL) gene, partial cds; chloroplast

-AACTTAACAGCTTCAATTATTGGTAACGTATTTGGGTTCAAAGCAGTTAAAGCTTTACGTCTTGAAGACATGCGTATTCCTTATGCATACTTAAAAACTTTCCAAGGACCTGCTACTGGTGTAGTTGTTGAACGTGAACGTTTAGACAAATTCGGAAGACCTTTACTTGGTGCAACTGTAAAACCAAAACTAGGTTTATCAGGTAAAAACTATGGACGTGTGGTATTCGAAGGTTTAAAAGGTGGTTTAGACTTCTTAAAAGACGACGAAAACATCAACTCTCAACCTTTCATGCGTTGGCGTGAACG

>MW665522.1 Eustigmatophyceae sp. strain WTwin_8/9_T-6m6.8 ribulose-1,5-bis-phosphate carboxylase/oxygenase, large subunit (rbcL) gene, partial cds; plastid

-AACTTAACAGCTTCAATTATTGGTAACGTATTTGGGTTCAAAGCAGTTAAAGCTTTACGTCTTGAAGACATGCGTATTCCTTATGCATACTTAAAAACTTTCCAAGGACCTGCTACTGGTGTAGTTGTTGAACGTGAACGTTTAGACAAATTCGGAAGACCTTTACTTGGTGCAACTGTAAAACCAAAACTAGGTTTATCAGGTAAAAACTATGGACGTGTGGTATTCGAAGGTTTAAAAGGTGGTTTAGATTTCTTAAAAGACGACGAAAACATCAACTCTCAACCTTTCATGCGTTGGCGTGAACG

>MW665505.1 Eustigmatophyceae sp. strain UP3_5/31-17m ribulose-1,5-bis-phosphate carboxylase/oxygenase, large subunit (rbcL) gene, partial cds; plastid

-AACTTAACAGCTTCAATTATTGGTAACGTATTTGGGTTCAAAGCAGTTAAAGCTTTACGTCTTGAAGACATGCGTATTCCTTATGCATACTTAAAAACTTTCCAAGGACCTGCTACTGGAGTAGTTGTTGAACGTGAACGTTTAGACAAATTCGGAAGACCTTTACTTGGTGCAACTGTAAAACCAAAACTAGGTTTATCAGGTAAAAACTATGGACGTGTGGTATTCGAAGGTTTAAAAGGTGGTTTAGACTTCTTAAAAGACGACGAAAACATCAACTCTCAACCTTTCATGCGTTGGCGTGAACG

>MW665500.1 Pseudellipsoidion sp. strain Tow_8/9_T-1w ribulose-1,5-bis-phosphate carboxylase/oxygenase, large subunit (rbcL) gene, partial cds; plastid

-AACTTAACAGCTTCAATTATTGGTAACGTTTTTGGATTCAAAGCAGTTAAAGCATTACGTCTTGAAGATATGCGTATTCCTTATGCTTACTTAAAAACTTTCCAAGGTCCTGCGACAGGTATCGTTGTAGAACGTGAACGTTTAGATAAGTTTGGAAGACCTTTCTTAGGTGCAACTGTAAAACCAAAACTAGGTTTATCAGGTAAAAACTATGGTCGTGTTGTATTTGAAGGTTTAAAAGGTGGTTTAGACTTCTTAAAAGATGACGAAAACATTAACTCTCAACCTTTCATGCGTTGGCGCGAACG

>MW665492.1 Eustigmatophyceae sp. strain Itas_8/9_T-1w ribulose-1,5-bis-phosphate carboxylase/oxygenase, large subunit (rbcL) gene, partial cds; plastid

-AACTTAACAGCTTCAATTATTGGTAACGTATTTGGGTTCAAAGCAGTTAAAGCTTTACGTCTTGAAGACATGCGTATTCCTTATGCATACTTAAAAACTTTCCAAGGACCTGCTACTGGTGTAGTTGTTGAACGTGAACGTTTAGACAAGTTCGGAAGACCTTTACTTGGTGCAACTGTAAAACCAAAACTAGGTTTATCAGGTAAAAACTATGGACGTGTGGTATTCGAAGGTTTAAAAGGTGGTTTAGACTTCTTAAAAGACGACGAAAACATCAACTCTCAACCTTTCATGCGTTGGCGTGAACG

>MW665468.1 Pseudellipsoidion sp. strain BogD_8/9_T-3m ribulose-1,5-bis-phosphate carboxylase/oxygenase, large subunit (rbcL) gene, partial cds; plastid

-AACTTAACAGCTTCAATCATTGGTAACGTTTTTGGATTCAAAGCAGTTAAAGCATTACGTCTTGAAGATATGCGTATTCCTTATGCTTACTTAAAAACTTTCCAAGGTCCTGCAACAGGTATCGTTGTAGAACGTGAACGTTTAGATAAGTTTGGAAGACCTTTCTTAGGTGCAACTGTAAAACCAAAACTAGGTTTATCAGGTAAAAACTATGGTCGTGTTGTATTTGAAGGTTTAAAAGGTGGTTTAGACTTCTTAAAAGATGACGAAAACATTAACTCTCAACCTTTCATGCGTTGGCGCGAACG

>KX354386.1 Eustigmatophyceae sp. Tow 9/21 P-2w ribulose-1,5-bisphosphate carboxylase/oxygenase large subunit (rbcL) gene, partial cds; chloroplast

-AACTTAACAGCTTCAATTATTGGTAACGTTTTTGGATTCAAAGCAGTTAAAGCATTACGTCTTGAAGATATGCGTATTCCTTATGCTTACTTAAAAACTTTCCAAGGTCCTGCGACAGGTATCGTTGTAGAACGTGAACGTTTAGATAAGTTTGGAAGACCTTTCTTAGGTGCAACTGTAAAACCAAAACTAGGTTTATCAGGTAAAAACTATGGTCGTGTTGTATTTGAAGGTTTAAAAGGTGGTTTAGACTTCTTAAAAGATGATGAAAACATCAATTCTCAACCTTTCATGCGTTGGCGCGAACG

>MN401193.1 Munda aquilonaris strain ACOI 2424B ribulose-1,5-bisphosphate carboxylase/oxygenase large subunit (rbcL) gene, partial cds; chloroplast

-AACTTAACAGCTTCAATCATTGGTAACGTTTTTGGATTCAAAGCAGTTAAAGCATTACGTCTTGAAGATATGCGTATGCCTTACGCTTATTTAAAAACTTTCCAAGGTCCTGCAACAGGTATCGTTGTAGAACGTGAACGTTTAGATAAGTTTGGAAGACCATTCTTAGGTGCAACTGTAAAACCAAAACTAGGTTTATCTGGTAAAAACTACGGTCGTGTTGTATTCGAAGGTTTAAAAGGTGGTTTAGACTTCTTAAAAGATGACGAAAACATTAACTCTCAACCATTCATGCGTTGGCGCGAACG

>KX354385.1 Eustigmatophyceae sp. Tow 8/18 T-12d ribulose-1,5-bisphosphate carboxylase/oxygenase large subunit (rbcL) gene, partial cds; chloroplast

-AACTTAACAGCTTCAATCATTGGTAACGTTTTTGGATTCAAAGCAGTTAAAGCATTACGTCTTGAAGATATGCGTATTCCTTATGCTTACTTAAAAACTTTCCAAGGTCCTGCAACAGGTATCGTTGTAGAACGTGAACGTTTAGATAAGTTTGGAAGACCTTTCTTAGGTGCAACTGTAAAACCAAAATTAGGTTTATCTGGTAAAAACTACGGTCGTGTTGTATTTGAAGGTTTAAAAGGTGGTTTAGACTTCTTAAAAGATGACGAAAACATCAACTCTCAACCTTTCATGCGTTGGCGCGAACG

>MW665516.1 Pseudellipsoidion sp. strain WTwin_8/9_T-10m6.8 ribulose-1,5-bis-phosphate carboxylase/oxygenase, large subunit (rbcL) gene, partial cds; plastid

-AACTTAACAGCTTCAATCATTGGTAACGTTTTTGGATTCAAAGCAGTTAAAGCATTACGTCTTGAAGATATGCGTATTCCTTATGCTTACTTAAAAACTTTCCAAGGTCCTGCTACAGGTATCGTTGTAGAACGTGAACGTTTAGATAAGTTTGGAAGACCTTTCTTAGGTGCAACTGTAAAACCAAAATTAGGTTTATCAGGTAAAAACTATGGTCGTGTTGTGTTCGAAGGGCTAAAAGGTGGTTTAGACTTCTTAAAAGATGACGAAAACATTAACTCTCAACCTTTCATGCGTTGGCGCGAACG

>MN401199.1 Neomonodus sp. strain CAUP Q301 ribulose-1,5-bisphosphate carboxylase/oxygenase large subunit (rbcL) gene, partial cds; chloroplast

-AACTTAACAGCTTCAATTATCGGTAACGTATTTGGATTCAAAGCAGTAAAAGCATTACGTCTTGAAGATATGCGTCTTCCTTATGCTTACTTAAAAACTTTCCAAGGTCCTGCAACAGGTATCGTTGTAGAACGTGAACGTTTAGATAAGTTCGGAAGACCTTTCTTAGGTGCAACTGTAAAACCAAAATTAGGTTTATCTGGTAAAAACTACGGTCGTGTTGTATTTGAAGGTTTAAAAGGTGGTTTAGACTTCTTAAAAGATGACGAAAACATTAACTCTCAACCATTCATGCGTTGGCGCGAACG

>MW665472.1 Eustigmataceae sp. strain BogD_8/9_T-7m ribulose-1,5-bis-phosphate carboxylase/oxygenase, large subunit (rbcL) gene, partial cds; plastid

-AACTTAACAGCTTCAATCATTGGTAACGTATTTGGTTTCAAAGCAGTTAAGGCATTACGTCTTGAAGATATGCGTATTCCTTATGCATACTTAAAAACTTTCCAAGGTCCAGCTACTGGGGTAGTTGTTGAACGTGAACGTTTAGACAAGTTTGGAAGACCTCTACTTGGAGCAACTGTAAAACCAAAACTTGGTTTATCAGGTAAAAACTACGGTCGTGTTGTATTCGAAGGTTTAAAAGGTGGTTTAGACTTCTTAAAAGATGACGAAAACATCAACTCACAACCATTCATGCGTTGGCGTGAACG

>MW665506.1 Neomonodus sp. strain UP3_5/31-1m ribulose-1,5-bis-phosphate carboxylase/oxygenase, large subunit (rbcL) gene, partial cds; plastid

-AACTTAACAGCTTCAATTATCGGTAACGTATTTGGATTCAAAGCAGTAAAAGCATTACGTCTTGAAGATATGCGTCTTCCTTATGCTTACTTAAAAACTTTCCAAGGTCCTGCAACAGGTATCGTTGTAGAACGTGAACGTTTAGATAAATTCGGAAGACCTTTCTTAGGTGCAACTGTAAAACCAAAATTAGGTTTATCTGGTAAAAACTACGGTCGTGTTGTATTTGAAGGTTTAAAAGGTGGTCTAGACTTCTTAAAAGATGACGAAAACATTAACTCTCAACCATTCATGCGTTGGCGCGAACG

>MW665469.1 Eustigmataceae sp. strain BogD_8/9_T-3m6.8 ribulose-1,5-bis-phosphate carboxylase/oxygenase, large subunit (rbcL) gene, partial cds; plastid

-AACTTAACAGCTTCAATCATTGGTAACGTATTTGGTTTCAAAGCAGTTAAGGCATTACGTCTTGAAGATATGCGTATCCCTTATGCATACTTAAAAACTTTCCAAGGTCCTGCTACTGGGGTAGTTGTTGAACGTGAACGTTTAGACAAGTTCGGAAGACCTTTACTTGGAGCAACTGTAAAACCAAAATTAGGTTTATCAGGTAAAAACTACGGTCGTGTTGTATTCGAAGGTTTAAAAGGTGGTTTAGACTTCTTAAAAGATGACGAAAACATCAACTCACAACCTTTCATGCGTTGGCGCGAACG

>MN447638.1 Pseudellipsoidion sp. strain Beav4/26 T-6w ribulose-1,5-bisphosphate carboxylase/oxygenase large subunit (rbcL) gene, partial cds; plastid

-AACTTAACAGCTTCAATCATTGGTAACGTTTTTGGATTCAAAGCAGTTAAAGCATTACGTCTTGAAGATATGCGTATTCCTTATGCTTACTTAAAAACTTTCCAAGGTCCTGCAACAGGTATCGTTGTAGAACGTGAACGTTTAGATAAGTTTGGAAGACCTTTCTTAGGTGCAACTGTAAAACCAAAATTAGGTTTATCTGGTAAAAACTACGGTCGTGTTGTATTTGAAGGTTTAAAAGGTGGTTTAGACTTCTTAAAAGATGATGAAAACATCAACTCTCAACCTTTCATGCGTTGGCGCGAACG

>MW665470.1 Eustigmataceae sp. strain BogD_8/9_T-4m ribulose-1,5-bis-phosphate carboxylase/oxygenase, large subunit (rbcL) gene, partial cds; plastid

-AACTTAACAGCTTCAATTATTGGTAACGTATTTGGGTTTAAAGCTGTTAAAGCTTTACGTCTTGAAGACATGCGTCTTCCATATGCATACCTAAAAACTTTCCAAGGACCTGCTACAGGTATAGTTGTTGAACGTGAACGTTTAGATAAGTTCGGAAGACCTTTACTTGGAGCAACTGTAAAACCAAAACTTGGTTTATCAGGTAAAAACTATGGTCGTGTTGTATTCGAAGGTTTAAAAGGTGGTTTAGACTTCTTAAAAGATGACGAAAACATCAACTCTCAACCTTTCATGCGTTGGCGTGAACG

>MW665467.1 Neomonodaceae sp. strain BogD_8/9_T-1m6.8 ribulose-1,5-bis-phosphate carboxylase/oxygenase, large subunit (rbcL) gene, partial cds; plastid

-AACTTAACAGCTTCAATTATTGGTAACGTTTTTGGATTCAAAGCAGTTAAAGCATTACGTCTTGAAGACATGCGTATTCCTTATGCTTATTTAAAAACTTTCCAAGGTCCTGCAACAGGTATCGTTGTAGAACGTGAACGTTTAGATAAATTTGGAAGACCTTTCTTAGGTGCAACTGTAAAACCAAAATTAGGTTTATCAGGTAAAAACTACGGTCGTGTTGTATTTGAAGGTTTAAAAGGTGGTTTAGACTTCTTAAAAGATGATGAAAACATCAACTCTCAAGCTTTCATGCGTTGGCGCGAACG

>MN401196.1 Neomonodus sp. strain ACOI 2437 ribulose-1,5-bisphosphate carboxylase/oxygenase large subunit (rbcL) gene, partial cds; chloroplast

-AACTTAACAGCTTCAATTATCGGTAACGTATTTGGATTCAAAGCAGTAAAAGCATTACGTCTTGAAGATATGCGTCTTCCTTATGCTTACTTAAAAACTTTCCAAGGTCCTGCAACAGGTATCGTTGTAGAACGTGAACGTTTAGATAAGTTCGGAAGACCTTTCTTAGGTGCAACTGTAAAACCAAAATTAGGTTTATCTGGTAAAAACTACGGTCGTGTTGTATTTGAAGGTTTAAAAGGTGGTTTAGACTTCTTAAAAGATGACGAAAACATTAACTCTCAACCATTCATGCGTTGGCGCGAACG

>MW665471.1 Neomonodaceae sp. strain BogD_8/9_T-6m ribulose-1,5-bis-phosphate carboxylase/oxygenase, large subunit (rbcL) gene, partial cds; plastid

-AACTTAACAGCTTCAATTATTGGTAACGTTTTTGGATTCAAAGCAGTTAAAGCATTACGTCTTGAAGATATGCGTATTCCTTATGCTTACTTAAAAACTTTCCAAGGTCCTGCAACAGGTATCGTTGTAGAACGTGAACGTTTAGATAAGTTCGGAAGACCTTTCTTAGGTGCAACTGTAAAACCAAAATTAGGTTTATCAGGTAAAAACTACGGTCGTGTTGTATTTGAAGGTTTAAAAGGTGGTTTAGACTTCTTAAAAGATGACGAAAACATTAACTCTCAACCATTCATGCGTTGGCGCGAACG

>OL703604.1:11429-12836 Emiliania huxleyi strain RCC4030 plastid, complete genome

TAACTTAACAGCATCAATTATTGGTAACATCTTCGGTTTCAAGGCTGTAAAGGCTCTTCGTCTTGAAGATATGCGTTTCCCTGTAGCTCTACTTAAGACTTACCAAGGTCCTGCAACAGGTGTTGTTGTTGAGCGTGAGCGTATGGATAAGTTCGGTCGTCCACTTCTAGGTGCAACAGTTAAGCCTAAGCTAGGTCTTTCTGGTAAGAACTATGGTCGTGTAGTATTCGAAGGTCTAAAAGGTGGTCTAGACTTCCTTAAGGATGATGAGAACATTAACTCACAGCCATTCATGCGTTACCGTGAGCG

>OL703602.1:11429-12836 Emiliania huxleyi strain RCC4002 plastid, complete genome

TAACTTAACAGCATCAATTATTGGTAACATCTTCGGTTTCAAGGCTGTAAAGGCTCTTCGTCTTGAAGATATGCGTTTCCCTGTAGCTCTACTTAAGACTTACCAAGGTCCTGCAACAGGTGTTGTTGTTGAGCGTGAGCGTATGGATAAGTTCGGTCGTCCACTTCTAGGTGCAACAGTTAAGCCTAAGCTAGGTCTTTCTGGTAAGAACTATGGTCGTGTAGTATTCGAAGGTCTAAAAGGTGGTCTAGACTTCCTTAAGGATGATGAGAACATTAACTCACAGCCATTCATGCGTTACCGTGAGCG

>OL703599.1:11429-12836 Emiliania huxleyi strain NZEH plastid, complete genome

TAACTTAACAGCATCAATTATTGGTAACATCTTCGGTTTCAAGGCTGTAAAGGCTCTTCGTCTTGAAGATATGCGTTTCCCTGTAGCTCTACTTAAGACTTACCAAGGTCCTGCAACAGGTGTTGTTGTTGAGCGTGAGCGTATGGATAAGTTCGGTCGTCCACTTCTAGGTGCAACAGTTAAGCCTAAGCTAGGTCTTTCTGGTAAGAACTATGGTCGTGTAGTATTCGAAGGTCTAAAAGGTGGTCTAGACTTCCTTAAGGATGATGAGAACATTAACTCACAGCCATTCATGCGTTACCGTGAGCG

>OL703598.1:11429-12836 Emiliania huxleyi strain M217 plastid, complete genome

TAACTTAACAGCATCAATTATTGGTAACATCTTCGGTTTCAAGGCTGTAAAGGCTCTTCGTCTTGAAGATATGCGTTTCCCTGTAGCTCTACTTAAGACTTACCAAGGTCCTGCAACAGGTGTTGTTGTTGAGCGTGAGCGTATGGATAAGTTCGGTCGTCCACTTCTAGGTGCAACAGTTAAGCCTAAGCTAGGTCTTTCTGGTAAGAACTATGGTCGTGTAGTATTCGAAGGTCTAAAAGGTGGTCTAGACTTCCTTAAGGATGATGAGAACATTAACTCACAGCCATTCATGCGTTACCGTGAGCG

>OL703596.1:11429-12836 Emiliania huxleyi strain EH2 plastid, complete genome

TAACTTAACAGCATCAATTATTGGTAACATCTTCGGTTTCAAGGCTGTAAAGGCTCTTCGTCTTGAAGATATGCGTTTCCCTGTAGCTCTACTTAAGACTTACCAAGGTCCTGCAACAGGTGTTGTTGTTGAGCGTGAGCGTATGGATAAGTTCGGTCGTCCACTTCTAGGTGCAACAGTTAAGCCTAAGCTAGGTCTTTCTGGTAAGAACTATGGTCGTGTAGTATTCGAAGGTCTAAAAGGTGGTCTAGACTTCCTTAAGGATGATGAGAACATTAACTCACAGCCATTCATGCGTTACCGTGAGCG

>OL703595.1:11429-12836 Emiliania huxleyi strain CCMP379 plastid, complete genome

TAACTTAACAGCATCAATTATTGGTAACATCTTCGGTTTCAAGGCTGTAAAGGCTCTTCGTCTTGAAGATATGCGTTTCCCTGTAGCTCTACTTAAGACTTACCAAGGTCCTGCAACAGGTGTTGTTGTTGAGCGTGAGCGTATGGATAAGTTCGGTCGTCCACTTCTAGGTGCAACAGTTAAGCCTAAGCTAGGTCTTTCTGGTAAGAACTATGGTCGTGTAGTATTCGAAGGTCTAAAAGGTGGTCTAGACTTCCTTAAGGATGATGAGAACATTAACTCACAGCCATTCATGCGTTACCGTGAGCG

>JX292160.1:1-1408 Emiliania huxleyi ribulose-1,5-bisphosphate carboxylase/oxygenase large subunit (rbcL) gene, partial cds; chloroplast

TAACTTAACAGCATCAATTATTGGTAACATCTTCGGTTTCAAGGCTGTAAAGGCTCTTCGTCTTGAAGATATGCGTTTCCCTGTAGCTCTACTTAAGACTTACCAAGGTCCTGCAACAGGTGTTGTTGTTGAGCGTGAGCGTATGGATAAGTTCGGTCGTCCACTTCTAGGTGCAACAGTTAAGCCTAAGCTAGGTCTTTCTGGTAAGAACTATGGTCGTGTAGTATTCGAAGGTCTAAAAGGTGGTCTAGACTTCCTTAAGGATGATGAGAACATTAACTCACAGCCATTCATGCGTTACCGTGAGCG

>OL703594.1:11429-12836 Emiliania huxleyi strain CCMP371 plastid, complete genome

TAACTTAACAGCATCAATTATTGGTAACATCTTCGGTTTCAAGGCTGTAAAGGCTCTTCGTCTTGAAGATATGCGTTTCCCTGTAGCTCTACTTAAGACTTACCAAGGTCCTGCAACAGGTGTTGTTGTTGAGCGTGAGCGTATGGATAAGTTCGGTCGTCCACTTCTAGGTGCAACAGTTAAGCCTAAGCTAGGTCTTTCTGGTAAGAACTATGGTCGTGTAGTATTCGAAGGTCTAAAAGGTGGTCTAGACTTCCTTAAGGATGATGAGAACATTAACTCACAGCCATTCATGCGTTACCGTGAGCG

>OL703593.1:11429-12836 Emiliania huxleyi strain AWI1516 plastid, complete genome

TAACTTAACAGCATCAATTATTGGTAACATCTTCGGTTTCAAGGCTGTAAAGGCTCTTCGTCTTGAAGATATGCGTTTCCCTGTAGCTCTACTTAAGACTTACCAAGGTCCTGCAACAGGTGTTGTTGTTGAGCGTGAGCGTATGGATAAGTTCGGTCGTCCACTTCTAGGTGCAACAGTTAAGCCTAAGCTAGGTCTTTCTGGTAAGAACTATGGTCGTGTAGTATTCGAAGGTCTAAAAGGTGGTCTAGACTTCCTTAAGGATGATGAGAACATTAACTCACAGCCATTCATGCGTTACCGTGAGCG

>OL703590.1:11429-12836 Emiliania huxleyi strain 92E plastid, complete genome

TAACTTAACAGCATCAATTATTGGTAACATCTTCGGTTTCAAGGCTGTAAAGGCTCTTCGTCTTGAAGATATGCGTTTCCCTGTAGCTCTACTTAAGACTTACCAAGGTCCTGCAACAGGTGTTGTTGTTGAGCGTGAGCGTATGGATAAGTTCGGTCGTCCACTTCTAGGTGCAACAGTTAAGCCTAAGCTAGGTCTTTCTGGTAAGAACTATGGTCGTGTAGTATTCGAAGGTCTAAAAGGTGGTCTAGACTTCCTTAAGGATGATGAGAACATTAACTCACAGCCATTCATGCGTTACCGTGAGCG

>OL322706.1:11431-12838 Emiliania huxleyi strain RCC1216 plastid, complete genome

TAACTTAACAGCATCAATTATTGGTAACATCTTCGGTTTCAAGGCTGTAAAGGCTCTTCGTCTTGAAGATATGCGTTTCCCTGTAGCTCTACTTAAGACTTACCAAGGCCCTGCAACAGGTGTTGTTGTTGAGCGTGAGCGTATGGATAAGTTCGGTCGTCCACTTCTAGGTGCAACAGTTAAGCCGAAGCTAGGTCTTTCTGGTAAGAACTATGGTCGTGTAGTATTCGAAGGTCTAAAAGGTGGTCTAGACTTCCTTAAGGATGATGAGAATATTAACTCGCAGCCATTCATGCGTTACCGTGAGCG

>OL703605.1:11428-12835 Emiliania huxleyi strain Van556 plastid, complete genome

TAACTTAACAGCATCAATTATTGGTAACATCTTCGGTTTCAAGGCTGTAAAGGCTCTTCGTCTTGAAGATATGCGTTTCCCTGTAGCTCTACTTAAGACTTACCAAGGTCCTGCAACAGGTGTTGTTGTTGAGCGTGAGCGTATGGATAAGTTCGGTCGTCCACTTCTAGGTGCAACAGTTAAGCCTAAGCTAGGTCTTTCTGGTAAGAACTATGGTCGTGTAGTATTCGAAGGTCTAAAAGGTGGTCTAGACTTCCTTAAGGATGATGAGAACATTAACTCACAGCCATTCATGCGTTACCGTGAGCG

>OL703601.1:11428-12835 Emiliania huxleyi strain RCC1253 plastid, complete genome

TAACTTAACAGCATCAATTATTGGTAACATCTTCGGTTTCAAGGCTGTAAAGGCTCTTCGTCTTGAAGATATGCGTTTCCCTGTAGCTCTACTTAAGACTTACCAAGGTCCTGCAACAGGTGTTGTTGTTGAGCGTGAGCGTATGGATAAGTTCGGTCGTCCACTTCTAGGTGCAACAGTTAAGCCTAAGCTAGGTCTTTCTGGTAAGAACTATGGTCGTGTAGTATTCGAAGGTCTAAAAGGTGGTCTAGACTTCCTTAAGGATGATGAGAACATTAACTCACAGCCATTCATGCGTTACCGTGAGCG

>OL703600.1:11428-12835 Emiliania huxleyi strain RCC174 plastid, complete genome

TAACTTAACAGCATCAATTATTGGTAACATCTTCGGTTTCAAGGCTGTAAAGGCTCTTCGTCTTGAAGATATGCGTTTCCCTGTAGCTCTACTTAAGACTTACCAAGGTCCTGCAACAGGTGTTGTTGTTGAGCGTGAGCGTATGGATAAGTTCGGTCGTCCACTTCTAGGTGCAACAGTTAAGCCTAAGCTAGGTCTTTCTGGTAAGAACTATGGTCGTGTAGTATTCGAAGGTCTAAAAGGTGGTCTAGACTTCCTTAAGGATGATGAGAACATTAACTCACAGCCATTCATGCGTTACCGTGAGCG

>OL703597.1:11428-12835 Emiliania huxleyi strain L plastid, complete genome

TAACTTAACAGCATCAATTATTGGTAACATCTTCGGTTTCAAGGCTGTAAAGGCTCTTCGTCTTGAAGATATGCGTTTCCCTGTAGCTCTACTTAAGACTTACCAAGGTCCTGCAACAGGTGTTGTTGTTGAGCGTGAGCGTATGGATAAGTTCGGTCGTCCACTTCTAGGTGCAACAGTTAAGCCTAAGCTAGGTCTTTCTGGTAAGAACTATGGTCGTGTAGTATTCGAAGGTCTAAAAGGTGGTCTAGACTTCCTTAAGGATGATGAGAACATTAACTCACAGCCATTCATGCGTTACCGTGAGCG

>OL703592.1:11428-12835 Emiliania huxleyi strain ARC30-1 plastid, complete genome

TAACTTAACAGCATCAATTATTGGTAACATCTTCGGTTTCAAGGCTGTAAAGGCTCTTCGTCTTGAAGATATGCGTTTCCCTGTAGCTCTACTTAAGACTTACCAAGGTCCTGCAACAGGTGTTGTTGTTGAGCGTGAGCGTATGGATAAGTTCGGTCGTCCACTTCTAGGTGCAACAGTTAAGCCTAAGCTAGGTCTTTCTGGTAAGAACTATGGTCGTGTAGTATTCGAAGGTCTAAAAGGTGGTCTAGACTTCCTTAAGGATGATGAGAACATTAACTCACAGCCATTCATGCGTTACCGTGAGCG

>OL703591.1:11428-12835 Emiliania huxleyi strain 92F plastid, complete genome

TAACTTAACAGCATCAATTATTGGTAACATCTTCGGTTTCAAGGCTGTAAAGGCTCTTCGTCTTGAAGATATGCGTTTCCCTGTAGCTCTACTTAAGACTTACCAAGGTCCTGCAACAGGTGTTGTTGTTGAGCGTGAGCGTATGGATAAGTTCGGTCGTCCACTTCTAGGTGCAACAGTTAAGCCTAAGCTAGGTCTTTCTGGTAAGAACTATGGTCGTGTAGTATTCGAAGGTCTAAAAGGTGGTCTAGACTTCCTTAAGGATGATGAGAACATTAACTCACAGCCATTCATGCGTTACCGTGAGCG

>OL691557.1:11428-12835 Emiliania huxleyi strain B39 plastid, partial genome

TAACTTAACAGCATCAATTATTGGTAACATCTTCGGTTTCAAGGCTGTAAAGGCTCTTCGTCTTGAAGATATGCGTTTCCCTGTAGCTCTACTTAAGACTTACCAAGGTCCTGCAACAGGTGTTGTTGTTGAGCGTGAGCGTATGGATAAGTTCGGTCGTCCACTTCTAGGTGCAACAGTTAAGCCTAAGCTAGGTCTTTCTGGTAAGAACTATGGTCGTGTAGTATTCGAAGGTCTAAAAGGTGGTCTAGACTTCCTTAAGGATGATGAGAACATTAACTCACAGCCATTCATGCGTTACCGTGAGCG

>D45845.1:1-1330 Emiliania huxleyi plastid rbcL gene for large subunit of ribulose-1,5-bisphosphate carboxylase/oxygenase (Rubisco), partial cds

TAACTTAACAGCATCAATTATTGGTAACATCTTCGGTTTCAAGGCTGTAAAGGCTCTTCGTCTTGAAGATATGCGTTTCCCTGTAGCTCTACTTAAGACTTACCAAGGTCCTGCAACAGGTGTTGTTGTTGAGCGTGAGCGTATGGATAAGTTCGGTCGTCCACTTCTAGGTGCAACAGTTAAGCCTAAGCTAGGTCTTTCTGGTAAGAACTATGGTCGTGTAGTATTCGAAGGTCTAAAAGGTGGTCTAGACTTCCTTAAGGATGATGAGAACATTAACTCACAGCCATTCATGCGTTACCGTGAGCG

>OL691556.1:11428-12835 Emiliania huxleyi culture RCC:175 strain RCC175 plastid, complete genome

TAACTTAACAGCATCAATTATTGGTAACATCTTCGGTTTCAAGGCTGTAAAGGCTCTTCGTCTTGAAGATATGCGTTTCCCTGTAGCTCTACTTAAGACTTACCAAGGTCCTGCAACAGGTGTTGTTGTTGAGCGTGAGCGTATGGATAAGTTCGGTCGTCCACTTCTAGGTGCAACAGTTAAGCCTAAGCTAGGTCTTTCTGGTAAGAACTATGGTCGTGTAGTATTCGAAGGTCTAAAAGGTGGTCTAGACTTCCTTAAGGATGATGAGAACATTAACTCACAGCCATTCATGCGTTACCGTGAGCG

>OL691558.1:11429-12836 Emiliania huxleyi strain M219 plastid, partial genome

TAACTTAACAGCATCAATTATTGGTAACATCTTCGGTTTCAAGGCTGTAAAGGCTCTTCGTCTTGAAGATATGCGTTTCCCTGTAGCTCTACTTAAGACTTACCAAGGTCCTGCAACAGGTGTTGTTGTTGAGCGTGAGCGTATGGATAAGTTCGGTCGTCCACTTCTAGGTGCAACAGTTAAGCCTAAGCTAGGTCTTTCTGGTAAGAACTATGGTCGTGTAGTATTCGAAGGTCTAAAAGGTGGTCTAGACTTCCTTAAGGATGATGAGAACATTAACTCACAGCCATTCATGCGTTACCGTGAGCG

>OL703603.2:11429-12836 Emiliania huxleyi strain RCC4028 plastid, complete genome

TAACTTAACAGCATCAATTATTGGTAACATCTTCGGTTTCAAGGCTGTAAAGGCTCTTCGTCTTGAAGATATGCGTTTCCCTGTAGCTCTACTTAAGACTTACCAAGGTCCTGCAACAGGTGTTGTTGTTGAGCGTGAGCGTATGGATAAGTTCGGTCGTCCACTTCTAGGTGCAACAGTTAAGCCTAAGCTAGGTCTTTCTGGTAAGAACTATGGTCGTGTAGTATTCGAAGGTCTAAAAGGTGGTCTAGACTTCCTTAAGGATGATGAGAACATTAACTCACAGCCATTCATGCGTTACCGTGAGCG

>JN022705.1:11429-12836 Emiliania huxleyi culture-collection CCMP:1516 plastid, complete genome

TAACTTAACAGCATCAATTATTGGTAACATCTTCGGTTTCAAGGCTGTAAAGGCTCTTCGTCTTGAAGATATGCGTTTCCCTGTAGCTCTACTTAAGACTTACCAAGGTCCTGCAACAGGTGTTGTTGTTGAGCGTGAGCGTATGGATAAGTTCGGTCGTCCACTTCTAGGTGCAACAGTTAAGCCTAAGCTAGGTCTTTCTGGTAAGAACTATGGTCGTGTAGTATTCGAAGGTCTAAAAGGTGGTCTAGACTTCCTTAAGGATGATGAGAACATTAACTCACAGCCATTCATGCGTTACCGTGAGCG

>AY741371.1:11429-12836 Emiliania huxleyi strain CCMP 373 chloroplast, complete genome

TAACTTAACAGCATCAATTATTGGTAACATCTTCGGTTTCAAGGCTGTAAAGGCTCTTCGTCTTGAAGATATGCGTTTCCCTGTAGCTCTACTTAAGACTTACCAAGGTCCTGCAACAGGTGTTGTTGTTGAGCGTGAGCGTATGGATAAGTTCGGTCGTCCACTTCTAGGTGCAACAGTTAAGCCTAAGCTAGGTCTTTCTGGTAAGAACTATGGTCGTGTAGTATTCGAAGGTCTAAAAGGTGGTCTAGACTTCCTTAAGGATGATGAGAACATTAACTCACAGCCATTCATGCGTTACCGTGAGCG

>OL703611.1:11429-12836 Gephyrocapsa parvula strain RCC4034 plastid, complete genome

TAACTTAACAGCATCAATTATTGGTAACATCTTCGGTTTCAAGGCTGTAAAGGCTCTTCGTCTTGAAGATATGCGTTTCCCTGTAGCTCTACTTAAGACTTACCAAGGTCCTGCAACAGGTGTTGTTGTTGAGCGTGAGCGTATGGATAAGTTCGGTCGTCCACTTCTAGGTGCAACAGTTAAGCCTAAGCTAGGTCTTTCTGGTAAGAACTATGGTCGTGTAGTATTCGAAGGTCTAAAAGGTGGTCTAGACTTCCTTAAGGATGATGAGAACATTAACTCACAGCCATTCATGCGTTACCGTGAGCG

>OL703607.1:11429-12836 Gephyrocapsa oceanica strain RCC3711 plastid, complete genome

TAACTTAACAGCATCAATTATTGGTAACATCTTCGGTTTCAAGGCTGTAAAGGCTCTTCGTCTTGAAGATATGCGTTTCCCTGTAGCTCTACTTAAGACTTACCAAGGTCCTGCAACAGGTGTTGTTGTTGAGCGTGAGCGTATGGATAAGTTCGGTCGTCCACTTCTAGGTGCAACAGTTAAGCCTAAGCTAGGTCTTTCTGGTAAGAACTATGGTCGTGTAGTATTCGAAGGTCTAAAAGGTGGTCTAGACTTCCTTAAGGATGATGAGAACATTAACTCACAGCCATTCATGCGTTACCGTGAGCG

>OL703606.1:11429-12836 Gephyrocapsa oceanica strain RCC1296 plastid, complete genome

TAACTTAACAGCATCAATTATTGGTAACATCTTCGGTTTCAAGGCTGTAAAGGCTCTTCGTCTTGAAGATATGCGTTTCCCTGTAGCTCTACTTAAGACTTACCAAGGTCCTGCAACAGGTGTTGTTGTTGAGCGTGAGCGTATGGATAAGTTCGGTCGTCCACTTCTAGGTGCAACAGTTAAGCCTAAGCTAGGTCTTTCTGGTAAGAACTATGGTCGTGTAGTATTCGAAGGTCTAAAAGGTGGTCTAGACTTCCTTAAGGATGATGAGAACATTAACTCACAGCCATTCATGCGTTACCGTGAGCG

>NC_063782.1:11429-12836 Gephyrocapsa oceanica strain RCC1296 plastid, complete genome

TAACTTAACAGCATCAATTATTGGTAACATCTTCGGTTTCAAGGCTGTAAAGGCTCTTCGTCTTGAAGATATGCGTTTCCCTGTAGCTCTACTTAAGACTTACCAAGGTCCTGCAACAGGTGTTGTTGTTGAGCGTGAGCGTATGGATAAGTTCGGTCGTCCACTTCTAGGTGCAACAGTTAAGCCTAAGCTAGGTCTTTCTGGTAAGAACTATGGTCGTGTAGTATTCGAAGGTCTAAAAGGTGGTCTAGACTTCCTTAAGGATGATGAGAACATTAACTCACAGCCATTCATGCGTTACCGTGAGCG

>NC_063785.1:11429-12836 Gephyrocapsa parvula strain RCC4033 plastid, complete genome

TAACTTAACAGCATCAATTATTGGTAACATCTTCGGTTTCAAGGCTGTAAAGGCTCTTCGTCTTGAAGATATGCGTTTCCCTGTAGCTCTACTTAAGACTTACCAAGGTCCTGCAACAGGTGTTGTTGTTGAGCGTGAGCGTATGGATAAGTTCGGTCGTCCACTTCTAGGTGCAACAGTTAAGCCTAAGCTAGGTCTTTCTGGTAAGAACTATGGTCGTGTAGTATTCGAAGGTCTAAAAGGTGGTCTAGACTTCCTTAAGGATGATGAGAACATTAACTCACAGCCATTCATGCGTTACCGTGAGCG

>OL703608.1:11428-12835 Gephyrocapsa muellerae strain RCC3370 plastid, complete genome

TAACTTAACAGCATCAATTATTGGTAACATCTTCGGTTTCAAGGCTGTAAAGGCTCTTCGTCTTGAAGATATGCGTTTCCCTGTAGCTCTACTTAAGACTTACCAAGGTCCTGCAACAGGTGTTGTTGTTGAGCGTGAGCGTATGGATAAGTTCGGTCGTCCACTTCTAGGTGCAACAGTTAAGCCTAAGCTAGGTCTTTCTGGTAAGAACTATGGTCGTGTAGTATTCGAAGGTCTAAAAGGTGGTCTAGACTTCCTTAAGGATGATGAGAACATTAACTCACAGCCATTCATGCGTTACCGTGAGCG

>NC_063784.1:11429-12836 Gephyrocapsa ericsonii strain RCC4032 plastid, complete genome

TAACTTAACAGCATCAATTATTGGTAACATCTTCGGTTTCAAGGCTGTAAAGGCTCTTCGTCTTGAAGATATGCGTTTCCCTGTAGCTCTACTTAAGACTTACCAAGGTCCTGCAACAGGTGTTGTTGTTGAGCGTGAACGTATGGATAAGTTCGGTCGTCCACTTCTAGGTGCAACAGTTAAGCCTAAGCTAGGTCTTTCTGGTAAGAACTATGGTCGTGTAGTATTCGAAGGTCTAAAAGGTGGTCTAGACTTCCTTAAGGATGATGAGAACATTAACTCACAGCCATTCATGCGTTACCGTGAGCG

>OL703610.1:11429-12836 Gephyrocapsa parvula strain RCC4033 plastid, complete genome

TAACTTAACAGCATCAATTATTGGTAACATCTTCGGTTTCAAGGCTGTAAAGGCTCTTCGTCTTGAAGATATGCGTTTCCCTGTAGCTCTACTTAAGACTTACCAAGGTCCTGCAACAGGTGTTGTTGTTGAGCGTGAGCGTATGGATAAGTTCGGTCGTCCACTTCTAGGTGCAACAGTTAAGCCTAAGCTAGGTCTTTCTGGTAAGAACTATGGTCGTGTAGTATTCGAAGGTCTAAAAGGTGGTCTAGACTTCCTTAAGGATGATGAGAACATTAACTCACAGCCATTCATGCGTTACCGTGAGCG

>OL703609.1:11429-12836 Gephyrocapsa ericsonii strain RCC4032 plastid, complete genome

TAACTTAACAGCATCAATTATTGGTAACATCTTCGGTTTCAAGGCTGTAAAGGCTCTTCGTCTTGAAGATATGCGTTTCCCTGTAGCTCTACTTAAGACTTACCAAGGTCCTGCAACAGGTGTTGTTGTTGAGCGTGAACGTATGGATAAGTTCGGTCGTCCACTTCTAGGTGCAACAGTTAAGCCTAAGCTAGGTCTTTCTGGTAAGAACTATGGTCGTGTAGTATTCGAAGGTCTAAAAGGTGGTCTAGACTTCCTTAAGGATGATGAGAACATTAACTCACAGCCATTCATGCGTTACCGTGAGCG

>NC_063783.1:11428-12835 Gephyrocapsa muellerae strain RCC3370 plastid, complete genome

TAACTTAACAGCATCAATTATTGGTAACATCTTCGGTTTCAAGGCTGTAAAGGCTCTTCGTCTTGAAGATATGCGTTTCCCTGTAGCTCTACTTAAGACTTACCAAGGTCCTGCAACAGGTGTTGTTGTTGAGCGTGAGCGTATGGATAAGTTCGGTCGTCCACTTCTAGGTGCAACAGTTAAGCCTAAGCTAGGTCTTTCTGGTAAGAACTATGGTCGTGTAGTATTCGAAGGTCTAAAAGGTGGTCTAGACTTCCTTAAGGATGATGAGAACATTAACTCACAGCCATTCATGCGTTACCGTGAGCG

>AB043630.1:1-1330 Gephyrocapsa oceanica plastid rbcL gene for ribulose-1,5-bisphosphate carboxylase/oxygenase large subunit, partial cds

TAACTTAACAGCATCAATTATTGGTAACATCTTCGGTTTCAAGGCTGTAAAGGCTCTTCGTCTTGAAGATATGCGTTTCCCTGTAGCTCTACTTAAGACTTACCAAGGTCCTGCAACAGGTGTTGTTGTTGAGCGTGAGCGTATGGATAAGTTCGGTCGTCCACTTCTAGGTGCAACAGTTAAGCCTAAGCTAGGTCTTTCTGGTAAGAACTATGGTCGTGTAGTATTCGAAGGTCTAAAAGGTGGTCTAGACTTCCTTAAGGATGATGAGAACATTAACTCACAGCCATTCATGCGTTACCGTGAGCG

>AB043693.1:1-1330 Isochrysis galbana plastid rbcL gene for ribulose-1,5-bisphosphate carboxylase/oxygenase large subunit, partial cds

TAACCTAACTGCATCAATCATTGGTAACATCTTCGGTTTCAAGGCTGTTAAGGCTCTTCGTCTTGAAGATATGCGTATGCCAGTTGCTCTATTAAAGACTTACCAAGGTCCAGCAACAGGTCTTGTTGTTGAGCGTGAGCGTCTAGATAAGTTCGGTCGTCCACTTCTAGGTGCAACAGTTAAGCCTAAATTAGGTCTTTCTGGTAAGAACTATGGTCGTGTAGTATTCGAAGGTCTTAAAGGTGGTCTAGACTTCCTTAAGGATGATGAGAACATTAACTCACAACCATTTATGCGTTACCGTGAGCG

>AB043629.1:1-1330 Umbilicosphaera sibogae var. foliosa plastid rbcL gene for ribulose-1,5-bisphosphate carboxylase/oxygenase large subunit, partial cds

TAACTTAACAGCGTCAATTATTGGTAACATTTTCGGTTTCAAGGCTGTTAAGGCTCTAAGACTTGAAGATATGCGTATGCCAGTAGCACTTCTTAAGACGTACCAAGGTCCAGCAACAGGTCTTATCGTAGAGCGTGAGCGTCTAGATAAGTTTGGTCGTCCACTTTTAGGTGCAACAGTTAAGCCTAAGCTAGGTCTTTCTGGTAAGAACTATGGTCGTGTAGTATTCGAAGGTCTTAAAGGTGGTCTAGACTTCCTTAAGGATGATGAGAACATTAACTCACAACCATTCATGCGTTACAGAGAGCG

>D45843.1:1-1330 Umbilicosphaera sibogae var. foliosa plastid rbcL gene for large subunit of ribulose-1,5-bishosphate carboxylase/oxygenase (Rubisco), partial cds

TAACTTAACAGCGTCAATTATTGGTAACATTTTCGGTTTCAAGGCTGTTAAGGCTCTAAGACTTGAAGATATGCGTATGCCAGTAGCACTTCTTAAGACGTACCAAGGTCCAGCAACAGGTCTTATCGTAGAGCGTGAGCGTCTAGATAAGTTTGGTCGTCCACTTTTAGGTGCAACAGTTAAGCCTAAGCTAGGTCTTTCTGGTAAGAACTATGGTCGTGTAGTATTCGAAGGTCTTAAAGGTGGTCTAGACTTCCTTAAGGATGATGAGAACATTAACTCACAACCATTCATGCGTTACAGAGAGCG

>D11140.1:370-1777 Pleurochrysis carterae chloroplast genes for ribulose-1,5-bisphosphate carboxylase/oxygenase large and small subunits, complete cds

TAACCTAACTGCGTCTATTATTGGTAACATTTTCGGTTTCAAGGCTGTAAAAGCTCTAAGACTTGAAGATATGCGTATGCCATACGCACTACTAAAGACTTACCAAGGTCCAGCTACTGGCCTAATCGTAGAGCGAGAGCGTCTAGATAAATTCGGTCGCCCACTACTAGGTGCAACTGTTAAGCCTAAGCTAGGTCTTTCAGGTAAAAACTACGGTCGTGTAGTATTCGAAGGTCTTAAAGGTGGTCTAGATTTCCTTAAGGATGATGAGAACATTAACTCACAGCCTTTCATGCGTTACCGCGAGCG

>AB043691.1:1-1330 Umbilicosphaera sibogae plastid rbcL gene for ribulose-1,5-bisphosphate carboxylase/oxygenase large subunit, partial cds

TAACTTAACAGCTTCAATTATTGGTAACATTTTCGGTTTCAAGGCTGTTAAAGCTCTAAGACTTGAAGATATGCGTATGCCAGTTGCACTTCTTAAGACGTACCAAGGTCCAGCAACAGGTCTTATTGTAGAGCGTGAGCGTCTAGATAAGTTTGGTCGTCCACTTTTAGGTGCAACAGTTAAGCCTAAGCTAGGTCTTTCTGGTAAGAACTATGGTCGTGTAGTATTCGAAGGTCTTAAAGGTGGTTTAGACTTCCTTAAGGATGATGAGAACATTAACTCACAACCATTCATGCGTTACAGAGAGCG

>AB043628.1:1-1330 Calyptrosphaera sphaeroidea plastid rbcL gene for ribulose-1,5-bisphosphate carboxylase/oxygenase large subunit, partial cds

TAACCTAACTGCTTCAATTATCGGTAACATTTTCGGTTTCAAGGCTGTTAAGGCTCTAAGACTTGAAGATATGCGTATGCCATACGCTCTACTTAAGACTTACCAAGGTCCTGCAACTGGTCTAGTTGTAGAGCGTGAGCGTCTAGACAAATTTGGTCGTCCACTTCTAGGTGCTACAGTTAAACCTAAGCTAGGTCTTTCTGGTAAGAACTATGGTCGTGTAGTATTCGAAGGTCTTAAAGGTGGTCTAGACTTCCTTAAGGATGATGAGAATATTAACTCACAGCCATTCATGCGTTACAGAGAGCG

>D45842.1:1-1330 Calyptrosphaera sphaeroidea plastid rbcL gene for large subunit of ribulose-1,5-bisphosphate carboxylase/oxgenase (Rubisco), partial cds

TAACCTAACTGCTTCAATTATCGGTAACATTTTCGGTTTCAAGGCTGTTAAGGCTCTAAGACTTGAAGATATGCGTATGCCATACGCTCTACTTAAGACTTACCAAGGTCCTGCAACTGGTCTAGTTGTAGAGCGTGAGCGTCTAGACAAATTTGGTCGTCCACTTCTAGGTGCTACAGTTAAACCTAAGCTAGGTCTTTCTGGTAAGAACTATGGTCGTGTAGTATTCGAAGGTCTTAAAGGTGGTCTAGACTTCCTTAAGGATGATGAGAATATTAACTCACAGCCATTCATGCGTTACAGAGAGCG

>AB043632.1:1-1328 Chrysochromulina hirta plastid rbcL gene for ribulose-1,5-bisphosphate carboxylase/oxygenase large subunit, partial cds

TAACCTAACTGCATCTATTATTGGTAACATCTTCGGTTTTAAGGCTGTAAAGGCACTTAGACTAGAAGATATGCGTTTCCCAGTTGCACTACTTAAGACTTACCAAGGTCCTGCGACTGGTCTAATTGTAGAGCGTGAGCGTATGGATAAGTTTGGTCGTCCACTACTAGGTGCTACAGTTAAGCCTAAGCTAGGTCTTTCTGGTAAGAACTATGGTCGTGTAGTATTCGAAGGTCTTAAAGGTGGTCTTGATTTCCTTAAGGATGATGAGAACATTAACTCTCAACCATTCATGCGTTACAGAGAGCG

>D45846.1:1-1328 Chrysochromulina hirta plastid rbcL gene for large subunit of ribulose-1,5-bisphosphate carboxylase/oxygenase (Rubisco), partial cds

TAACCTAACTGCATCTATTATTGGTAACATCTTCGGTTTTAAGGCTGTAAAGGCACTTAGACTAGAAGATATGCGTTTCCCAGTTGCACTACTTAAGACTTACCAAGGTCCTGCGACTGGTCTAATTGTAGAGCGTGAGCGTATGGATAAGTTTGGTCGTCCACTACTAGGTGCTACAGTTAAGCCTAAGCTAGGTCTTTCTGGTAAGAACTATGGTCGTGTAGTATTCGAAGGTCTTAAAGGTGGTCTTGATTTCCTTAAGGATGATGAGAACATTAACTCTCAACCATTCATGCGTTACAGAGAGCG

>HQ656827.1:1-1330 Chrysotila lamellosa culture-collection PCC:465 ribulose-1,5-bisphosphate carboxylase/oxygenase large subunit (rbcL) gene, partial cds; chloroplast

TAACTTAACTGCGTCGATTATTGGTAACATTTTTGGTTTCAAGGCTGTTAAAGCTCTTCGTCTTGAAGATATGCGTATGCCGGTTGCTCTATTAAAGACATACCAAGGTCCTGCAACAGGTCTTGTTGTTGAGCGTGAACGTCTAGATAAGTTCGGTCGTCCACTTTTAGGTGCAACAGTTAAGCCGAAGCTAGGTCTTTCAGGTAAAAACTATGGTCGTGTAGTATTCGAAGGTCTTAAAGGTGGTTTAGATTTCCTAAAGGATGAGGAAAACATTAACTCACAGCCATTTATGCGTTACCGCGAGCG

>AB043696.1:1-1328 Dicrateria rotunda plastid rbcL gene for ribulose-1,5-bisphosphate carboxylase/oxygenase large subunit, partial cds

AAACTTAACTGCGTCTATTATCGGAAACATTTTCGGTTTCAAGGCTGTTAAAGCTCTAAGACTTGAAGATATGCGTTTCCCATATGCGCTACTTAAGACTTTCCAAGGTCCTGCAACTGGTCTAGTTGTAGAGCGTGAGCGTATGGATAAGTTTGGTCGTCCTCTTCTAGGTGCTACTGTAAAGCCTAAGTTAGGTCTTTCAGGTAAGAACTACGGTCGTGTAGTATTCGAAGGTCTTAAAGGTGGTCTTGACTTCCTAAAGGATGACGAGAACATTAACTCGCAGCCTTTCATGCGTTACAGAGAGCG

>AB043695.1:1-1328 Chrysochromulina alifera plastid rbcL gene for ribulose-1,5-bisphosphate carboxylase/oxygenase large subunit, partial cds

TAACCTAACTGCATCAATTATCGGTAACATCTTCGGTTTCAAAGCTGTTAAGGCTCTTCGTCTTGAAGATATGCGTTTCCCTTATGCACTTCTTAAGACTTTCCAAGGTCCTGCGACTGGTCTAGTTGTAGAGCGTGAGCGTATGGATAAGTTTGGTCGTCCACTTCTAGGTGCTACTGTAAAGCCTAAGCTAGGTCTTTCAGGTAAGAACTATGGTCGTGTAGTATTCGAAGGTCTTAAAGGTGGTCTAGACTTCCTTAAGGATGATGAGAACATTAACTCACAGCCATTCATGCGTTACAGAGAGCG

>HQ656833.1:1-1330 Coccolithus pelagicus culture-collection PCC:182g ribulose-1,5-bisphosphate carboxylase/oxygenase large subunit (rbcL) gene, partial cds; chloroplast

TAACCTAACTGCTTCAATTATTGGTAACATTTTTGGTTTCAAGGCTGTTAAGTCTCTAAGACTTGAAGATATGCGTATGCCATACGCACTTCTTAAAACTTACCAAGGTCCTGCAACAGGTCTAGTAGTAGAGCGTGAGCGACTGGATAAGTTCGGTCGTCCACTTCTAGGTGCTACAGTTAAGCCTAAGCTAGGTCTTTCTGGTAAGAACTATGGTCGTGTAGTATTCGAAGGTCTAAAAGGTGGTCTAGACTTCCTTAAAGATGATGAGAACATTAACTCACAACCGTTCATGCGTTACAGAGAGCG

>HQ656831.1:1-1330 Pleurochrysis placolithoides culture-collection PCC:604 ribulose-1,5-bisphosphate carboxylase/oxygenase large subunit (rbcL) gene, partial cds; chloroplast

TAACCTAACTGCGTCTATCATTGGTAACATTTTCGGTTTCAAGGCTGTAAAAGCTCTAAGACTTGAAGATATGCGTATGCCATACGCACTACTAAAGACTTACCAAGGTCCAGCTACTGGTCTAGTTGTAGAGCGTGAGCGTCTAGATAAGTTCGGTCGTCCACTATTAGGTGCAACTGTTAAGCCTAAGCTAGGTCTTTCAGGTAAGAACTACGGTCGTGTAGTATTCGAGGGTCTTAAAGGTGGTCTAGACTTCCTTAAGGATGATGAGAACATTAACTCACAACCTTTCATGCGTTACCGTGAGCG

>HQ656829.1:1-1330 Isochrysis galbana culture-collection PCC:8 ribulose-1,5-bisphosphate carboxylase/oxygenase large subunit (rbcL) gene, partial cds; chloroplast

TAACCTAACTGCATCAATTATTGGTAACATTTTTGGTTTCAAGGCTGTTAAAGCTCTTCGTCTTGAAGATATGCGTATGCCAGTTGCTTTATTAAAAACATACCAAGGTCCTGCAACAGGTCTTGTTGTTGAGCGTGAGCGTCTAGATAAGTTTGGTCGTCCACTTTTAGGTGCAACGGTTAAACCTAAGCTAGGTCTTTCTGGTAAGAACTATGGTCGTGTAGTATTCGAAGGTCTTAAAGGTGGTTTAGACTTCCTAAAAGATGACGAGAACATTAACTCACAACCATTCATGCGTTACCGTGAGCG

>AB043690.1:1-1330 Calcidiscus leptoporus plastid rbcL gene for ribulose-1,5-bisphosphate carboxylase/oxygenase large subunit, partial cds

TAACTTAACAGCGTCAATTATTGGTAACATCTTCGGTTTCAAGGCTGTTAAGTCTCTAAGACTTGAAGATATGCGTATGCCATATGCACTTCTTAAAACGTACCAAGGTCCTGCATGTGGACTTATTGTAGAGCGTGAGCGTCTAGACAAGTTTGGTCGTCCATTACTAGGTGCAACAGTTAAGCCAAAGCTAGGTCTTTCTGGTAAGAACTATGGTCGTGTAGTATTCGAAGGTCTTAAAGGTGGTCTAGACTTCCTTAAGGATGATGAGAACATTAACTCACAACCATTTATGCGTTACAGAGAGCG

>AB043689.1:1-1330 Cruciplacolithus neohelis plastid rbcL gene for ribulose-1,5-bisphosphate carboxylase/oxygenase large subunit, partial cds

TAACCTAACTGCTTCAATTATCGGTAACATTTTCGGTTTCAAGGCTGTTAAGGCTCTAAGACTTGAAGATATGCGTTTACCGTACGCACTGCTTAAGACTTACCAAGGTCCTGCAACTGGTATGGTTGTAGAGCGTGAGCGTCTAGATAAGTTTGGTCGTCCACTTTTAGGTGCTACAGTTAAGCCTAAATTAGGTCTTTCTGGTAAGAACTATGGTCGTGTAGTATTTGAAGGTCTTAAAGGTGGTCTAGACTTCCTTAAGGATGATGAGAACATTAACTCACAGCCGTTCATGCGTTACAGAGAGCG

>AB043692.1:1-1328 Helicosphaera carteri plastid rbcL gene for ribulose-1,5-bisphosphate carboxylase/oxygenase large subunit, partial cds

TAACCTTACTGCATCGATTATTGGTAACATTTTCGGTTTCAAGGCAGTTAAAGCTCTAAGACTTGAGGATATGCGTATGCCAGTTGCTCTATTAAAGACTTACCAAGGTCCAGCTACAGGTCTAGTTGTTGAGCGTGAGCGTCTAGACAAGTTTGGTCGTCCACTACTAGGTGCAACAGTTAAGCCTAAGCTAGGTCTTTCAGGAAAGAACTATGGTCGTGTAGTATTCGAAGGTCTAAAAGGAGGTCTAGATTTCCTTAAGGATGATGAGAACATCAACTCACAACCATTCATGCGTTACAGAGAGCG

>AB043694.1:1-1330 Chrysochromulina parva plastid rbcL gene for ribulose-1,5-bisphosphate carboxylase/oxygenase large subunit, partial cds

TAACCTAACTGCTTCGATCATTGGTAACATTTTTGGTTTCAAAGCCGTTAAAGCACTTCGTCTTGAAGATATGCGTTTCCCTTACGCACTACTTAAGACTTACCAAGGCCCTGCAACTGGTCTAGTTGTAGAGCGTGAGCGTATGGATAAGTTTGGTCGTCCTCTTCTAGGTGCTACAGTAAAGCCTAAGCTAGGTCTTTCTGGTAAGAACTACGGTCGTGTAGTATTCGAGGGTCTTAAAGGTGGTCTAGATTTCCTTAAGGATGATGAGAACATTAACTCACAACCATTCATGCGTTACAGAGAGCG

>AB043688.1:1-1330 Pleurochrysis haptonemofera plastid rbcL gene for ribulose-1,5-bisphosphate carboxylase/oxygenase large subunit, partial cds

TAACCTAACTGCGTCTATTATTGGTAACATCTTCGGCTTCAAAGCTGTAAAAGCTCTAAGACTTGAAGATATGCGTATGCCATACGCGCTACTAAAAACTTACCAAGGTCCAGCTACTGGTCTAATCGTAGAGCGTGAGCGTCTAGATAAGTTCGGTCGTCCACTACTAGGTGCAACTGTTAAGCCTAAGCTAGGTCTTTCAGGTAAGAACTACGGTCGTGTAGTATTCGAAGGTCTTAAAGGTGGTCTAGATTTCCTTAAGGACGACGAGAACATTAACTCACAACCTTTCATGCGTTACCGCGAGCG

>AB043697.1:1-1328 Chrysochromulina sp. TKB8936 plastid rbcL gene for ribulose-1,5-bisphosphate carboxylase/oxygenase large subunit, partial cds

TAACCTAACTGCATCTATTATTGGTAACATCTTCGGTTTCAAAGCTGTAAAAGCTCTTAGACTAGAAGATATGCGTTTCCCTGTTGCACTATTAAAGACTTACCAAGGTCCTGCTTGTGGTCTAATCGTAGAGCGTGAGCGTATGGATAAGTTTGGTCGTCCTCTTCTAGGTGCGACTGTAAAGCCTAAGCTTGGTCTTTCTGGTAAGAACTATGGTCGTGTAGTATTCGAAGGTCTTAAGGGTGGTCTTGACTTCCTAAAGGATGATGAGAACATTAACTCTCAGCCATTCATGCGTTACAGAGAGCG

>KC900889.1:99447-100852 Phaeocystis globosa strain Pg-G(A) chloroplast, complete genome

TAACCTAACTGCTTCAATTATTGGTAACATCTTCGGATTCAAAGCTGTTAAAGCTCTAAGATTAGAAGATATGAGAATGCCAGTTGCACTATTAAAGACTTACCAAGGTCCTGCTTGTGGTCTAATTGTAGAGCGTGAGAGAATGGATAAGTTTGGTCGTCCTCTACTTGGTGCTACAGTTAAGCCTAAGCTAGGTCTTTCTGGTAAGAACTACGGTCGTGTAGTATTCGAAGGTCTTAAAGGTGGTCTTGATTTCCTTAAGGATGATGAGAACATTAACTCACAACCGTTCATGCGTTACAGAGAGCG

>JN117275.2:96916-98321 Phaeocystis antarctica strain CCMP1374 plastid, complete genome

TAACCTAACTGCTTCAATTATTGGTAACATTTTCGGATTCAAAGCTGTTAAAGCTCTAAGATTAGAAGATATGAGAATGCCAGTTGCACTATTAAAGACTTACCAAGGTCCTGCTTGTGGTCTAATTGTAGAGCGTGAGAGAATGGATAAGTTTGGTCGTCCTCTACTTGGTGCTACAGTTAAGCCTAAATTAGGTCTTTCTGGTAAGAACTATGGTCGTGTAGTATTCGAAGGTCTTAAAGGTGGTCTTGATTTCCTTAAGGATGATGAGAACATTAACTCACAACCATTCATGCGTTACAGAGAGCG

>MT471334.1:10-1415 Phaeocystis globosa strain CNS00093 chloroplast, complete genome

TAACCTAACTGCTTCAATTATTGGTAACATCTTCGGATTCAAAGCTGTTAAAGCTCTAAGATTAGAAGATATGAGAATGCCAGTTGCACTACTAAAGACTTACCAAGGTCCTGCTTGTGGTCTAATTGTAGAGCGTGAGAGAATGGACAAGTTTGGTCGTCCTCTACTTGGTGCTACAGTTAAGCCTAAGCTAGGTCTTTCTGGTAAGAACTATGGTCGTGTAGTATTCGAAGGTCTTAAAGGTGGTCTTGATTTCCTTAAGGATGATGAGAACATTAACTCACAACCGTTCATGCGTTACAGAGAGCG

>MT471333.1:10-1415 Phaeocystis globosa strain CNS00087 chloroplast, complete genome

TAACCTAACTGCTTCAATTATTGGTAACATCTTCGGATTCAAAGCTGTTAAAGCTCTAAGATTAGAAGATATGAGAATGCCAGTTGCACTACTAAAGACTTACCAAGGTCCTGCTTGTGGTCTAATTGTAGAGCGTGAGAGAATGGACAAGTTTGGTCGTCCTCTACTTGGTGCTACAGTTAAGCCTAAGCTAGGTCTTTCTGGTAAGAACTATGGTCGTGTAGTATTCGAAGGTCTTAAAGGTGGTCTTGATTTCCTTAAGGATGATGAGAACATTAACTCACAACCGTTCATGCGTTACAGAGAGCG

>MT471332.1:10-1415 Phaeocystis globosa strain CNS00080 chloroplast, complete genome

TAACCTAACTGCTTCAATTATTGGTAACATCTTCGGATTCAAAGCTGTTAAAGCTCTAAGATTAGAAGATATGAGAATGCCAGTTGCACTACTAAAGACTTACCAAGGTCCTGCTTGTGGTCTAATTGTAGAGCGTGAGAGAATGGACAAGTTTGGTCGTCCTCTACTTGGTGCTACAGTTAAGCCTAAGCTAGGTCTTTCTGGTAAGAACTATGGTCGTGTAGTATTCGAAGGTCTTAAAGGTGGTCTTGATTTCCTTAAGGATGATGAGAACATTAACTCACAACCGTTCATGCGTTACAGAGAGCG

>MT471331.1:10-1415 Phaeocystis globosa strain CNS00079 chloroplast, complete genome

TAACCTAACTGCTTCAATTATTGGTAACATCTTCGGATTCAAAGCTGTTAAAGCTCTAAGATTAGAAGATATGAGAATGCCAGTTGCACTACTAAAGACTTACCAAGGTCCTGCTTGTGGTCTAATTGTAGAGCGTGAGAGAATGGACAAGTTTGGTCGTCCTCTACTTGGTGCTACAGTTAAGCCTAAGCTAGGTCTTTCTGGTAAGAACTATGGTCGTGTAGTATTCGAAGGTCTTAAAGGTGGTCTTGATTTCCTTAAGGATGATGAGAACATTAACTCACAACCGTTCATGCGTTACAGAGAGCG

>MN927526.1:1-1391 Phaeocystis globosa isolate CNS00088 ribulose-1,5-bisphosphate carboxylase/oxygenase large subunit (rbcL) gene, partial cds; mitochondrial

TAACCTAACTGCTTCAATTATTGGTAACATCTTCGGATTCAAAGCTGTTAAAGCTCTAAGATTAGAAGATATGAGAATGCCAGTTGCACTACTAAAGACTTACCAAGGTCCTGCTTGTGGTCTAATTGTAGAGCGTGAGAGAATGGACAAGTTTGGTCGTCCTCTACTTGGTGCTACAGTTAAGCCTAAGCTAGGTCTTTCTGGTAAGAACTATGGTCGTGTAGTATTCGAAGGTCTTAAAGGTGGTCTTGATTTCCTTAAGGATGATGAGAACATTAACTCACAACCGTTCATGCGTTACAGAGAGCG

>MN927525.1:1-1391 Phaeocystis globosa isolate CNS00087 ribulose-1,5-bisphosphate carboxylase/oxygenase large subunit (rbcL) gene, partial cds; mitochondrial

TAACCTAACTGCTTCAATTATTGGTAACATCTTCGGATTCAAAGCTGTTAAAGCTCTAAGATTAGAAGATATGAGAATGCCAGTTGCACTACTAAAGACTTACCAAGGTCCTGCTTGTGGTCTAATTGTAGAGCGTGAGAGAATGGACAAGTTTGGTCGTCCTCTACTTGGTGCTACAGTTAAGCCTAAGCTAGGTCTTTCTGGTAAGAACTATGGTCGTGTAGTATTCGAAGGTCTTAAAGGTGGTCTTGATTTCCTTAAGGATGATGAGAACATTAACTCACAACCGTTCATGCGTTACAGAGAGCG

>MN927524.1:1-1391 Phaeocystis globosa isolate CNS00086 ribulose-1,5-bisphosphate carboxylase/oxygenase large subunit (rbcL) gene, partial cds; mitochondrial

TAACCTAACTGCTTCAATTATTGGTAACATCTTCGGATTCAAAGCTGTTAAAGCTCTAAGATTAGAAGATATGAGAATGCCAGTTGCACTACTAAAGACTTACCAAGGTCCTGCTTGTGGTCTAATTGTAGAGCGTGAGAGAATGGACAAGTTTGGTCGTCCTCTACTTGGTGCTACAGTTAAGCCTAAGCTAGGTCTTTCTGGTAAGAACTATGGTCGTGTAGTATTCGAAGGTCTTAAAGGTGGTCTTGATTTCCTTAAGGATGATGAGAACATTAACTCACAACCGTTCATGCGTTACAGAGAGCG

>MN927523.1:1-1391 Phaeocystis globosa isolate CNS00085 ribulose-1,5-bisphosphate carboxylase/oxygenase large subunit (rbcL) gene, partial cds; mitochondrial

TAACCTAACTGCTTCAATTATTGGTAACATCTTCGGATTCAAAGCTGTTAAAGCTCTAAGATTAGAAGATATGAGAATGCCAGTTGCACTACTAAAGACTTACCAAGGTCCTGCTTGTGGTCTAATTGTAGAGCGTGAGAGAATGGACAAGTTTGGTCGTCCTCTACTTGGTGCTACAGTTAAGCCTAAGCTAGGTCTTTCTGGTAAGAACTATGGTCGTGTAGTATTCGAAGGTCTTAAAGGTGGTCTTGATTTCCTTAAGGATGATGAGAACATTAACTCACAACCGTTCATGCGTTACAGAGAGCG

>MN927522.1:1-1391 Phaeocystis globosa isolate CNS00084 ribulose-1,5-bisphosphate carboxylase/oxygenase large subunit (rbcL) gene, partial cds; mitochondrial

TAACCTAACTGCTTCAATTATTGGTAACATCTTCGGATTCAAAGCTGTTAAAGCTCTAAGATTAGAAGATATGAGAATGCCAGTTGCACTACTAAAGACTTACCAAGGTCCTGCTTGTGGTCTAATTGTAGAGCGTGAGAGAATGGACAAGTTTGGTCGTCCTCTACTTGGTGCTACAGTTAAGCCTAAGCTAGGTCTTTCTGGTAAGAACTATGGTCGTGTAGTATTCGAAGGTCTTAAAGGTGGTCTTGATTTCCTTAAGGATGATGAGAACATTAACTCACAACCGTTCATGCGTTACAGAGAGCG

>MN927521.1:1-1391 Phaeocystis globosa isolate CNS00083 ribulose-1,5-bisphosphate carboxylase/oxygenase large subunit (rbcL) gene, partial cds; mitochondrial

TAACCTAACTGCTTCAATTATTGGTAACATCTTCGGATTCAAAGCTGTTAAAGCTCTAAGATTAGAAGATATGAGAATGCCAGTTGCACTACTAAAGACTTACCAAGGTCCTGCTTGTGGTCTAATTGTAGAGCGTGAGAGAATGGACAAGTTTGGTCGTCCTCTACTTGGTGCTACAGTTAAGCCTAAGCTAGGTCTTTCTGGTAAGAACTATGGTCGTGTAGTATTCGAAGGTCTTAAAGGTGGTCTTGATTTCCTTAAGGATGATGAGAACATTAACTCACAACCGTTCATGCGTTACAGAGAGCG

>MN927520.1:1-1391 Phaeocystis globosa isolate CNS00082 ribulose-1,5-bisphosphate carboxylase/oxygenase large subunit (rbcL) gene, partial cds; mitochondrial

TAACCTAACTGCTTCAATTATTGGTAACATCTTCGGATTCAAAGCTGTTAAAGCTCTAAGATTAGAAGATATGAGAATGCCAGTTGCACTACTAAAGACTTACCAAGGTCCTGCTTGTGGTCTAATTGTAGAGCGTGAGAGAATGGACAAGTTTGGTCGTCCTCTACTTGGTGCTACAGTTAAGCCTAAGCTAGGTCTTTCTGGTAAGAACTATGGTCGTGTAGTATTCGAAGGTCTTAAAGGTGGTCTTGATTTCCTTAAGGATGATGAGAACATTAACTCACAACCGTTCATGCGTTACAGAGAGCG

>MN927519.1:1-1391 Phaeocystis globosa isolate CNS00081 ribulose-1,5-bisphosphate carboxylase/oxygenase large subunit (rbcL) gene, partial cds; mitochondrial

TAACCTAACTGCTTCAATTATTGGTAACATCTTCGGATTCAAAGCTGTTAAAGCTCTAAGATTAGAAGATATGAGAATGCCAGTTGCACTACTAAAGACTTACCAAGGTCCTGCTTGTGGTCTAATTGTAGAGCGTGAGAGAATGGACAAGTTTGGTCGTCCTCTACTTGGTGCTACAGTTAAGCCTAAGCTAGGTCTTTCTGGTAAGAACTATGGTCGTGTAGTATTCGAAGGTCTTAAAGGTGGTCTTGATTTCCTTAAGGATGATGAGAACATTAACTCACAACCGTTCATGCGTTACAGAGAGCG

>MN927518.1:1-1391 Phaeocystis globosa isolate CNS00080 ribulose-1,5-bisphosphate carboxylase/oxygenase large subunit (rbcL) gene, partial cds; mitochondrial

TAACCTAACTGCTTCAATTATTGGTAACATCTTCGGATTCAAAGCTGTTAAAGCTCTAAGATTAGAAGATATGAGAATGCCAGTTGCACTACTAAAGACTTACCAAGGTCCTGCTTGTGGTCTAATTGTAGAGCGTGAGAGAATGGACAAGTTTGGTCGTCCTCTACTTGGTGCTACAGTTAAGCCTAAGCTAGGTCTTTCTGGTAAGAACTATGGTCGTGTAGTATTCGAAGGTCTTAAAGGTGGTCTTGATTTCCTTAAGGATGATGAGAACATTAACTCACAACCGTTCATGCGTTACAGAGAGCG

>MN927517.1:1-1391 Phaeocystis globosa isolate CNS00079 ribulose-1,5-bisphosphate carboxylase/oxygenase large subunit (rbcL) gene, partial cds; mitochondrial

TAACCTAACTGCTTCAATTATTGGTAACATCTTCGGATTCAAAGCTGTTAAAGCTCTAAGATTAGAAGATATGAGAATGCCAGTTGCACTACTAAAGACTTACCAAGGTCCTGCTTGTGGTCTAATTGTAGAGCGTGAGAGAATGGACAAGTTTGGTCGTCCTCTACTTGGTGCTACAGTTAAGCCTAAGCTAGGTCTTTCTGGTAAGAACTATGGTCGTGTAGTATTCGAAGGTCTTAAAGGTGGTCTTGATTTCCTTAAGGATGATGAGAACATTAACTCACAACCGTTCATGCGTTACAGAGAGCG

>MN927512.1:1-1391 Phaeocystis globosa isolate CNS00073 ribulose-1,5-bisphosphate carboxylase/oxygenase large subunit (rbcL) gene, partial cds; mitochondrial

TAACCTAACTGCTTCAATTATTGGTAACATCTTCGGATTCAAAGCTGTTAAAGCTCTAAGATTAGAAGATATGAGAATGCCAGTTGCACTACTAAAGACTTACCAAGGTCCTGCTTGTGGTCTAATTGTAGAGCGTGAGAGAATGGACAAGTTTGGTCGTCCTCTACTTGGTGCTACAGTTAAGCCTAAGCTAGGTCTTTCTGGTAAGAACTATGGTCGTGTAGTATTCGAAGGTCTTAAAGGTGGTCTTGATTTCCTTAAGGATGATGAGAACATTAACTCACAACCGTTCATGCGTTACAGAGAGCG

>MN927511.1:1-1391 Phaeocystis globosa isolate CNS00072 ribulose-1,5-bisphosphate carboxylase/oxygenase large subunit (rbcL) gene, partial cds; mitochondrial

TAACCTAACTGCTTCAATTATTGGTAACATCTTCGGATTCAAAGCTGTTAAAGCTCTAAGATTAGAAGATATGAGAATGCCAGTTGCACTACTAAAGACTTACCAAGGTCCTGCTTGTGGTCTAATTGTAGAGCGTGAGAGAATGGACAAGTTTGGTCGTCCTCTACTTGGTGCTACAGTTAAGCCTAAGCTAGGTCTTTCTGGTAAGAACTATGGTCGTGTAGTATTCGAAGGTCTTAAAGGTGGTCTTGATTTCCTTAAGGATGATGAGAACATTAACTCACAACCGTTCATGCGTTACAGAGAGCG

>MN927510.1:1-1391 Phaeocystis globosa isolate CNS00070 ribulose-1,5-bisphosphate carboxylase/oxygenase large subunit (rbcL) gene, partial cds; mitochondrial

TAACCTAACTGCTTCAATTATTGGTAACATCTTCGGATTCAAAGCTGTTAAAGCTCTAAGATTAGAAGATATGAGAATGCCAGTTGCACTACTAAAGACTTACCAAGGTCCTGCTTGTGGTCTAATTGTAGAGCGTGAGAGAATGGACAAGTTTGGTCGTCCTCTACTTGGTGCTACAGTTAAGCCTAAGCTAGGTCTTTCTGGTAAGAACTATGGTCGTGTAGTATTCGAAGGTCTTAAAGGTGGTCTTGATTTCCTTAAGGATGATGAGAACATTAACTCACAACCGTTCATGCGTTACAGAGAGCG

>MN927509.1:1-1391 Phaeocystis globosa isolate CNS00068 ribulose-1,5-bisphosphate carboxylase/oxygenase large subunit (rbcL) gene, partial cds; mitochondrial

TAACCTAACTGCTTCAATTATTGGTAACATCTTCGGATTCAAAGCTGTTAAAGCTCTAAGATTAGAAGATATGAGAATGCCAGTTGCACTACTAAAGACTTACCAAGGTCCTGCTTGTGGTCTAATTGTAGAGCGTGAGAGAATGGACAAGTTTGGTCGTCCTCTACTTGGTGCTACAGTTAAGCCTAAGCTAGGTCTTTCTGGTAAGAACTATGGTCGTGTAGTATTCGAAGGTCTTAAAGGTGGTCTTGATTTCCTTAAGGATGATGAGAACATTAACTCACAACCGTTCATGCGTTACAGAGAGCG

>MN927508.1:1-1391 Phaeocystis globosa isolate CNS00069 ribulose-1,5-bisphosphate carboxylase/oxygenase large subunit (rbcL) gene, partial cds; mitochondrial

TAACCTAACTGCTTCAATTATTGGTAACATCTTCGGATTCAAAGCTGTTAAAGCTCTAAGATTAGAAGATATGAGAATGCCAGTTGCACTACTAAAGACTTACCAAGGTCCTGCTTGTGGTCTAATTGTAGAGCGTGAGAGAATGGACAAGTTTGGTCGTCCTCTACTTGGTGCTACAGTTAAGCCTAAGCTAGGTCTTTCTGGTAAGAACTATGGTCGTGTAGTATTCGAAGGTCTTAAAGGTGGTCTTGATTTCCTTAAGGATGATGAGAACATTAACTCACAACCGTTCATGCGTTACAGAGAGCG

>MT471330.1:10-1415 Phaeocystis globosa strain CNS00076 chloroplast, complete genome

TAACCTAACTGCTTCAATTATTGGTAACATCTTCGGATTCAAAGCTGTTAAAGCTCTAAGATTAGAAGATATGAGAATGCCAGTTGCACTATTAAAGACTTACCAAGGTCCTGCTTGTGGTCTAATCGTAGAGCGTGAGAGAATGGATAAGTTTGGTCGTCCTCTACTTGGTGCTACAGTTAAGCCTAAGCTAGGTCTTTCTGGTAAGAACTACGGTCGTGTAGTATTCGAAGGTCTTAAAGGTGGTCTTGATTTCCTTAAGGATGATGAGAACATTAACTCACAACCGTTCATGCGTTACAGAGAGCG

>MT471329.1:10-1415 Phaeocystis globosa strain CNS00075 chloroplast, complete genome

TAACCTAACTGCTTCAATTATTGGTAACATCTTCGGATTCAAAGCTGTTAAAGCTCTAAGATTAGAAGATATGAGAATGCCAGTTGCACTATTAAAGACTTACCAAGGTCCTGCTTGTGGTCTAATCGTAGAGCGTGAGAGAATGGATAAGTTTGGTCGTCCTCTACTTGGTGCTACAGTTAAGCCTAAGCTAGGTCTTTCTGGTAAGAACTACGGTCGTGTAGTATTCGAAGGTCTTAAAGGTGGTCTTGATTTCCTTAAGGATGATGAGAACATTAACTCACAACCGTTCATGCGTTACAGAGAGCG

>MT471328.1:10-1415 Phaeocystis globosa strain CNS00067 chloroplast, complete genome

TAACCTAACTGCTTCAATTATTGGTAACATCTTCGGATTCAAAGCTGTTAAAGCTCTAAGATTAGAAGATATGAGAATGCCAGTTGCACTATTAAAGACTTACCAAGGTCCTGCTTGTGGTCTAATCGTAGAGCGTGAGAGAATGGATAAGTTTGGTCGTCCTCTACTTGGTGCTACAGTTAAGCCTAAGCTAGGTCTTTCTGGTAAGAACTACGGTCGTGTAGTATTCGAAGGTCTTAAAGGTGGTCTTGATTTCCTTAAGGATGATGAGAACATTAACTCACAACCGTTCATGCGTTACAGAGAGCG

>MT471327.1:10-1415 Phaeocystis globosa strain CNS00066 chloroplast, complete genome

TAACCTAACTGCTTCAATTATTGGTAACATCTTCGGATTCAAAGCTGTTAAAGCTCTAAGATTAGAAGATATGAGAATGCCAGTTGCACTATTAAAGACTTACCAAGGTCCTGCTTGTGGTCTAATCGTAGAGCGTGAGAGAATGGATAAGTTTGGTCGTCCTCTACTTGGTGCTACAGTTAAGCCTAAGCTAGGTCTTTCTGGTAAGAACTACGGTCGTGTAGTATTCGAAGGTCTTAAAGGTGGTCTTGATTTCCTTAAGGATGATGAGAACATTAACTCACAACCGTTCATGCGTTACAGAGAGCG

>MT471326.1:10-1415 Phaeocystis globosa strain CNS00065 chloroplast, complete genome

TAACCTAACTGCTTCAATTATTGGTAACATCTTCGGATTCAAAGCTGTTAAAGCTCTAAGATTAGAAGATATGAGAATGCCAGTTGCACTATTAAAGACTTACCAAGGTCCTGCTTGTGGTCTAATCGTAGAGCGTGAGAGAATGGATAAGTTTGGTCGTCCTCTACTTGGTGCTACAGTTAAGCCTAAGCTAGGTCTTTCTGGTAAGAACTACGGTCGTGTAGTATTCGAAGGTCTTAAAGGTGGTCTTGATTTCCTTAAGGATGATGAGAACATTAACTCACAACCGTTCATGCGTTACAGAGAGCG

>MT471325.1:10-1415 Phaeocystis globosa strain CNS00064 chloroplast, complete genome

TAACCTAACTGCTTCAATTATTGGTAACATCTTCGGATTCAAAGCTGTTAAAGCTCTAAGATTAGAAGATATGAGAATGCCAGTTGCACTATTAAAGACTTACCAAGGTCCTGCTTGTGGTCTAATCGTAGAGCGTGAGAGAATGGATAAGTTTGGTCGTCCTCTACTTGGTGCTACAGTTAAGCCTAAGCTAGGTCTTTCTGGTAAGAACTACGGTCGTGTAGTATTCGAAGGTCTTAAAGGTGGTCTTGATTTCCTTAAGGATGATGAGAACATTAACTCACAACCGTTCATGCGTTACAGAGAGCG

>MT471324.1:10-1415 Phaeocystis globosa strain CNS00063 chloroplast, complete genome

TAACCTAACTGCTTCAATTATTGGTAACATCTTCGGATTCAAAGCTGTTAAAGCTCTAAGATTAGAAGATATGAGAATGCCAGTTGCACTATTAAAGACTTACCAAGGTCCTGCTTGTGGTCTAATCGTAGAGCGTGAGAGAATGGATAAGTTTGGTCGTCCTCTACTTGGTGCTACAGTTAAGCCTAAGCTAGGTCTTTCTGGTAAGAACTACGGTCGTGTAGTATTCGAAGGTCTTAAAGGTGGTCTTGATTTCCTTAAGGATGATGAGAACATTAACTCACAACCGTTCATGCGTTACAGAGAGCG

>MT471323.1:10-1415 Phaeocystis globosa strain CNS00062 chloroplast, complete genome

TAACCTAACTGCTTCAATTATTGGTAACATCTTCGGATTCAAAGCTGTTAAAGCTCTAAGATTAGAAGATATGAGAATGCCAGTTGCACTATTAAAGACTTACCAAGGTCCTGCTTGTGGTCTAATCGTAGAGCGTGAGAGAATGGATAAGTTTGGTCGTCCTCTACTTGGTGCTACAGTTAAGCCTAAGCTAGGTCTTTCTGGTAAGAACTACGGTCGTGTAGTATTCGAAGGTCTTAAAGGTGGTCTTGATTTCCTTAAGGATGATGAGAACATTAACTCACAACCGTTCATGCGTTACAGAGAGCG

>MN927516.1:1-1391 Phaeocystis globosa isolate CNS00078 ribulose-1,5-bisphosphate carboxylase/oxygenase large subunit (rbcL) gene, partial cds; mitochondrial

TAACCTAACTGCTTCAATTATTGGTAACATCTTCGGATTCAAAGCTGTTAAAGCTCTAAGATTAGAAGATATGAGAATGCCAGTTGCACTATTAAAGACTTACCAAGGTCCTGCTTGTGGTCTAATCGTAGAGCGTGAGAGAATGGATAAGTTTGGTCGTCCTCTACTTGGTGCTACAGTTAAGCCTAAGCTAGGTCTTTCTGGTAAGAACTACGGTCGTGTAGTATTCGAAGGTCTTAAAGGTGGTCTTGATTTCCTTAAGGATGATGAGAACATTAACTCACAACCGTTCATGCGTTACAGAGAGCG

>MN927515.1:1-1391 Phaeocystis globosa isolate CNS00077 ribulose-1,5-bisphosphate carboxylase/oxygenase large subunit (rbcL) gene, partial cds; mitochondrial

TAACCTAACTGCTTCAATTATTGGTAACATCTTCGGATTCAAAGCTGTTAAAGCTCTAAGATTAGAAGATATGAGAATGCCAGTTGCACTATTAAAGACTTACCAAGGTCCTGCTTGTGGTCTAATCGTAGAGCGTGAGAGAATGGATAAGTTTGGTCGTCCTCTACTTGGTGCTACAGTTAAGCCTAAGCTAGGTCTTTCTGGTAAGAACTACGGTCGTGTAGTATTCGAAGGTCTTAAAGGTGGTCTTGATTTCCTTAAGGATGATGAGAACATTAACTCACAACCGTTCATGCGTTACAGAGAGCG

>MN927514.1:1-1391 Phaeocystis globosa isolate CNS00076 ribulose-1,5-bisphosphate carboxylase/oxygenase large subunit (rbcL) gene, partial cds; mitochondrial

TAACCTAACTGCTTCAATTATTGGTAACATCTTCGGATTCAAAGCTGTTAAAGCTCTAAGATTAGAAGATATGAGAATGCCAGTTGCACTATTAAAGACTTACCAAGGTCCTGCTTGTGGTCTAATCGTAGAGCGTGAGAGAATGGATAAGTTTGGTCGTCCTCTACTTGGTGCTACAGTTAAGCCTAAGCTAGGTCTTTCTGGTAAGAACTACGGTCGTGTAGTATTCGAAGGTCTTAAAGGTGGTCTTGATTTCCTTAAGGATGATGAGAACATTAACTCACAACCGTTCATGCGTTACAGAGAGCG

>MN927513.1:1-1391 Phaeocystis globosa isolate CNS00075 ribulose-1,5-bisphosphate carboxylase/oxygenase large subunit (rbcL) gene, partial cds; mitochondrial

TAACCTAACTGCTTCAATTATTGGTAACATCTTCGGATTCAAAGCTGTTAAAGCTCTAAGATTAGAAGATATGAGAATGCCAGTTGCACTATTAAAGACTTACCAAGGTCCTGCTTGTGGTCTAATCGTAGAGCGTGAGAGAATGGATAAGTTTGGTCGTCCTCTACTTGGTGCTACAGTTAAGCCTAAGCTAGGTCTTTCTGGTAAGAACTACGGTCGTGTAGTATTCGAAGGTCTTAAAGGTGGTCTTGATTTCCTTAAGGATGATGAGAACATTAACTCACAACCGTTCATGCGTTACAGAGAGCG

>MN927507.1:1-1391 Phaeocystis globosa isolate CNS00067 ribulose-1,5-bisphosphate carboxylase/oxygenase large subunit (rbcL) gene, partial cds; mitochondrial

TAACCTAACTGCTTCAATTATTGGTAACATCTTCGGATTCAAAGCTGTTAAAGCTCTAAGATTAGAAGATATGAGAATGCCAGTTGCACTATTAAAGACTTACCAAGGTCCTGCTTGTGGTCTAATCGTAGAGCGTGAGAGAATGGATAAGTTTGGTCGTCCTCTACTTGGTGCTACAGTTAAGCCTAAGCTAGGTCTTTCTGGTAAGAACTACGGTCGTGTAGTATTCGAAGGTCTTAAAGGTGGTCTTGATTTCCTTAAGGATGATGAGAACATTAACTCACAACCGTTCATGCGTTACAGAGAGCG

>MN927506.1:1-1391 Phaeocystis globosa isolate CNS00066 ribulose-1,5-bisphosphate carboxylase/oxygenase large subunit (rbcL) gene, partial cds; mitochondrial

TAACCTAACTGCTTCAATTATTGGTAACATCTTCGGATTCAAAGCTGTTAAAGCTCTAAGATTAGAAGATATGAGAATGCCAGTTGCACTATTAAAGACTTACCAAGGTCCTGCTTGTGGTCTAATCGTAGAGCGTGAGAGAATGGATAAGTTTGGTCGTCCTCTACTTGGTGCTACAGTTAAGCCTAAGCTAGGTCTTTCTGGTAAGAACTACGGTCGTGTAGTATTCGAAGGTCTTAAAGGTGGTCTTGATTTCCTTAAGGATGATGAGAACATTAACTCACAACCGTTCATGCGTTACAGAGAGCG

>MN927505.1:1-1391 Phaeocystis globosa isolate CNS00065 ribulose-1,5-bisphosphate carboxylase/oxygenase large subunit (rbcL) gene, partial cds; mitochondrial

TAACCTAACTGCTTCAATTATTGGTAACATCTTCGGATTCAAAGCTGTTAAAGCTCTAAGATTAGAAGATATGAGAATGCCAGTTGCACTATTAAAGACTTACCAAGGTCCTGCTTGTGGTCTAATCGTAGAGCGTGAGAGAATGGATAAGTTTGGTCGTCCTCTACTTGGTGCTACAGTTAAGCCTAAGCTAGGTCTTTCTGGTAAGAACTACGGTCGTGTAGTATTCGAAGGTCTTAAAGGTGGTCTTGATTTCCTTAAGGATGATGAGAACATTAACTCACAACCGTTCATGCGTTACAGAGAGCG

>MN927504.1:1-1391 Phaeocystis globosa isolate CNS00064 ribulose-1,5-bisphosphate carboxylase/oxygenase large subunit (rbcL) gene, partial cds; mitochondrial

TAACCTAACTGCTTCAATTATTGGTAACATCTTCGGATTCAAAGCTGTTAAAGCTCTAAGATTAGAAGATATGAGAATGCCAGTTGCACTATTAAAGACTTACCAAGGTCCTGCTTGTGGTCTAATCGTAGAGCGTGAGAGAATGGATAAGTTTGGTCGTCCTCTACTTGGTGCTACAGTTAAGCCTAAGCTAGGTCTTTCTGGTAAGAACTACGGTCGTGTAGTATTCGAAGGTCTTAAAGGTGGTCTTGATTTCCTTAAGGATGATGAGAACATTAACTCACAACCGTTCATGCGTTACAGAGAGCG

>MN927503.1:1-1391 Phaeocystis globosa isolate CNS00063 ribulose-1,5-bisphosphate carboxylase/oxygenase large subunit (rbcL) gene, partial cds; mitochondrial

TAACCTAACTGCTTCAATTATTGGTAACATCTTCGGATTCAAAGCTGTTAAAGCTCTAAGATTAGAAGATATGAGAATGCCAGTTGCACTATTAAAGACTTACCAAGGTCCTGCTTGTGGTCTAATCGTAGAGCGTGAGAGAATGGATAAGTTTGGTCGTCCTCTACTTGGTGCTACAGTTAAGCCTAAGCTAGGTCTTTCTGGTAAGAACTACGGTCGTGTAGTATTCGAAGGTCTTAAAGGTGGTCTTGATTTCCTTAAGGATGATGAGAACATTAACTCACAACCGTTCATGCGTTACAGAGAGCG

>AY955041.1 Micromonas pusilla strain CCMP493 ribulose-1,5-bisphosphate carboxylase/oxygenase large subunit (rbcL) gene, partial cds; chloroplast

TAACCTTTTCACATCAATTGTAGGTAACGTATTCGGTTTCAAGGCACTTCGTGCACTTCGTTTAGAGGACTTACGTATTCCTGTAGCTTACTGTAAGACTTTCCAAGGAGCTCCTCACGGAATCCAGGTTGAGCGTGATAAGTTAAACAAGTACGGACGTTCACTTCTTGGATGTACTATTAAGCCTAAGCTTGGACTTTCTGCTAAGAACTACGGTCGTGCAGTATACGAGTGTCTTCGTGGTGGACTTGACTTCACTAAGGATGACGAGAACGTAAACTCACAGCCATTCATGCGTTGGCGTGACCG

>AY955036.1 Micromonas pusilla strain CCMP488 ribulose-1,5-bisphosphate carboxylase/oxygenase large subunit (rbcL) gene, partial cds; chloroplast

TAACCTTTTCACATCAATTGTAGGTAACGTATTCGGTTTCAAGGCACTTCGTGCACTTCGTTTAGAGGACTTACGTATTCCTGTAGCTTACTGTAAGACTTTCCAAGGAGCTCCTCACGGAATCCAGGTTGAGCGTGATAAGTTAAACAAGTACGGACGTTCACTTCTTGGATGTACTATTAAGCCTAAGCTTGGACTTTCTGCTAAGAACTACGGTCGTGCAGTATACGAGTGTCTTCGTGGTGGACTTGACTTCACTAAGGATGACGAGAACGTAAACTCACAGCCATTCATGCGTTGGCGTGACCG

>AY955034.1 Micromonas pusilla strain CCMP1764 ribulose-1,5-bisphosphate carboxylase/oxygenase large subunit (rbcL) gene, partial cds; chloroplast

TAACCTTTTCACATCAATTGTAGGTAACGTATTCGGTTTCAAGGCACTTCGTGCACTTCGTTTAGAGGACTTACGTATTCCTGTAGCTTACTGTAAGACTTTCCAAGGAGCTCCTCACGGAATCCAGGTTGAGCGTGATAAGTTAAACAAGTACGGACGTTCACTTCTTGGATGTACTATTAAGCCTAAGCTTGGACTTTCTGCTAAGAACTACGGTCGTGCAGTATACGAGTGTCTTCGTGGTGGACTTGACTTCACTAAGGATGACGAGAACGTAAACTCACAGCCATTCATGCGTTGGCGTGACCG

>AY955040.1 Micromonas pusilla strain CCMP492 ribulose-1,5-bisphosphate carboxylase/oxygenase large subunit (rbcL) gene, partial cds; chloroplast

TAACCTTTTCACATCAATTGTAGGTAACGTATTCGGTTTCAAGGCACTTCGTGCACTTCGTTTAGAGGACTTACGTATTCCTGTAGCTTACTGTAAGACTTTCCAAGGAGCTCCTCACGGAATCCAGGTTGAGCGTGATAAGTTAAACAAGTACGGACGTTCACTTCTTGGATGTACTATTAAGCCTAAGCTTGGACTTTCTGCTAAGAACTACGGTCGTGCAGTATACGAGTGTCTTCGTGGTGGACTTGACTTCACTAAGGATGACGAGAACGTAAACTCACAGCCATTCATGCGTTGGCGTGACCG

>AY955037.1 Micromonas pusilla strain CCMP489 ribulose-1,5-bisphosphate carboxylase/oxygenase large subunit (rbcL) gene, partial cds; chloroplast

TAACCTTTTCACATCAATTGTAGGTAACGTATTCGGTTTCAAGGCACTTCGTGCACTTCGTTTAGAGGACTTACGTATTCCTGTAGCTTACTGTAAGACTTTCCAAGGAGCTCCTCACGGAATCCAGGTTGAGCGTGATAAGTTAAACAAGTACGGACGTTCACTTCTTGGATGTACTATTAAGCCTAAGCTTGGACTTTCTGCTAAGAACTACGGTCGTGCAGTATACGAGTGTCTTCGTGGTGGACTTGACTTCACTAAGGATGACGAGAACGTAAACTCACAGCCATTCATGCGTTGGCGTGACCG

>AY955033.1 Micromonas pusilla strain CCMP1723 ribulose-1,5-bisphosphate carboxylase/oxygenase large subunit (rbcL) gene, partial cds; chloroplast

TAACCTTTTCACATCAATTGTAGGTAACGTATTCGGTTTCAAGGCACTTCGTGCACTTCGTTTAGAGGACTTACGTATTCCTGTAGCTTACTGTAAGACTTTCCAAGGAGCTCCTCACGGAATTCAGGTTGAGCGTGATAAGTTAAACAAGTACGGACGTTCACTTCTTGGATGTACTATTAAGCCTAAGCTTGGACTTTCTGCTAAGAACTACGGTCGTGCAGTATACGAGTGTCTTCGTGGTGGACTTGACTTCACTAAGGATGACGAGAACGTAAACTCACAGCCATTCATGCGTTGGCGTGACCG

>AY955046.1 Micromonas pusilla strain NEPCC29 ribulose-1,5-bisphosphate carboxylase/oxygenase large subunit (rbcL) gene, partial cds; chloroplast

TAACCTTTTCACTTCAATTGTAGGTAACGTATTCGGTTTCAAGGCACTTCGTGCACTCCGTTTAGAGGATTTACGTATTCCTGTAGCTTACTGTAAGACTTTCCAAGGAGCTCCTCACGGAATCCAGGTTGAGCGTGATAAGTTAAACAAGTACGGACGTGGACTTCTTGGATGTACTATTAAGCCTAAGCTTGGACTTTCTGCTAAGAACTACGGTCGTGCAGTATACGAGTGTCTCCGTGGTGGACTTGACTTCACTAAGGATGATGAGAACGTAAACTCACAGCCATTCATGCGTTGGCGTGACCG

>AY955044.1 Micromonas pusilla strain CS222 ribulose-1,5-bisphosphate carboxylase/oxygenase large subunit (rbcL) gene, partial cds; chloroplast

TAACCTTTTCACTTCAATTGTAGGTAACGTATTCGGTTTCAAGGCACTTCGTGCACTCCGTTTAGAGGATTTACGTATTCCTGTAGCTTACTGTAAGACTTTCCAAGGAGCTCCTCACGGAATCCAGGTTGAGCGTGATAAGTTAAACAAGTACGGACGTGGACTTCTTGGATGTACTATTAAGCCTAAGCTTGGACTTTCTGCTAAGAACTACGGTCGTGCAGTATACGAGTGTCTCCGTGGTGGACTTGACTTCACTAAGGATGATGAGAACGTAAACTCACAGCCATTCATGCGTTGGCGTGACCG

>AY955030.1 Micromonas pusilla strain CCMP1195 ribulose-1,5-bisphosphate carboxylase/oxygenase large subunit (rbcL) gene, partial cds; chloroplast

TAACCTTTTCACTTCAATTGTAGGTAACGTATTCGGTTTCAAGGCACTTCGTGCACTCCGTTTAGAGGATTTACGTATTCCTGTAGCTTACTGTAAGACTTTCCAAGGAGCTCCTCACGGAATCCAGGTTGAGCGTGATAAGTTAAACAAGTACGGACGTGGACTTCTTGGATGTACTATTAAGCCTAAGCTTGGACTTTCTGCTAAGAACTACGGTCGTGCAGTATACGAGTGTCTCCGTGGTGGACTTGACTTCACTAAGGATGATGAGAACGTAAACTCACAGCCATTCATGCGTTGGCGTGACCG

>AY955032.1 Micromonas pusilla strain CCMP1646 ribulose-1,5-bisphosphate carboxylase/oxygenase large subunit (rbcL) gene, partial cds; chloroplast

TAACCTTTTCACATCTATTGTAGGTAACGTATTTGGTTTCAAAGCACTTCGTGCACTTCGTTTAGAAGATTTACGTATTCCAGTAGCTTACTGTAAGACTTTCCAGGGTGCTCCTCACGGAATCCAGGTTGAGCGTGATAAGTTAAACAAGTACGGACGTGGTCTCCTTGGATGTACTATTAAGCCTAAGCTTGGACTTTCTGCTAAGAACTACGGTCGTGCAGTATACGAGTGTCTTCGTGGTGGTCTTGACTTCACTAAGGATGATGAGAACGTAAACTCTCAGCCATTCATGCGTTGGCGTGACCG

>AY955038.1 Micromonas pusilla strain CCMP490 ribulose-1,5-bisphosphate carboxylase/oxygenase large subunit (rbcL) gene, partial cds; chloroplast

TAACCTTTTCACATCAATTGTAGGTAACGTATTCGGATTCAAGGCACTTCGTGCACTCCGTTTAGAGGACTTACGTATTCCAGTAGCTTACTGTAAGACTTTCCAGGGTGCTCCTCACGGAATCCAGGTTGAGCGTGATAAGTTAAACAAGTACGGACGTGGTCTTCTTGGATGTACTATTAAGCCTAAGCTTGGACTTTCTGCTAAGAACTACGGTCGTGCAGTATACGAGTGTCTTCGTGGTGGTCTTGACTTCACGAAGGATGATGAGAACGTAAACTCTCAGCCATTCATGCGTTGGCGTGACCG

>AY955031.1 Micromonas pusilla strain CCMP1545 ribulose-1,5-bisphosphate carboxylase/oxygenase large subunit (rbcL) gene, partial cds; chloroplast

TAACCTTTTCACATCAATTGTAGGTAACGTATTCGGATTCAAGGCACTTCGTGCACTCCGTTTAGAGGACTTACGTATTCCAGTAGCTTACTGTAAGACTTTCCAGGGTGCTCCTCACGGAATCCAGGTTGAGCGTGATAAGTTAAACAAGTACGGACGTGGTCTTCTTGGATGTACTATTAAGCCTAAGCTTGGACTTTCTGCTAAGAACTACGGTCGTGCAGTATACGAGTGTCTTCGTGGTGGTCTTGACTTCACGAAGGATGATGAGAACGTAAACTCTCAGCCATTCATGCGTTGGCGTGACCG

>AY955035.1 Micromonas pusilla strain CCMP2099 ribulose-1,5-bisphosphate carboxylase/oxygenase large subunit (rbcL) gene, partial cds; chloroplast

GAACCTTTTCACATCTATTGTAGGTAACGTATTTGGTTTTAAAGCACTTCGTGCACTTCGTTTAGAAGATTTACGTATTCCAGTAGCTTACTGTAAGACTTTCCAAGGTGCTCCACACGGAATCCAGGTTGAGCGTGATAAGTTAAACAAGTACGGGCGTGGTCTCCTTGGATGTACTATTAAGCCTAAGCTTGGACTTTCTGCTAAGAACTACGGTCGTGCAGTATACGAGTGTCTTCGTGGTGGTCTTGACTTCACTAAGGATGATGAGAACGTAAACTCTCAGCCATTCATGCGTTGGCGTGACCG

>EU380525.1 Micromonas sp. SYD ribulose-15-bisphosphate carboxylase/oxygenase large subunit (rbcL) gene, partial cds; plastid

TAACCTTTTCACATCAATTGTAGGTAACGTATTCGGATTCAAGGCACTTCGTGCTCTCCGTTTAGAGGACTTACGTATTCCAGTAGCTTACTGTAAGACTTTCCAGGGTGCTCCTCACGGAATCCAGGTTGAACGTGACAAATTAAACAAATATGGTCGTGGTCTTTTAGGTTGTACAATCAAACCTAAATTAGGACTTTCAGCTAAAAACTACGGTCGTGCAGTTTATGAATGTTTACGTGGTGGTCTTGACTTTACTAAAGACGACGAAAACGTAAACTCACAACCATTCATGCGTTGGCGTGACCG

>U30276.1 Micromonas pusilla ribulose-1,5-bisphosphate carboxylase/oxygenase large subunit (rbcL) gene, chloroplast gene encoding chloroplast protein, partial cds

TAACCTTTTCACATCTATTGTAGGTAACGTATTCGGATTCAAGGCACTTCGTCGACTTCGTTTAGAGGATTTACGTATTCCTGTAGCTTACTGTAAGACTTTCGTTGGAGCTCCTCACGGAATCCAGGTTGAGCGTGATAAGTTAAACAAGTACGGACGTGGTCTCCTTGGATGTACTATTAAGCCTAAGCTTGGACTTTCTGCTAAGAACTACGGTCGTGCAGTATACGAGTGTCTTCGTGGTGGTCTTGACTTCACGAAGGATGATGAGAACGTAAACTCTCAGCCATTCATGCGTTGGCGTGACCG

>AB491660.1 Prasinococcus capsulatus chloroplast rbcL gene for ribulose-1,5-bisphosphate carboxylase/oxygenase large subunit, partial cds, strain: MBIC11011

TAACCTATTCACTTCAATTGTAGGTAACGTATTTGGATTCAAGGCTCTACGTGCTCTACGTCTTGAGGATCTACGTATTCCTCCTGCTTACTGTAAGACTTTCGTAGGACCTCCTCACGGAATTCAAGTTGAGCGTGATAAGCTAAACAAGTATGGACGTGGACTTCTAGGATGTACAATTAAGCCTAAACTAGGTCTTTCTGCAAAGAACTATGGTCGTGCTGTATATGAGTGTCTTCGTGGTGGACTTGACTTTACGAAGGATGATGAGAACGTAAACTCACAACCATTCATGCGTTGGCGTGATCG

>U30278.1 Mantoniella squamata ribulose-1,5-bisphosphate carboxylase/oxygenase large subunit (rbcL) gene, chloroplast gene encoding chloroplast protein, partial cds

TAACCTTTTCACATCAATCGTTGGTAACGTATTTGGTTTTAAGGCCCTTCGTGCACTTCGTTTAGAAGATTTACGTATTCCTCCAGCATACTGTAAGACTTTCTTCGGTCCTCCACACGGTATCCAGGTTGAGCGTGACCGTCTAAACAAGTATGGTCGTCCTCTACTAGGTTGTACTATTAAGCCAAAGCTTGGACTTTCGGCTAAGAACTACGGTCGTGCCGTTTACGAATGTCTTCGTGGTGGTCTTGACTTCACTAAGGATGATGAGAACGTAACTTCACAGCCATTCGTGCGCTGGAGAGACCG

>AB491664.1 Prasinoderma coloniale chloroplast rbcL gene for ribulose-1,5-bisphosphate carboxylase/oxygenase large subunit, partial cds, strain: MBIC10720

TAACCTATTTACTTCAATTGTTGGTAACGTATTTGGTTTTAAGGCTTTACGTTCACTTCGTCTAGAAGATTTACGTATTCCTCCAGCTTACGTTAAGACTTTCACTGGTCCTCCTCATGGTATTCAGGTTGAGCGTGATAAGCTAAACAAGTACGGACGTGCTCTACTAGGTTGTACTATTAAGCCTAAGCTAGGTCTTTCTGCTAAGAACTACGGTCGTGCAGTATACGAATGTCTTCGTGGTGGTCTTGATTTCACTAAGGATGATGAGAACGTAAACTCTCAGCCATTCATGCGTTGGCGTGATCG

>HM770958.1 Pseudomuriella engadinensis strain UTEX 57 ribulose-1,5-bisphosphate carboxylase/oxygenase large subunit (rbcL) gene, partial cds; chloroplast

AAACCTATTCACTTCAATTGTAGGTAACGTATTTGGTTTCAAAGCTCTTCGTGCACTTCGTCTAGAAGATCTTCGTATTCCTCCAGCTTACGTTAAAACTTTCCAAGGACCTCCACACGGTATTCAATCTGAACGTGACAAACTAAACAAATACGGACGTGGCTTTTTAGGTTGTACAATTAAGCCTAAACTAGGTTTATCAGCTAAAAACTACGGACGTGCATGTTACGAGTGTTTACGTGGTGGTCTTGACTTCACTAAAGATGACGAAAACGTAAACTCACAACCATTCATGCGTTGGAGAGACCG

>EF587479.1 Chlamydomonas moewusii strain UTEX 97 ribulose-1,5-bisphosphate carboxylase/oxygenase large subunit (rbcL) gene, complete cds; chloroplast

TAACTTATTTACATCTATTGTAGGTAACGTATTTGGTTTCAAAGCTTTACGTGCACTACGTCTAGAAGATTTACGTATCCCTCCAGCATATGTTAAAACATTCTCTGGACCTCCACACGGTATCCAAGTAGAACGTGACAAAATTAACAAATACGGACGTGGTCTTTTAGGTTGTACAATTAAGCCTAAATTAGGTCTTTCAGCTAAAAACTATGGACGTGCTGTTTACGAATGTCTACGTGGTGGACTTGACTTTACTAAGGATGACGAAAACGTAAACTCACAACCATTCATGCGTTGGCGTGACCG

>MF101234.1 Uncultured Chlamydomonas ribulose-1,5-bisphosphate carboxylase/oxygenase large subunit (rbcL) gene, partial cds; chloroplast

TAACATGTTCACATCTATTGTAGGTAACGTATTTGGTTTCAAAGCGTTACGTGCTCTACGTTTAGAAGACCTTCGTATCCCTGTAGCTTACGCTAAAACATTCTCAGGTCCTCCACACGGTATCCAAGTAGAACGTGACAAAATTAACAAATACGGTCGTGGTCTATTAGGTTGTACTATTAAACCAAAACTAGGTCTTTCAGCTAAAAACTACGGTCGTGCTGTTTACGAATGTTTACGTGGTGGTTTAGACTTCACTAAAGATGACGAAAACGTAAACTCACAACCATTCATGCGTTGGCGTGACCG

>AB127988.1 Chlamydomonas parkeae chloroplast rbcL gene for large subunit of ribulose-1,5-bisphosphate carboxylase/oxygenase, partial cds, strain:NIES-440

TAACCTTTTCACTTCAATTGTAGGTAACGTTTTCGGTTTCAAAGCTTTACGTGCTCTTCGTTTAGAAGATTTACGTATTTCTAGTGCTTACTGTAAAACTTTCTTAGGCCCACCTCACGGTATCCAAGTTGAACGTGACAAACTAAACAAATACGGTCGTGGTTTATTAGGTTGTACAATTAAACCTAAATTAGGTCTTTCTGCTAAAAACTACGGTCGTGCTGTGTACGAATGTTTACGTGGTGGTTTAGACTTCACTAAAGATGACGAAAACGTTAACTCTCAACCATTCATGCGTTGGAGAGACCG

>U30275.1 Bathycoccus prasinos ribolose-1,5-bisphosphate carboxylase/oxygenase large subunit (rbcL) gene, chloroplast gene encoding chloroplast protein, partial cds

TAACCTTTTCACATCAATGGTAGGTAACGTATTTGGCTTTAAAGCTCTTCGTGCTCTACGTCTAGAAGACTTACGTATTCCTGCAGCATACTGTAAGACTTTCCAAGGTCCTCCACACGGTATTCAGGTTGAGCGTGACNNNNTAAACAAATATGGTCGTCCTCTACTAGGTTGTACAATTAAAGCAAAAGCTGGACTTTCTGCTAANAACTACGGTCGTGCAGAATACGAATGTCTTCGTGGTGAACTTGACTTCACTAAGGATGACGAAGACGTAAACTCACAGCCATTCATGCGTTGGCGTGACCG

>U30277.1 Mamiella sp. ribulose-1,5-bisphosphate carboxylase/oxygenase large subunit (rbcL) gene, chloroplast gene encoding chloroplast protein, partial cds

AAACCTTTTCACATCAATTGTTGGTAACGTATTTGGTTTCAAAGCNNTTCGTGCACTTCGTTTAGAAGATTTACGTATCCCGCCAGCATACTGTAAGACTTTCGTTGGTGCGCCACACGGAATTCAGGTTGAGCGTGATAAGTTAAACAAGTATGGTCGTGGTCTCCTTGGATGTACTATTAAGCCAAAGCTTGGACTTTCTGCGAAGAACTATGGACGTGCAGTATACGAGTGTCTCCGTGGTGGACTTGACTTCACAAAGGATGATGAGAACGTAAACTCACAGCCATTCATGCGTTGGCGTGACCG

>KX139562.1 Choricystis parasitica isolate TB-2 ribulose-1,5-bisphosphate carboxylase/oxygenase large subunit gene, partial cds; chloroplast

TCACTTTTCA-CTTCAATCGTTGGTAACGTATTCGGTTTCAAAGCTTTACGTGCTTTACGTTTAGAAGATTTACGTATTCCACAAGCTTACGCTAAAACTTTCCAAGGTGCTCCGCACGGTATTCAAGTTGAACGTGACAAACTAAACAAATATGGTCGTGGTTTATTAGGTTGTACAATTAAACCAAAATTAGGTTTATCAGCTAAAAACTATGGTCGTGCAGTATACGAATGTTTACGCGGTGGTCTTGACTTCACGAAAGATGATGAAAACGTTACTTCACAACCATTCATGCGTTGGAGAGACCG

>KX139563.1 Choricystis parasitica isolate HB-1 ribulose-1,5-bisphosphate carboxylase/oxygenase large subunit gene, partial cds; chloroplast

TCACTTTTCA-CTTCAATCGTTGGTAACGTATTCGGTTTCAAAGCTTTACGTGCTTTACGTTTAGAAGATTTACGTATTCCACAAGCTTACGCTAAAACTTTCCAAGGTGCTCCGCACGGTATTCAAGTTGAACGTGACAAACTAAACAAATATGGTCGTGGTTTATTAGGTTGTACAATTAAACCAAAATTAGGTTTATCAGCTAAAAACTATGGTCGTGCAGTATACGAATGTTTACGCGGTGGTCTTGACTTCACGAAAGATGATGAAAACGTTACTTCACAACCATTCATGCGTTGGAGAGACCG

>KX139561.1 Choricystis parasitica isolate TB-1 ribulose-1,5-bisphosphate carboxylase/oxygenase large subunit gene, partial cds; chloroplast

TCACTTTTCA-CTTCAATCGTTGGTAACGTATTCGGTTTCAAAGCTTTACGTGCTTTACGTTTAGAAGATTTACGTATTCCACAAGCTTACGCTAAAACTTTCCAAGGTGCTCCGCACGGTATTCAAGTTGAACGTGACAAACTAAACAAATATGGTCGTGGTTTATTAGGTTGTACAATTAAACCAAAATTAGGTTTATCAGCTAAAAACTATGGTCGTGCAGTATACGAATGTTTACGCGGTGGTCTTGACTTCACGAAAGATGATGAAAACGTTACTTCACAACCATTCATGCGTTGGAGAGACCG

>KX139564.1 Choricystis parasitica isolate HB-2 ribulose-1,5-bisphosphate carboxylase/oxygenase large subunit gene, partial cds; chloroplast

TCACTTTTCA-CTTCAATCGTTGGTAACGTATTCGGTTTCAAAGCTTTACGTGCTTTACGTTTAGAAGATTTACGTATTCCACAAGCTTACGCTAAAACTTTCCAAGGTGCTCCGCACGGTATTCAAGTTGAACGTGACAAACTAAACAAATATGGTCGTGGTTTATTAGGTTGTACAATTAAACCAAAATTAGGTTTATCAGCTAAAAACTATGGTCGTGCAGTATACGAATGTTTACGCGGTGGTCTTGACTTCACGAAAGATGATGAAAACGTTACTTCACAACCATTCATGCGTTGGAGAGACCG

>EF455924.1:559-1928 Tribonema minus strain SAG 2168 psbA-rbcL intergenic spacer, partial sequence; ribulose-1,5-bisphosphate carboxylase/oxygenase large subunit (rbcL) gene, complete cds; rbcL-rbcS intergenic spacer, complete sequence; and ribulose-1,5-bisphosphate carboxylase/oxygenase small subunit (rbcS) gene, partial cds; chloroplast

AAACTTAACGGCATCTATTATTGGTAACGTTTTCGGCTTCAAAGCTGTTAAAGCATTACGTTTAGAAGATATGCGAATTCCTTTTGCATACTTAAAAACTTTCCAAGGTCCTGCTACAGGTCTAGTTGTAGAAAGAGAAAGAATGGACAAATTTGGTCGTCCATTATTAGGTGCAACTGTAAAACCAAAACTAGGTCTTTCAGGTAAAAACTATGGTCGTGTAGTATATGAAGGTCTTCGTGGTGGTCTTGATTTCTTAAAAGATGATGAAAACATCAACTCTCAACCATTCATGCGTTGGAGAGAGCG

>EF455928.1:688-2057 Tribonema regulare strain SAG 2164 PsbA (psbA) gene, partial cds; psbA-rbcL intergenic spacer, complete sequence; ribulose-1,5-bisphosphate carboxylase/oxygenase large subunit (rbcL) gene, complete cds; rbcL-rbcS intergenic spacer, complete sequence; and ribulose-1,5-bisphosphate carboxylase/oxygenase small subunit (rbcS) gene, partial cds; chloroplast

AAACTTAACGGCATCTATTATTGGTAACGTTTTCGGCTTCAAAGCTGTTAAAGCATTACGTTTAGAAGATATGCGAATTCCTTTTGCATACTTAAAAACTTTCCAAGGTCCTGCTACAGGTCTAATTGTAGAAAGAGAAAGAATGGACAAATTTGGTCGTCCATTATTAGGTGCAACTGTAAAACCAAAACTAGGTCTTTCAGGTAAAAACTATGGTCGTGTAGTATATGAAGGTCTTCGTGGTGGACTTGATTTCTTAAAAGATGATGAAAACATCAACTCTCAACCATTCATGCGTTGGAGAGAGCG

>MW176117.1:689-2058 Tribonema sp. strain SAG 2603 photosystem II protein D1 (psbA) gene, partial cds; ribulose-1,5-bisphosphate carboxylase/oxygenase large subunit (rbcL) gene, complete cds; and ribulose-1,5-bisphosphate carboxylase/oxygenase small subunit (rbcS) gene, partial cds; plastid

AAACTTAACGGCATCTATTATTGGTAACGTTTTCGGCTTCAAAGCTGTTAAAGCATTACGTTTAGAAGATATGCGAATTCCTTTTGCATACTTAAAAACTTTCCAAGGTCCTGCTACAGGTCTAATTGTAGAAAGAGAAAGAATGGACAAATTTGGTCGTCCATTATTAGGTGCAACTGTAAAACCAAAACTAGGTCTTTCAGGTAAAAACTATGGTCGTGTAGTATATGAAGGTCTTCGTGGTGGTCTTGATTTCTTAAAAGATGATGAAAACATCAACTCTCAACCATTCATGCGTTGGAGAGAGCG

>MW176118.1:631-2000 Tribonema sp. strain SAG 2604 photosystem II protein D1 (psbA) gene, partial cds; ribulose-1,5-bisphosphate carboxylase/oxygenase large subunit (rbcL) gene, complete cds; and ribulose-1,5-bisphosphate carboxylase/oxygenase small subunit (rbcS) gene, partial cds; plastid

AAACTTAACGGCATCTATTATTGGTAACGTTTTCGGCTTCAAAGCTGTTAAAGCATTACGTTTAGAAGATATGCGAATTCCTTTTGCATACTTAAAAACTTTCCAAGGTCCTGCTACAGGTCTAATTGTAGAAAGAGAAAGAATGGACAAATTTGGTCGTCCATTATTAGGTGCAACTGTAAAACCAAAACTAGGTCTTTCAGGTAAAAACTATGGTCGTGTAGTATATGAAGGTCTTCGTGGTGGTCTTGATTTCTTAAAAGATGATGAAAACATCAACTCTCAACCATTCATGCGTTGGAGAGAGCG

>EF460489.1:700-2069 Tribonema sp. SAG 2165 psbA gene, partial sequence; psbA-rbcL intergenic spacer, complete sequence; ribulose-1,5-bisphosphate carboxylase/oxygenase large subunit (rbcL) gene, complete cds; rbcL-rbcS intergenic spacer, complete sequence; and rbcS gene, partial sequence; chloroplast

AAACTTAACGGCATCTATTATTGGTAACGTTTTCGGCTTCAAAGCTGTTAAAGCATTACGTTTAGAAGATATGCGAATTCCTTTTGCATACTTAAAAACTTTCCAAGGTCCTGCTACAGGTCTAGTTGTAGAAAGAGAAAGAATGGACAAATTTGGTCGTCCATTATTAGGTGCAACTGTAAAACCAAAACTAGGTCTTTCAGGTAAAAACTATGGTCGTGTAGTATATGAAGGTCTTCGTGGTGGTCTTGATTTCTTAAAAGATGATGAAAACATCAACTCTCAACCATTCATGCGTTGGAGAGAGCG

>EF460490.1:448-1817 Tribonema sp. SAG 2174 psbA-rbcL intergenic spacer, partial sequence; ribulose-1,5-bisphosphate carboxylase/oxygenase large subunit (rbcL) gene, complete cds; rbcL-rbcS intergenic spacer, complete sequence; and ribulose-1,5-bisphosphate carboxylase/oxygenase small subunit (rbcS) gene, partial cds; chloroplast

GAACTTAACGGCATCTATTATTGGTAACGTTTTCGGCTTCAAAGCTGTTAAAGCATTACGTTTAGAAGATATGCGAATTCCTTTTGCATACTTAAAAACTTTCCAAGGTCCTGCTACAGGTCTAATTGTAGAAAGAGAAAGAATGGACAAATTTGGTCGTCCATTATTAGGTGCAACTGTAAAACCAAAACTAGGTCTTTCAGGTAAAAACTATGGTCGTGTAGTATATGAAGGTCTTCGTGGTGGTCTTGATTTCTTAAAAGATGATGAAAACATCAACTCTCAACCATTCATGCGTTGGAGAGAGCG

>EF455944.1:690-2059 Tribonema sp. SAG 2175 PsbA (psbA) gene, partial cds; psbA-rbcL intergenic spacer, complete sequence; ribulose-1,5-bisphosphate carboxylase/oxygenase large subunit (rbcL) gene, complete cds; rbcL-rbcS intergenic spacer, complete sequence; and ribulose-1,5-bisphosphate carboxylase/oxygenase small subunit (rbcS) gene, partial cds; chloroplast

GAACTTAACGGCATCTATTATTGGTAACGTTTTCGGCTTCAAAGCTGTTAAAGCATTACGTTTAGAAGATATGCGAATTCCTTTTGCATACTTAAAAACTTTCCAAGGTCCTGCTACAGGTCTAATTGTAGAAAGAGAAAGAATGGACAAATTTGGTCGTCCATTATTAGGTGCAACTGTAAAACCAAAACTAGGTCTTTCAGGTAAAAACTATGGTCGTGTAGTATATGAAGGTCTTCGTGGTGGTCTTGATTTCTTAAAAGATGATGAAAACATCAACTCTCAACCATTCATGCGTTGGAGAGAGCG

>EF455950.1:447-1816 Tribonema sp. SAG 2176 psbA-rbcL intergenic spacer, partial sequence; ribulose-1,5-bisphosphate carboxylase/oxygenase large subunit (rbcL) gene, complete cds; rbcL-rbcS intergenic spacer, complete sequence; and ribulose-1,5-bisphosphate carboxylase/oxygenase small subunit (rbcS) gene, partial cds; chloroplast

GAACTTAACGGCATCTATTATTGGTAACGTTTTCGGCTTCAAAGCTGTTAAAGCATTACGTTTAGAAGATATGCGAATTCCTTTTGCATACTTAAAAACTTTCCAAGGTCCTGCTACAGGTCTAATTGTAGAAAGAGAAAGAATGGACAAATTTGGTCGTCCATTATTAGGTGCAACTGTAAAACCAAAACTAGGTCTTTCAGGTAAAAACTATGGTCGTGTAGTATATGAAGGTCTTCGAGGTGGTCTTGATTTCTTAAAAGATGATGAAAACATCAACTCTCAACCATTCATGCGTTGGAGAGAGCG

>EF460493.1:451-1820 Tribonema viride strain SAG 2167 psbA-rbcL intergenic spacer, partial sequence; ribulose-1,5-bisphosphate carboxylase/oxygenase large subunit (rbcL) gene, complete cds; rbcL-rbcS intergenic spacer, complete sequence; and ribulose-1,5-bisphosphate carboxylase/oxygenase small subunit (rbcS) gene, partial cds; chloroplast

AAACTTAACGGCATCTATTATTGGTAACGTTTTCGGCTTCAAAGCTGTTAAAGCATTACGTTTAGAAGATATGCGAATTCCTTTTGCATACTTAAAGACTTTCCAAGGTCCTGCTACAGGTCTAATTGTAGAAAGAGAAAGAATGGACAAATTTGGTCGTCCATTATTAGGTGCAACTGTAAAACCAAAACTAGGTCTTTCAGGTAAAAACTATGGTCGTGTAGTATATGAAGGTCTTCGTGGTGGTCTTGATTTCTTAAAAGATGATGAAAACATCAACTCTCAACCATTCATGCGTTGGAGAGAGCG

>EF455966.1:621-1990 Tribonema viride strain SAG 23.94 psbA-rbcL intergenic spacer, complete sequence; ribulose-1,5-bisphosphate carboxylase/oxygenase large subunit (rbcL) gene, complete cds; rbcL-rbcS intergenic spacer, complete sequence; and ribulose-1,5-bisphosphate carboxylase/oxygenase small subunit (rbcS) gene, partial cds; chloroplast

AAACTTAACGGCATCTATTATTGGTAACGTTTTCGGCTTCAAAGCTGTTAAAGCATTACGTTTAGAAGATATGCGAATTCCTTTTGCATACTTAAAAACTTTCCAAGGTCCTGCTACAGGTCTAATTGTAGAAAGAGAAAGAATGGACAAATTTGGTCGTCCATTATTAGGTGCAACTGTAAAACCAAAACTAGGTCTTTCAGGTAAAAACTATGGTCGTGTAGTATATGAAGGTCTTCGTGGTGGTCTTGATTTCTTAAAAGATGATGAAAACATCAACTCTCAACCATTCATGCGTTGGAGAGAGCG

>MK482703.1:721-2090 Ophiocytium capitatum strain ACOI 1329 ribulose-1,5-bisphosphate carboxylase/oxygenase large subunit (rbcL) gene, complete cds; and ribulose-1,5-bisphosphate carboxylase/oxygenase small subunit (rbcS) gene, partial cds; plastid

TAACTTAACAGCATCTATTATTGGTAACGTTTTCGGCTTCAAAGCTGTTAAAGCATTACGTTTAGAAGATATGCGAATTCCTTTTGCATACTTAAAAACTTTCCAAGGTCCTGCTACAGGTTTAATTGTAGAAAGAGAAAGAATGGACAAATTTGGTCGTCCATTATTAGGTGCTACTGTAAAACCAAAATTAGGTCTTTCAGGTAAAAACTACGGTCGTGTAGTATATGAAGGTCTACGTGGTGGTCTTGACTTCTTAAAAGATGACGAAAACATCAACTCTCAACCATTCATGCGTTGGAGAGAGCG

>MK482701.1:509-1878 Ophiocytium capitatum strain PACC 8735 ribulose-1,5-bisphosphate carboxylase/oxygenase large subunit (rbcL) gene, complete cds; and ribulose-1,5-bisphosphate carboxylase/oxygenase small subunit (rbcS) gene, partial cds; plastid

GAACTTAACAGCATCTATTATTGGTAACGTTTTCGGTTTCAAAGCTGTTAAAGCATTACGTTTAGAAGATATGCGAATTCCTTTTGCATACTTAAAAACTTTCCAAGGTCCTGCGACAGGTTTAATTGTAGAAAGAGAAAGAATGGACAAATTTGGTCGTCCATTATTAGGTGCTACTGTAAAACCAAAATTAGGTCTTTCAGGTAAAAACTACGGTCGTGTAGTATATGAAGGTCTTCGTGGTGGTCTTGACTTCTTAAAAGATGATGAAAACATCAACTCTCAACCATTCATGCGTTGGAGAGAGCG

>EF455959.1:670-2039 Ophiocytium capitatum strain SAG 7.83 psbA-rbcL intergenic spacer, partial sequence; ribulose-1,5-bisphosphate carboxylase/oxygenase large subunit (rbcL) gene, complete cds; rbcL-rbcS intergenic spacer, complete sequence; and ribulose-1,5-bisphosphate carboxylase/oxygenase small subunit (rbcS) gene, partial cds; chloroplast

GAACTTAACAGCATCTATTATTGGTAACGTTTTCGGTTTCAAAGCTGTTAAAGCATTACGTTTAGAAGATATGCGAATTCCTTTTGCATACTTAAAAACTTTCCAAGGTCCTGCGACAGGTTTAATTGTAGAAAGAGAAAGAATGGACAAATTTGGTCGTCCATTATTAGGTGCTACTGTAAAACCAAAATTAGGTCTTTCAGGTAAAAACTACGGTCGTGTAGTATATGAAGGTCTTCGTGGTGGTCTTGACTTCTTAAAAGATGATGAAAACATCAACTCTCAACCATTCATGCGTTGGAGAGAGCG

>MK482696.1:664-2033 Ophiocytium mucronatum strain ACOI 1514 ribulose-1,5-bisphosphate carboxylase/oxygenase large subunit (rbcL) gene, complete cds; and ribulose-1,5-bisphosphate carboxylase/oxygenase small subunit (rbcS) gene, partial cds; plastid

AAATTTAACAGCATCTATTATTGGTAACGTTTTCGGCTTCAAAGCTGTTAAAGCATTACGTTTAGAAGATATGCGAATTCCTTTTGCATACTTAAAAACTTTCCAAGGTCCAGCTACAGGTTTAGTTGTAGAAAGAGAAAGAATGGACAAATTCGGTCGTCCATTCTTAGGTGCTACTGTAAAACCAAAATTAGGTCTTTCTGGTAAAAACTACGGTCGTGTAGTATATGAAGGTCTTCGTGGTGGTCTTGATTTCTTAAAAGATGATGAAAACATTAACTCTCAACCATTCATGCGTTGGAGAGAACG

>MK482699.1:1384-2753 Ophiocytium parvulum strain ACOI 2954 ribulose-1,5-bisphosphate carboxylase/oxygenase large subunit (rbcL) gene, complete cds; and ribulose-1,5-bisphosphate carboxylase/oxygenase small subunit (rbcS) gene, partial cds; plastid

AAACTTAACAGCATCTATCATTGGTAACGTTTTCGGCTTCAAAGCTGTTAAAGCATTACGTTTAGAAGATATGCGAATTCCTTTTGCATACTTAAAAACTTTCCAAGGTCCAGCTACTGGTTTAATTGTAGAAAGAGAAAGAATGGACAAATTTGGTCGTCCATTATTAGGTGCTACTGTAAAACCAAAATTAGGTCTTTCAGGTAAAAACTACGGTCGTGTAGTATATGAAGGTCTTCGTGGTGGTCTTGATTTCTTAAAAGATGATGAAAACATTAACTCTCAACCATTCATGCGTTGGAGAGAGCG

>MK482698.1:1384-2753 Ophiocytium parvulum strain ACOI 1517 ribulose-1,5-bisphosphate carboxylase/oxygenase large subunit (rbcL) gene, complete cds; and ribulose-1,5-bisphosphate carboxylase/oxygenase small subunit (rbcS) gene, partial cds; plastid

AAACTTAACAGCATCTATCATTGGTAACGTTTTCGGCTTCAAAGCTGTTAAAGCATTACGTTTAGAAGATATGCGAATTCCTTTTGCATACTTAAAAACTTTCCAAGGTCCAGCTACTGGTTTAATTGTAGAAAGAGAAAGAATGGACAAATTTGGTCGTCCATTATTAGGTGCTACTGTAAAACCAAAATTAGGTCTTTCAGGTAAAAACTACGGTCGTGTAGTATATGAAGGTCTTCGTGGTGGTCTTGATTTCTTAAAAGATGATGAAAACATTAACTCTCAACCATTCATGCGTTGGAGAGAGCG

>AJ579570.1:36-1389 Botrydiopsis intercedens chloroplast partial rbcL gene for ribulose-1,5-bisphosphate carboxylase/oxygenase

AAACTTAACAGCATCAATCATTGGTAACGTTTTCGGTTTCAAAGCTGTTAAAGCATTACGTTTAGAAGATATGCGAATTCCTTTTGCATACTTAAAAACTTTCCAAGGTCCTGCTACAGGTTTAATCGTAGAAAGAGAAAGAATGGACAAATTTGGCCGTCCTTTACTAGGTGCTACTGTAAAACCAAAATTAGGTCTTTCAGGTAAAAACTACGGTCGTGTAGTATATGAAGGTCTTCGCGGTGGTCTTGACTTCTTAAAAGATGATGAAAACATTAACTCTCAACCATTCATGCGTTGGAGAGAACG

>AJ579564.1:34-1391 Botryochloris sp. 'Southern Victoria Land' chloroplast partial rbcL gene for ribulose-1,5-bisphosphate carboxylase/oxygenase

AAACTTAACAGCATCAATTATCGGAAACGTTTTTGGTTTCAAAGCTGTTAAAGCTTTACGTTTAGAAGATATGCGAATTCCTTTTGCATACTTAAAAACTTTCCAAGGTCCTGCTACAGGTTTAATTGTAGAAAGAGAAAGAATGGACAAATTTGGCCGTCCTTTACTAGGTGCTACTGTAAAACCGAAATTAGGTCTTTCAGGTAAAAACTACGGTCGTGTAGTATACGAAGGTCTTCGCGGTGGTCTTGACTTCTTAAAAGATGATGAAAACATTAACTCTCAACCATTCATGCGTTGGAGAGAACG

>EF460492.1:621-1990 Bumilleria klebsiana strain SAG 2158 PsbA (psbA) gene, partial cds; psbA-rbcL intergenic spacer, complete sequence; ribulose-1,5-bisphosphate carboxylase/oxygenase large subunit (rbcL) gene, complete cds; rbcL-rbcS intergenic spacer, complete sequence; and ribulose-1,5-bisphosphate carboxylase/oxygenase small subunit (rbcS) gene, partial cds; chloroplast

AAACTTAACTGCATCTATTATTGGTAACGTTTTCGGCTTCAAAGCTGTTAAAGCATTACGTTTAGAAGATATGCGAATTCCTTTTGCATACTTAAAAACTTTCCAAGGTCCTGCTACAGGTCTAATCGTAGAAAGAGAAAGAATGGACAAATTCGGTCGTCCATTATTAGGTGCAACTGTAAAACCAAAATTAGGTCTTTCAGGTAAAAACTACGGTCGTGTAGTATATGAAGGTCTTCGTGGTGGTCTTGACTTCTTAAAAGATGATGAAAACATTAACTCTCAACCATTCATGCGTTGGAGAGAGCG

>EF455982.1:664-2033 Bumilleria sicula strain SAG 808-1 PsbA (psbA) gene, partial cds; psbA-rbcL intergenic spacer, complete sequence; ribulose-1,5-bisphosphate carboxylase/oxygenase large subunit (rbcL) gene, complete cds; rbcL-rbcS intergenic spacer, complete sequence; and ribulose-1,5-bisphosphate carboxylase/oxygenase small subunit (rbcS) gene, partial cds; chloroplast

AAACTTAACTGCATCTATTATTGGTAACGTTTTCGGCTTCAAAGCTGTTAAAGCATTACGTTTAGAAGATATGCGAATTCCTTTTGCATACTTAAAAACTTTCCAAGGTCCTGCTACAGGTCTAATCGTAGAAAGAGAAAGAATGGACAAATTCGGTCGTCCATTATTAGGTGCAACTGTAAAACCAAAATTAGGTCTTTCAGGTAAAAACTACGGTCGTGTAGTATATGAAGGTCTTCGTGGTGGTCTTGACTTCTTAAAAGATGATGAAAACATTAACTCTCAACCATTCATGCGTTGGAGAGAACG

>AJ874707.1:28-1371 Bumilleria sicula plastid partial rbcL gene for ribulose bisphosphate carboxylase large chain

AAACTTAACTGCATCTATTATTGGTAACGTTTTCGGCTTCAAAGCTGTTAAAGCATTACGTTTAGAAGATATGCGAATTCCTTTTGCATACTTAAAAACTTTCCAAGGTCCTGCTACAGGTCTAATCGTAGAAAGAGAAAGAATGGACAAATTCGGTCGTCCATTATTAGGTGCAACTGTAAAACCAAAATTAGGTCTTTCAGGTAAAAACTACGGTCGTGTAGTATATGAAGGTCTTCGTGGTGGTCTTGACTTCTTAAAAGATGATGAAAACATTAACTCTCAACCATTCATGCGTTGGAGAGAACG

>EF426792.1:613-1982 Bumilleria sp. SAG 2157 PsbA (psbA) gene, partial cds; psbA-rbcL intergenic spacer, complete sequence; ribulose-1,5-bisphosphate carboxylase/oxygenase large subunit (rbcL) gene, complete cds; rbcL-rbcS intergenic spacer, complete sequence; and ribulose-1,5-bisphosphate carboxylase/oxygenase small subunit (rbcS) gene, partial cds; chloroplast

AAACTTAACTGCATCTATTATTGGTAACGTTTTCGGCTTCAAAGCTGTTAAAGCATTACGTTTAGAAGATATGCGAATTCCTTTTGCATACTTAAAAACTTTCCAAGGTCCTGCTACAGGTCTAATCGTAGAAAGAGAAAGAATGGACAAATTCGGTCGTCCATTATTAGGTGCAACTGTAAAACCAAAATTAGGTCTTTCAGGTAAAAACTACGGTCGTGTAGTATATGAAGGTCTTCGTGGTGGTCTTGACTTCTTAAAAGATGATGAAAACATTAACTCTCAACCATTCATGCGTTGGAGAGAGCG

>EF455927.1:696-2065 Bumilleria sp. SAG 2159 PsbA (psbA) gene, partial cds; psbA-rbcL intergenic spacer, complete sequence; ribulose-1,5-bisphosphate carboxylase/oxygenase large subunit (rbcL) gene, complete cds; rbcL-rbcS intergenic spacer, complete sequence; and ribulose-1,5-bisphosphate carboxylase/oxygenase small subunit (rbcS) gene, partial cds; chloroplast

AAACTTAACTGCATCTATTATTGGTAACGTTTTCGGCTTCAAAGCTGTTAAAGCGTTACGTTTAGAAGATATGCGAATTCCTTTTGCATACTTAAAAACTTTCCAAGGTCCTGCTACAGGTCTAATCGTAGAAAGGGAAAGAATGGACAAATTCGGTCGTCCATTATTAGGTGCAACTGTAAAACCAAAATTAGGTCTTTCAGGTAAAAACTACGGTCGTGTAGTATATGAAGGTCTTCGTGGTGGTCTTGACTTCTTAAAAGATGATGAAAACATTAACTCTCAACCATTCATGCGTTGGAGAGAACG

>EF460494.1:714-2083 Bumilleria sp. SAG 2160 psbA gene, partial sequence; psbA-rbcL intergenic spacer, complete sequence; ribulose-1,5-bisphosphate carboxylase/oxygenase large subunit (rbcL) gene, complete cds; rbcL-rbcS intergenic spacer, complete sequence; and rbcS gene, partial sequence; chloroplast

AAACTTAACTGCATCTATTATTGGTAACGTTTTCGGCTTCAAAGCTGTTAAAGCGTTACGTTTAGAAGATATGCGAATTCCTTTTGCATACTTAAAAACTTTCCAAGGTCCTGCTACAGGTCTAATCGTAGAAAGGGAAAGAATGGACAAATTCGGTCGTCCATTATTAGGTGCAACTGTAAAACCAAAATTAGGTCTTTCAGGTAAAAACTACGGTCGTGTAGTATATGAAGGTCTTCGTGGTGGTCTTGACTTCTTAAAAGATGATGAAAACATTAACTCTCAACCATTCATGCGTTGGAGAGAACG

>EF426793.1:674-2043 Bumilleriopsis cf. filiformis strain SAG 2161 PsbA (psbA) gene, partial cds; psbA-rbcL intergenic spacer, complete sequence; ribulose-1,5-bisphosphate carboxylase/oxygenase large subunit (rbcL) gene, complete cds; rbcL-rbcS intergenic spacer, complete sequence; and ribulose-1,5-bisphosphate carboxylase/oxygenase small subunit (rbcS) gene, partial cds; chloroplast

AAACTTAACTGCATCTATTATTGGTAACGTTTTCGGCTTCAAAGCTGTTAAAGCATTACGTTTAGAAGATATGCGAATTCCTTTTGCATACTTAAAAACTTTCCAAGGTCCTGCTACAGGTCTAATCGTAGAAAGAGAAAGAATGGACAAATTCGGTCGTCCATTATTAGGTGCAACTGTAAAACCAAAATTAGGTCTTTCAGGTAAAAACTACGGTCGTGTAGTATATGAAGGTCTTCGTGGTGGTCTTGACTTCTTAAAAGATGATGAAAACATTAACTCTCAACCATTCATGCGTTGGAGAGAACG

>EF431851.1:660-2029 Bumilleriopsis filiformis strain SAG 809-2 PsbA (psbA) gene, partial cds; psbA-rbcL intergenic spacer, complete sequence; ribulose-1,5-bisphosphate carboxylase/oxygenase large subunit (rbcL) gene, complete cds; rbcL-rbcS intergenic spacer, complete sequence; and ribulose-1,5-bisphosphate carboxylase/oxygenase small subunit (rbcS) gene, partial cds; chloroplast

AAACTTAACTGCATCTATTATTGGTAACGTTTTCGGCTTCAAAGCTGTTAAAGCATTACGTTTAGAAGATATGCGAATTCCTTTTGCATACTTAAAAACTTTCCAAGGTCCTGCTACAGGTCTAATCGTAGAAAGAGAAAGAATGGACAAATTCGGTCGTCCATTATTAGGTGCAACTGTAAAACCAAAATTAGGTCTTTCAGGTAAAAACTACGGTCGTGTAGTATATGAAGGTCTTCGTGGTGGTCTTGACTTCTTAAAAGATGATGAAAACATTAACTCTCAACCATTCATGCGTTGGAGAGAACG

>AJ874703.1:1-1370 Bumilleriopsis filiformis plastid partial rbcL gene for ribulose bisphosphate carboxylase large chain

AAACTTAACTGCATCTATTATTGGTAACGTTTTCGGCTTCAAAGCTGTTAAAGCATTACGTTTAGAAGATATGCGAATTCCTTTTGCATACTTAAAAACTTTCCAAGGTCCTGCTACAGGTCTAATCGTAGAAAGAGAAAGAATGGACAAATTCGGTCGTCCATTATTAGGTGCAACTGTAAAACCAAAATTAGGTCTTTCAGGTAAAAACTACGGTCGTGTAGTATATGAAGGTCTTCGTGGTGGTCTTGACTTCTTAAAAGATGATGAAAACATTAACTCTCAACCATTCATGCGTTGGAGAGAACG

>EF455978.1:630-1999 Pseudobumilleriopsis pyrenoidosa strain SAG 69.90 PsbA (psbA) gene, partial cds; psbA-rbcL intergenic spacer, complete sequence; ribulose-1,5-bisphosphate carboxylase/oxygenase large subunit (rbcL) gene, complete cds; rbcL-rbcS intergenic spacer, complete sequence; and ribulose-1,5-bisphosphate carboxylase/oxygenase small subunit (rbcS) gene, partial cds; chloroplast

AAACTTAACTGCATCTATTATTGGTAACGTTTTCGGCTTCAAAGCTGTTAAAGCCTTACGTTTAGAAGATATGCGAATTCCTTTTGCATACTTAAAAACTTTCCAAGGTCCTGCTACAGGTCTAATCGTAGAAAGAGAAAGAATGGACAAATTCGGTCGTCCATTATTAGGTGCAACTGTAAAACCAAAATTAGGTCTTTCAGGTAAAAACTACGGTCGTGTAGTATATGAAGGTCTTCGTGGTGGTCTTGACTTTTTAAAAGATGATGAAAACATTAACTCTCAACCATTCATGCGTTGGAGAGAACG

>EF431850.1:601-1970 Bumilleriopsis sp. SAG 57.94 PsbA (psbA) gene, partial cds; psbA-rbcL intergenic spacer, complete sequence; ribulose-1,5-bisphosphate carboxylase/oxygenase large subunit (rbcL) gene, complete cds; rbcL-rbcS intergenic spacer, complete sequence; and ribulose-1,5-bisphosphate carboxylase/oxygenase small subunit (rbcS) gene, partial cds; chloroplast

AAACTTAACTGCATCTATTATTGGTAACGTTTTCGGCTTCAAAGCTGTTAAAGCCTTACGTTTAGAAGATATGCGAATTCCTTTTGCATACTTAAAAACTTTCCAAGGTCCTGCTACAGGTCTAATTGTAGAAAGAGAAAGAATGGACAAATTCGGTCGTCCATTATTAGGTGCAACTGTAAAACCAAAATTAGGTCTTTCAGGTAAAAACTACGGTCGTGTAGTATATGAAGGTCTTCGTGGTGGTCTTGACTTTTTAAAAGATGATGAAAACATTAACTCTCAACCATTCATGCGTTGGAGAGAACG

>AJ874705.1:28-1397 Bumilleriopsis sp. SAG 57.94 plastid partial rbcL gene for ribulose bisphosphate carboxylase large chain

AAACTTAACTGCATCTATTATTGGTAACGTTTTCGGCTTCAAAGCTGTTAAAGCCTTACGTTTAGAAGATATGCGAATTCCTTTTGCATACTTAAAAACTTTCCAAGGTCCTGCTACAGGTCTAATTGTAGAAAGAGAAAGAATGGACAAATTCGGTCGTCCATTATTAGGTGCAACTGTAAAACCAAAATTAGGTCTTTCAGGTAAAAACTACGGTCGTGTAGTATATGAAGGTCTTCGTGGTGGTCTTGACTTTTTAAAAGATGATGAAAACATTAACTCTCAACCATTCATGCGTTGGAGAGAACG

>EF455963.1:618-1987 Bumilleriopsis sp. SAG 58.94 PsbA (psbA) gene, partial cds; psbA-rbcL intergenic spacer, complete sequence; ribulose-1,5-bisphosphate carboxylase/oxygenase large subunit (rbcL) gene, complete cds; rbcL-rbcS intergenic spacer, complete sequence; and ribulose-1,5-bisphosphate carboxylase/oxygenase small subunit (rbcS) gene, partial cds; chloroplast

AAACTTAACTGCATCTATTATTGGTAACGTTTTCGGCTTCAAAGCTGTTAAAGCGTTACGTTTAGAAGATATGCGAATTCCTTTTGCATACTTAAAAACTTTCCAAGGTCCTGCTACAGGTCTAATCGTAGAAAGAGAAAGAATGGACAAATTCGGTCGTCCATTATTAGGTGCAACTGTAAAACCAAAATTAGGTCTTTCAGGTAAAAACTACGGTCGTGTAGTATATGAAGGTCTTCGTGGTGGTCTTGACTTCTTAAAAGATGATGAAAACATTAACTCTCAACCATTCATGCGTTGGAGAGAACG

>AJ874704.1:1-1370 Bumilleriopsis sp. SAG 58.94 plastid partial rbcL gene for ribulose bisphosphate carboxylase large chain

AAACTTAACTGCATCTATTATTGGTAACGTTTTCGGCTTCAAAGCTGTTAAAGCGTTACGTTTAGAAGATATGCGAATTCCTTTTGCATACTTAAAAACTTTCCAAGGTCCTGCTACAGGTCTAATCGTAGAAAGAGAAAGAATGGACAAATTCGGTCGTCCATTATTAGGTGCAACTGTAAAACCAAAATTAGGTCTTTCAGGTAAAAACTACGGTCGTGTAGTATATGAAGGTCTTCGTGGTGGTCTTGACTTCTTAAAAGATGATGAAAACATTAACTCTCAACCATTCATGCGTTGGAGAGAACG

>MK792448.1:599-1968 Xanthonema bristolianum strain SAG 2570 PsbA (psbA) gene, partial cds; psbA-rbcL intergenic spacer, complete sequence; ribulose-1,5-bisphosphate carboxylase/oxygenase large subunit (rbcL) gene, complete cds; rbcL-rbcS intergenic spacer, complete sequence; and ribulose-1,5-bisphosphate carboxylase/oxygenase small subunit (rbcS) gene, partial cds; chloroplast

TAACTTAACAGCATCTATTATTGGTAACGTTTTCGGCTTCAAAGCTGTTAAAGCATTACGTTTAGAAGATATGCGAATTCCATTTGCATACTTAAAAACTTTCCAAGGTCCTGCTACAGGTCTAATTGTAGAAAGAGAAAGAATGGACAAATTTGGTCGTCCTTTCTTAGGTGCTACTGTAAAACCAAAATTAGGTCTTTCAGGTAAAAACTACGGTCGTGTAGTATATGAAGGTCTACGTGGTGGTCTTGACTTCTTAAAAGATGATGAAAACATTAACTCTCAACCATTCATGCGTTGGAGAGAGCG

>MW176119.1:551-1920 Xanthonema bristolianum strain SAG 2599 photosystem II protein D1 (psbA) gene, partial cds; ribulose-1,5-bisphosphate carboxylase/oxygenase large subunit (rbcL) gene, complete cds; and ribulose-1,5-bisphosphate carboxylase/oxygenase small subunit (rbcS) gene, partial cds; plastid

TAACTTAACAGCATCTATTATTGGTAACGTTTTCGGCTTCAAAGCTGTTAAAGCATTACGTTTAGAAGATATGCGAATTCCATTTGCATACTTAAAAACTTTCCAAGGTCCTGCTACAGGTCTAATTGTAGAAAGAGAAAGAATGGACAAATTTGGTCGTCCTTTCTTAGGTGCTACTGTAAAACCAAAATTAGGTCTTTCAGGTAAAAACTACGGTCGTGTAGTATATGAAGGTCTACGTGGTGGTCTTGACTTCTTAAAAGATGATGAAAACATTAACTCTCAACCATTCATGCGTTGGAGAGAGCG

>MK804168.1:583-1952 Xanthonema bristolianum strain SAG 2579 Photosystem II protein D1 (psbA) gene, partial cds; ribulose-1,5-bisphosphate carboxylase/oxygenase large subunit (rbcL) gene, complete cds; and ribulose-1,5-bisphosphate carboxylase/oxygenase small subunit (rbcS) gene, partial cds; plastid

TAACTTAACAGCATCTATTATTGGTAACGTTTTCGGCTTCAAAGCTGTTAAAGCATTACGTTTAGAAGATATGCGAATTCCATTTGCATACTTAAAAACTTTCCAAGGTCCTGCTACAGGTCTAATTGTAGAAAGAGAAAGAATGGACAAATTTGGTCGTCCTTTCTTAGGTGCTACTGTAAAACCAAAATTAGGTCTTTCAGGTAAAAACTACGGTCGTGTAGTATATGAAGGTCTACGTGGTGGTCTTGACTTCTTAAAAGATGATGAAAACATTAACTCTCAACCATTCATGCGTTGGAGAGAGCG

>MK804167.1:592-1961 Xanthonema bristolianum strain SAG 2576 Photosystem II protein D1 (psbA) gene, partial cds; ribulose-1,5-bisphosphate carboxylase/oxygenase large subunit (rbcL) gene, complete cds; and ribulose-1,5-bisphosphate carboxylase/oxygenase small subunit (rbcS) gene, partial cds; plastid

TAACTTAACAGCATCTATTATTGGTAACGTTTTCGGCTTCAAAGCTGTTAAAGCATTACGTTTAGAAGATATGCGAATTCCATTTGCATACTTAAAAACTTTCCAAGGTCCTGCTACAGGTCTAATTGTAGAAAGAGAAAGAATGGACAAATTTGGTCGTCCTTTCTTAGGTGCTACTGTAAAACCAAAATTAGGTCTTTCAGGTAAAAACTACGGTCGTGTAGTATATGAAGGTCTACGTGGTGGTCTTGACTTCTTAAAAGATGATGAAAACATTAACTCTCAACCATTCATGCGTTGGAGAGAGCG

>MK804166.1:563-1932 Xanthonema bristolianum strain SAG 2577 Photosystem II protein D1 (psbA) gene, partial cds; ribulose-1,5-bisphosphate carboxylase/oxygenase large subunit (rbcL) gene, complete cds; and ribulose-1,5-bisphosphate carboxylase/oxygenase small subunit (rbcS) gene, partial cds; plastid

TAACTTAACAGCATCTATTATTGGTAACGTTTTCGGCTTCAAAGCTGTTAAAGCATTACGTTTAGAAGATATGCGAATTCCATTTGCATACTTAAAAACTTTCCAAGGTCCTGCTACAGGTCTAATTGTAGAAAGAGAAAGAATGGACAAATTTGGTCGTCCTTTCTTAGGTGCTACTGTAAAACCAAAATTAGGTCTTTCAGGTAAAAACTACGGTCGTGTAGTATATGAAGGTCTACGTGGTGGTCTTGACTTCTTAAAAGATGATGAAAACATTAACTCTCAACCATTCATGCGTTGGAGAGAGCG

>AJ874331.1:28-1397 Xanthonema bristolianum plastid partial rbcL gene for ribulose bisphosphate carboxylase large chain

TAACTTAACAGCATCTATTATTGGTAACGTTTTCGGCTTCAAAGCTGTTAAAGCATTACGTTTAGAAGATATGCGAATTCCATTTGCATACTTAAAAACTTTCCAAGGTCCTGCTACAGGTCTAATTGTAGAAAGAGAAAGAATGGACAAATTTGGTCGTCCTTTCTTAGGTGCTACTGTAAAACCAAAATTAGGTCTTTCAGGTAAAAACTACGGTCGTGTAGTATATGAAGGTCTACGTGGTGGTCTTGACTTCTTAAAAGATGATGAAAACATTAACTCTCAACCATTCATGCGTTGGAGAGAGCG

>EF455955.1:555-1924 Xanthonema bristolianum strain CCALA 516 PsbA (psbA) gene, partial cds; psbA-rbcL intergenic spacer, complete sequence; ribulose-1,5-bisphosphate carboxylase/oxygenase large subunit (rbcL) gene, complete cds; rbcL-rbcS intergenic spacer, complete sequence; and ribulose-1,5-bisphosphate carboxylase/oxygenase small subunit (rbcS) gene, partial cds; chloroplast

TAACTTAACAGCATCTATTATTGGTAACGTTTTCGGCTTCAAAGCTGTTAAAGCATTACGTTTAGAAGATATGCGAATTCCATTTGCATACTTAAAAACTTTCCAAGGTCCTGCTACAGGTCTAATTGTAGAAAGAGAAAGAATGGACAAATTTGGTCGTCCTTTCTTAGGTGCTACTGTAAAACCAAAATTAGGTCTTTCAGGTAAAAACTACGGTCGTGTAGTATATGAAGGTCTACGTGGTGGTCTTGACTTCTTAAAAGATGATGAAAACATTAACTCTCAACCATTCATGCGTTGGAGAGAGCG

>MW176120.1:566-1935 Xanthonema bristolianum strain SAG 2600 photosystem II protein D1 (psbA) gene, partial cds; ribulose-1,5-bisphosphate carboxylase/oxygenase large subunit (rbcL) gene, complete cds; and ribulose-1,5-bisphosphate carboxylase/oxygenase small subunit (rbcS) gene, partial cds; plastid

TAACTTAACAGCATCTATTATTGGTAACGTTTTCGGCTTCAAAGCTGTTAAAGCATTACGTTTAGAAGATATGCGAATTCCATTTGCATACTTAAAAACTTTCCAAGGTCCTGCTACAGGTCTAATTGTAGAAAGAGAAAGAATGGACAAATTTGGTCGTCCTTTCTTAGGTGCTACTGTAAAACCAAAATTAGGTCTTTCAGGTAAAAACTACGGTCGTGTAGTATATGAAGGTCTACGTGGTGGTCTTGACTTCTTAAAAGATGATGAAAACATTAACTCTCAACCATTCATGCGTTGGAGAGAGCG

>AF084612.1:49-1418 Xanthonema debile ribulose 1,5-bisphosphate carboxylase/oxygenase large subunit (rbcL) gene, chloroplast gene encoding chloroplast protein, partial cds

TAACTTAACAGCATCTATCATTGGTAACGTTTTCGGCTTCAAAGCTGTTAAAGCATTACGTTTAGAAGATATGCGAATTCCATTTGCATACTTAAAAACTTTCCAAGGTCCTGCTACAGGCCTAATTGTAGAAAGAGAAAGAATGGACAAATTCGGTCGTCCTTTCTTAGGTGCTACTGTAAAACCAAAATTAGGTCTTTCAGGTAAAAACTACGGTCGTGTAGTATATGAAGGTCTACGTGGTGGTCTTGACTTCTTAAAAGATGATGAAAACATTAACTCTCAACCATTCATGCGTTGGAGAGAACG

>EF455920.1:601-1970 Heterothrix debilis strain UTEX 155 PsbA (psbA) gene, partial cds; psbA-rbcL intergenic spacer, complete sequence; ribulose-1,5-bisphosphate carboxylase/oxygenase large subunit (rbcL) gene, complete cds; rbcL-rbcS intergenic spacer, complete sequence; and ribulose-1,5-bisphosphate carboxylase/oxygenase small subunit (rbcS) gene, partial cds; chloroplast

TAACTTAACAGCATCTATCATTGGTAACGTTTTCGGCTTCAAAGCTGTTAAAGCATTACGTTTAGAAGATATGCGAATTCCATTTGCATACTTAAAAACTTTCCAAGGTCCTGCTACAGGCCTAATTGTAGAAAGAGAAAGAATGGACAAATTCGGTCGTCCTTTCTTAGGTGCTACTGTAAAACCAAAATTAGGTCTTTCAGGTAAAAACTACGGTCGTGTAGTATATGAAGGTCTACGTGGTGGTCTTGACTTCTTAAAAGATGATGAAAACATTAACTCTCAACCATTCATGCGTTGGAGAGAACG

>EF455975.1:594-1963 Xanthonema debile strain SAG 836-1 PsbA (psbA) gene, partial cds; psbA-rbcL intergenic spacer, complete sequence; ribulose-1,5-bisphosphate carboxylase/oxygenase large subunit (rbcL) gene, complete cds; rbcL-rbcS intergenic spacer, complete sequence; and ribulose-1,5-bisphosphate carboxylase/oxygenase small subunit (rbcS) gene, partial cds; chloroplast

GAACTTAACAGCATCTATTATTGGTAACGTTTTCGGCTTCAAAGCTGTTAAAGCATTACGTTTAGAAGATATGCGAATTCCATTTGCATACTTAAAAACTTTCCAAGGTCCTGCTACAGGTCTAATTGTAGAAAGAGAAAGAATGGACAAATTCGGTCGTCCTTTCTTAGGTGCTACTGTAAAACCAAAATTAGGTCTTTCAGGTAAAAACTACGGTCGTGTAGTATATGAAGGTCTACGTGGTGGTCTTGACTTCTTAAAAGATGATGAAAACATTAACTCTCAACCATTCATGCGTTGGAGAGAACG

>EF455937.1:593-1962 Bumilleria exilis strain CCAP 808/2 PsbA (psbA) gene, partial cds; psbA-rbcL intergenic spacer, complete sequence; ribulose-1,5-bisphosphate carboxylase/oxygenase large subunit (rbcL) gene, complete cds; rbcL-rbcS intergenic spacer, complete sequence; and ribulose-1,5-bisphosphate carboxylase/oxygenase small subunit (rbcS) gene, partial cds; chloroplast

TAACTTAACAGCATCTATCATTGGTAACGTTTTCGGCTTCAAAGCTGTTAAAGCATTACGTTTAGAAGATATGCGAATTCCATTTGCATACTTAAAAACTTTCCAAGGTCCTGCTACAGGTCTAATTGTAGAAAGAGAAAGAATGGACAAATTCGGTCGTCCTTTCTTAGGTGCTACTGTAAAACCAAAATTAGGTCTTTCAGGTAAAAACTACGGTCGTGTAGTATATGAAGGTCTACGTGGTGGTCTTGACTTCTTAAAAGATGATGAAAACATTAACTCTCAACCATTCATGCGTTGGAGAGAGCG

>EF455929.1:587-1956 Xanthonema exile strain SAG 2180 PsbA (psbA) gene, partial cds; psbA-rbcL intergenic spacer, complete sequence; ribulose-1,5-bisphosphate carboxylase/oxygenase large subunit (rbcL) gene, complete cds; rbcL-rbcS intergenic spacer, complete sequence; and ribulose-1,5-bisphosphate carboxylase/oxygenase small subunit (rbcS) gene, partial cds; chloroplast

TAACTTAACAGCATCTATTATTGGTAACGTTTTCGGCTTCAAAGCTGTTAAAGCATTACGTTTAGAAGATATGCGAATTCCATTTGCATACTTAAAAACTTTCCAAGGTCCTGCTACAGGTCTAATTGTAGAAAGAGAAAGAATGGACAAATTTGGTCGTCCTTTCTTAGGTGCTACTGTAAAACCAAAATTAGGTCTTTCAGGTAAAAACTACGGTCGTGTAGTATATGAAGGTCTACGTGGTGGTCTTGACTTCTTAAAAGATGATGAAAACATTAACTCTCAACCATTCATGCGTTGGAGAGAGCG

>EF455939.1:518-1887 Xanthonema hormidioides strain CCAP 836/2 PsbA (psbA) gene, partial cds; psbA-rbcL intergenic spacer, complete sequence; ribulose-1,5-bisphosphate carboxylase/oxygenase large subunit (rbcL) gene, complete cds; rbcL-rbcS intergenic spacer, complete sequence; and ribulose-1,5-bisphosphate carboxylase/oxygenase small subunit (rbcS) gene, partial cds; chloroplast

GAACTTAACAGCATCTATTATTGGTAACGTTTTCGGCTTCAAAGCTGTTAAAGCATTACGTTTAGAAGATATGCGAATTCCATTTGCATACTTAAAAACTTTCCAAGGTCCTGCTACAGGTCTAATTGTAGAAAGAGAAAGAATGGACAAATTCGGTCGTCCTTTCTTAGGTGCTACTGTAAAACCAAAATTAGGTCTTTCAGGTAAAAACTACGGTCGTGTAGTATATGAAGGTCTACGTGGTGGTCTTGACTTCTTAAAAGATGATGAAAACATTAACTCTCAACCATTCATGCGTTGGAGAGAACG

>EF455973.1:626-1995 Xanthonema solidum strain SAG 836-4 PsbA (psbA) gene, partial cds; psbA-rbcL intergenic spacer, complete sequence; ribulose-1,5-bisphosphate carboxylase/oxygenase large subunit (rbcL) gene, complete cds; rbcL-rbcS intergenic spacer, complete sequence; and ribulose-1,5-bisphosphate carboxylase/oxygenase small subunit (rbcS) gene, partial cds; chloroplast

TAACTTAACAGCATCTATCATTGGTAACGTTTTCGGCTTCAAAGCTGTTAAAGCTTTACGTTTAGAAGATATGCGAATTCCATATGCATACTTAAAAACTTTCCAAGGTCCTGCTACAGGTCTAATTGTGGAAAGAGAAAGAATGGACAAATTCGGTCGTCCTTTCTTAGGTGCTACTGTAAAACCAAAATTAGGTCTTTCAGGTAAAAACTACGGTCGTGTAGTATATGAAGGTCTACGTGGTGGTCTTGATTTCTTAAAAGATGATGAAAACATTAACTCTCAACCATTCATGCGTTGGAGAGAGCG

>AJ874711.1:28-1397 Xanthonema sp. 421 plastid partial rbcL gene for ribulose bisphosphate carboxylase large chain

TAATTTAACAGCATCTATTATTGGTAACGTTTTCGGCTTCAAAGCTGTTAAAGCATTACGTTTAGAAGATATGCGAATTCCATTTGCATACTTAAAAACTTTCCAAGGTCCTGCTACAGGTCTAATTGTGGAAAGAGAAAGAATGGACAAATTCGGTCGTCCCTTCTTAGGTGCTACTGTAAAACCAAAATTAGGTCTTTCAGGTAAAAACTACGGTCGTGTAGTATATGAAGGTCTACGTGGTGGTCTTGACTTCTTAAAAGATGATGAAAACATTAACTCTCAACCATTCATGCGTTGGAGAGAACG

>EF455940.1:602-1971 Xanthonema sp. CCAP 836/5 PsbA (psbA) gene, partial cds; psbA-rbcL intergenic spacer, complete sequence; ribulose-1,5-bisphosphate carboxylase/oxygenase large subunit (rbcL) gene, complete cds; rbcL-rbcS intergenic spacer, complete sequence; and ribulose-1,5-bisphosphate carboxylase/oxygenase small subunit (rbcS) gene, partial cds; chloroplast

TAACTTAACAGCATCTATTATTGGTAACGTTTTCGGCTTCAAAGCTGTTAAAGCATTACGTTTAGAAGATATGCGAATTCCATTTGCATACTTAAAAACTTTCCAAGGTCCTGCTACAGGTCTAATTGTAGAAAGAGAAAGAATGGACAAATTTGGTCGTCCTTTCTTAGGTGCTACTGTAAAACCAAAATTAGGTCTTTCAGGTAAAAACTACGGTCGTGTAGTATATGAAGGTCTACGTGGTGGTCTTGACTTCTTAAAAGATGATGAAAACATTAACTCTCAACCATTCATGCGTTGGAGAGAGCG

>EF426796.1:567-1936 Xanthonema sp. SAG 2179 PsbA (psbA) gene, partial cds; psbA-rbcL intergenic spacer, complete sequence; ribulose-1,5-bisphosphate carboxylase/oxygenase large subunit (rbcL) gene, complete cds; rbcL-rbcS intergenic spacer, complete sequence; and ribulose-1,5-bisphosphate carboxylase/oxygenase small subunit (rbcS) gene, partial cds; chloroplast

GAACTTAACAGCATCTATTATTGGTAACGTTTTCGGCTTCAAAGCTGTTAAAGCATTACGTTTAGAAGATATGCGAATTCCATTTGCATACTTAAAAACTTTCCAAGGTCCTGCTACAGGTCTAATTGTAGAAAGAGAAAGAATGGACAAATTCGGTCGTCCTTTCTTAGGTGCTACTGTAAAACCAAAATTAGGTCTTTCAGGTAAAAACTACGGTCGTGTAGTATATGAAGGTCTACGTGGTGGTCTTGACTTCTTAAAAGATGATGAAAACATTAACTCTCAACCATTCATGCGTTGGAGAGAACG

>EF455931.1:603-1972 Xanthonema sp. SAG 2184 PsbA (psbA) gene, partial cds; psbA-rbcL intergenic spacer, complete sequence; ribulose-1,5-bisphosphate carboxylase/oxygenase large subunit (rbcL) gene, complete cds; rbcL-rbcS intergenic spacer, complete sequence; and ribulose-1,5-bisphosphate carboxylase/oxygenase small subunit (rbcS) gene, partial cds; chloroplast

TAACTTAACAGCATCTATCATTGGTAACGTTTTCGGCTTCAAAGCTGTTAAAGCATTACGTTTAGAAGATATGCGAATTCCATTTGCATACTTAAAAACTTTCCAAGGTCCTGCTACAGGTCTAATTGTAGAAAGAGAAAGAATGGACAAATTTGGTCGTCCTTTCTTAGGTGCTACTGTAAAACCAAAATTAGGTCTTTCAGGTAAAAACTACGGTCGTGTAGTATATGAAGGTCTACGTGGTGGTCTTGACTTCTTAAAAGATGATGAAAACATTAACTCTCAACCATTCATGCGTTGGAGAGAGCG

>AJ874334.1:28-1397 Xanthonema tribonematoides plastid partial rbcL gene for ribulose bisphosphate carboxylase large chain

AAACTTAACTGCATCTATTATTGGTAACGTTTTCGGCTTCAAAGCTGTTAAAGCATTACGTTTAGAAGATATGCGAATTCCTTTTGCATACTTAAAAACTTTCCAAGGTCCTGCTACAGGTCTAATCGTAGAAAGAGAAAGAATGGACAAATTCGGTCGTCCATTATTAGGTGCAACTGTAAAACCAAAATTAGGTCTTTCAGGTAAAAACTACGGTCGTGTAGTATATGAAGGTCTTCGTGGTGGTCTTGACTTCTTAAAAGATGATGAAAACATTAACTCTCAACCATTCATGCGTTGGAGAGAGCG
